# Getting Started with Machine Learning for Experimental Biochemists and Other Molecular Scientists

**DOI:** 10.1101/2024.10.23.619819

**Authors:** Matthew J. K. Vince, Kristin A. Hughes, Anastasiya Buzuk, Deborah L. Perlstein, Lauren A. Viarengo-Baker, Adrian Whitty

**Affiliations:** Department of Chemistry, Boston University, Boston, MA 02215, USA; Relay Therapeutics, 399 Binney Street, Cambridge, MA 02142, USA

**Author notes:** These authors contributed equally (Matthew J. K. Vince, Kristin A. Hughes, Anastasiya Buzuk).

**Keywords:** Clustering, PCA, PLSR, PLSDA

## Abstract

Machine learning (ML) is rapidly gaining traction in many areas of experimental molecular science for elucidating relationships and patterns in large or complex data sets. Historically, ML was largely the preserve of those with specialized training in fields such as statistics or cheminformatics. Increasingly, however, ML methodologies are becoming part of the standard toolkit for experimental scientists across a range of disciplines. Lowering the barrier of entry to these ML techniques, for scientists without a significant background in computer science or statistics, is important to broadening access to these powerful methods. Here we provide detailed, step by step tutorials for performing four ML methods that are particularly useful for applications in biochemistry, cell biology, and drug discovery: hierarchical clustering, Principal Component Analysis (PCA), Partial Least-Squares Discriminant Analysis (PLSDA), and Partial Least-Squares Regression (PLSR). The protocols are written for the widely used software MATLAB, but no prior experience with MATLAB is required to use them. We include an explanation of each step, pitched at a level to be understood by investigators without any prior experience with ML, MATLAB, or any kind of coding. We also highlight the scientific issues pertaining to selecting and scaling the data to be analyzed, and describe controls to test the validity of the results obtained. Throughout, we emphasize the relationship between the scientific question and how to choose data and methods that will allow it to be addressed in a meaningful way. Our aim is to provide a basic introduction that will equip experimental chemical biologists and other chemical and biomedical scientists with the knowledge required to use ML to aid in the design of experiments, the formulation and data-driven testing of hypotheses, and the analysis of experimental data.

**Basic Protocol 1:** Clustering

**Basic Protocol 2:** Principal Component Analysis (PCA)

**Basic Protocol 3:** Partial Least Squares Discriminant Analysis (PLSDA)

**Basic Protocol 4:** Partial Least Squares Regression (PLSR)

## I. INTRODUCTION

Machine learning (ML) is rapidly emerging as an essential tool for the molecular scientist. Searching PubMed for ‘machine learning’ returns only 58 articles published in the year 2000 and 718 such articles for 2010, while for 2023 this number grows to over 33,000. Over this period, applications of ML have expanded from specialized subfields of molecular bioscience, such as cheminformatics, bioinformatics, and the various ‘omics’ disciplines, to demonstrate powerful utility in virtually every area of molecular science. Consequently, basic literacy in ML is now required to understand a growing portion of the literature, and the ability to use ML is rapidly evolving from a niche expertise into an expectation for a well-trained experimental scientist.

A major barrier to learning ML is that the relevant literature tends to assume a knowledge of statistics and computer science that, while trivial to the specialist, may be lacking in the typical experimental scientist. Articles describing how to perform ML analyses that are pitched to readers without a computational background are hard to find. While developing an understanding of the key statistical concepts is unquestionably essential for the effective use of ML methods, when first approaching ML the use of unfamiliar terms or concepts can stymie the progress of even a motivated learner. This language barrier also exists in oral communications; personal experience has shown that it can be exceedingly difficult for an ML aficionado with expertise in computer science to effectively communicate to a nonspecialist even with high effort and goodwill on both sides. The protocols described herein were developed specifically to overcome this barrier. The goal is to enable experimental scientists not only to use the ML methods described in a technical sense, but also to understand each method well enough to design meaningful and rigorous analyses and appropriately interpret the results.

In offering this article, we wish to emphasize that we do not consider ourselves to be expert in Machine learning. Rather, we are experimental scientists who have worked to learn certain ML techniques, and to understand their operation and usage, much as we would learn an experimental method needed to advance our research. We believe that having approached the subject from this angle positions us well to pass on what we have learned to other experimental scientists. However, as learners ourselves, we are aware that our understanding may by incorrect or incomplete in places. It is our hope that, if readers detect any errors or imperfect understandings in the following article, they will be kind enough to notify us so we can make appropriate corrections.

### What is Machine Learning?

The term ‘Machine learning’ encompasses a diverse collection of methods for organizing and visualizing complex data to reveal non-obvious relationships and patterns, as well as for building models to reveal how input data relate to measurable or knowable outputs. Many ML methods are straightforward statistical manipulations for which the relationship of the input data to the final results can be fully understood and explained. All of the methods described in this article fall into this ‘explainable’ ML category. Other ML methods, such as Artificial Neural Networks and other ‘Deep Learning’ approaches, can generate highly accurate models to describe complex data and relationships, but generally do not provide any accessible explanation for precisely how the outcome derives from the input data and properties used. We have found the classical statistics-based ML methods of use for our goal of gaining insight into the principles, properties, or variables that govern a particular phenomenon, whereas Deep Learning might be more appropriate if the goal were solely to develop an accurate predictive model.

An additional dichotomy is whether the goal of the ML analysis is to explore the internal relationships between items in a data set or, alternatively, to model or predict some measurable output or behavior. Methods that do the former are termed ‘unsupervised’ (see Glossary), because there is no external outcome that the model is constrained to explain. In contrast, ML techniques that determine how the properties of items in a data set relate to a measurable outcome are termed ‘supervised’ methods (**Figure 1**).

**Figure 1.**
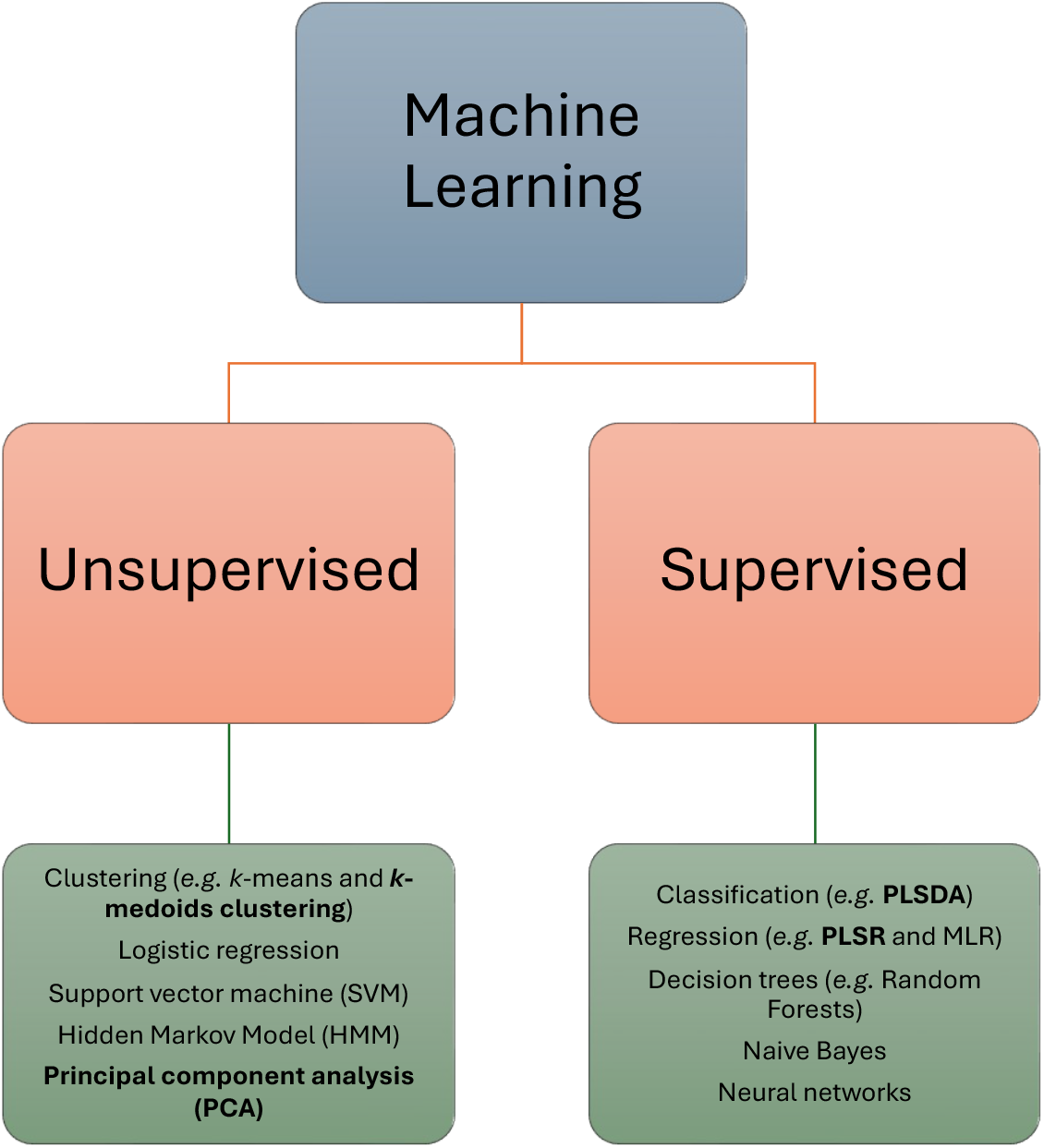
Categorization of ML methods as Supervised or Unsupervised, with selected examples of each. Methods described in this article are highlighted in bold. Abbreviations are: PLSR, partial least squares regression; PLSDA, partial least squares discriminant analysis; and MLR, multiple linear regression.

Here we present step-by-step protocols for implementing a set of supervised and unsupervised ML methods that we have found particularly useful in our research in protein biochemistry, small molecule ligand discovery, and analysis of protein-protein and protein-ligand interactions.

### Basic Protocol 1

describes how to take a set of elements (*e.g*. molecules, cells, etc.) and cluster them based on their similarity with respect to a chosen set of characteristics or properties. Clustering is a simple yet powerful method for establishing and graphically illustrating degrees of relatedness, particularly when the characteristics that are being used to assess the relatedness are relatively simple and easily understandable, and are not themselves the object of the analysis. A familiar application of clustering is in the multiple sequence alignment of protein or DNA sequences from different organisms, which allows construction of a dendrogram that illustrates the evolutionary relatedness among the sequences, but clustering has far broader applications than this. We describe three approaches to clustering that cover different purposes and generate different kinds of output.

### Basic Protocol 2

shows how to perform <u>Principal Component Analysis (PCA)</u>, a venerable and widely used method for determining and illustrating the relatedness between items with respect to a large number of characteristics or properties. PCA identifies a smaller number of composite properties (‘principal components’ (PC)) that capture the most important information contained in the original data set. By reducing the number of dimensions (*i.e*. properties or axes) to be considered, the method allows the information to be manipulated and graphically represented more simply. PCA has been used, for example, to integrate a complex set of intracellular signaling responses to elucidate how stimulation with different combinations of growth factors lead to cell proliferation versus cell death (Janes *et al*., 2006; Janes & Yaffe, 2006), or to compare the molecular structures of macrocyclic compounds to identify the subset of properties that characterize oral macrocyclic drugs (Viarengo-Baker *et al*., 2021).

### Basic Protocol 3

describes the method of <u>Partial Least-Squares Discriminant Analysis (PLSDA)</u>. This method is closely related to PLSR, but for situations in which the behavior to be explained is a categorical outcome, such as whether a cell survives or dies, rather than a quantity that can be expressed on a continuous numerical scale. For example, we used PLSDA to model how the physicochemical properties of certain residues in an enzyme correlate with whether the host bacterium grows naturally at low, moderate, or high temperatures, and showed that this model can identify amino acid substitutions that can increase the thermal stability of the protein (Muellers *et al*., 2023). ML methods that place items into categories in this way are called ‘classifiers’ (**Figure 1**).

### Basic Protocol 4

describes the method of <u>Partial Least-Squares Regression (PLSR)</u>. This method is related to PCA in that it condenses partially overlapping properties or characteristics into a smaller number of distinct ‘components’ but, unlike PCA, it is designed to determine how these properties combine to explain some measurable outcome or activity. Thus, PLSR can be used to create models that describe the relationship between a set of input properties and a measured outcome or behavior, and these models can then be used to predict the outcome for new cases based on their properties. The model is constructed so as to minimize the discrepancy between the predicted outcome for a particular item and the actual known outcome, akin to how linear regression minimizes the discrepancy between the best-fit line and the underlying experimental data, but for PLSR working in high dimensionality (*i.e*. considering many characteristics at once rather than a single x-axis variable). As examples, we have used PLSR to elucidate which molecular properties among a set of compounds correlate with permeability through a lipid membrane (Rzepiela *et al*., 2022), and which geometric properties among a set of molecules most influence affinity for binding to a target protein (Ortet *et al*., 2021).

For each protocol, we provide annotation and explanations designed to allow the nonspecialist investigator to understand (a) what each method does and how it works, (b) how to choose what data to include for a meaningful analysis, (c) how to properly curate the input data to eliminate unwanted biases, (d) how to assess the validity of the results, including control ‘experiments’ that can identify erroneous or misleading outcomes, and (e) how to interpret the results to obtain meaningful information about the molecules, process, or phenomenon under study. These issues are independent of which software is used to do the analysis. Thus, although the protocols described below are written for execution in MATLAB, the article may also have value for investigators using other software by providing an accessible introduction to ML for experimental scientists together with practical tips for how to set up a scientifically meaningful analysis and correctly interpret the results.

### The article is designed such that all the information required to execute a given method is contained within the corresponding protocol. It is not necessary to first complete earlier protocols in order to follow later ones

Basic Protocols (1), (2a), and (4), encompassing clustering, PCA, and PLSR, were originally developed as part of a four-week module for a graduate-level class in Quantitative Biochemistry, and were refined and expanded over several iterations of this course. The students consistently achieved good success in learning how to execute and understand the methods, and for multiple students it provided a starting point from which they went on to successfully use these or other ML methods in their PhD research. Importantly, having acquired proficiency with these initial methods, the students were then positioned to independently learn new ML techniques as needed for their research. Basic Protocol 1c (*k*-medoids clustering) and Basic Protocol 3 (PLSDA) are examples of additional methods that individual students learned and implemented during their research.

#### The Test System: A set of Cyclic Peptides with Known Membrane Permeabilities

We chose to illustrate the different ML methods by showing how each can be used to analyze the properties of a set of chemical compounds; specifically, a set of cyclic peptides. The unsupervised ML methods show how the compounds can be compared to each other and categorized with respect to these properties. In contrast, the supervised methods explore how these properties relate to the ability of the compounds to passively permeate through a lipid membrane. Membrane permeability is a key attribute for a compound to access intracellular targets or to be orally bioavailable, but is incompletely understood and difficult to predict. This is especially true for macrocyclic compounds such as cyclic peptides, and is currently an active area of investigation (Matsson *et al*., 2016; Cao *et al*., 2024)

In these worked examples, we use data downloaded from the recently published online database of cyclic peptide membrane permeability, CycPeptMPDB (http://cycpeptmpdb.com) (Li *et al*., 2023). CycPeptMPDB allows free access to independently deposited data on macrocyclic peptides, including an extensive set of pre-calculated physicochemical property values for each peptide as well as its experimentally measured membrane permeability value. Permeability values are reported from a range of different assay methods, but most often were measured using the parallel artificial membrane permeability assay (PAMPA) (Kansy *et al*., 1998). Due to the simplicity of PAMPA results, which do not require consideration of receptor-mediated efflux, this data type was selected for use in our protocol examples. We, and others, typically use the log_10_ of the permeability rather than the permeability itself because differences in the logarithmically transformed values are proportional to differences in the free energy barrier for passing through the membrane.

### We provide .xlsx files containing the data used in each protocol in the Supplementary Information so readers can perform the protocols without having to find and download the data from the original source

We also include instructions for how the original data were prepared and formatted for use, to aid users who wish to apply these protocols to other kinds of data of interest to them, in the Supplementary Information (Data_prepared_SI.docx).

#### How to Use These Protocols

The protocols described below are written for execution in MATLAB. **No prior experience with MATLAB is assumed**. However, users who are entirely unfamiliar with this program are recommended to first complete MATLAB’s free, self-paced online course, ‘MATLAB Onramp’ (https://MATLABacademy.mathworks.com/details/MATLAB-onramp/gettingstarted).

Readers are recommended to begin by reading the introductory material for the protocol in question to understand what the method aims to achieve and, at a conceptual level, how it works. Readers should then execute the protocol step by step, typing or pasting each command into the MatLab Command window and pressing return, to see what it does. By reading the associated comments and comparing your results with those provided in the protocol, you should aim to understand, for each step: (i) What is the purpose of the step? (ii) What data are used as input for the step, and how are these formatted? (iii) What is the output from the step, and what form does this output take? (iv) What was the overall effect of the step, and how can you tell whether or not it worked as intended? Experimenting with how the results of a given step are affected if a user-defined parameter or input value is changed can also be informative.

All commands included in each protocol are additionally provided as MATLAB script editor files (.mlx) in the supplemental documentation, which can be opened in MATLAB to give a functioning script that requires only input of appropriate data. These scripts obviate the need for step-by-step operation of the protocols, allowing the methods to be executed much more quickly. Our intent is that a reader who understands points (i)-(iv) from the preceding paragraph will be able to modify these scripts to use the methods for different purposes using the reader’s own data.

## II. GLOSSARY OF RELEVANT MACHINE LEARNING TERMINOLOGY

**Table 1.**
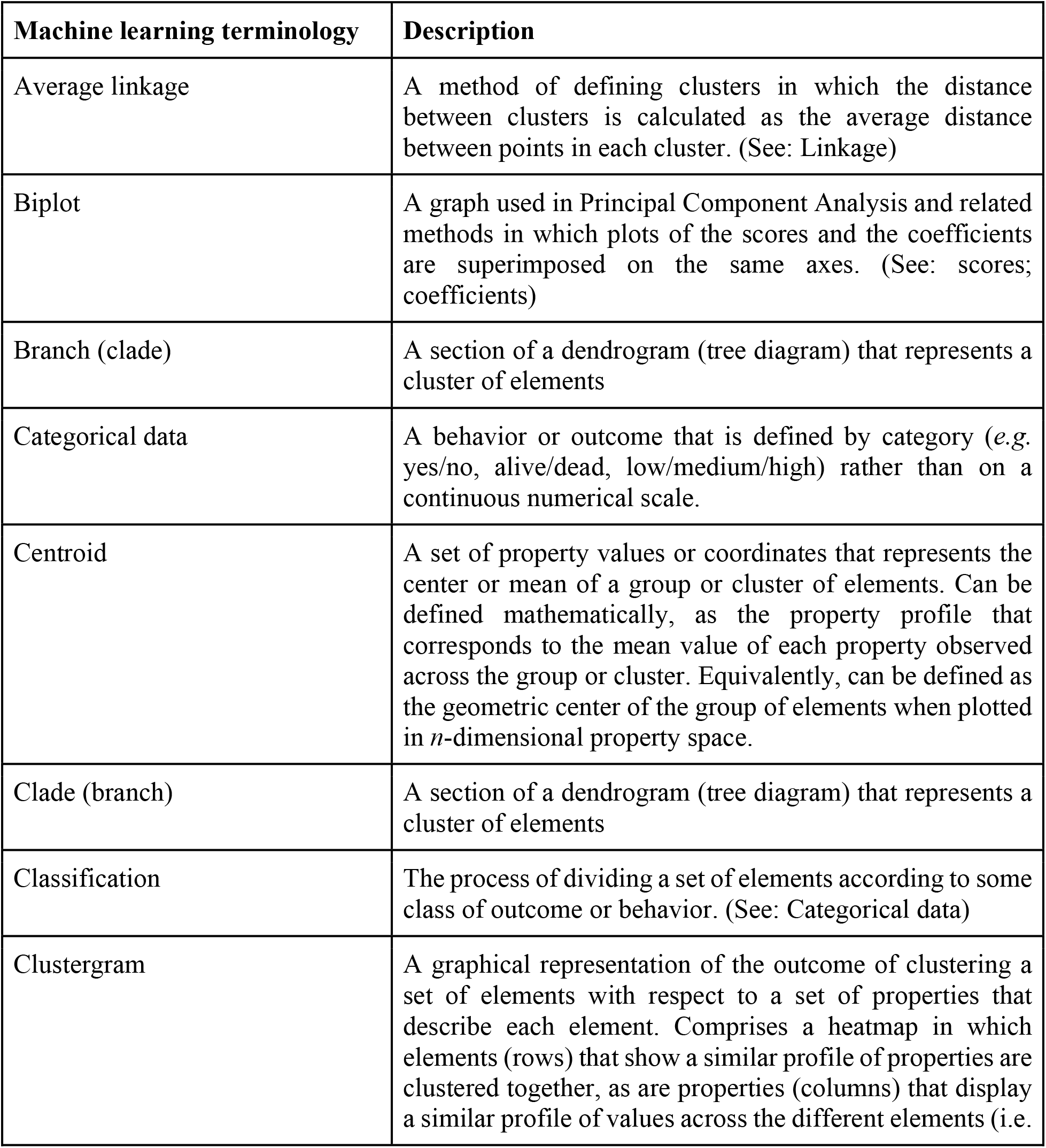
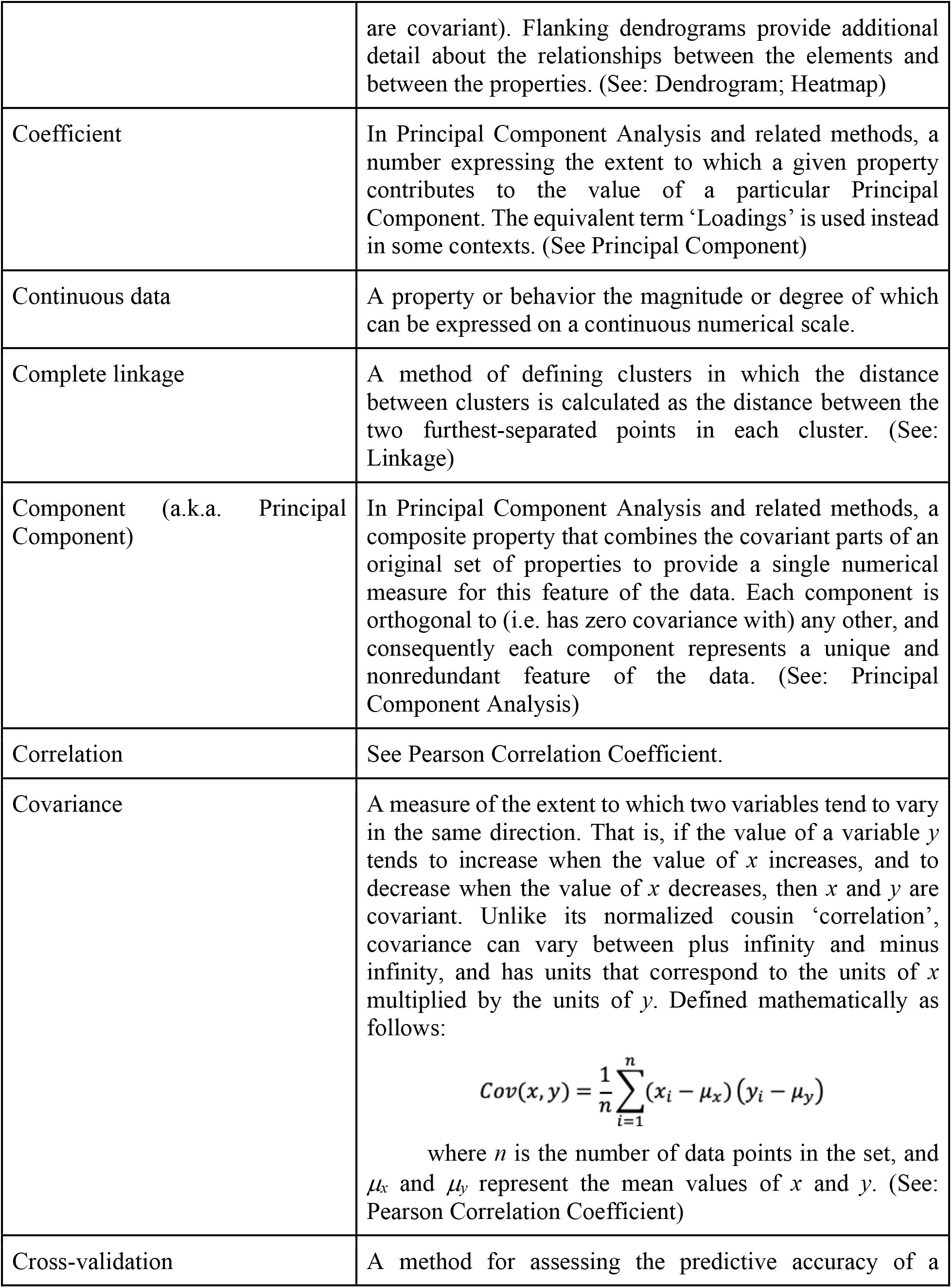
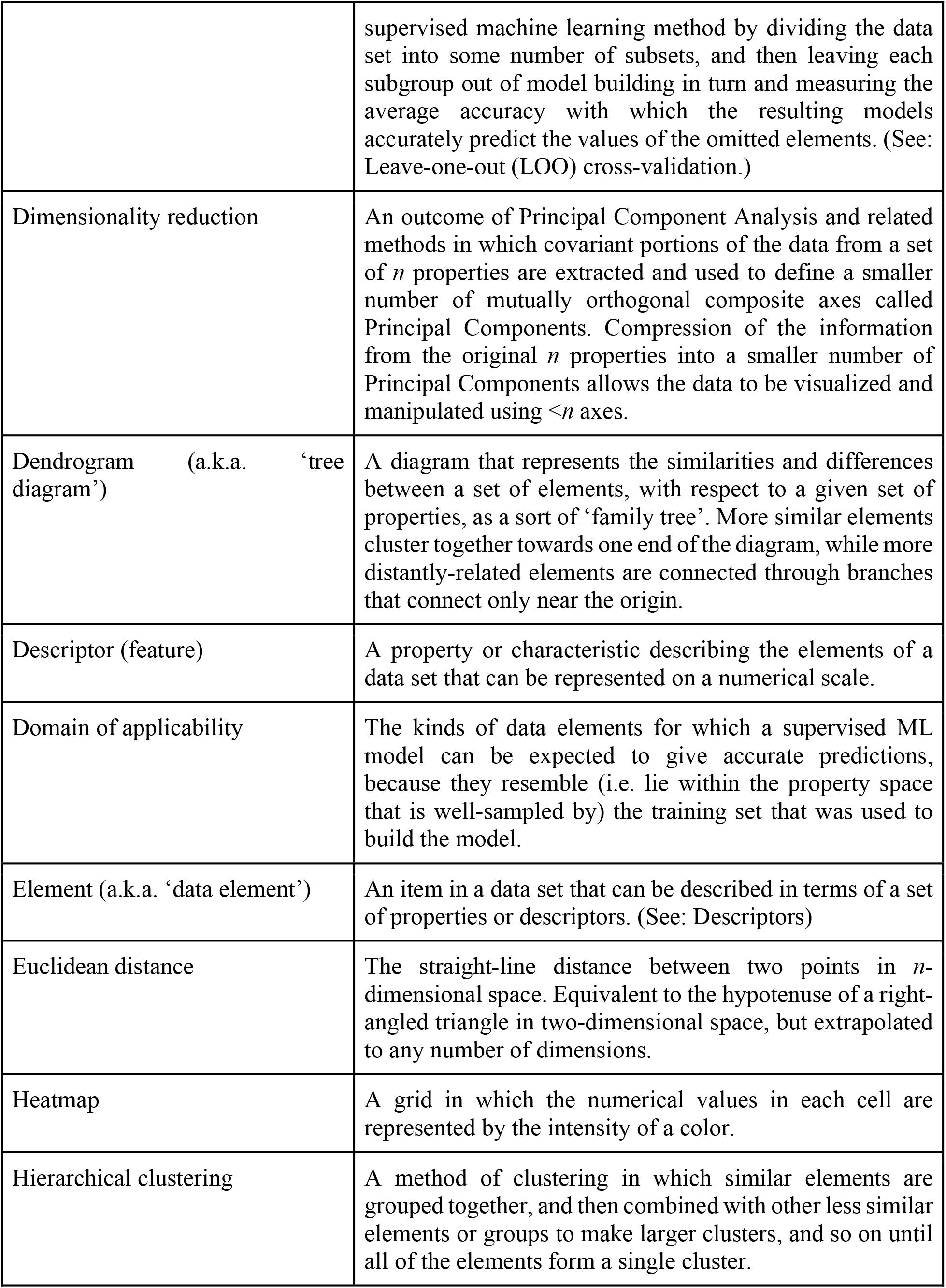
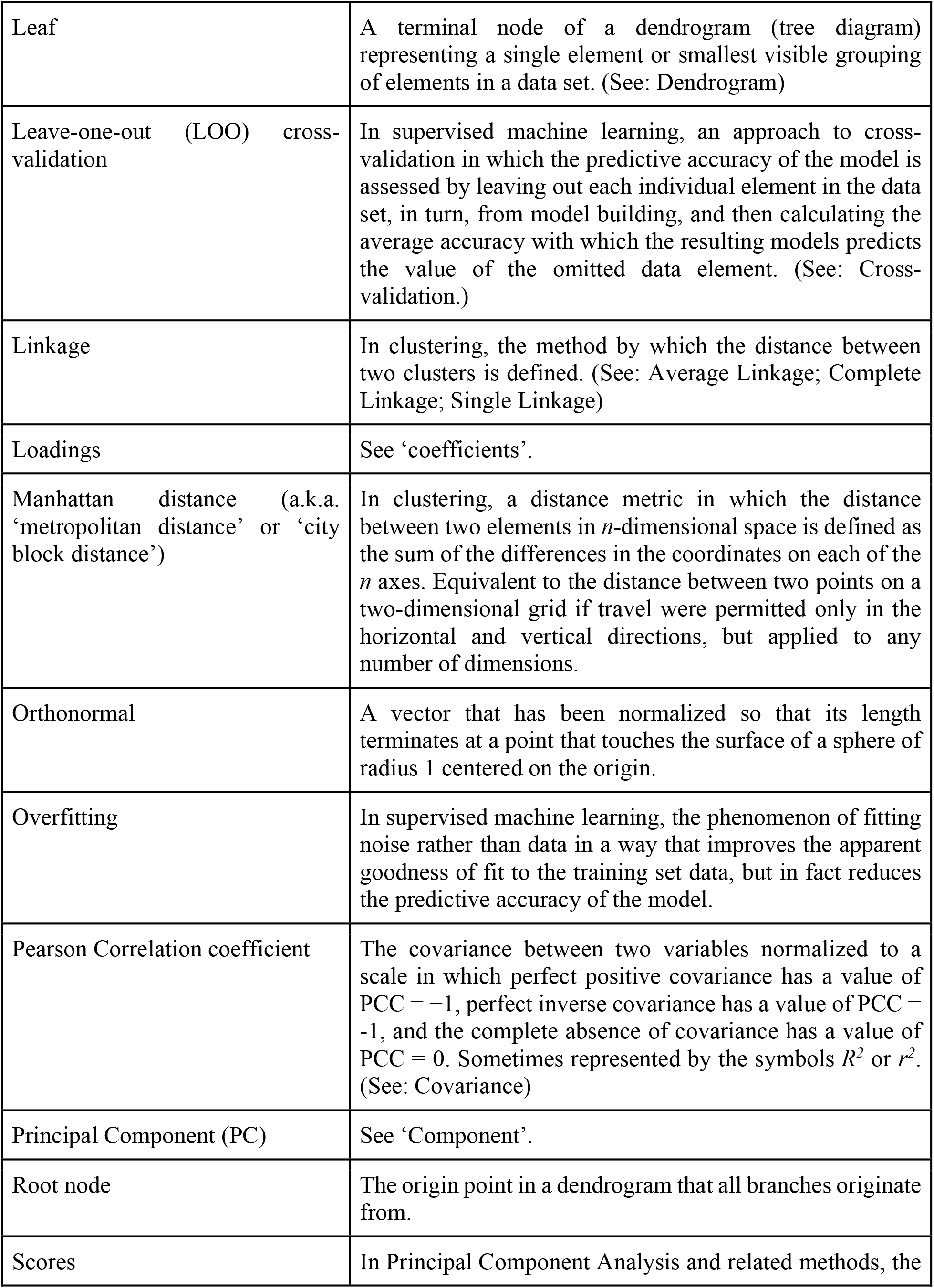
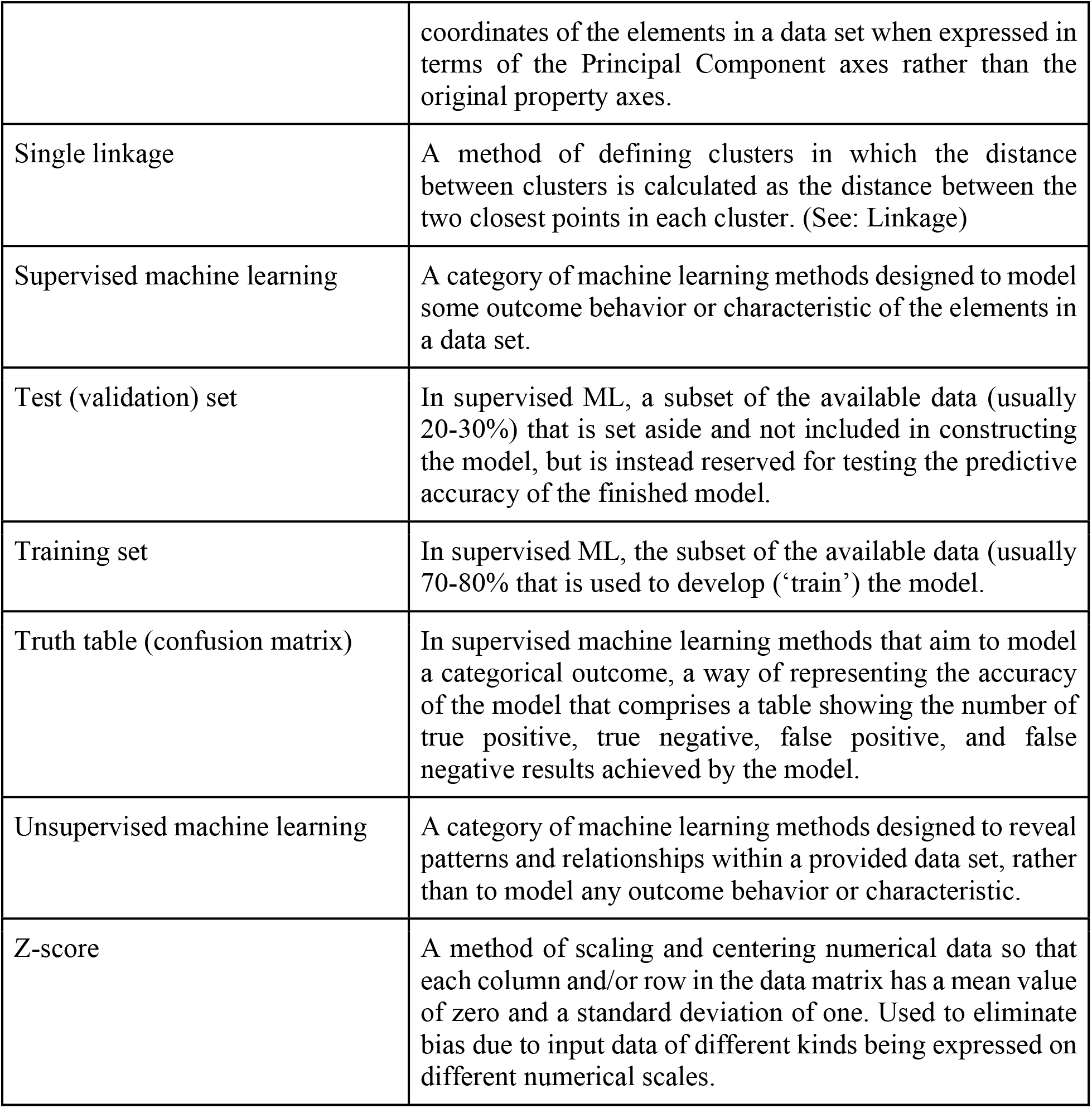
Important ML terminology discussed in this article, and the ML protocol(s) in which each is used.

| Machine learning terminology | Description |
| --- | --- |
| Average linkage | A method of defining clusters in which the distance between clusters is calculated as the average distance between points in each cluster. (See: Linkage) |
| Biplot | A graph used in Principal Component Analysis and related methods in which plots of the scores and the coefficients are superimposed on the same axes. (See: scores; coefficients) |
| Branch (clade) | A section of a dendrogram (tree diagram) that represents a cluster of elements |
| Categorical data | A behavior or outcome that is defined by category (*e.g.* yes/no, alive/dead, low/medium/high) rather than on a continuous numerical scale. |
| Centroid | A set of property values or coordinates that represents the center or mean of a group or cluster of elements. Can be defined mathematically, as the property profile that corresponds to the mean value of each property observed across the group or cluster. Equivalently, can be defined as the geometric center of the group of elements when plotted in $n$-dimensional property space. |
| Clade (branch) | A section of a dendrogram (tree diagram) that represents a cluster of elements |
| Classification | The process of dividing a set of elements according to some class of outcome or behavior. (See: Categorical data) |
| Clustergram | A graphical representation of the outcome of clustering a set of elements with respect to a set of properties that describe each element. Comprises a heatmap in which elements (rows) that show a similar profile of properties are clustered together, as are properties (columns) that display a similar profile of values across the different elements (*i.e.* |
|  | are covariant). Flanking dendrograms provide additional detail about the relationships between the elements and between the properties. (See: Dendrogram; Heatmap) |
| Coefficient | In Principal Component Analysis and related methods, a number expressing the extent to which a given property contributes to the value of a particular Principal Component. The equivalent term ‘Loadings’ is used instead in some contexts. (See Principal Component) |
| Continuous data | A property or behavior the magnitude or degree of which can be expressed on a continuous numerical scale. |
| Complete linkage | A method of defining clusters in which the distance between clusters is calculated as the distance between the two furthest-separated points in each cluster. (See: Linkage) |
| Component (a.k.a. Principal Component) | In Principal Component Analysis and related methods, a composite property that combines the covariant parts of an original set of properties to provide a single numerical measure for this feature of the data. Each component is orthogonal to (i.e. has zero covariance with) any other, and consequently each component represents a unique and nonredundant feature of the data. (See: Principal Component Analysis) |
| Correlation | See Pearson Correlation Coefficient. |
| Covariance | A measure of the extent to which two variables tend to vary in the same direction. That is, if the value of a variable $y$ tends to increase when the value of $x$ increases, and to decrease when the value of $x$ decreases, then $x$ and $y$ are covariant. Unlike its normalized cousin ‘correlation’, covariance can vary between plus infinity and minus infinity, and has units that correspond to the units of $x$ multiplied by the units of $y$. Defined mathematically as follows:<br>$Cov(x, y) = \frac{1}{n} \sum_{i=1}^n (x_i - \mu_x)(y_i - \mu_y)$<br>where $n$ is the number of data points in the set, and $\mu_x$ and $\mu_y$ represent the mean values of $x$ and $y$. (See: Pearson Correlation Coefficient) |
| Cross-validation | A method for assessing the predictive accuracy of a |
|  | supervised machine learning method by dividing the data set into some number of subsets, and then leaving each subgroup out of model building in turn and measuring the average accuracy with which the resulting models accurately predict the values of the omitted elements. (See: Leave-one-out (LOO) cross-validation.) |
| Dimensionality reduction | An outcome of Principal Component Analysis and related methods in which covariant portions of the data from a set of $n$ properties are extracted and used to define a smaller number of mutually orthogonal composite axes called Principal Components. Compression of the information from the original $n$ properties into a smaller number of Principal Components allows the data to be visualized and manipulated using $<n$ axes. |
| Dendrogram (a.k.a. ‘tree diagram’) | A diagram that represents the similarities and differences between a set of elements, with respect to a given set of properties, as a sort of ‘family tree’. More similar elements cluster together towards one end of the diagram, while more distantly-related elements are connected through branches that connect only near the origin. |
| Descriptor (feature) | A property or characteristic describing the elements of a data set that can be represented on a numerical scale. |
| Domain of applicability | The kinds of data elements for which a supervised ML model can be expected to give accurate predictions, because they resemble (i.e. lie within the property space that is well-sampled by) the training set that was used to build the model. |
| Element (a.k.a. ‘data element’) | An item in a data set that can be described in terms of a set of properties or descriptors. (See: Descriptors) |
| Euclidean distance | The straight-line distance between two points in $n$-dimensional space. Equivalent to the hypotenuse of a right-angled triangle in two-dimensional space, but extrapolated to any number of dimensions. |
| Heatmap | A grid in which the numerical values in each cell are represented by the intensity of a color. |
| Hierarchical clustering | A method of clustering in which similar elements are grouped together, and then combined with other less similar elements or groups to make larger clusters, and so on until all of the elements form a single cluster. |
| Leaf | A terminal node of a dendrogram (tree diagram) representing a single element or smallest visible grouping of elements in a data set. (See: Dendrogram) |
| Leave-one-out (LOO) cross-validation | In supervised machine learning, an approach to cross-validation in which the predictive accuracy of the model is assessed by leaving out each individual element in the data set, in turn, from model building, and then calculating the average accuracy with which the resulting models predicts the value of the omitted data element. (See: Cross-validation.) |
| Linkage | In clustering, the method by which the distance between two clusters is defined. (See: Average Linkage; Complete Linkage; Single Linkage) |
| Loadings | See ‘coefficients’. |
| Manhattan distance (a.k.a. ‘metropolitan distance’ or ‘city block distance’) | In clustering, a distance metric in which the distance between two elements in $n$-dimensional space is defined as the sum of the differences in the coordinates on each of the $n$ axes. Equivalent to the distance between two points on a two-dimensional grid if travel were permitted only in the horizontal and vertical directions, but applied to any number of dimensions. |
| Orthonormal | A vector that has been normalized so that its length terminates at a point that touches the surface of a sphere of radius 1 centered on the origin. |
| Overfitting | In supervised machine learning, the phenomenon of fitting noise rather than data in a way that improves the apparent goodness of fit to the training set data, but in fact reduces the predictive accuracy of the model. |
| Pearson Correlation coefficient | The covariance between two variables normalized to a scale in which perfect positive covariance has a value of $PCC = +1$, perfect inverse covariance has a value of $PCC = -1$, and the complete absence of covariance has a value of $PCC = 0$. Sometimes represented by the symbols $R^2$ or $r^2$. (See: Covariance) |
| Principal Component (PC) | See ‘Component’. |
| Root node | The origin point in a dendrogram that all branches originate from. |
| Scores | In Principal Component Analysis and related methods, the |
|  | coordinates of the elements in a data set when expressed in terms of the Principal Component axes rather than the original property axes. |
| Single linkage | A method of defining clusters in which the distance between clusters is calculated as the distance between the two closest points in each cluster. (See: Linkage) |
| Supervised machine learning | A category of machine learning methods designed to model some outcome behavior or characteristic of the elements in a data set. |
| Test (validation) set | In supervised ML, a subset of the available data (usually 20-30%) that is set aside and not included in constructing the model, but is instead reserved for testing the predictive accuracy of the finished model. |
| Training set | In supervised ML, the subset of the available data (usually 70-80% that is used to develop ('train') the model. |
| Truth table (confusion matrix) | In supervised machine learning methods that aim to model a categorical outcome, a way of representing the accuracy of the model that comprises a table showing the number of true positive, true negative, false positive, and false negative results achieved by the model. |
| Unsupervised machine learning | A category of machine learning methods designed to reveal patterns and relationships within a provided data set, rather than to model any outcome behavior or characteristic. |
| Z-score | A method of scaling and centering numerical data so that each column and/or row in the data matrix has a mean value of zero and a standard deviation of one. Used to eliminate bias due to input data of different kinds being expressed on different numerical scales. |

## III. REQUIRED SOFTWARE

For the ML methods described herein we use MATLAB (Paluszek and Thomas, 2020). MATLAB can be purchased and downloaded from MathWorks at the following link: https://www.mathworks.com/licensecenter/licenses/add?s_tid=ML_mod_pers_cwl. There are many other software options for performing the analyses we describe, including the freeware R (Lantz, 2019). Whatever the software used the steps are analogous, and the discussion of descriptor selection, data preparation, data validation and analysis, and troubleshooting are applicable regardless of the product used.

*The protocols were written using MATLAB R2023a. We cannot guarantee that changes to MATLAB in future releases won’t result in minor deviations from the precise commands shown here. If a command does not work as written, readers are advised to check the MATLAB online documentation for the command to see whether the syntax has changed since version R2023a*.

Users will also need to download and install some additional MATLAB Toolboxes, as described below.

*While MATLAB does not always automatically restart upon downloading a toolbox, it is a good practice to restart MATLAB after downloading and setting the path to ensure all changes take effect*.

### MATLAB toolboxes required for All Protocols

All protocols require installation of the *Statistics and Machine Learning* Toolbox and the *Clustering* Toolbox.

*If you are downloading MATLAB for the first time, toolboxes will be offered during the initial startup of the program and can be selected at that time. In the case that MATLAB is already downloaded, these toolboxes may be separately downloaded and installed from the following links:*

1. Statistics and Machine Learning Toolbox: https://www.mathworks.com/products/statistics.html
2. Clustering Toolbox: https://www.mathworks.com/MATLABcentral/fileexchange/7486-clustering-toolbox

### Additional MATLAB toolbox required for Basic Protocol 1a (Clustering)

The clustering analyses additionally require the *Bioinformatics ToolBox*. This toolbox includes the modules for phylogenetic analysis using the dendrogram function. It can be downloaded from: https://www.mathworks.com/products/bioinfo.html.

### Additional MATLAB toolbox required for Basic Protocol 3 (PLSDA)

The PLSDA analysis additionally requires the *Classification ToolBox*, as described by Balabio and Consonni’s work (Ballabio and Consonni, 2013). This toolbox represents a collection of MATLAB modules for supervised pattern recognition developed by The University of Milan-Bicocca’s Chemometric and QSAR Research Group. It can be downloaded from: https://michem.unimib.it/download/MATLAB-toolboxes/classification-toolbox-for-MATLAB/.

After downloading a Toolbox, If the file is compressed (*e.g*. a ZIP file), it should be uncompressed using appropriate software (*e.g*. ‘Unarchiver’ available at https://theunarchiver.com/). It may also be necessary to set the path so MATLAB can connect to the toolbox. To do this, open MATLAB and go to the “Home” tab, and select ‘Set Path’. In the Set Path dialog box, click on ‘Add Folder’. Navigate to the location where you extracted your toolbox, select the folder, and click OK. After adding the folder, click ‘Save’ and then ‘Close’ in the ‘Set Path’ dialog box.

*Ensure you select the folder containing the toolbox files directly. Adding an outer folder (e.g*., *a top-level folder created during extraction) may not work correctly*.

*To confirm that MATLAB recognizes the toolbox, try using one of the functions from the toolbox. For example, type ‘help plsdacompsel’ in the Command window and check if MATLAB recognizes it without error*.

*For additional information on how to set paths, see MATLAB help https://www.mathworks.com/help/matlab/matlab_env/add-remove-or-reorder-folders-on-the-search-path.html*

## IV. DATA DOWNLOAD AND PREPARATION

The ML methods described herein require the input data to be provided as a numeric matrix, conventionally called ‘X’, which comprises the numerical values of *n* features or characteristics for each of *m* items or elements to be analyzed. For example, in the protocols described below, the input data comprise the values of *n* molecular properties for a set of *m* macrocyclic peptides. The input matrix, X, therefore comprises an *m* x *n* matrix where each column is a different molecular property and each row is a different macrocyclic peptide. The cells of this matrix each contain the numerical value of a particular property for one peptide. This format of rows that contain the items to be analyzed and columns that contain the properties to be considered in the analysis is fairly standard for ML.

For all protocols, the left-most column (*i.e*. Column 1) of the input data matrix X contains the ID numbers of the *m* compounds. The other cells in any given row contain the descriptor values pertaining to the compound named in column 1 of that row. The top row of X contains the names of the *n* molecular descriptors, such that each column in X is headed by a descriptor name and contains the values of that descriptor for each compound in the set. Basic Protocols 3 and 4 also require an output matrix, *y*, comprising *m* rows and 1 column. The output matrix *y* contains the measured membrane permeability values of permeability class (Permeable/Impermeable) that is being modeled in these two protocols.

For use in the following protocols, the data should be formatted as shown in **Figure 2** to facilitate subsequent import into MATLAB. The data may be prepared in a program such as Microsoft Excel and saved in a variety of file formats, *e.g*. .xlsx or .csv.

**Figure 2.**
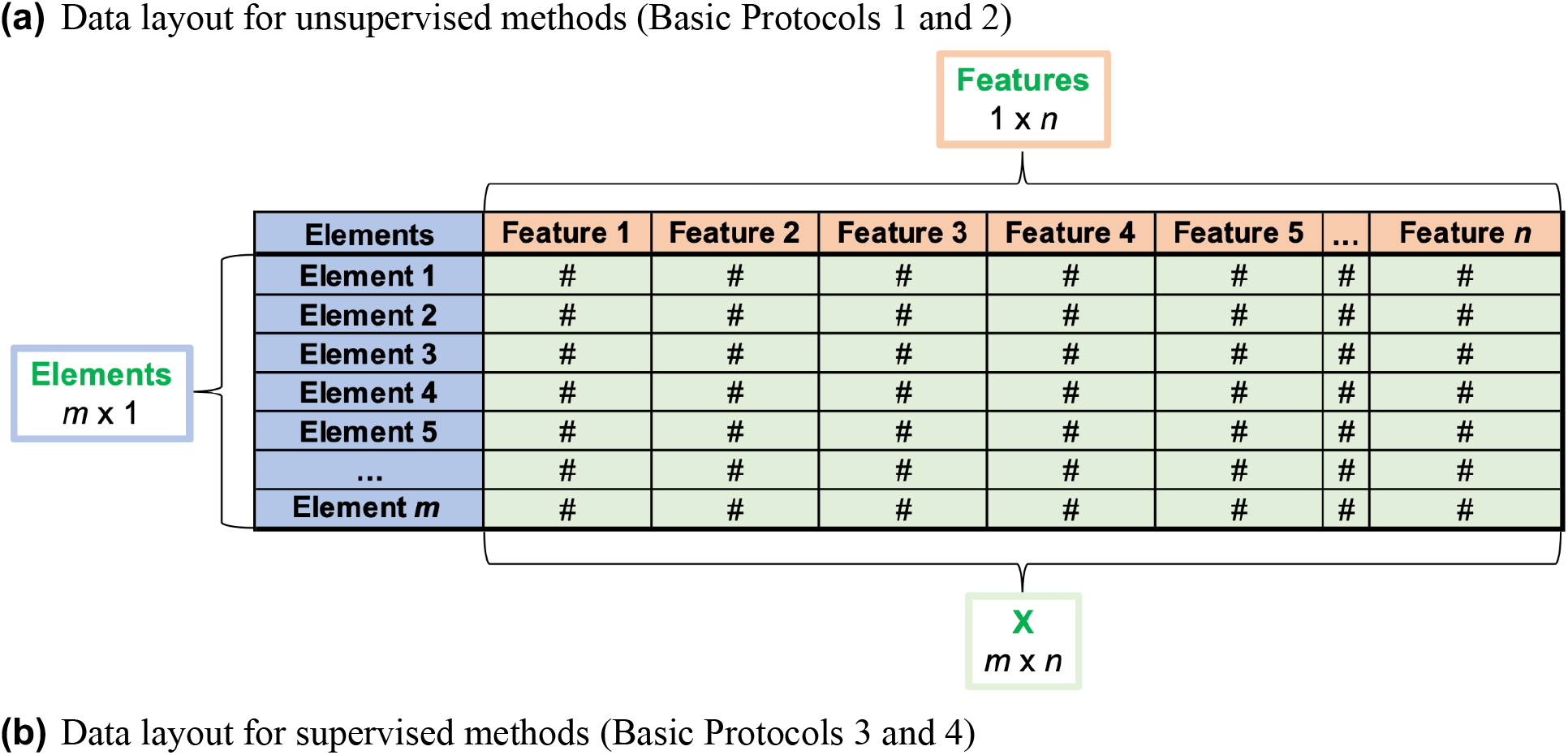
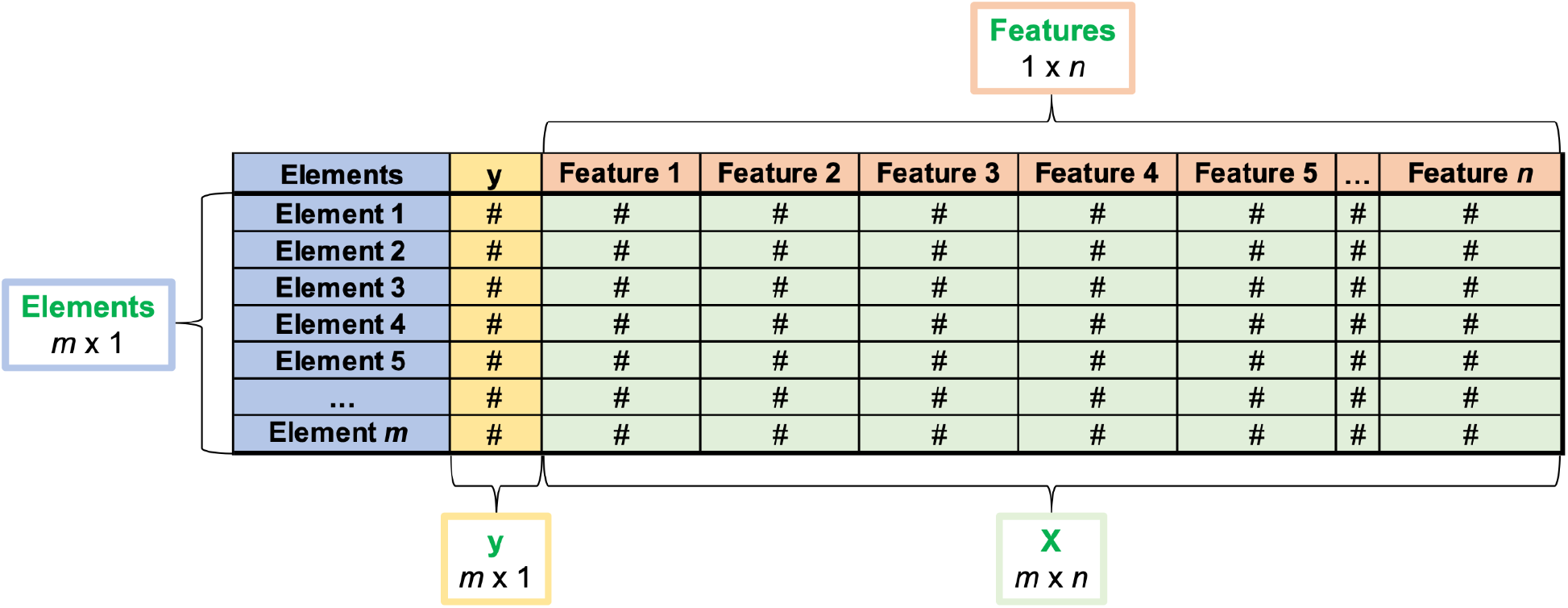
Data layout for import into MATLAB. **(a)** The unsupervised ML methods described in Basic Protocols 1 and 2 require an input data file, conventionally called X, that contains *m* rows, one for each item or element in the set to be analyzed, and *n* columns, one for each feature or property to be used to compare the elements. An additional column to the left of the numerical data should contain the names of the items or elements that the numbers in each row describe. Similarly, an additional row at the top of the matrix should contain the names of the feature that the numerical values in each column quantify. **(b)** The supervised ML methods described in Basic Protocols 3 and 4 additionally require an output matrix, conventionally called *y*, that contains the experimentally determined behavior to be modeled. In the examples shown here, *y* comprises a single column that contains *m* values, one for each element. This column is placed to the left of the *m* x *n* number matrix, immediately to the right of the list of element names. Output matrices that comprise more than one column of values can also be used, but the examples illustrated in Basic Protocols 3 and 4 involve modeling only a single behavior represented by a single column of values.

All data files needed to perform the protocols described in this article are provided as supplementary files and are already formatted as shown in **Figure 2**. Details of how these data files were prepared from the raw files downloaded from the original source, the CycPeptMPDB database (Li *et al*., 2023), are provided in the Supplementary File called ‘Data_prepared_SI.docx’. Readers who create their own data files for use with these protocols should take care to make sure that these external files are given the same names we use in the protocols below (shown in blue) or, alternatively, should modify the commands shown below to match the names they give the external files.

## V. How to Use these Protocols

In the following protocols, commands to be typed/pasted into MATLAB are shown in **black and bolded**. Any parts of the command that require the user to choose and enter a value or setting are colored **red**. External data files (*e.g*. .xlsx, .csv, etc.) to be uploaded as part of the command are colored blue. Any variables created using MATLAB’s ‘Import Data’ function that are stored in the MATLAB workspace are colored **green**. These colors are solely to help users to understand these protocols. You do not need to use any particular text colors when typing in the Command window; MATLAB uses text colors for different purposes and will assign its own colors to the text you enter.

### Helpful MATLAB Tips & Tricks

- Copying and pasting commands from certain word processing software can sometimes result in an error message: *‘Error: The input character is not valid in MATLAB statements or expressions.’* Most often, this error results from the fact that MATLAB only accepts apostrophes and not ‘smart apostrophes’, as are sometimes the default *e.g*. in Microsoft Word. To fix this issue you can change the settings in the word processing software to disable smart apostrophes. Alternatively, working directly in the MATLAB Command window, you can simply delete the apostrophes and retype them.
- When importing different subsets of rows/columns from an external file to create multiple separate MATLAB data objects, each newly created MATLAB object must be renamed before more selections are imported from the same external file. This is because the name for the newly created object that MATLAB initially assigns is the name of the external file from which the data were imported. Consequently, if the created object is not immediately renamed, the next set of rows/columns imported from the same external file will be given the same default name and will overwrite the first.
- MATLAB requires that a variable name starts with a letter followed, if desired, by additional characters comprising letters, numbers or underscores. The name is case sensitive. Further information on naming MATLAB files can be found at the following link: https://www.mathworks.com/help/MATLAB/MATLAB_prog/variable-names.html
- When working in the Command window, previously used commands can be accessed and re-used without having to type them in again. This is done by pressing the ‘up’ arrow on the keyboard, when in the Command window, and selecting the desired command from the dropdown menu that appears. Multiple commands can be selected by clicking and dragging accordingly. The commands can be edited, *e.g*. to correct an error or change a setting, before pressing enter to execute the command.
- Many MATLAB commands take the form [variable_name(s)] = command(data_object_name(s)). For example, the zscore command (about which more later) has the form

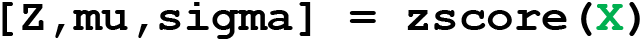

The effect is to perform the operation ‘zscore’ on the data in the data variable named X, and deposit different aspects of the results into three new variables called ‘Z’, ‘mu’, and ‘sigma’. The labels ‘Z’, ‘mu’ and ‘sigma’ are arbitrary, and are simply names that are often chosen for the variables created by this command. Any valid filenames (e.g. [A,B,C]) could be used in the left-hand side of this command. Most MATLAB commands share this flexibility with regard to the names of the new variables they create.

- Adding a semicolon after a command instructs MATLAB not to display the results of the operation in the Command window. If the semicolon is omitted, the command may return many lines of results as it displays all of the numbers and text in any data variables that were created. Adding the semicolon silences this display, so the results are stored as data variables in the Workspace window but are not also printed out in full in the Command window.
- In MATLAB, placing a period immediately before a mathematical operation, as in ‘.*’ or ‘./’, instructs that the operation be performed separately on each value in the matrix being operated on. If the period is omitted, the command would instead perform matrix multiplication, matrix division, etc. with different results.
- To save the workspace at any time, select the ‘Save Workspace’ option under the Home tab. This will save the workspace as a .mat file. We suggest saving the workspace after importing the required data files into the workspace, and again after completing the protocol, using different filenames to distinguish the two files.
- When a command creates a plot or figure, the figure opens automatically in a separate MATLAB Figure window. The figure can be edited in this window by selecting the ‘Tools’ tab and then ‘Edit plot’, which will open a window with various editing options. If the Figure window is obscuring other MATLAB windows you need to access, for example to enter formatting commands into the Command window, it can be dragged to one side or minimized. Do not close the figure file while entering successive formatting commands.
- Figures can be saved in several different file formats. Having formatted the figure to give the desired appearance, it can be saved as an image file (*e.g*. jpg). However, we recommend also saving each figure as a modifiable MATLAB figure file (.fig), in case you wish to edit its appearance at a later date. The file format is chosen from the ‘File’ tab by selecting ‘Save As’ choosing the desired output file type. We recommend you save each figure as a .fig file as soon as it has been formatted as desired.
- MATLAB provides detailed documentation about any function or command available in MATLAB, including its usage, input and output arguments, examples, and related functions. To get help on a specific function, type ‘doc’ followed by the name of the function in the Command window and press Enter. Doing this will open a new window containing full documentation on the command and how to use it. Help with a function can also be obtained directly in the Command window by entering ‘help’ followed by the command name.

## VI. UNSUPERVISED ML METHODS: Clustering and Principal Component Analysis (PCA)

Unsupervised ML methods analyze a data set to identify relationships and patterns among the different items in the set. A common goal of unsupervised learning is to group the items in the data set into clusters, such that items that are similar to each other with respect to the properties of interest are clustered together while more dissimilar items are placed in different clusters. In essence, clustering calculates the degree of relatedness of each item in the data set to each other item with respect to a set of properties of interest. Clustering thereby provides a means to evaluate which items can be grouped together as sharing broadly similar properties, and enables the representation of these groupings and subgroupings in an easily interpretable two-dimensional graph. There are many different algorithms for clustering, and each involves certain decisions the user must make about how relatedness is quantified and how the groups are formed, as described in more detail in the next section. In this article, we describe two different approaches to clustering, one designed to reveal the internal relationships in the data with no preconceived notion of how many groups might exist, and the other situations in which the user wishes to cluster the data into a preselected number of groups.

Principal Component Analysis (PCA) is another venerable yet powerful and widely used unsupervised ML method. The distinctive feature of PCA is that it has the effect of compressing the input data to eliminate any redundant information. Thus, if the original input data comprises the values of *n* properties for each of *m* items in the set, PCA will identify properties that contain substantial covariance and combine them into a smaller set of new composite properties, called Principal Components (PCs), that contain the same information. PCA orders the PCs such that the component that contains the most statistical variance is numbered PC1, the PC containing the second largest share of variance is numbered PC2, and so on. Because of this data compression and ordering of PCs, it is often possible to capture most of the total information (variance) present in the original *n* x *m* input data matrix in just the first two or three PCs, allowing the data to be visualized using 2D or 3D plots with minimal loss of information. Analysis of such plots identifies which items in the set are more or less similar to which other items, as similar items will cluster together while dissimilar items will appear far apart. By examining which of the original properties contributed most strongly to the first few PCs, it is also possible to understand which of the original properties contribute most to differentiating the items in the data set from one another.

Both clustering and PCA reveal relationships between items in the data set with respect to a specified set of properties. But clustering considers each property to contribute independently to differentiating the items, while PCA takes into account and eliminates any redundancy due to covariance between the original properties. Clustering is therefore appropriate for visualizing the similarities and differences between items in cases where there is high confidence that all of the included properties included in the analysis are individually and separately relevant. PCA, on the other hand, is useful for identifying the minimum number of distinct and nonredundant features that differentiate the items in the data set, and for identifying which kinds of properties comprise each feature.

More detailed descriptions of clustering and PCA are provided in the corresponding sections, below.

## BASIC PROTOCOL 1: Clustering

### Purpose of method

To compare a set of items using a list of properties or characteristics, with the goals of revealing how the items organize into groups and subgroups based on similarities in their characteristics, and quantifying how similar or different any item is to any other item in the set with respect to those characteristics. Here we show how to cluster a set of chemical compounds with respect to 139 structural and physicochemical properties, and how to identify a representative subset of any chosen size from among these compounds.

### Background

Clustering is a branch of unsupervised machine learning that reveals patterns within a data set based on the similarities or differences between items (of any kind) with respect to a user-chosen set of numerical properties or characteristics. Items that are more similar with respect to the included characteristics will tend to cluster together, while items with very different characteristics will appear far apart. Therefore, the outcome of clustering depends on both the items being clustered and the characteristics that are used to assess their degree of similarity or difference.

If clustering *m* different items with respect to *n* characteristics, the data to be used for clustering can be represented as a numerical matrix with *m* rows (one for each item in the set), and *n* columns (one for each property or characteristic to be considered) (**Figure 1.1)**. Each cell in this *m* x *n* matrix contains the numerical value of a given property for one of the items. In principle, one way of representing such a data set would be to plot each item as a point on a set of *n* Cartesian axes (one axis for each property), with the coordinates of each point corresponding to the item’s numerical value for each property. For example, if only two properties are being considered, then each item can be represented as a point on a set of x-y axes, with the x-coordinate corresponding to the value of property 1 and the y-coordinate the value of property 2. Of course, if the number of properties is greater than three, such a graph would have many mutually orthogonal axes and thus many dimensions, making it impossible or very difficult to accurately represent in the two dimensions of a flat page or screen. Simplifying the analysis and representation of such high dimensionality data is one valuable outcome of clustering.

**Figure 1.1.**
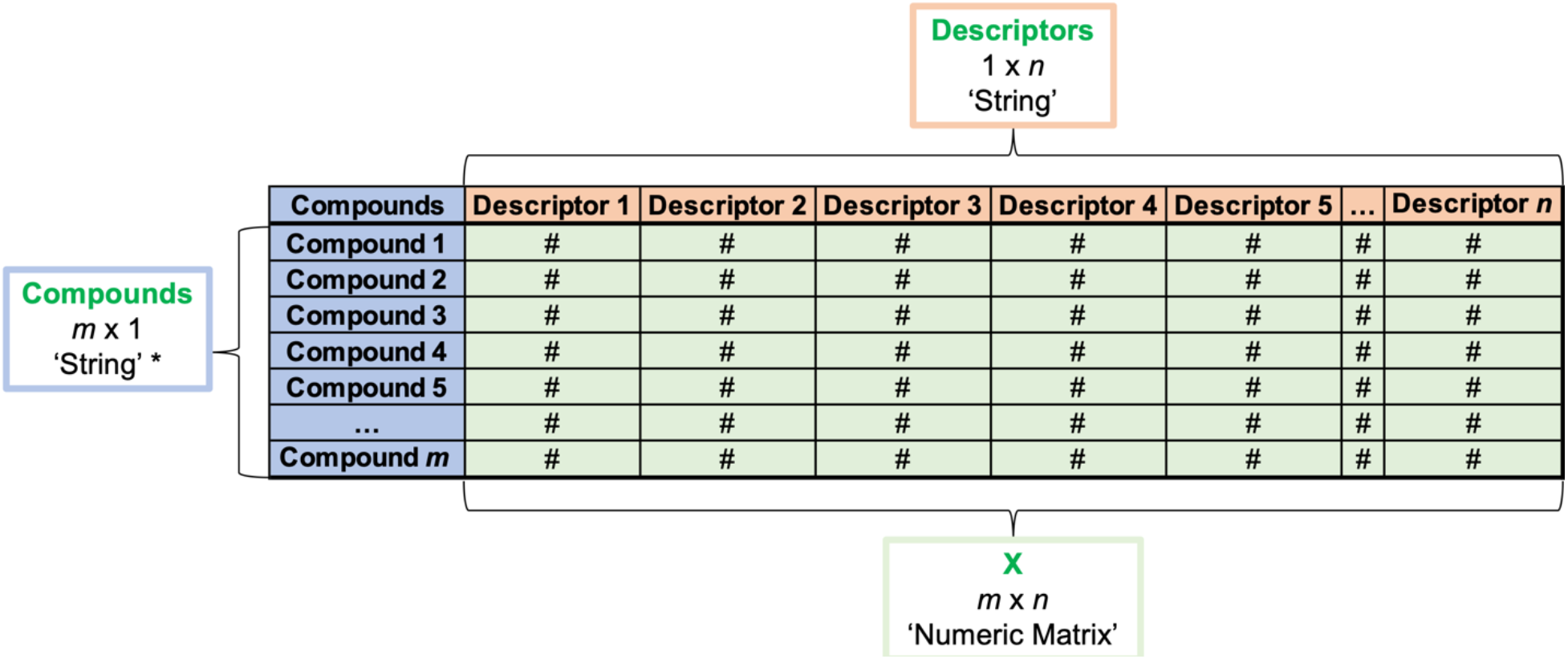
Example of a data set to be used for clustering. Each row represents a different item in the set (in this case, a different chemical compound), while each column represents a different property or characteristic of the item (for chemical compounds, these are often called molecular ‘descriptors’). Each cell in the matrix contains a number that is the value of a given property for a given compound. The numerical values, descriptor names, and compound names are imported into MATLAB separately, to give MATLAB data variables of the types and dimensions shown. *For Basic Protocol 1b, if the compound names take the form of ID numbers, they must be imported as a numeric matrix.

When the input data are considered in terms of the position of each item on this set of *n* axes, it is self-evident that the data points representing items with nearly similar values for all of the properties will tend to lie close together in this *n*-dimensional property space, while items with highly dissimilar property values will be far apart. In general, clustering algorithms work by calculating the pairwise distances between each of the data points (items) in this *n*-dimensional property space, and then grouping the items in terms of which are close to which. Additional factors that determine the outcome of clustering are therefore how the pairwise distance in *n*-dimensional space is quantified, and how the distance between two groups of points is defined (called the ‘linkage’).

#### Distance metrics

##### Euclidean distance

A commonly used measure of the distance between two data points in *n*-dimensional space is simply the straight-line distance. For example, the distance between two data points, (1) and (2), in two-dimensional *x*-*y* space is the hypotenuse of a right-angled triangle where one of the short sides is parallel to the *x*-axis and the other is parallel to the *y*-axis (**Figure 3**). Using the Pythagorean Theorem this distance can be calculated as *d* = sqrt([*x*_*2*_*-x*_*1*_]^2^ + [*y*_*2*_*-y*_*1*_]^2^). The straight line distance between two points, *i* and *j*, in *n*-dimensional space can similarly be calculated as *d* = Sqrt(SUM[*x*_*i*_*-x*_*j*_]^2^) for *x* = 1 to *n*. This straight-line distance is commonly termed the ‘Euclidean distance’.

##### Manhattan distance

When dealing with very large data sets, the computational time required to calculate all the pairwise Euclidean distances can become large. An alternative way of defining the distance between two points that is computationally cheaper to calculate is the so-called ‘Manhattan’ or ‘City block’ distance. Using this metric, the distance between the points is simply defined as the differences between the coordinate values on each axis summed over all axes, as if one were traveling from one point to the other but could only move parallel to a given axis and cannot cut across diagonally (hence the analogy with navigating around Manhattan).

There is no universal right answer as to which distance metric to use in clustering. Indeed, other distance metrics beyond the two mentioned here can also be used. Generally speaking, items with similar properties will appear close together in *n*-dimensional space using either of the above metrics, compared to items with property values that are very different. In our own work we tend to use Euclidean distance as our default, due to its conceptual simplicity, and only consider Manhattan distance for very large data sets if the computation were to became inconveniently slow.

##### Linkage methods

Clustering involves decisions about how groups of data points might be further grouped to generate larger superclusters. Such decisions require a method for defining the distance between two groups of data points. One approach would be to calculate the ‘center of mass’ (centroid) of each cluster, and use the distance between these centroids as a measure of how closely the two clusters are related. This approach is called ‘average linkage’. Alternatively, one might define the distance between two clusters as the distance separating the single pair of data points, one from each cluster, that lie closest together in *n*-dimensional space. This closest neighbor approach is called ‘single linkage’. In contrast, the distance between two clusters might be defined as the distance separating the most distant individual data points from each cluster. Calculating the separation in this manner, which gives a distance that fully encompasses all members of both clusters, is called ‘complete linkage’. Different linkage methods tend to give clusters with different characteristics, as summarized below in Table 2.

As was the case with the different distance metrics, there is no universal right answer to what is the best linkage to use when clustering; the choice may be dictated by the goal of the clustering analysis, but is often somewhat arbitrary. If in doubt, the best thing is to test the results of different choices of distance metric and linkage, and see which results best reveal the patterns and relationships in your data that you are interested in.

There are many conceptually distinct approaches to clustering. For the purposes of this article we will restrict ourselves to two, ‘hierarchical clustering’ and ‘*k*-medoids clustering’.

***Hierarchical Clustering*** gathers the elements of the data set into pairs or small groups that have similar properties (i.e. lie close together in *n*-dimensional property space). It then relates these groups to other nearby groups or elements to form larger groups, then further clusters these larger groups together with their nearest neighbors into even larger groups, and so on until all of the items are in a single group (**Figure 1.2A**). The calculation of which items or groups are close enough together to be grouped, at any given step of this process, is done using your chosen distance metric and linkage. The above process can be done either by starting with the individual items and iteratively grouping them together based on proximity, as described above (‘agglomerative clustering’), or by starting with the full set of items and iteratively dividing and subdividing them until each point is in its own group (‘divisive clustering’). In either case, the result is organization of the items into a hierarchical set of groups and subgroups that can be represented as a tree-like structure called a ‘dendrogram’ (**Figure 1.2B**). In the dendrogram, at the lowest level of grouping each individual item occupies its own separate group, called a ‘leaf’, and the combination of items or groups together at ‘nodes’ creates ‘branches’ that further combine to ultimately encompass all items in the set.

**Figure 1.2.**
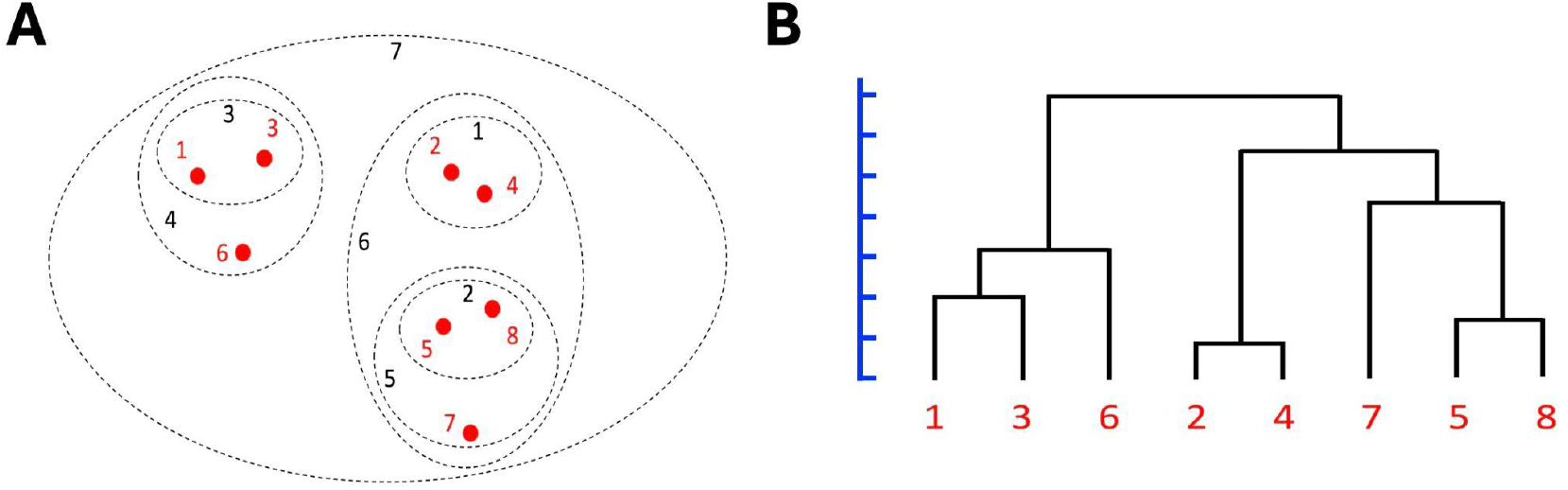
Illustration of how items can be hierarchically clustered based on their proximities in property space. **(A)** An input set of 8 items (numbered red dots), graphed so as to represent their relative positions in some hypothetical property space. If using an agglomerative clustering algorithm, step 1 (black numeral 1) is to identify the closest two data points, 2 and 4, and combine them into a single group. Items 5 and 8 (step 2), and items 1 and 3 (step 3), are also quite close neighbors and so are similarly grouped as pairs. Increasing the distance threshold further established item 6 as a neighbor of the group (1,3) (step 4), and so these three items become grouped together, and so on until all items are included in a single group. **(B)** The results of the clustering in (A) represented as a dendrogram, in which the height of each node indicates the distance in separating the relevant items or groups in (A). A distance scale is included (blue) to highlight that the branch lengths have quantitative meaning, though an explicit scale is often omitted in practice.

In interpreting the results of hierarchical clustering, two key principles apply. (i) The length of a given branch is not arbitrary, but rather indicates the distance between the connected items or groups in the original *n*-dimensional feature space. Specifically, the position of a given node with respect to the distance axis indicates the separation (using the chosen distance metric) between the items or groups that the node connects. Thus, for example, in **Figure 1.2B** the node connecting items 1 and 3 is roughly 2-fold higher on the distance scale than the node that connects items 2 and 4, indicating that items 1 and 3 are roughly 2-fold further apart in the original *n*-dimensional feature space than are items 2 and 4 (see **Figure 1.2A**). Similarly, the node that connects the group (2,4) to the group (7,5,8) is quite high on the diagram, indicating that these two groups are quite far apart in feature space. (ii) The horizontal arrangement of the branches is arbitrary. Thus, even though in **Figure 1.2B** the leaf containing item 6 is closer to that containing item 3 than to the leaf containing item 1, this does not imply that items 3 and 6 are closer together in the original feature space. A useful way to think about this is to consider the dendrogram as if it were a child’s mobile, in which each node is a pivot point that can freely swing around horizontally. Thus, a dendrogram in which the leaves containing items 1 and 3 could freely interchange to create an equally valid dendrogram. Thus, it is important to resist the temptation to assume that the dendrogram in **Figure 1.2B** implies that items 1 and 8 have the greatest mutual distance in the set as, though a 180° rotation about two of the nodes, an equally valid dendrogram could be drawn that would have these two items as immediate neighbors. For a dendrogram oriented as shown in **Figure 1.2B**, distance information is contained only in the vertical axis; the arrangement of the leaves and branches in the horizontal direction is largely arbitrary and contains no information about the relatedness of the elements concerned. The fact that the dendrogram, by itself, does not tell us which individual member of any given group, e.g. (1,3), is closest to any other item or group, such as item 6 illustrates that some information is lost upon clustering, for the gain of achieving a simple two-dimensional representation of the data.

Importantly, dendrograms can be shown in a variety of equivalent orientations and shapes. For example, the dendrogram in Figure 1.2B could equivalently be portrayed horizontally, with the branches horizontal and the leaves at the extreme left or right. Triangular and circular forms are also commonly encountered. In interpreting a dendrogram, the important things are to establish the orientation of the axis that represents distance, which may be vertical, horizontal, or from the center to the periphery, and to understand that the diagram can rotate at any node and thus the proximity of the leaves in the diagram do not indicate anything about the degree of relatedness of the corresponding items. A set of excellent puzzles and exercises on how to interpret dendrograms has been published (Baum, 2008) (https://www.ebi.ac.uk/sites/ebi.ac.uk/files/content.ebi.ac.uk/materials/2014/140602_prague/tree_thinking_tests.pdf), which we recommend highly.

***k-means and k-medoids clustering*** are related methods for subdividing a set of items into some predefined number (*k*) of groups based on their similarity with respect to a set of characteristics or properties. The method of *k*-means clustering divides a set of items into *k* groups, where *k* is pre-selected by the user as the desired number of groups, based on their proximity in *n*-dimensional property space, and outputs the mean values of the properties of the items that make up each group. *k*-medoids clustering similarly divides a set of items into *k* groups based on their similarity with respect to a set of properties, but while *k*-means clustering outputs average values of the properties that are characteristic of each group, *k*-medoids clustering identifies the specific item from each group that has properties closest to the average properties for that group.

Hierarchical clustering and *k*-means or *k*-medoid clustering all share the characteristic that the relationships the clustering reveals depends strongly on the properties or characteristics used for the analysis. For example, clustering a set of humans based on height, versus based on age, or number of books read, etc. will likely give very different results; any pair of individuals might appear very similar, and be clustered together, or very different, depending on the characteristic(s) used to describe them in the clustering exercise. Thus, for example, when using *k*-medoids clustering to identify a representative subset of compounds from a larger collection, the result will depend very strongly on which molecular properties of the compounds are used as input data, and generally much less strongly on operational decisions such as distance metric or linkage method.

A key difference between hierarchical clustering and *k*-means or *k*-medoid clustering is that, for the former, it is not necessary to choose beforehand how many groups are required. In hierarchical clustering, the resulting dendrogram (**Figure 1.2B**) can be sliced perpendicularly at any desired level to separate the items into the desired number of groups or to give a desired average group size or similarity.

## BASIC PROTOCOL 1a: Clustering to create a dendrogram

### Purpose of method

To subdivide a set of elements, based on the characteristics of each element with respect to a defined list of properties, and represent the results in the form of a tree diagram (‘dendrogram’) that graphically illustrates which elements in the set are more closely or less closely related to which others. This method is used for problems in which relatedness is usefully illustrated in the form of a ‘family tree’. For example, a common use is to compare a set of homologous protein or DNA sequences to generate a tree diagram illustrating the evolutionary relatedness of their organisms of origin.

### Background

The method takes as input an *m* x *n* matrix in which each row represents a different element of the data set (in our case, a different compound), and each column represents a different property that describes the elements. Each cell in the matrix contains the numerical values of a particular property for a particular element. The numerical values of the *n* properties can be considered as coordinates that define the location of each element in a notional *n*-dimensional property space. The degree of relatedness between any two objects can be quantified in terms of the distance between their positions in this *n*-dimensional space, defined using whichever distance metric the investigator chooses (see Protocol 1 Introduction). The closer any two elements are, the more similar they are considered to be with respect to the set of properties included in the model. In the clustering method described here, the distances between the elements in the set are represented as a dendrogram, as illustrated in **Figure 1.2B**. This type of clustering is designed to reveal the intrinsic structure of the data – that is, how each element relates to the others – without any preconceived idea of how many groups or clusters might be expected. In this sense, it differs from the techniques of *k*-means and *k*-medoids clustering, which instead perform the task of subdividing the elements into a pre-determined number of clusters. Instead, with the method described here the output dendrogram can be assessed at any desired distance threshold to achieve a desired number of clusters or average cluster size.

The results of this clustering method will depend strongly on the properties or characteristics used for the analysis and how they are scaled, as described in the Introduction. The particular set of elements included in the clustering will also affect the results, in the sense that clustering different randomly selected subsets of elements from the same parent set may give dendrograms that look somewhat different. The results will generally depend less strongly on operational decisions such as distance metric or linkage method.

#### Before you start

Be sure you have installed the necessary toolboxes mentioned in the ‘REQUIRED SOFTWARE’ section. If using your own dataset, make sure you have formatted the data for this analysis as described in the ‘DATA DOWNLOAD AND PREPARATION’ section.

##### 1. Gather the input data

Details of the data download and preparation are given above in Section IV, and spreadsheets containing the data required to perform the protocols are included in the supplementary file ‘Data_Clustering_and_PCA.xlsx’. For this protocol, these comprise a set of 201 data elements (compounds), a set of 139 molecular features (descriptors), and the numerical values of each descriptor for each compound, provided here in an Excel spreadsheet, on a single page with no gaps or breaks between cells. The supplementary file ‘Data_Clustering_and_PCA.xlsx’ contains the data already formatted in this way, in the sheet named ‘data_Clustering_PCA’.

*If you are using your own data rather than the supplementary file provided, be sure to format your external data files as described in ‘Section IV. DATA DOWNLOAD AND PREPARATION, and to change the external filenames (blue text) in the following steps to match the names you have assigned to your files*.

##### 2. Identify the file that contains the data to be imported into MATLAB

i. Starting with a clear workspace, click on the Import Data button in the MATLAB Home tab (top of screen), navigate to where you have saved the Supplementary File ‘Data_Clustering_and_PCA.xlsx’, and selecting the tab labeled ‘data_Clustering_PCA’ at the bottom of the Data Import window. Be sure you are on the correct sheet before you begin importing data.

##### 3. Import the numeric values that make up the body of the input data

i. In your open Data Import window, select the cells that contain the numerical data. Only the descriptor values (area shaded green in ‘data_Clustering_PCA’) should be selected for import in this step; <u>do not select the column headers in row 1 or the compound names in column 1</u>. The selected data should comprise 201 rows (reflecting the 201 compounds in the data set) and 139 columns (reflecting the 139 molecular descriptors we are using for this analysis), and should include only numbers.
ii. Go to ‘Output Type’ at the top of the Data Import window and select ‘Numeric Matrix’ from the drop-down menu, then go to the Import Selection button at the top right of the Import Data window and click the green check mark (or select ‘Import Data’ from the drop-down menu). You should see a new data object with dimensions 201×139 appear in the Workspace window of your MATLAB desktop (you may need to move the Import Data window aside to see it).
iii. Rename the 201 × 139 workspace variable. For the purpose of this exercise, you should call the new workspace variable **X**. You can do this in the open Data Import window by double clicking on the default name (same as the original Excel filename) that appears just below the top row of the window and typing in the desired new name. Alternatively, you can click on the newly created workspace variable in the MATLAB Workspace window to highlight its default name, and then type in the new name, **X**.

##### 4. Import the column headings (descriptor names) and row headings (compound identifiers)

In the Data Import window, select the cells in the top row that contain the column headings (*i.e*. the names of the 139 molecular descriptors). <u>Do not include cell A1, as the contents are not a descriptor name</u>. At the top of the Data Import window change the ‘Output Type’ to ‘String Array’, type in the box just above row 1 to assign the name **Descriptors** to the new data object, then click on ‘Import Data’. A new data object with this name, of size 1 × 139, will appear in your Workspace window. To import the compound identifiers, select the cells in the first column that contain the compound names. <u>Again, do not include cell A1, as this is not a compound name</u>. Change the ‘Output Type’ to ‘String Array’, assign the name **Compounds** to the new data object, then click on ‘Import Data’. A new data object with this name, of size 201 × 1, will appear in your Workspace window.

> *Data objects created in MATLAB can be renamed after the fact, as described in Step 3(iii), but it is good practice to assign the desired name in the Data Import window before creating the object. The reason is that the default name that MATLAB assigns to new data objects is the name of the external file from which the data were imported, making it easy to accidentally over-write a different data object previously imported from the same external file if it was not assigned a unique name upon creation*.

##### 5. Save the MATLAB workspace that now contains the imported data as a .mat file

*This file can be reopened to give a MATLAB workspace identical to the state that was saved*.

##### 6. Make a box plot of the input descriptor values

> *Making a boxplot allows you to look at the distribution of values for each of the properties in the data set. The y-axis shows the names or numerical identifiers of each of the properties included in the analysis (i.e. the 139 molecular descriptor), while the x-axis shows the distribution of numerical values for each property for all items in the data set (i.e. the compounds). The distribution of values for each property is shown as a horizontal ‘box and whisker’ plot beside the descriptor name. Note that, because the values are all plotted using a common x-axis scale, the distribution of values for properties that vary over only a small numerical range will appear compressed in the horizontal direction compared to the distribution for properties that span a larger proportion of the x-axis scale*.

Type or copy the following text into the Command window, to the right of the prompt (>>), pressing the Return key at the end of each line:

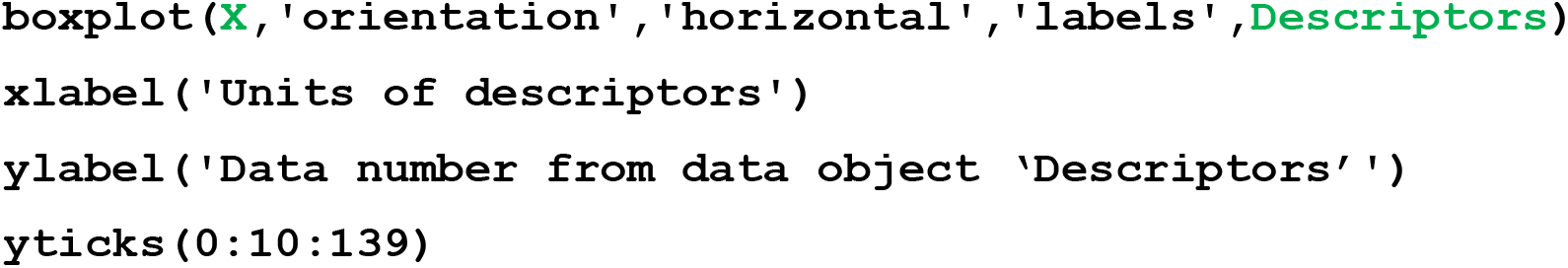
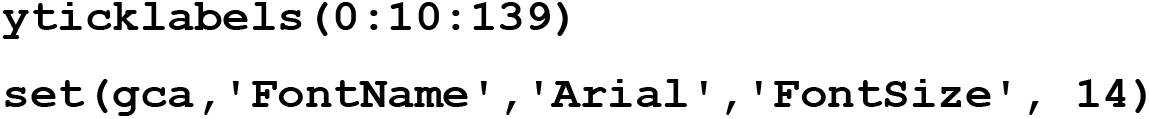

A plot resembling **Figure 1.3** will appear.

**Figure 1.3:**
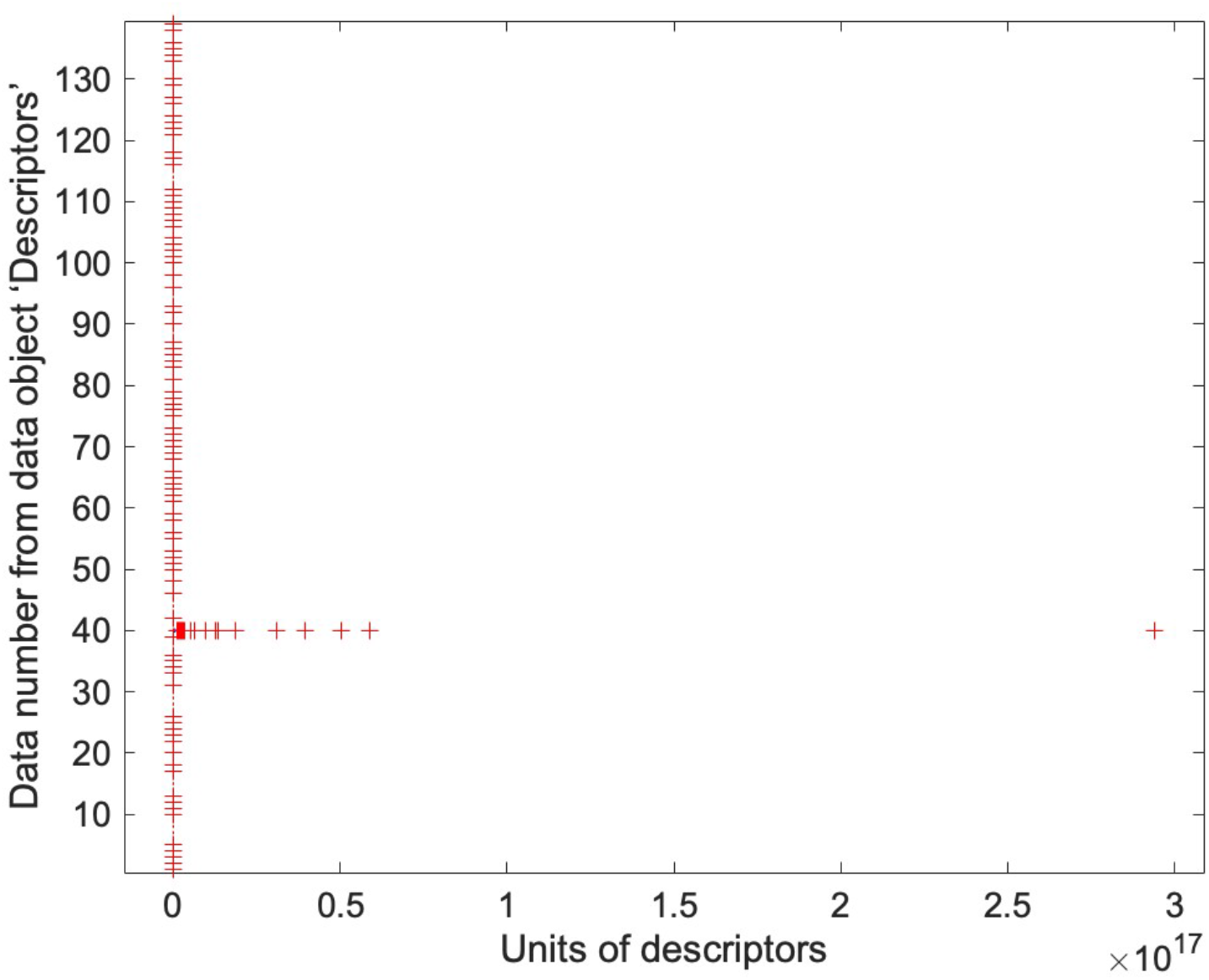
Box plot showing the distribution of numerical values of the 201 compounds in the data set (*x*-axis) with respect to each of the 139 molecular descriptors (*y*-axis). To avoid the crowding of descriptor names as labels on the *y*-axis, for visualization purposes we have substituted the descriptor names with numerical values from 1-139 indicating their order from left to right in the original Excel data sheet.

Save the box plot by going to the File menu <u>in the Figure window</u> and selecting ‘Save’ and the file format ‘MATLAB figure (*.fig)’. This will save the file as a MATLAB figure file that can be edited later.

> *As described in the Figure caption, the plot generated by the command shown above will show descriptor names to the left rather than numbers. To create a plot that looks exactly like **Figure 1.3**, the above command must be modified so that the MATLAB workspace variable ‘****Descriptors****’, which contains a list of the descriptor names, is replaced with the name of a different variable, which one might name e.g. ‘****Descriptor_Numbers****’, that comprises a 1 × 139 vector containing the numbers from 1-139. This new variable must be separately created and named before it can be called on in a command*.
>
> *As formatted using the commands shown here, the box plot shows the 25th-75th percentile range of the values present for each property as a horizontal box with a central mark to indicate the median value and whiskers extending to the left and right to indicate the low and high extremes of the non-outlier data points*.
>
> *Individual values that the algorithm considers to be outliers from the distribution are plotted as* x *symbols. These features are better seen in* ***Figure 1.4***, *where the descriptor ranges have been normalized*.
>
> *The boxplot in* ***Figure 1.3*** *shows that, for the data we are using here, the values of one descriptor (number 40) span a far larger numerical range (from zero to 3 × 10*^*17*^*) than any other descriptor, such that the value ranges for all of the other descriptors appear compressed to near zero. Consequently, except for highlighting this single fact, the box plot for this data set is less informative than is typically the case. The numerical scales for the different descriptors can be normalized to a common scale by ‘z-scoring’ the data, as described in Step 7, below*.

##### 7. Z-score (scale and center) the X data values

> *When input data values have very different numerical scales, as is the case here for descriptor 40 versus the other descriptors, this can be problematic for clustering and other unsupervised ML methods. The distances between the items in n-dimensional property space (i.e. the overall numerical differences between the compounds) will be dominated by differences in the property values having the largest numerical variance (largest range), giving these descriptors an outsized influence compared to other properties that could be as or more important functionally. In such cases, it can be useful to normalize the data to make the numerical ranges more comparable. One commonly used approach for doing this is called ‘z-scoring’. When data are z-scored, the mean value of the numbers in each column is subtracted from all values in the column, and the results are then divided by the standard deviation of the values in that column*.
>
>
> 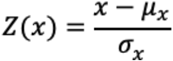
>
>
> *The result is to center the values in each column on zero and scale the variance to a standard deviation (SD) of exactly 1. A discussion of the pros and cons of z-scoring, and when it is appropriate for a particular type of analysis, is provided in the Commentary section (Section VI). But some manner of scaling is clearly required for the data we are using here, and so we will z-score the data to achieve this end*.

Z-score the data in **X** to normalize the descriptor values in each column to have mean = 0 and SD = 1, by entering following the command in the Command window:

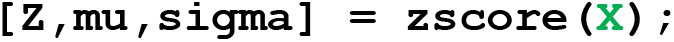

> *Three new variables will be created in the MATLAB Workspace window, called ‘Z’, ‘mu’, and ‘sigma’. ‘Z’ is an m x n matrix (in this case 201 × 139) that contains the descriptor values from* ***X*** *normalized to each have mean = 0 and SD = 1. The variables ‘mu’ and ‘sigma’ are 139 × 1 vectors that contain, respectively, the mean and the SD values of each of the 139 columns in the original data matrix* ***X***.
>
> *Variations of the* ***zscore*** *command can be used to normalize across the rows of* ***X*** *as well as, or instead, of down the columns. Type ‘****help zscore****’ into the MATLAB Command window for further details*.

###### Important

> *Commands such as z-score can sometimes generate results for some positions in the answer matrix that appear as ‘NaN’ (‘Not a Number’). ‘NaN’ entries result from invalid mathematical operations such as division by zero, and their presence can prevent the proper operation of some subsequent commands. If you are using the data sets provided with this article, no NaN values will be generated. However, if you are following this protocol with your own data set, for which NaN results could occur, ‘NaN’ results can be identified and replaced by zeroes with the following command:* `Z(isnan(Z)) = 0`, *or, more generally*, `variable_name(isnan(variable_name)) = 0;`.

##### 8. Make a box plot of the normalized descriptor values

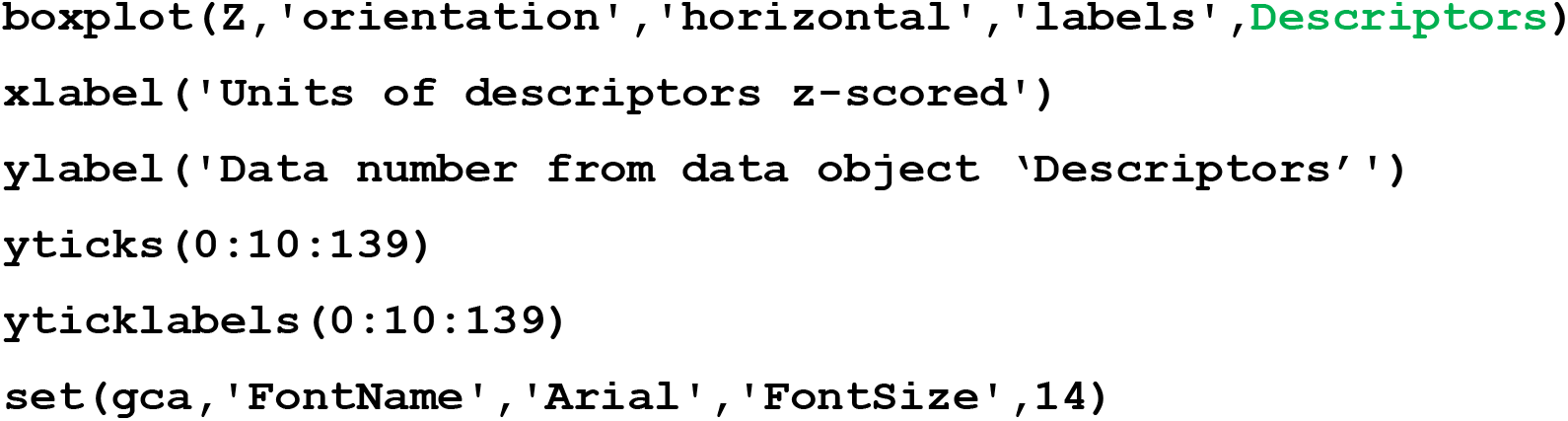

> *Note: the above command is identical to that used in Step 6, except specifying ‘Z’ rather than* ***X*** *as the input data for the ‘boxplot’ command*.

A new Figure window will appear containing a plot that resembles **Figure 1.4**.

**Figure 1.4.**
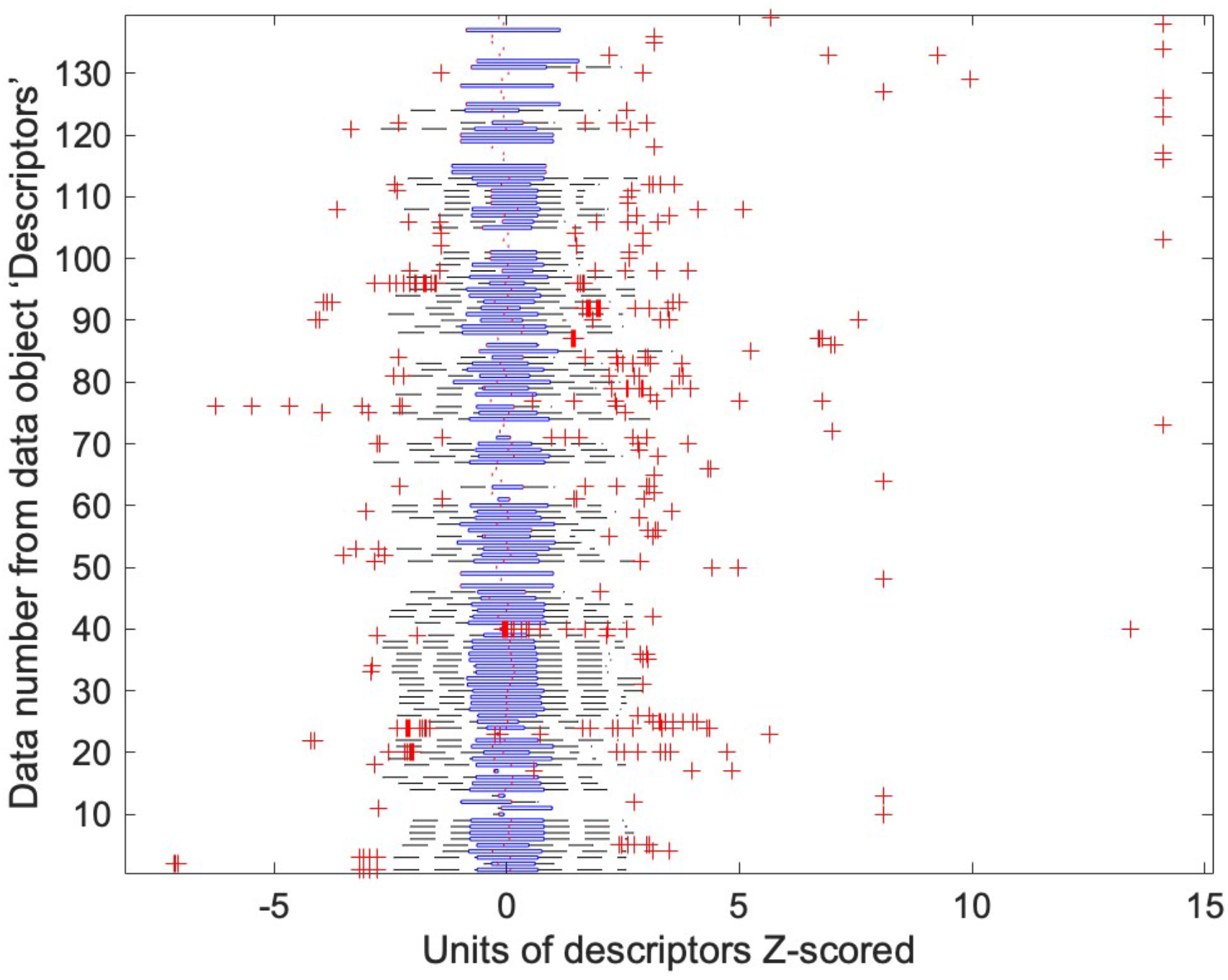
Box plot showing the distribution of numerical values of the 201 compounds in the data set (*x*-axis) with respect to each of the 139 molecular descriptors (*y*-axis). To avoid the crowding of descriptor names as labels on the y-axis, for visualization purposes we have substituted the descriptor names with numerical values from 1-139 indicating their order from left to right in the original excel sheet (see note under Step 5 for explanation). Note that, after z-scoring, all of the descriptors show values distributed around a mean of zero, with roughly similar interquartile ranges due to SD having been fixed at unity.

##### 9. Perform hierarchical clustering on the unscaled data in X

> *We will use the ‘linkage’ command, choosing average as our linkage method and Euclidean distance as our distance metric (see Background for explanations of these terms)*.

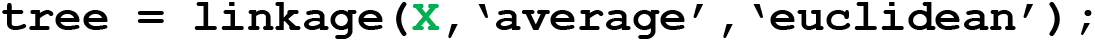

A new variable, which we have chosen to call ‘tree’, will appear in the Workspace window.

> *The variable ‘tree’ contains three columns and* m*-1 rows, where* m *is the number of items in the set (in our case, 201). This matrix contains the results of the hierarchical clustering. Specifically, each row of columns 1 and 2 contains the numerical identifier of a different item or clustered group of items from the data set, while the corresponding row of column 3 contains the distance separating this pair of items/clusters in* n*-dimensional property space. Individual items are assigned numbers from 1 to* m, *and each pair or group of items is assigned a number in the order they are identified, starting at* m*+1*.
>
> *The scale on which the distance between items/groups is expressed corresponds to the distance in* n*-dimensional space. Consequently, if the original descriptor values are expressed in units that result in large numerical differences between items, the separation between items and groups in ‘tree’ will tend to have large numerical values. If the data have been z-scored prior to clustering, most data points will lie within 1-2 units of the center of the property space, and so the distances in ‘tree’ will mostly have single-digit values*.
>
> *In the ‘linkage’ command, single linkage is the default linkage type and Euclidean distance is the default distance metric. Thus, in this example, specifying ‘average’ is required as part of the command, but it is not necessary to specify ‘euclidean’ which could be omitted from the command text. To view the different linkage methods, distance metrics, and other options for ‘linkage’, type ‘Doc linkage’ in the Command window of MATLAB*.

##### 10. Plot the results of the clustering of the original, unscaled data X as a dendrogram

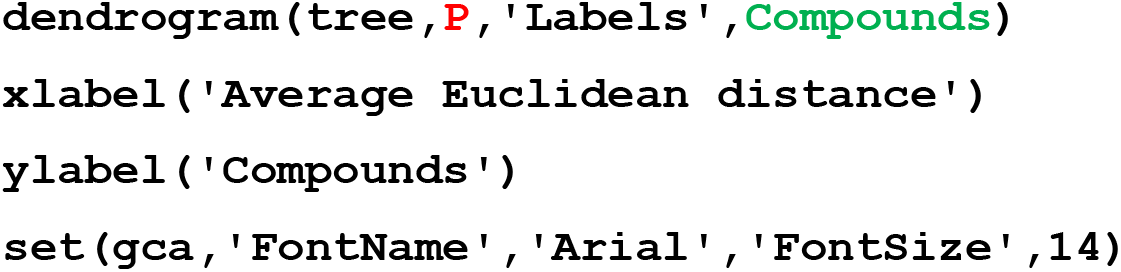

> *In this command*, ***P*** *is a user-defined parameter that specifies the maximum number of ‘leaves’ (i.e. terminal points of branches) to be shown in the plot. In this instance, we set* ***P*** *= 0*.
>
> *Setting* ***P*** *= 0 instructs that all possible leaves should be plotted, corresponding to one leaf for each element in the data set. Entering a value for* ***P*** *between 0 and 201 (i.e. the total number of compounds in our set) would plot a ‘pruned’ version of the dendrogram truncated to show only the number of leaves specified*. ***P*** *can range from 2 up to the total number of elements*, m, *in the data set (in our case, 201). Setting* ***P*** *=* m *is equivalent to setting P = 0, as both instruct that no pruning should be done. If* ***P*** *is set to a value greater than zero and less than* m, *a given leaf in the dendrogram may represent a group rather than an individual element. Although decreasing the number of branches truncates the amount of data shown, the resulting simplification of the plot can aid visualization of the relationships between items within the data set*.

A new Figure window will appear that contains the results in the data object ‘tree’ plotted as a dendrogram (**Figure 1.5**).

**Figure 1.5:**
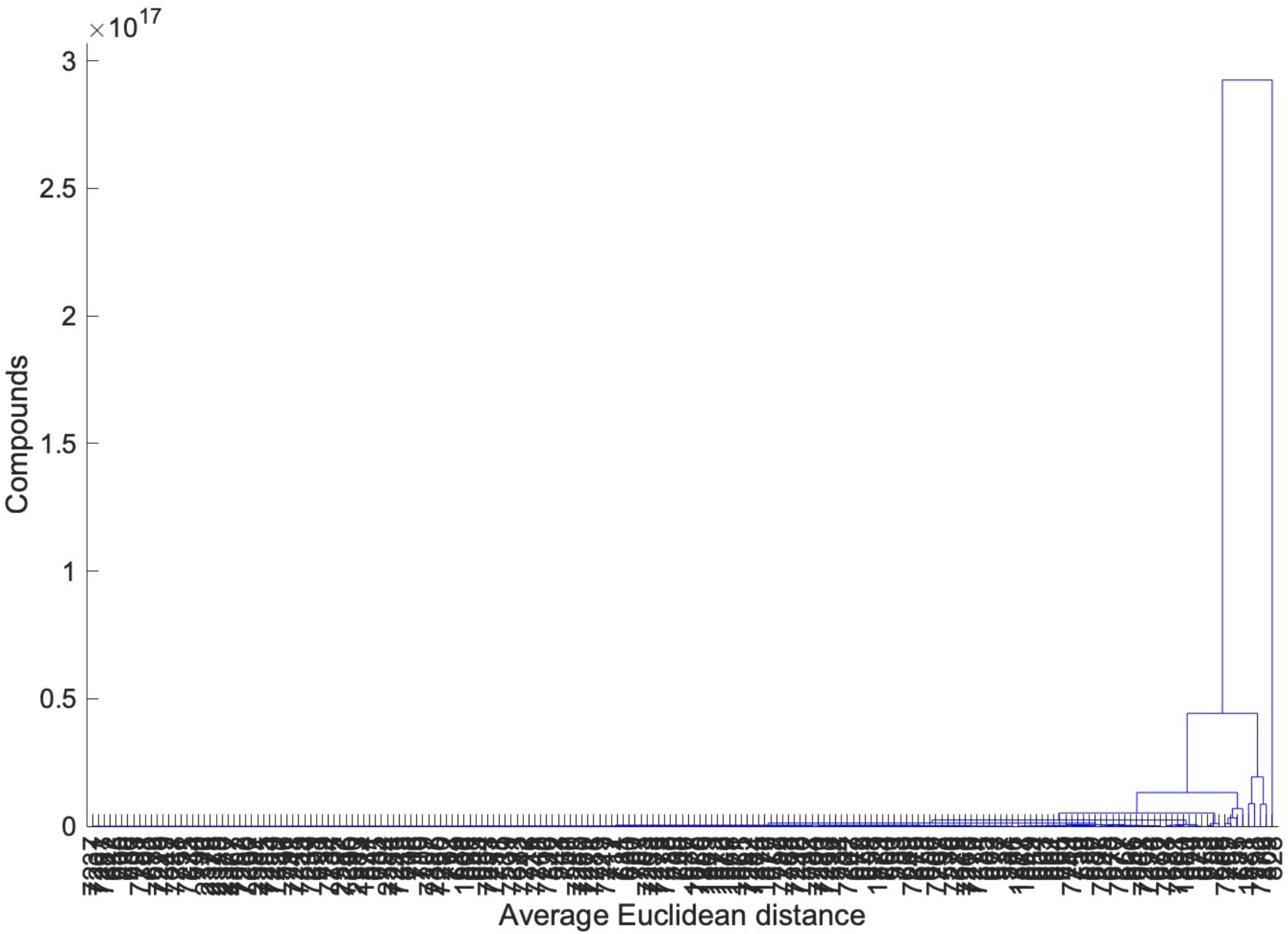
Dendrogram showing how the compounds cluster with respect to their values for the properties contained in **X**. The distance scale represents the Euclidean distance between any given pair of objects or clusters of objects in *n*-dimensional space, expressed in the original units used for the descriptor values in **X**. The horizontal lines connecting pairs of leaves or clusters are plotted at a height corresponding to the distance separating that pair of elements, the distance to be read off the distance (vertical) axis. In this particular case, one descriptor has a value range very much greater than any of the other descriptors, so that clusters defined by differences in the other descriptors are all compressed near zero on the distance scale. See Figure 1.4 to see how the data cluster if the descriptor values are first z-scored to normalize all their value ranges to SD = 1.

> *Due to the large number of leaves (201) in this dendrogram, the compound names overlap extensively and consequently are impossible to read. This problem can be solved by choosing a value of* ***P*** *that is >0 but substantially less than 201, to reduce the number of leaves and thus allow more room for each label, though with some loss of information as each leaf will no longer represent a single compound*.

##### 11. Cluster the z-scored data Z and plot the results as a dendrogram

> *The clustering done using the unscaled data in Step 9 results in a dendrogram (**Figure 1.5**) that is relatively uninformative, because the separations between the compounds in* n*-dimensional space are dominated by the value of a single descriptor that has a very large value range. One approach to solving this problem, with a view to better revealing the relatedness between the compounds, is to instead cluster the z-scored data contained in the variable Z that was created in Step 7 (**Figure 1.6**)*.

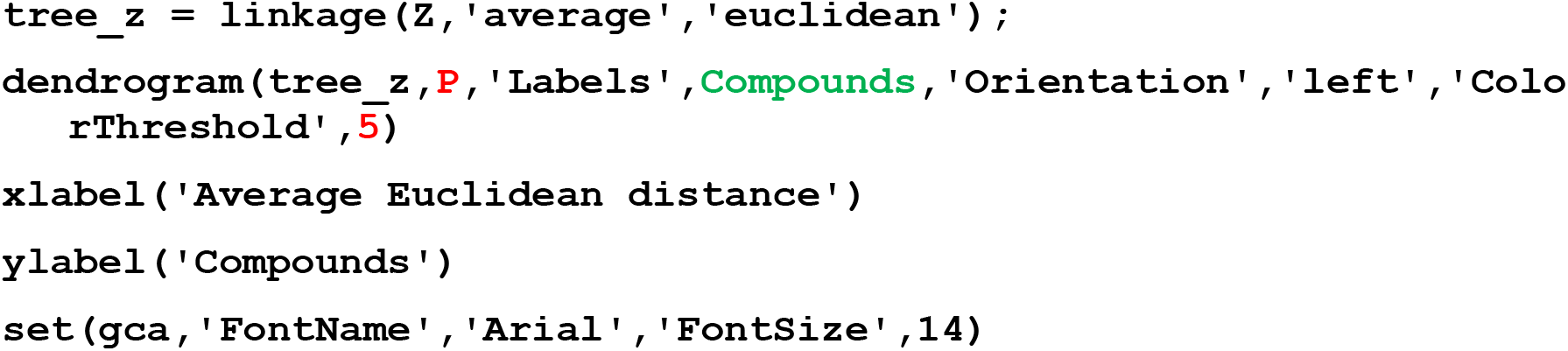

> *The new data variable created by this command has been named ‘tree_z’ to distinguish it from the variable ‘tree’ that was created in Step 8 using the unnormalized input data*.
>
> *The reader will note that, in the above, we have added a few more commands, to modify the format of the dendrogram:*

i. *The ‘Orientation’ command was used to change the orientation of the dendrogram from horizontal to vertical. Options for this command are ‘top’ (the default, as seen in **Figure 1.5**), ‘bottom’, ‘left’, and ‘right’*.
ii. *The ‘ColorThreshold’ command instructs the program to color the leaves/clusters to highlight groups of related items as well as outliers from those groups. The number* ***5*** *is the value we chose for a user-defined parameter*, ***T***, *that sets the threshold such that any leaves or clusters that are separated by <****T*** *units in n-dimensional space should be colored alike. If the ‘ColorThreshold’ command is omitted, or if a value for T is chosen that lies outside the range from zero to the maximum distance in ‘tree’, the dendrogram will have only one color*.

*For a full description of available formatting options, type ‘Doc linkage’ into the Command window. The dendrogram can also be modified using the ‘edit plot’ function in MATLAB. For further details see: https://www.mathworks.com/help/MATLAB/ref/plotedit.html*

**Figure 1.6:**
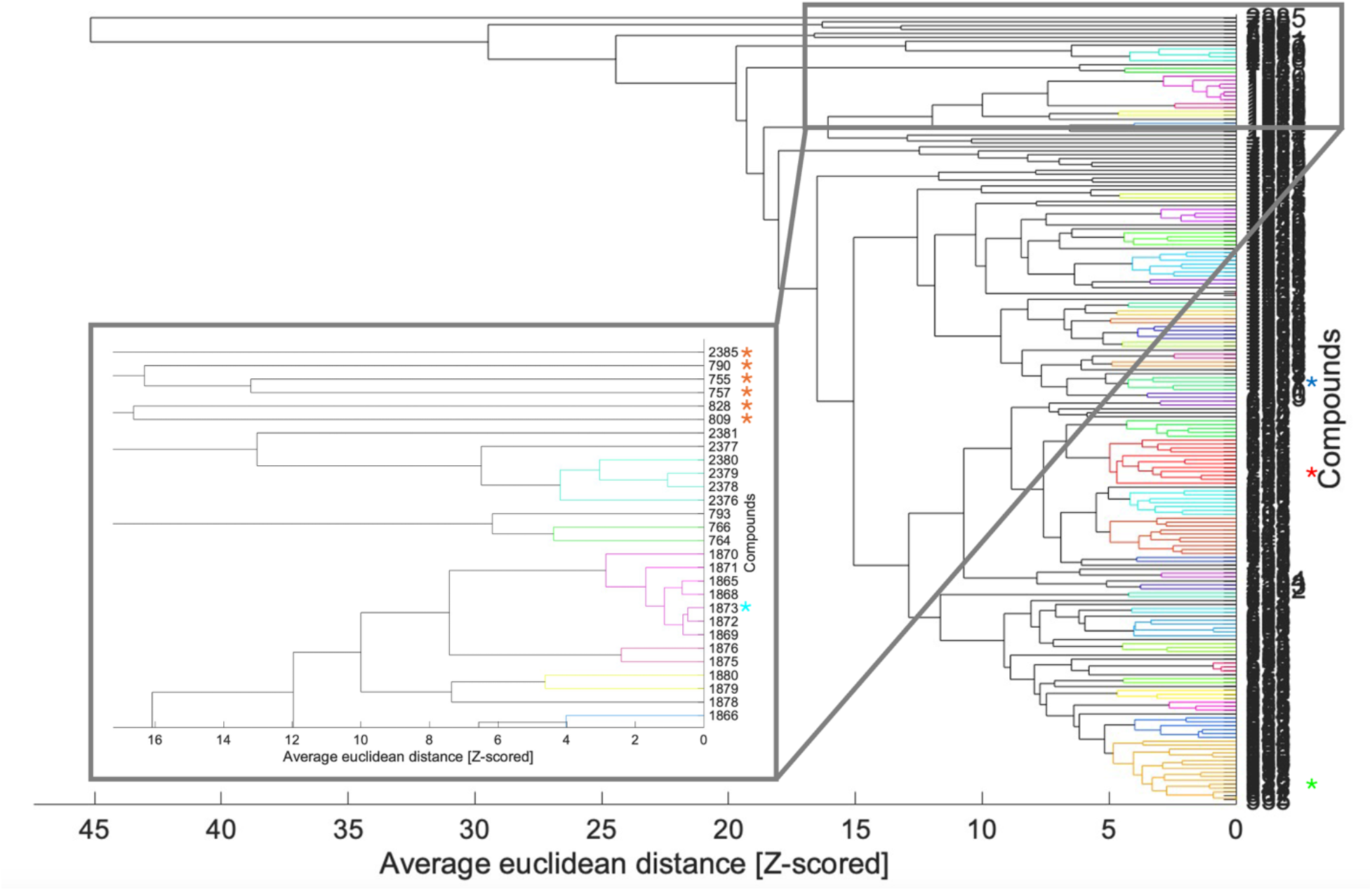
Dendrogram showing how the compounds cluster with respect to their z-scored values for the properties, contained in ‘Z’. The distance scale represents the Euclidean distance between any given pair of objects or clusters of objects in *n*-dimensional space, where 1 unit equals the standard deviation of the values in any single property. The leaves are colored such that compounds/clusters of compounds that lie within <5 units of each other in *n*-dimensional property space are colored alike. The vertical lines connecting pairs of leaves/clusters are plotted at a position corresponding to the distance separating that pair of elements, this distance to be read off the distance (horizontal) axis. The inset plot shows an expanded view of the first 30 leaves, illustrating how each leaf corresponds to a single compound in the data set. The red asterisks in the inset plot identify the six compounds that the dendrogram identifies as most extreme outliers, as defined by Euclidean distance from their neighbors.

***Figure 1.6*** *provides a two-dimensional representation of how the compounds cluster in the 139-dimensional property space. Recalling that it is the position of a branch origin on the* y*-axis that indicates the distances between the objects in the branches and leaves that originate at the point, we can see that compound ID2385 is by far the biggest outlier, being at the greatest distance from any other compound. Compounds 790, 755 and 757 are, as a group, also quite far from the remaining compounds, with compounds 828 and 829 being the next furthest from the bulk of compounds. These compounds are discussed further in later sections*.

##### 12. Map the elements in the data set to the leaves of the dendrogram

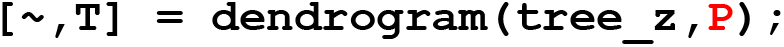

> ***Important:*** *use the same value for P that was used when creating the dendrogram of interest*.
>
> *The variable, ‘T’, generated by this command is a matrix of size* m *x 1 that identifies which of the* ***P*** *leaves in the dendrogram each data element (compound) belongs to*.
>
> *The leaves in the dendrogram are numbered from 1 to* ***P***, *working from left to right for horizontally oriented dendrograms and from the bottom up for vertically oriented ones. The first number in ‘T’ is the number of the leaf that contains the compound from Row 1 of the data object used in the ‘linkage’ command (in this case, ‘Z’). The second number in ‘T’ is the leaf number that contains the compound from row 2 of ‘Z’, and so on. ‘T’ thus contains a total of* m *numbers (one for each compound), with the numbers in ‘T’ having values between 1 and* ***P*** *(the total number of leaves in the dendrogram). If the dendrogram is not truncated (i.e*. ***P*** *=* m *or, equivalently*, ***P*** *= 0), every compound corresponds to a different leaf, and so every number in ‘T’ will be a different number between 1 and* m. *However, if 0 <* ***P*** *<* m, *the dendrogram will be truncated to some degree, such that some leaves will contain two or more compounds. In this case, the corresponding entries in ‘T’ will be repeats of the same leaf number*.
>
> *‘T’ can be queried to identify the compounds associated with any particular leaf in the truncated dendrogram by typing* **`find(T==#)`** *in the Command window, where* ***#*** *represents the leaf number of interest. The output from this command will be a list of numbers between 1 and m that indicate which rows in ‘Z’ contain compounds that map to the specified leaf in the dendrogram*.

##### 13. Save the final workspace as a .mat file

*This file can be reopened to give a MATLAB workspace identical to the state that was saved*.

## BASIC PROTOCOL 1b: Clustering to generate a clustergram

### Purpose of method

To reveal the similarities and differences among a set of *m* elements, with respect to a defined list of *n* properties, and represent the results in the form of a clustergram – an *m* x *n* heatmap in which property values are represented by colors/color intensity and similar elements cluster together as do covariant properties. A clustergram is a highly efficient way to visually represent the relationships among the elements and among the properties.

### Background

As with other types of clustering, the method shown here takes as input an *m* x *n* matrix in which each row represents a different element in the data set (in our case, a different compound), and each column represents a different property that describes the elements, and quantifies the degree of relatedness between any two objects in terms of the distance between their positions in *n*-dimensional property space (see Protocol 1 Introduction). The key difference between this method and that described in Basic Protocol 1a is in how the clustering results are represented. Here, the property values are representing as color intensities. For example, values near the high end of the observed range might be represented as bright red, those at the lower end by bright green, and intermediate values by lower intensity shades of the same colors, with values at the mean being represented by black or by white. Translating the property values into colors allows the input data matrix to be represented by an *m* x *n* grid of colored squares, providing a highly intuitive graphical representation of how the property values vary with respect to the mean across the different compounds (i.e. down a column) or, for a given compound, across all the properties (i.e. along a row). A representation of this kind is called a ‘heatmap’. Importantly, the elements (rows) and properties (columns) are not shown in their original order, but instead are clustered so that elements that share similar property profiles are clustered together in the heatmap, as are properties that display similar profiles across the different compounds (i.e. covariant properties). Dendrograms can be added to the element axis or the property axis to provide a yet more detailed and quantitative view of these relationships.

Like other clustering methods, the results of the clustering method described here will depend strongly on the properties or characteristics used for the analysis and how they are scaled, as described in the Introduction. The particular set of elements included in the clustering will also affect the results, in the sense that clustering different randomly selected subsets of elements from the same parent set may give heatmaps that look somewhat different. The results will generally depend less strongly on operational decisions such as distance metric or linkage method.

#### Before you start

Be sure you have installed the necessary toolboxes mentioned in the ‘REQUIRED SOFTWARE’ section. If using your own dataset, make sure you have formatted the data for this analysis as described in the ‘DATA DOWNLOAD AND PREPARATION’ section.

##### 1. Gather the input data

Details of the data download and preparation are given above in Section IV, and spreadsheets containing the data we use are included in the supplementary file ‘Data_Clustering_and_PCA.xlsx’’. In the worked example we use MATLAB as our software tool, though other software packages are available that can perform the same analysis. Readers using MATLAB can execute the protocol by performing each step as described below. For this protocol these comprise a set of 201 elements (compounds), a set of 139 molecular features (descriptors), and the numerical values of each descriptor for each compound (X data).

> *If you are using your own data rather than the supplementary file provided, be sure to format your external data files as described in ‘Section IV. DATA DOWNLOAD AND PREPARATION, and to change the external filenames (blue text) in the following steps to match the names you have assigned to your files*.

##### 2. Import the data into MATLAB

Click the ‘Import Data’ button in the MATLAB Home menu, and use the pop-up window to navigate to the Excel file from Step 1 that contains the input data. Readers using the supplemental Excel file ‘Data_Clustering_and_PCA.xlsx’ should use the sheet named ‘data_Clustering_PCA’.

##### 3. Import the numeric values that make up the body of the input data

i. In your open Data Import window, select the cells that contain the numerical data. Be sure to exclude any column headers or compound names from the selection. The selected data should comprise 201 rows (reflecting the 201 compounds in the data set) and 139 columns (reflecting the 139 molecular descriptors we are using for this analysis).
ii. Go to ‘Output Type’ at the top of the Data Import window and select ‘Numeric Matrix’ from the drop down menu, then go to the Import Selection button at the top right of the Import Data window and click the green check mark (or select ‘Import Data’ from the drop-down menu). You should see a new data object with dimensions 201×139 appear in the Workspace window of your MATLAB desktop (you may need to move the Import Data window aside to see it).
iii. Rename the 201 × 139 workspace variable. For the purpose of this exercise, you should call the new workspace variable **X**. You can do this in the open Data Import window by double clicking on the default name (same as the original Excel filename) that appears just below the top row of the window, and typing in the desired new name. Alternatively, you can click on the newly created workspace variable in the MATLAB Workspace window, to highlight its default name, and type in the new name, **X**.

##### 4. Import the column headings (descriptor names) and row headings (compound identifiers)

In the Data Import window, select the cells that contain the column headings (*i.e*. the names of the 139 molecular descriptors). Do not include cell A1, as this is not a descriptor name. At the top of the Data Import window change the ‘Output Type’ to ‘Cell Array’ with the ‘Text Options’ selected for ‘Cell Array of Character Vectors’ (please note if following along from Basic Protocol 1a this is a different import type than previously discussed), click on ‘Import Data’, and assign the name **Descriptors** to the new data object. A new data object with this name, of size 1 × 139, will appear in your Workspace window. In the Data Import window, select the cells that contain the row identifiers for each element found in the first column in the Data Import window. For the ‘Output Type’ select ‘Numeric Matrix’ (please note if following along from Basic Protocol 1a this is a different import type than previously discussed) into a data object called **Compounds** (again omitting cell A1). The result should be a new data object of size 201 × 1 in your Workspace window.

##### 5. Save the MATLAB workspace that now contains the imported data as a .mat file

> *This file can be reopened to give a MATLAB workspace identical to the state that was saved*.

##### 6. Z-score the data

Z-score the data in **X** to normalize the descriptor values in each column to have mean = 0 and SD = 1, by entering following the command in the Command window:

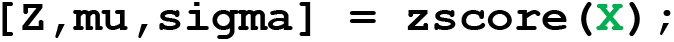

> *Three new variables will be created in the MATLAB Workspace window, called ‘Z’, ‘mu’, and ‘sigma’. ‘Z’ is an m x n matrix (in this case 201 × 139) that contains the descriptor values from* ***X*** *normalized to each have mean = 0 and SD = 1. The variables ‘mu’ and ‘sigma’ are 139 × 1 vectors that contain, respectively, the mean and the SD values of each of the 139 columns in the original data matrix* ***X***.

##### 7. Generate the clustergram

> *For the purpose of this protocol, we excluded one compound (compound ID* ***763****), because a heatmap created using this command can show only up to 200 rows. This was done by selecting the* ***X*** *variable in the MATLAB data object window, selecting the last row by right clicking on the row header (201), and then clicking on “Delete Row”. The same was done for the ‘Compounds’ variable to remove the corresponding compound ID, thereby keeping these two variables the same size*.

**cgo = clustergram(Z)**

*We name our output data variable ‘cgo’ as an abbreviation for ‘clustergram object’. The data variable ‘cgo’ contains all of the properties that define the appearance of the clustergram. These properties can be edited to modify how the clustergram looks, either by typing in new commands as shown in Step 8b, or by clicking on the clustergram itself in the Figure window and changing the desired property. To view all the properties of ‘cgo’, type* `get(cgo)`.

A new Figure window will appear containing a basic clustergram.

##### 8. Label the clustergram

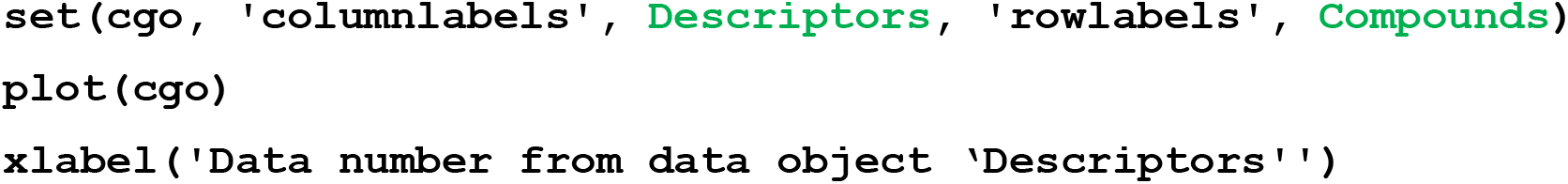
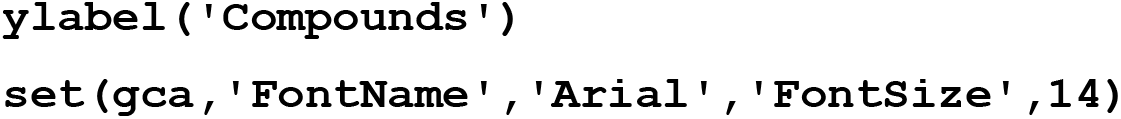

> *Other aspects of the appearance of the plot can be modified using the command* `set(cgo,’property’,’propertysetting’)`. *To see all possible options for plot features that can be changed, and the options that exist for each, that can be changed, enter* `setdisp(cgo)`. *You can also edit the plot using the ‘Edit Plot’ function in MATLAB. Further information can be found at: https://www.mathworks.com/help/MATLAB/ref/plotedit.html*
>
> *After entering the above command, the clustergram should appear as shown in* ***Figure 1.7A***.
>
> Figure 1.7: **(A)** Heatmap of z-scored descriptors and compounds compared. Red indicates values above the mean, black indicates values near the mean, green indicates values below the mean (see color bar at left). The compounds (rows) are clustered as shown by the dendrogram to the left of the heatmap, while the descriptors are clustered as shown by the dendrogram at the top. Thus, compounds that possess a similar profile of descriptor values will show similar color profiles horizontally, while descriptors that co-vary with each other will show similar color profiles vertically. **(B)** A close-up view of a portion of the plot from (A) showing how the dendrogram leaves correspond to individual compounds and descriptors. 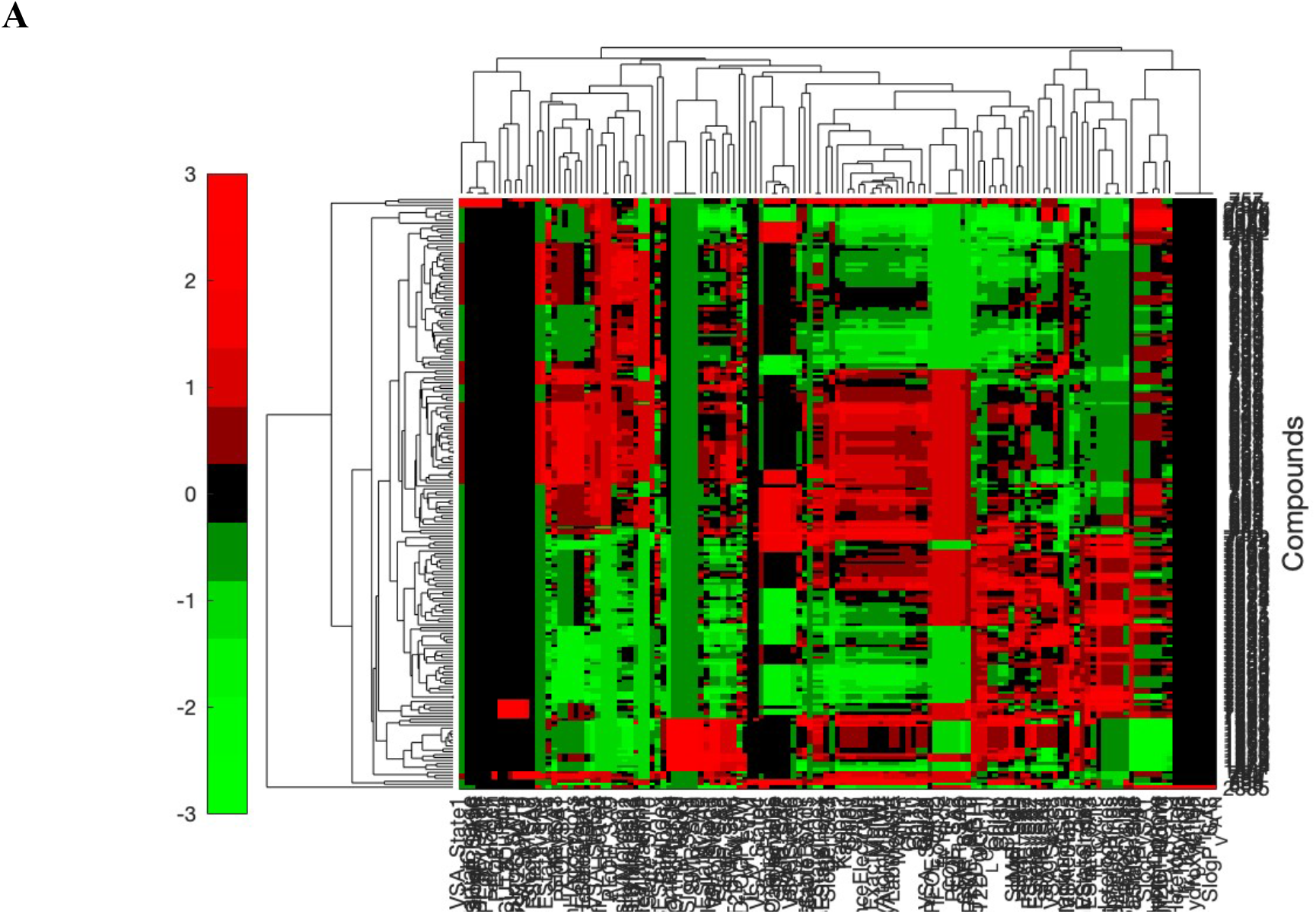 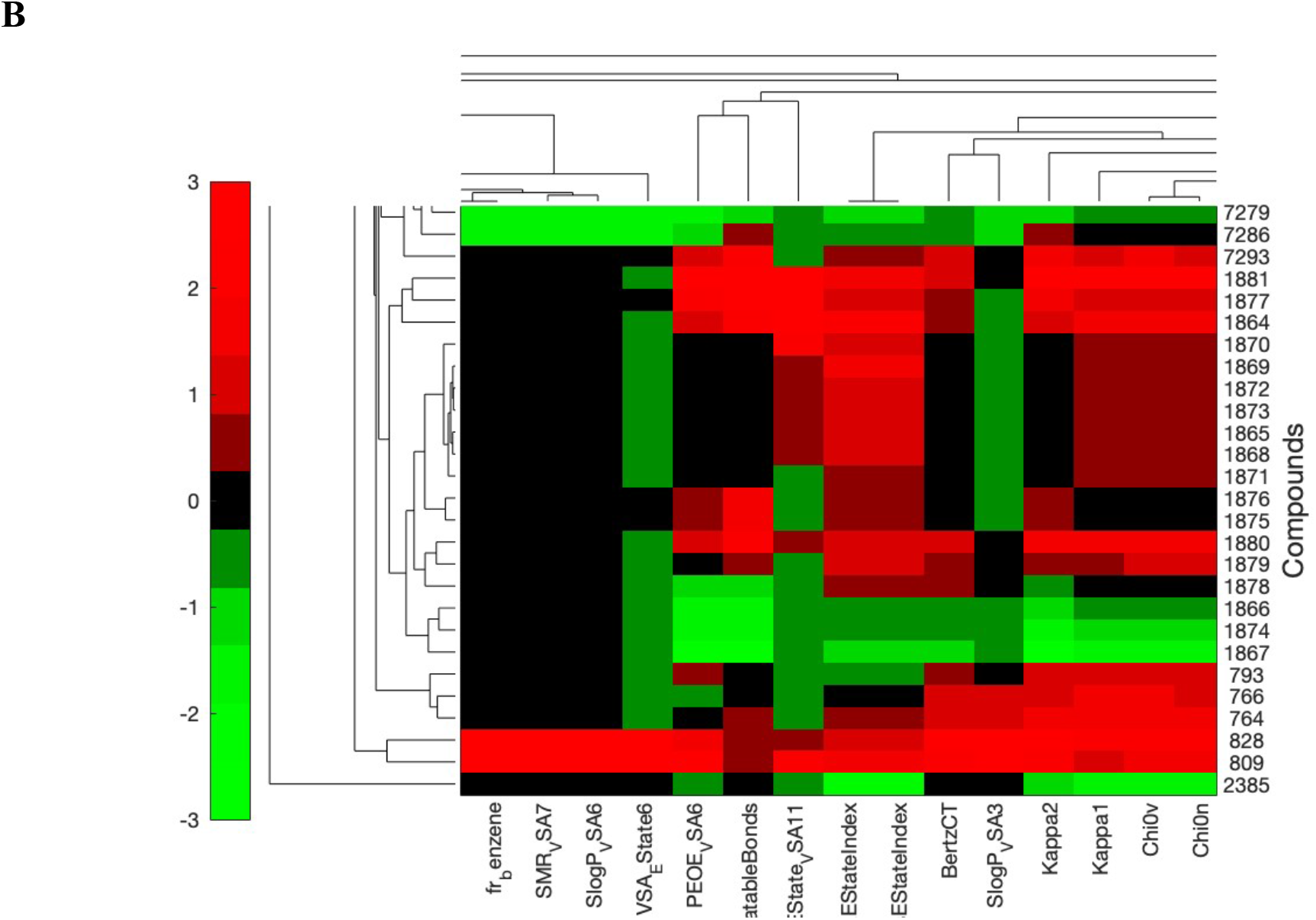
>
> ***Figure 1.7*** *provides another way to represent the distances between the compounds in 139-dimensional property space. The heatmap provides an intuitive and visually impactful way to see not only how the compounds cluster but also, to some extent, why they cluster as they do – that is, which descriptors contribute most strongly to distinguishing one compound cluster from another. The representation also provides information about the relationship between the descriptors; descriptors that cluster closely together are highly covariant, and thus reflect related information about the compounds’ structures and properties. A limitation to this kind of representation is that, if the number of elements (compounds) or properties (descriptors) is large enough that their labels cannot easily be read, as is the case in the figure shown here, it becomes difficult to extract information about individual elements or properties by visual inspection of the plot. In such case, for this level of information it is necessary to refer to the individual data variables contained in the data object ‘cgo’. For the purposes of visual representation, the element and property labels on the right hand and bottom edges of the plot can sometimes be replaced by terms that reflect a higher-level categorization, for example by designating groups of compounds as ‘Chemotype 1’, ‘Chemotype 2’, etc*., *or labeling subsets of descriptors according to the type of property they represent, e.g. ‘Polarity’, ‘Molecular size’, ‘Molecular flexibility’, etc. The extent to which such simplification is possible or useful depends on whether the elements and the properties cluster in ways that can be reasonably summarized in this fashion*.

##### 9. Save the final workspace as a .mat file

> *This file can be reopened to give a MATLAB workspace identical to the state that was saved*.

## BASIC PROTOCOL 1c: *k*-medoids clustering

### Purpose of method

The method of *k*-medoids clustering is one solution to the problem of how to select a representative subset of items from a larger set, as an alternative for example to choosing a random subset that might or might not be accurately representative of the full set. It is useful for tasks such as separating a set of items into a Test Set and a Training Set for the purpose of developing a predictive model using a supervised ML approach such as PLSDA or PLSR (see Basic Protocols 3 and 4).

### Background

The method of *k*-medoids clustering is conceptually related to the somewhat better-known *k*-means clustering. The latter method divides a set of items into *k* groups based on their proximity in *n*-dimensional property space, and outputs the mean values of the properties of the items that make up each group. As molecular scientists, however, we often want to use clustering to identify representative examples from a larger set, such as a subset of chemical compounds that is representative of a larger compound library. For these purposes, *k*-medoids clustering is our method of choice. Like *k*-means clustering, *k*-medoids clustering divides a set of items into *k* groups based on their similarity with respect to a set of properties. But while *k*-means clustering outputs average values of the properties characteristic of each group, *k*-medoids clustering instead identifies the actual item from each group that has properties closest to the average properties. Thus, while *k*-means clustering might tell you that the average value for e.g. the average number of hydrogen bond donors in a particular subgroup of compounds is 2.8, a value that no real compound can have, *k*-medoids clustering will identify *k* actual compounds from the set that best represent the average values of all the properties of the compounds in each of the *k* sub-groups.

Like all clustering methods, the results from *k*-medoids clustering depend strongly on the properties or characteristics used for the analysis. Thus, for example, when using *k*-medoids clustering to identify a representative subset of compounds from a larger collection, the result will depend very strongly on which molecular properties of the compounds are used as input data, and generally much less strongly on operational decisions such as distance metric or linkage method.

#### Before you start

Be sure you have installed the necessary toolboxes mentioned in the ‘REQUIRED SOFTWARE’ section. If using your own dataset, make sure you have formatted the data for this analysis as described in the ‘DATA DOWNLOAD AND PREPARATION’ section.

##### 1. Gather the input data

Details of the data download and preparation are given above in Section IV, and spreadsheets containing the data required to perform the protocols are included in the supplementary file ‘Data_Clustering_and_PCA.xlsx’. For this protocol, these comprise a set of 201 data elements (compounds), a set of 139 molecular features (descriptors), and the numerical values of each descriptor for each compound, provided here in an Excel spreadsheet, on a single page with no gaps or breaks between cells. The supplementary file ‘Data_Clustering_and_PCA.xlsx’ contains the data already formatted in this way, in the sheet named ‘data_Clustering_PCA’.

*If you are using your own data rather than the supplementary file provided, be sure to format your external data files as described in ‘Section IV. DATA DOWNLOAD AND PREPARATION, and to change the external filenames (blue text) in the following steps to match the names you have assigned to your files*.

##### 2. Identify the file that contains the data to be imported into MATLAB

i. Starting with a clear workspace, click on the Import Data button in the MATLAB Home tab (top of screen), navigate to where you have saved the Supplementary File ‘Data_Clustering_and_PCA.xlsx’, and selecting the tab labeled ‘data_Clustering_PCA’ at the bottom of the Data Import window. <u>Be sure you are on the correct sheet before you begin importing data</u>.

##### 3. Import the numeric values that make up the body of the input data

i. In your open Data Import window, select the cells that contain the numerical data. Only the descriptor values (area shaded green in ‘data_Clustering_PCA’) should be selected for import in this step; <u>do not select the column headers in row 1 or the compound names in column 1</u>. The selected data should comprise 201 rows (reflecting the 201 compounds in the data set) and 139 columns (reflecting the 139 molecular descriptors we are using for this analysis), and should include only numbers.
ii. Go to ‘Output Type’ at the top of the Data Import window and select ‘Numeric Matrix’ from the drop-down menu, then go to the Import Selection button at the top right of the Import Data window and click the green check mark (or select ‘Import Data’ from the drop-down menu). You should see a new data object with dimensions 201×139 appear in the Workspace window of your MATLAB desktop (you may need to move the Import Data window aside to see it).
iii. Rename the 201 × 139 workspace variable. For the purpose of this exercise, you should call the new workspace variable **X**. You can do this in the open Data Import window by double clicking on the default name (same as the original Excel filename) that appears just below the top row of the window and typing in the desired new name. Alternatively, you can click on the newly created workspace variable in the MATLAB Workspace window to highlight its default name, and then type in the new name, **X**.

##### 4. Import the row headings (compound identifiers)

In the Data Import window, select the cells in the first column that contain the compound names. <u>Again, do not include cell A1, as this is not a compound name</u>. Change the ‘Output Type’ to ‘String Array’, assign the name **Compounds** to the new data object, then click on ‘Import Data’. A new data object with this name, of size 201 × 1, will appear in your Workspace window.

> *Data objects created in MATLAB can be renamed after the fact, as described in Step 3(iii), but it is good practice to assign the desired name in the Data Import window before creating the object. The reason is that the default name that MATLAB assigns to new data objects is the name of the external file from which the data were imported, making it easy to accidentally over-write a different data object previously imported from the same external file if it was not assigned a unique name upon creation*.

##### 5. Save the MATLAB workspace that now contains the imported data as a .mat file

> *This file can be reopened to give a MATLAB workspace identical to the state that was saved*.

##### 6. Z-score the input data

(see Basic Protocol 1a, step 7 for additional information about z-scoring) Z-score the data in **X** to normalize the descriptor values in each column to have mean = 0 and SD = 1, by entering following the command in the Command window:

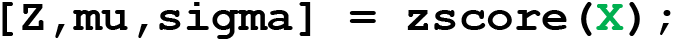

> *Three new variables will be created in the MATLAB Workspace window, called ‘Z’, ‘mu’, and ‘sigma’. ‘Z’ is an m x n matrix (in this case 201 × 139) that contains the descriptor values from* ***X*** *normalized to each have mean = 0 and SD = 1. The variables ‘mu’ and ‘sigma’ are 139 × 1 vectors that contain, respectively, the mean and the SD values of each of the 139 columns in the original data matrix* ***X***.
>
> *See Basic Protocol 1a, Step 7 for additional information about z-scoring*.

##### 7 Perform *k*-medoids clustering on the z-scored data

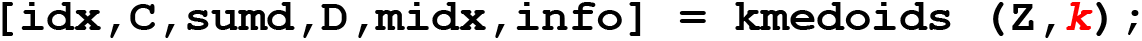

> *The variable* **k** *is the desired number of clusters – in this instance, the desired number of compounds to be selected for the representative subset. For the purpose of this protocol, we will use* **k *= 4***, *to cluster the compounds into 4 groups and identify the compound most representative of each cluster*.
>
> *The default distance metric used by the ‘kmedoids’ command is squared Euclidean distance (‘sqeuclidian’). Other distance metrics, such as city block, can be specified if desired. Type ‘Doc kmedoids’ into the Command window for details*.

Six new data variables will appear in the MATLAB workspace:

‘idx’ *An* m *x 1 vector (in our case 201 × 1, i.e. one row for each of the 201 compounds in Z) in which the single number in each row indicates which cluster, 1 through* **k**, *the compound in the corresponding row of Z belongs to*.

‘C’ *A* **k** *x* n *numerical matrix (in our case, 4 × 139) in which each row contains the values of the n descriptors that identify the median point of a given cluster in n-dimensional space, and each row represents a different cluster*.

‘sumd’ *A* **k** *x 1 vector containing the sum of the data point-to-medoid distance for each cluster*.

‘D’ *An* m *x* **k** *matrix in which each row gives the distances of a given data point (compound) from each of the* **k** *cluster medoids*.

‘midx’ *A* **k** *x 1 vector (in our case, 4 × 1) in which the single number in each row identifies the compound that is the medoid for each of the* **k** *clusters. The compounds are identified by their row number in Z. This is the result that identifies the subset of* **k** *compounds chosen by the method as the best representatives of each cluster*.

‘info’ *Information specifying the settings used in the ‘kmedoids’ command, organized in the form of a so-called ‘structure array’*.

##### 8. Identify which compounds belong to each cluster

The contents of ‘idx’ comprise a list of which cluster each compound in ‘Z’ belongs to, the compounds being identified by their row numbers in ‘Z’. A list of the compounds that belong to a given cluster, identified by compound name/ID number, can be generated as follows:

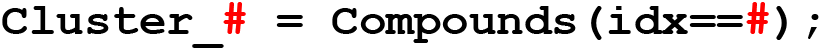

where # is the number of the cluster of interest (i.e., in this case, 1, 2, 3 or 4).

A new data variable will appear, named ‘Cluster_**#**’.

> *‘Cluster_****#****’ is a vector containing the compound names/ID numbers, taken from the data variable called ‘Compounds’ created in Step 4, for all the compounds that belong to cluster number #*.

##### 9. Identify the compounds most representative of each cluster

To identify the compounds listed in ‘midx’ as the four most representative of the four clusters into which we divided the 201-compound set, double click on ‘midx’ in the Workspace window. The contents of ‘midx’ reveal that these are the compounds found in rows 24, 155, 112, and 141 of the input data matrix ‘Z’.

The names/ID numbers for these four compounds can be identified using the following command:

**Medoids = Compounds(midx(:,1))**

This command returns the following compound IDs for the four compounds that represent the medoids (i.e. the most typical representative of) each of the four clusters into which we divided the set:

##### 10. Save the final workspace as a .mat file

> *This file can be reopened to give a MATLAB workspace identical to the state that was saved*.
>
> *For a description of how the compound clusters identified in steps 7 and 8 can be visualized by coloring data points in Principal Component space, see Basic Protocol 2, Step 17*.

### Understanding the Results of Basic Protocols 1a, b and c

Basic protocols 1a and 1b generate, respectively, a dendrogram and a heatmap illustrating the relationships between the compounds and/or the descriptors in the data set. The dendrogram comprises nested groups or clusters of compounds that diverge from one another into different branches. Each branches contain a subset of the elements (compounds) in the data set. The end of each branch, referred to as a leaf, contains an individual element. The vertical distance that connects two different leaves or branches is referred to as a node.

For illustrative purposes, in the figure below we have selected 3 compounds to compare from the dendrogram in **Figure 1.6**. Compounds ID1880 and ID7264 come from branches that are connected by a node with a relatively low value on the distance axis, giving the expectation that these two compounds will be relatively similar with respect to the properties in X. In contrast, compound ID828 was identified on the dendrogram as an outlier, indicating more different properties. Visual inspection of the chemical structures in **Figure 1.8** is consistent with these expectations, as ID828 is distinct from the other two compounds by virtue of possessing an extended segment of nonpeptidic structure in the macrocycle ring, as well as an N-ethel group and an unnatural amino acid.

**Figure 1.8.**
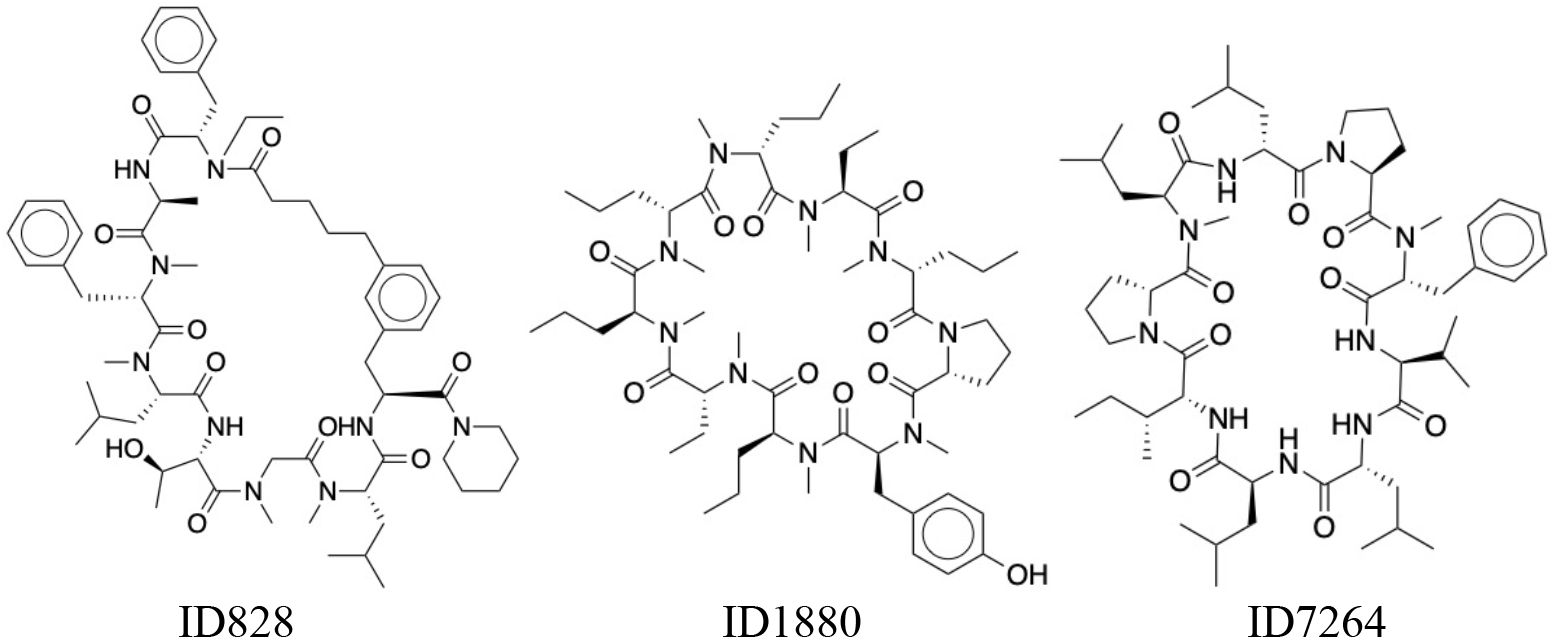
Chemical structures of two compounds that cluster together on the dendrogram in Figure 1.6, and one that was rated as an outlier.

Inspection of the dendrogram in Figure 1.6 identifies the following six compounds as the most extreme outliers, as measured by the position on the distance axis of the nodes that connect them to the other compounds:

The three compounds ID755, ID757, and ID790 were also classified as outliers with respect to the majority of compounds in the set, but clustered together indicating similarity with each other. These chemical structures of these compounds are shown in **Figure 1.9**.

**Figure 1.9.**
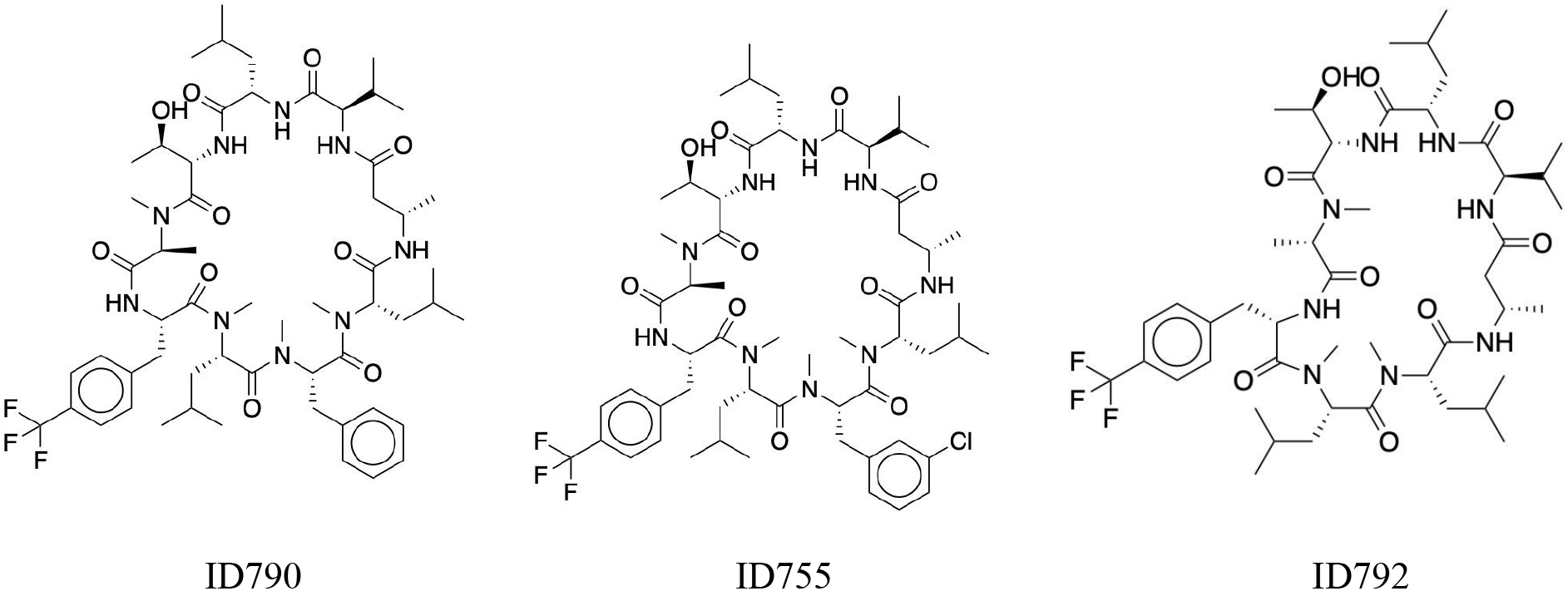
Chemical structures of three compounds that cluster together as outliers in the dendrogram in Figure 1.6.

## BASIC PROTOCOL 2: Principal Component Analysis (PCA)

### Purpose of Method

Principal Component Analysis (PCA) is a type of unsupervised machine learning. It is used to reveal patterns in complex data sets by reducing the dimensionality of the data, so that the information content can be visualized and analyzed in terms of a smaller number of dimensions known as Principal Components.

### Key concepts: Variance and covariance

<u>Variance:</u> For a set of *n* numerical values of some quantity *x*, the extent to which the individual values of *x* are distributed around the mean (designated *m*_*x*_ or just *m*) can be described by the standard deviation, or by a related measure called the <u>variance</u> that is given by the following equation:

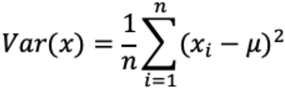

In words, the variance is the sum of the squares of the difference between each value and the mean (μ), divided by the total number of values. Variance is related to the standard deviation (σ) by Var(*x*) = σ^2^. Like standard deviation, the numerical value of the variance is sensitive to the numerical magnitude of the values of *x*, and thus to the units in which *x* is expressed. For example, for a set of lengths expressed in mm, both the variance and the standard deviation will have larger numerical values than if the same set of length measurements were expressed in units of cm or m.

<u>Covariance</u> is a metric, conceptually related to variance, that quantifies, for a set of (*x,y*) data points, how much the values of *x* are correlated with the values of *y*. For example, if one were to have a set of measurements of the heights and bodyweights for a random set of people, there would likely be some correlation between these variables. This is because very tall people, on average, are likely to be bigger and heavier than much shorter people. If one were to plot a graph of weight versus height for this set of people, it might therefore look something like the following (**Figure 2.1**):

**Figure 2.1.**
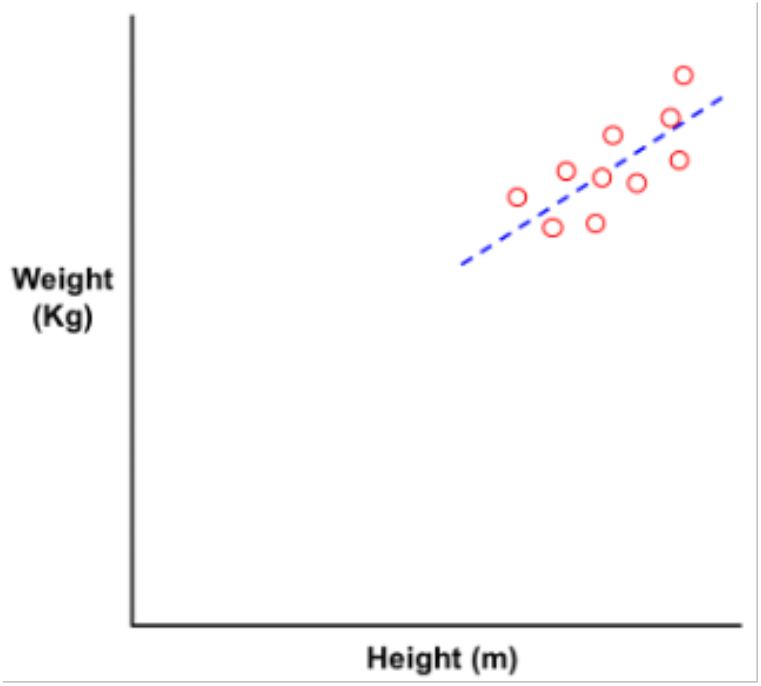
Hypothetical plot illustrating the partial covariance between height and weight that might be expected to exist for a random group of individuals (not real data).

One way to quantify the extent to which the *x* and *y* values in the above data set are correlated is to calculate their <u>covariance</u>, defined as follows:

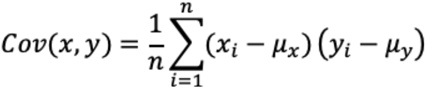

In words, for an *x,y* data set comprising *n* data points, the covariance between *x* and *y* is the difference between each *x* value and the mean of all the *x* values, multiplied by the corresponding difference for *y* (i.e. *y* – mean_y_) for the same data point, added together for all *n* data points, and then divided by *n* to get the mean value of the multiplied difference per data point.

If *x* and *y* are highly covariant – that is, if *x* tends to be large when *y* is large, and small when *y* is small – then the product (*x*_*i*_ – μ_*x*_)(*y*_*i*_ – μ_*y*_) will be a positive number, because (*x*_*i*_ – μ_*x*_) and (*y*_*i*_ – μ_*y*_) will tend to either both be positive or both negative. In contrast, if *x* and *y* are uncorrelated, so that a large value of *x* were equally likely to be associated with a large or a small value of *y*, the product (*x*_*i*_ – μ_*x*_)(*y*_*i*_ – μ_*y*_) will have a mix of positive or negative values, and over a large, randomly distributed data set will tend to zero. Finally, if *x* and *y* are anti-correlated – that is, if higher values of *x* tend to be associated with *lower* values of *y* – then (*x*_*i*_ – μ_*x*_) and (*y*_*i*_ – μ_*y*_) will tend to have opposite signs, and covariance will be negative. Thus, if the calculated covariance is positive, *x* and *y* are correlated to some extent, if zero they are uncorrelated, and if negative they are anti-correlated. The highest value of covariance will be found when *x* and *y* are exactly proportional, and the lowest (most negative value) when *x* is exactly proportional to minus *y*.

The numerical value of the covariance depends not only on the extent to which *x* and *y* are correlated, but also on the numerical magnitudes of the mean values of *x* and *y* and on the variances of the *x* and *y* values about their respective means. It is therefore not always obvious whether a particular positive value of the covariance represents a very high degree of covariance, as when *x* and *y* are exactly or very nearly proportional, or whether *x* and *y* simply trend together to a more limited extent. The covariance can be made more intuitively interpretable if it is normalized by dividing by the product σ_x_σ_y_, where σ represents standard deviation. This normalized covariance is called the Pearson Correlation Coefficient (PCC).

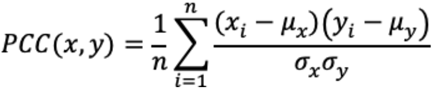

This function can also be expressed in the equivalent form in terms of the z-scored values of *x* and *y* (see Basic Protocol 1a, Step 7 for an explanation of z-scoring):

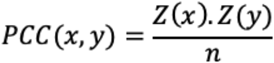

The PCC can have values between +1 and -1, with PCC = +1 indicating that the differences in *x* between any two data points are exactly proportional to the differences in *y*, PCC = 0 indicating *x* and *y* are uncorrelated, and PCC = -1 indicating that Δ*x* any Δ*y* are perfectly anti-correlated (i.e. Δ*x* is proportional to minus Δ*y*). A value of PCC between zero and 1 indicates that *x* and *y* are positively correlated to some extent, with a higher value indicating a stronger correlation. Values of PCC between zero and -1 similarly quantify the degree of anti-correlation (**Figure 2.2**).

**Figure 2.2.**
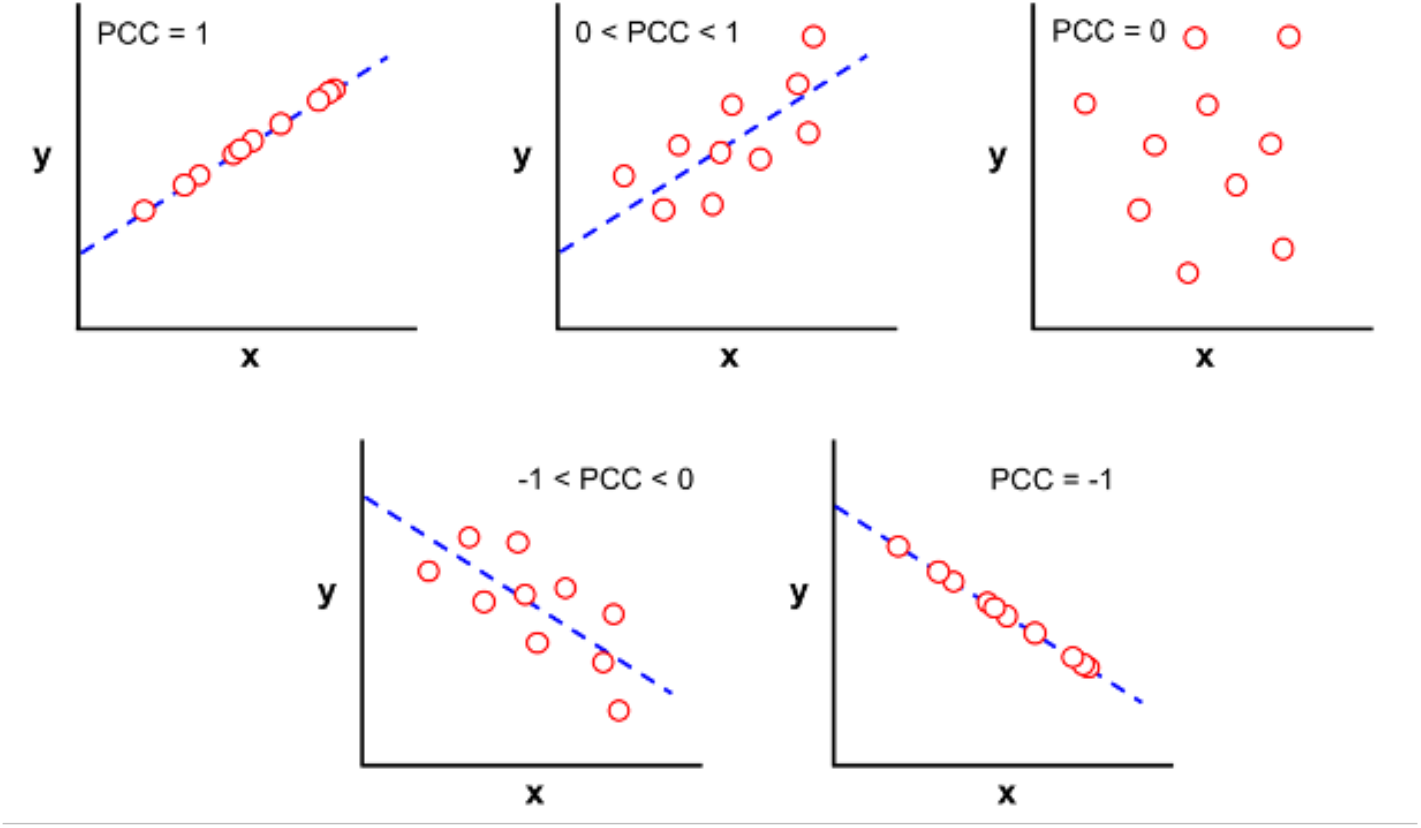
Hypothetical plots illustrating different degrees of covariance between two variables, *x* and *y*, and the value of the Pearson Correlation Coefficient (PCC) that would result (not real data).

### Principal Component Analysis

In the example discussed above, involving measurements of height and bodyweight for a set of *n* individuals, the information in this data set can be fully represented by a set of *n* (*x,y*) coordinates, where *x* is the height and *y* is the weight (or *vice versa*) of each individual. However, because height and weight are to some extent covariant, if we know the height of an individual, we are also in a better position to estimate their weight compared to if we just took a random guess. Because we know that height and weight are to some extent covariant, then we know that an individual with a height that is far above the mean is more likely than not to have a weight that is also above the mean. Indeed, if the correlation between height and weight were perfect (i.e. PCC = 1) then, if we knew an individual’s height, we could exactly specify their weight. In this extreme case of perfect covariance, we would only need to know one number, either height or weight, for each individual, to fully capture both pieces of information in the data set. Knowing both height and weight would in this case be fully redundant. Thus, *covariance reduces the amount of information that a data set contains, such that the information can, at least in principle, be represented using fewer bits of information*. <u>Principal Component Analysis</u> (PCA) is a statistical method for eliminating or reducing the redundancy in a data set that contains elements of covariance, so that the information content can be represented in a more concise – and often more easily interpretable – way. The more covariant the data, the greater the degree to which PCA can reduce it to the core of non-redundant information it contains.

### How PCA works

Consider the *x,y* dataset plotted in the figure below. Such a data set comprises two pieces of information – the value of a property *x* and of a second property *y* – for *n* data points. This information can be fully described by a set of *n* (*x,y*) coordinates, as represented for example in the scatter plot in **Figure 2.3**. For the hypothetical data shown in the figure, there is significant variance in both the *x* and the *y* values. Because the overall variance in the data is distributed between the two variables, the values for both variables must be known to fully account for all the variance in the data (**Figure 2.3**).

**Figure 2.3.**
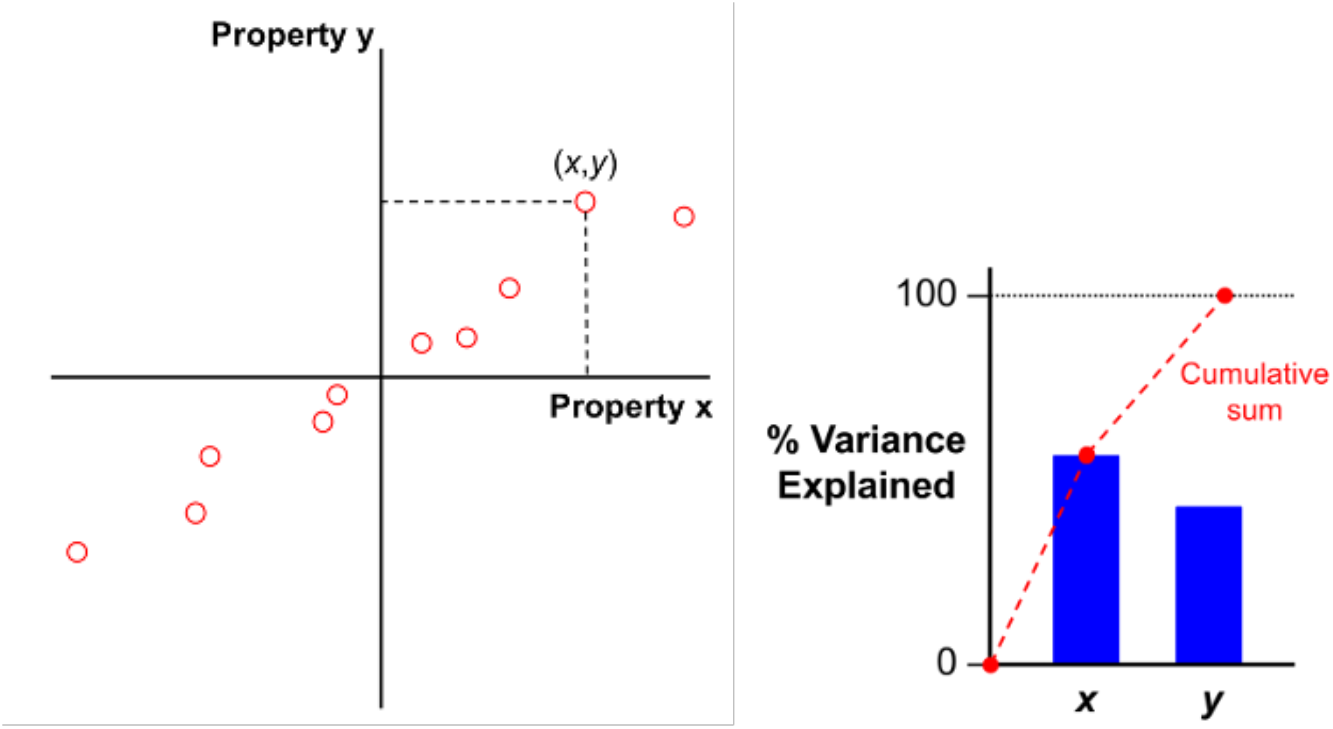
Hypothetical plot (not real data) illustrating that if two different properties, *x* and *y*, each contain substantial variance, then quantifying the total variance across a set of items requires knowledge of both property values for each item, corresponding to two dimensions of information.

It is clear from the figure that *x* and *y* are strongly covariant. Therefore, one could imagine defining a new system of coordinates, generated by rotating the original axes about the origin, such that the first axis (called Principal Component 1, or PC1) passes through the data points at an angle that captures as much of the variance as possible. A second new axis, PC2, at right angles to the first, could then capture the small amount of remaining variance that was not encompassed by PC1 (**Figure 2.4**). Using this new coordinate system, each data point can still be described by a coordinate pair, now (PC1,PC2 rather than *x,y*). But, crucially, almost all of the variance in the data set is now represented in the position of each data point along axis PC1 (i.e. in the value of the first coordinate, PC1), with very little left to be explained by PC2. Consequently, the data set could be reasonably represented using only the values of PC1, ignoring PC2 altogether, with very little loss of information. In summary, because *x* and *y* were highly covariant, representation of the data in the PC1,PC2 coordinate space instead of the original *x,y* coordinate space allowed most of the information content in the data to be compressed into a single coordinate, PC1.

**Figure 2.4.**
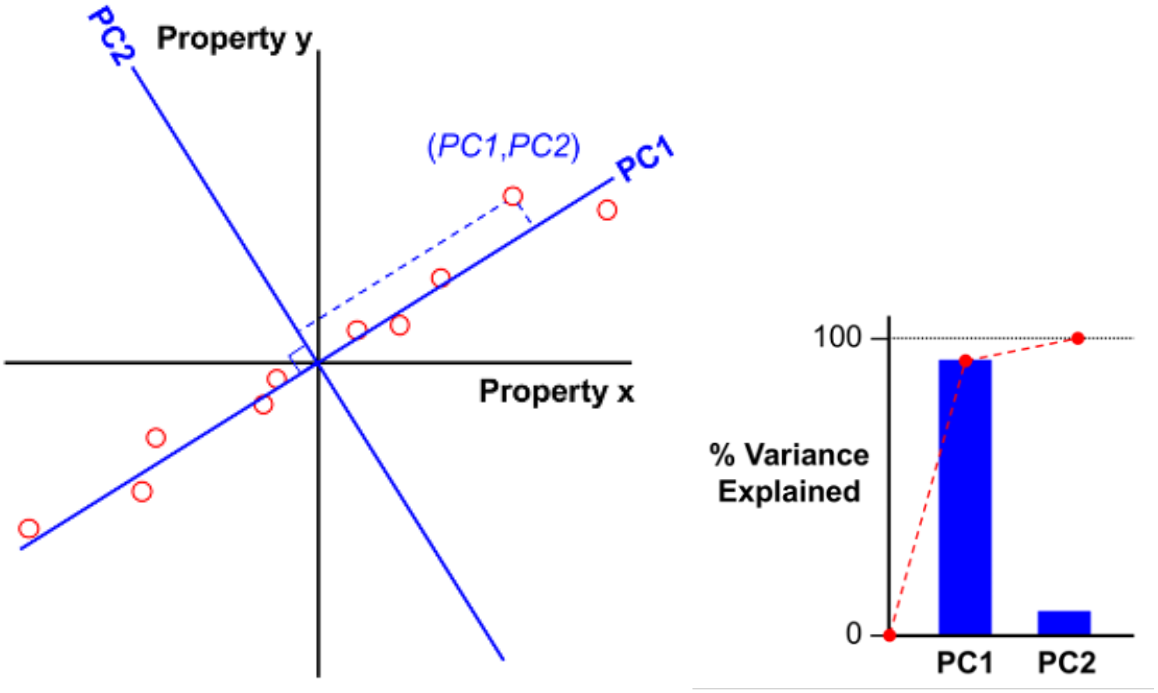
Plot illustrating, for the hypothetical *x,y* data from Figure 2.3, how PCA rotates the coordinate system used to describe the elements in the data set such that the greatest possible amount of variance is contained in the first axis, called PC1, with any residual variance captured by the second, orthogonal axis, called PC2. As a result, most of the information from the original *x,y* coordinates is now contained in a single axis, PC1, such that the data set can be described using just a single number per data point with minimal loss of information. PCA has thus reduced the dimensionality of the data from two dimensions to one, made possible by the covariance that exists between *x* and *y* in the original input data. In the same way, PCA can reduce the dimensionality of *n*-dimensional data sets, where the values of *n* different properties are provided for each data point, provided that there is some degree of covariance between some of the properties.

PCA performs the transformation described above for data sets containing any number of coordinates. For example, one can imagine for a group of individuals having more information than just their height and weight, but perhaps also their shoe size, the speed at which they can run 100 m, or any other number of quantifiable characteristics. In such a case, the data comprises a set of *n* characteristics per individual. To plot this data set would require using *n* axes, one for each property, making it difficult to visualize the relationships between the data points if *n* is a large number. PCA identifies a line through this *n*-dimensional property space that passes through the data points in such a way as to capture the most possible variance, as illustrated above for the two-dimensional case, and designates this new axis PC1. The data are then zeroed with respect to PC1 by subtracting out the variance captured in this coordinate, and the process is repeated by identifying another new axis, PC2, that is orthogonal to PC1 and captures as much as possible of the residual variance that remains in the subtracted data. This process is repeated *n* times to generate a set of *n* mutually orthogonal axes, designated PC1 to PC*n*, that together define a new coordinate system which fully describes the data set. Plotting or describing the data in terms of this new coordinate system is referred to as working in “Principal Component space”. The PC axis system differs from the original coordinate system in squeezing as much of the variance as possible into the lowest-numbered Principal Components. PCA thus reduces the dimensionality of the data set, allowing the data to be represented using fewer axes while minimizing loss of information. The extent to which this compression can be achieved depends on the degree of covariance that exists between the different characteristics in the original data set.

**Note:** An early step in the PCA algorithm is to center the values in each column of the data matrix (i.e. for each property) on zero. This is done by subtracting the mean value for a column from each value in that column. This procedure is necessary to center the data in *n*-dimensional property space around an origin that represents the center of mass of all the data points. Centering the data in this way ensures that a set of orthogonal Principal Component axes that pass through the origin will also pass through the center of mass of the data points, as is necessary if the PCs are to most efficiently capture all the variance in the data. Centering is not necessary if the data have previously been z-scored (see Basic Protocol 1a, Step 7).

### Data outputs from PCA

The coordinates of the data set in PC space are called the ‘scores’. Each of the *m* items in the data set will have a score for each PC, and the total number of PCs equals the number of original properties. Thus, the scores make up a matrix of size *m* x *n*, identical to the size of the original input data matrix, X. Unlike the input data, however, the PC axes are ordered such that PC1 contains the highest amount of variance that can be captured in a single linear dimension, PC2 contains the second highest variance, and so on. Therefore, when plotting or analyzing the data it is often possible to work with only the two or three top-ranked PCs, allowing visualization by plotting the data (scores) on two-dimensional or three-dimensional Cartesian axes with minimal loss of information. Among the outputs from PCA is a *n* x 1 vector containing the percentage of the total variance that is contained in each successive PC. The higher the percent variance explained, cumulatively, by the first two or three PCs, the less information is lost if the scores are plotted on a two- or three-dimensional scatter plot.

Information about the extent to which each of the original *n* properties contributes to PC1, PC2, etc. is given by a set of numbers called ‘coefficients’, or sometimes ‘loadings’. Each PC will have a set of coefficients, one for each of the original properties, that indicate how much that property is contributing to the PC. This information is contained in a matrix of size n x n, where the columns are the different PCs, from PC1 to PCn, and the rows are the original properties, with the cells being populated by the coefficient of a give property in a particular PC. A coefficient with a large positive value indicates that a larger value for that property will result in a substantially larger score in that PC. A coefficient with a small value, whether positive or negative, indicates that the value of the scores in that PC is relatively insensitive to the property. A large but negative coefficient indicates that a very negative value for the property in question will increase the scores in the PC. Thus, by looking for the properties that have the largest absolute value for the coefficients in a given PC indicates which properties are most influential in driving the scores in that PC, and the sign of the coefficient for a given property indicates whether that property is positively or inversely related to the scores. Evaluation of the coefficients can therefore help identify which set of the original properties a given PC most reflects.

The data output from PCA using the scripts provided in this protocol returns five main output files:

Component Coefficients: An *n* x *n* matrix showing how much each of the original *n* properties (rows) contributes to each of the *n* Principal Components (columns). In the standard MATLAB implementation of PCA, these coefficients are stored in a data matrix named “wcoeff”. For many uses, these coefficients must be normalized before they can be properly interpreted (specifically, the coefficient matrix must be made ‘orthonormal’). In MATLAB, the coefficients that have been made orthonormal are typically stored in a data vector called “coefforth”.

Scores: An *m* x *n* matrix that shows the coordinates of each of the original *m* items (rows) in each of the *n* PCs (columns). In MATLAB, these values are typically stored in a data matrix named “score”.

Component Variance: An *n* x 1 column vector indicating how much variance is explained by each PC. The variance is highest for PC1 (because PC1 was chosen precisely to capture the most variance possible), and incrementally lower for each subsequent PC. In MATLAB, these values are typically stored in a data matrix named “latent”.

Percent Variance Explained: An *n* x 1 column vector indicating what percentage of the total variance in the data is explained by each PC, calculated from the component variances described above. In MATLAB, these values are typically stored in a data matrix named “explained”.

> *Hotellings T-squared Statistic:* A measure of how far each data point lies from the center of mass of all the data points (i.e. from the origin), in PC space (Hotelling, 1931). Useful for identifying outliers. In MatLab, these coefficients are typically stored in a data matrix named “tsquared”.

## BASIC PROTOCOL 2: Principal Component Analysis (PCA)

### Before you start

Be sure you have installed the necessary toolboxes mentioned in the ‘REQUIRED SOFTWARE’ section. If using your own dataset, make sure you have formatted the data for this analysis as described in the ‘DATA DOWNLOAD AND PREPARATION’ section.

#### 1. Gather the input data

Details of the data download and preparation are given above in Section IV, and spreadsheets containing the data we use are included in the supplementary file ‘Data_Clustering_and_PCA.xlsx’. For this protocol these comprise a set of 201 data elements (compounds), a set of 139 molecular features (descriptors), and the numerical values of each descriptor for each compound. These are provided here in an Excel spreadsheet, on a single pa ge with no gaps or breaks between cells. The supplementary file ‘Data_Clustering_and_PCA.xlsx’ contains the data already formatted in this way, in the tab named ‘data_Clustering_PCA’.

> *If you are using your own data rather than the supplementary file provided, be sure to format your external data files as described in ‘Section IV. DATA DOWNLOAD AND PREPARATION, and to change the external filenames (blue text) in the following steps to match the names you have assigned to your files*.

#### 2. Identify the file that contain the data to be imported into MATLAB

Click the ‘Import Data’ button in the MATLAB Home menu, and use the pop-up window to navigate to the Excel file from Step 1 that contains the input data (‘data_Clustering_PCA’).

#### 3. Import the numeric values that make up the body of the input data

i. In your open Data Import window, select the cells that contain the numerical data. <u>Be sure to exclude any column headers or compound names from the selection</u>. The selected data should comprise 201 rows (reflecting the 201 compounds in the data set) and 139 columns (reflecting the 139 molecular descriptors we are using for this analysis). Under ‘Output Type’ select ‘Numeric Matrix’, then go to the Import Selection button at the top right of the Import Data window and click the green check mark (or select ‘Import Data’ from the drop-down menu). You should see a new data object with dimensions 201×139 appear in the Workspace window of your MATLAB desktop (you may need to move the Import Data window aside to see it).
ii. Rename the 201 × 139 workspace variable. For the purpose of this exercise, you should call the new workspace variable **X**. You can do this in the open Data Import window by double clicking on the default name (which will be same as the original Excel filename) that appears just below the top row of the window, and typing in the desired new name. Alternatively, you can click on the newly created workspace variable in the MATLAB Workspace WIndow, to highlight its default name, and type in the new name, **X**.

#### 4. Import the column headings (descriptor names) and row headings (compound identifiers)

In the Data Import window, select the cells in the top row that contain the column headings (*i.e*. the names of the 139 molecular descriptors). <u>Do not include cell A1, as this is not a descriptor name</u>. At the top of the Data Import window change the ‘Output Type’ to ‘String Array’, assign the name **Descriptors** to the new data object, then click on ‘Import Data’. A new data object with this name, of size 1 × 139, will appear in your Workspace window. To import the compound identifiers, select the cells in the first column that contain the compound names. <u>Again, do not include cell A1, as this is not</u> a compound name. Change the ‘Output Type’ to ‘String Array’, assign the name **Compounds** to the new data object, then click on ‘Import Data’. A new data object with this name, of size 201 × 1, will appear in your Workspace window.

> *Data objects created in MATLAB can be renamed after the fact, as described in Step 3(ii), but it is good practice to assign the desired name in the Data Import window before created the object. The reason is that the default name that MATLAB assigns to new data objects is the name of the external file from which the data were imported, making it easy to accidentally over-write a different data object previously imported from the same external file if it was not assigned a unique name upon creation*.

#### 5. Save the MATLAB workspace as a .mat file

#### 6. Z-score the input data

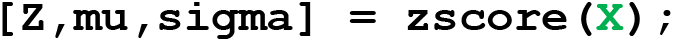

> *As described in more detail in Basic Protocol 1a, Step 7, this command creates three new data objects in the output window: a matrix of z-scores (*Z*), an* m *x 1 column vector*, mu, *that contains the mean values for each descriptor, and a second* m *x 1 matrix*, sigma, *that contains the standard deviations of the values for each descriptor*.
>
> *The labels ‘*Z*’, ‘*mu*’ and ‘*sigma*’ are arbitrary, and are simply names that are often chosen for these variables. Any valid filenames could be used on the left-hand side of this command*.
>
> *For further details on the zscore command, type ‘Doc zscore’ into the MATLAB Command window*.

#### 7. Calculate the covariance between the z-scored descriptor values

~~~
C = corr(Z,Z);
~~~

A new data variable called ‘C’, in the form of a data matrix of size *n* x *n*, will be created in the Workspace window, containing the Pearson Correlation Coefficient (PCC) for each descriptor with all 139 of the descriptors contained in ‘Z’.

> *In ‘C’, the cells that run diagonally from top left to bottom right contain the PCC values for each descriptor with itself, and so, by definition, will all contain the value 1. The off-diagonal cells contain the PCC values for each descriptor with respect to a different descriptor, and will lie between +1 (perfect correlation) through zero (no correlation), to -1 (perfect anticorrelation). The values in the off diagonal cells are mirrored about the diagonal because PCC for descriptor* i *with respect to descriptor* j *is identical to PCC for descriptor* j *with respect to descriptor* i. *Thus, either half of the matrix contains all of the unique information contained in ‘C’*.
>
> *The covariance matrix ‘C’ can be viewed most conveniently by pasting the data into Microsoft Excel and using ‘Conditional Formatting: New Rule: 3 color scale’ to color each cell according to the PCC value it contains. Such a plot is shown below (****Figure 2.5****) for a small section of ‘C’, for which we set the color scheme in Excel as follows: red for PCC = +1, white for PCC = 0, and blue for PCC = -1*.

**Figure 2.5.**
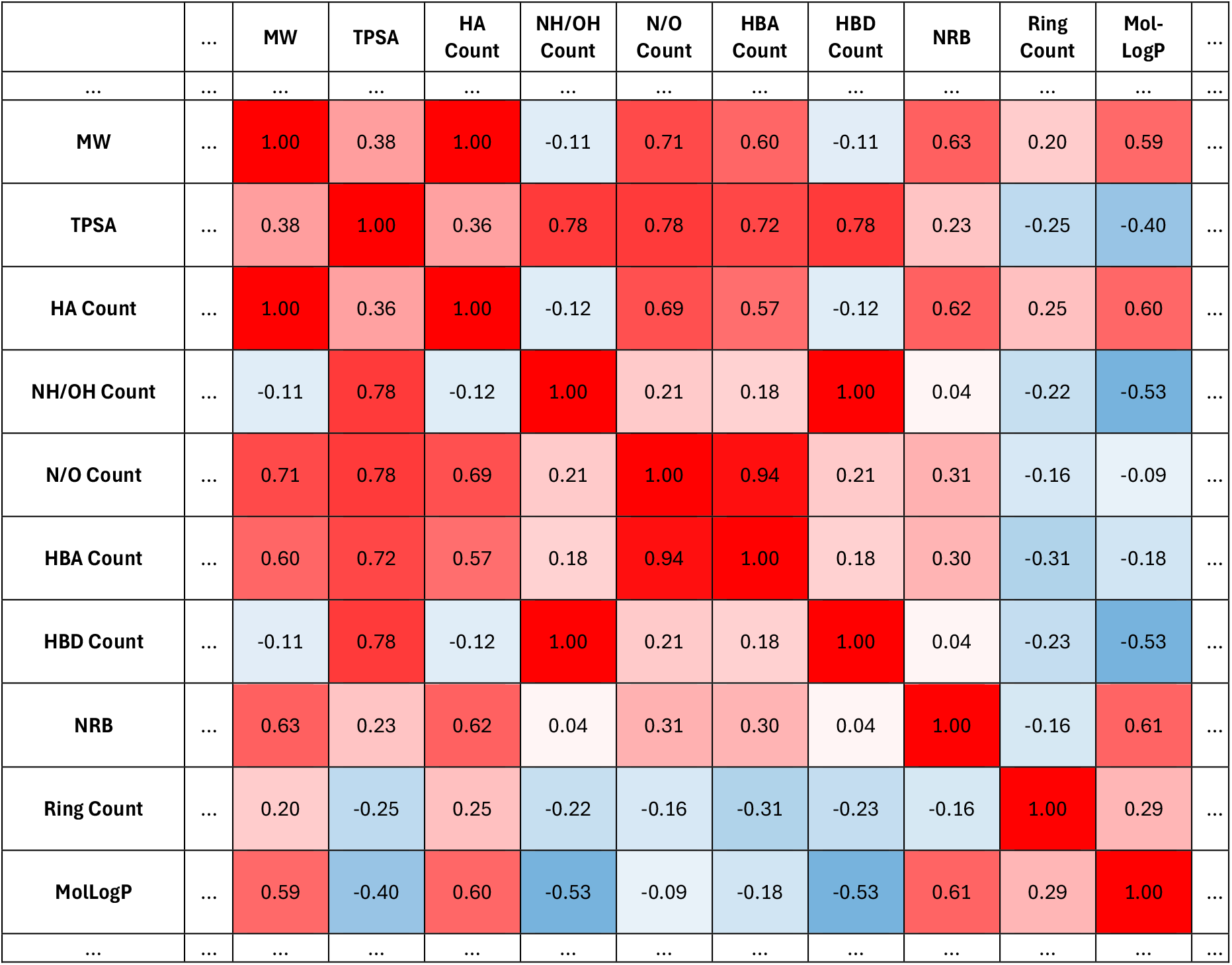
A portion of the covariance matrix ‘C’, created in Step 7, showing the covariance between the 10 molecular descriptors expressed in terms of the Pearson Correlation Coefficients (PCC). Cells are color-coded from intense red for PCC = 1 through white for PCC = 0 to intense blue for PCC = -1. Note that all cells on the diagonal, which represent PCC for each descriptor with itself, contain values of PCC = 1, as any set of numbers is perfectly covariant with itself. The PCC values in all off-diagonal cells have a plane of symmetry about the diagonal. Abbreviations are: MW, molecular weight; TPSA, topological polar surface area; HA, heavy atoms; HBA, hydrogen bond acceptor; HBD, hydrogen bond donor; NRB, number of rotatable bonds.

#### 8. Calculate the weights to apply to the descriptors in X to counter bias towards high-variance descriptors

> *While z-scoring is an effective way to scale and canter the data for some functions it is generally not recommended for PCA. The reason is that setting the variances of the different descriptors as equal in this way carries the risk that, for a descriptor the value of which remains roughly constant across the elements in the data set, these insignificant variations will become magnified as if they were as important as the more consequential variations in other descriptors. Nonetheless, it can be useful to weight the influence of the different descriptors to reduce the bias towards those with the largest numerical variance. Here, we show how this can be achieved by calculating a set of weighting factors corresponding to the reciprocal of the variance in each descriptor, which can be incorporated into the PCA calculation to counter the bias towards large-variance descriptors*.

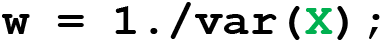

> *This command takes the reciprocal for the covariance in each column in ‘X’ and returns the result in a new data matrix, ‘w’, with dimensions of 1 x* n.

#### 9. Compute the Principal Components

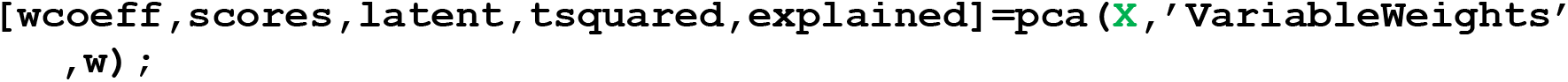

Five new data variables will be generated in the Workspace window: ‘wcoeff’ (size *n* x *n*, the weighted component coefficients), ‘scores’ (size *m* x *n*, the coordinates for each element of the data set in PC space), ‘latent’ (size *n* x 1, indicating how much of the total variance in **X** is explained by each Principal Component), ‘explained’ (size *n* x 1, the values in ‘latent’ expressed as percentages of the total variance), and ‘tsquared’ (size *m* x 1 column, the Hotelling’s T-squared statistic for each data element in **X**). See Introduction to Basic Protocol 2 for additional information about these outputs of PCA.

#### 10. Normalize the calculated coefficients

~~~
coefforth = diag(sqrt(w))*wcoeff;
~~~

*This command normalizes (or, more precisely, orthonormalizes) the coefficients contained in the data variable wcoeff such that each has the same vector length (of 1) when plotted in PC space. This process is necessary for some subsequent steps*.

#### 11. Use the percent variance explained by each Principal Component to choose how many Components to use for further analysis

(a) Plot the data in the variable ‘explained’ in the form of a Pareto chart (Figure 2.6)

~~~
pareto(explained,’MarkerFaceColor’,’blue’,’LineStyle’,’none’,
   ‘Marker’,’o’,’MarkerEdgeColor’,’none’)
xlabel(‘Principal Component’)
ylabel(‘Variance Explained (%)’)
set(gca,’FontName’,’Arial’,’FontSize’,14)
~~~

> *The ‘pareto’ command will plot the number of bars required to add to 95%, up to a maximum of 10 bars*.

(b) Use the Pareto chart (Figure 2.6) to choose how many Principal Components must be included to capture the bulk of the total variance in **X**.

**Figure 2.6:**
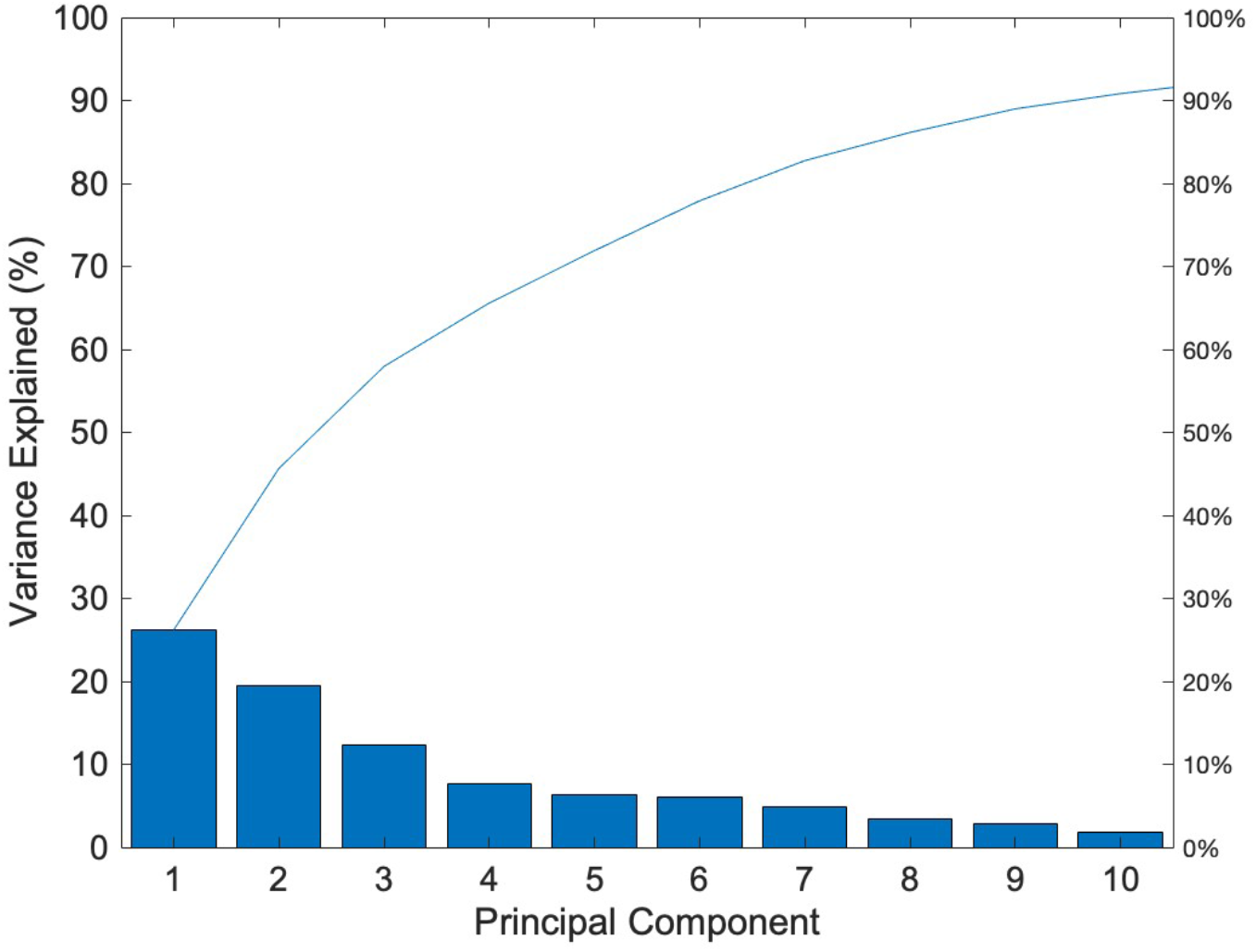
Pareto chart showing the percent of the total variance explained by each of the first 10 Principal Components (bars), with the cumulative sum of the percent variance explained shown as a solid line.

> *How many PCs to choose depends on the purpose of the analysis. In the current case, examination of* ***Figure 2.5*** *shows the first six or seven PCs collectively explain >80% of the total variance in* ***X***. *Thus, PCA has reduced the dimensionality of the data from the original 139-dimensional space (i.e. one Cartesian axis per descriptor) to ∼6 axes with minimal loss of information. For some purposes, therefore, PCs 1-6 would be a good choice to use for any additional calculations or quantitative analysis. One goal of PCA is to reduce the dimensionality of the input data set to allow it to be visualized in two or three dimensions. The more variance that is captured in the first two or three components, the more representative these 2D and 3D plots will be of the full data set. Figure 2.6 shows that PCs 1 and 2 together explain a little less than 50% of the total variance in* ***X***, *and adding PC3 raises the variance explained to ∼60% (the exact numbers can be found in ‘explained’). This is sufficient that plotting the data in two or three dimensions will give a reasonable representation of the information in* ***X***, *as the first PCs encompass the features that contribute most to differentiating the data elements from each other. However, it should be borne in mind that a non-negligible amount of the variance is not captured by PCs 1-3, and so some features that play lesser – but potentially still important – roles in distinguishing the data elements will be missing from these low-dimensionality representations*.

#### 12. Plot the PCA results in two dimensions

Plot the scores for the 201 elements (compounds) in the set with respect to Principal Components 1 and 2.

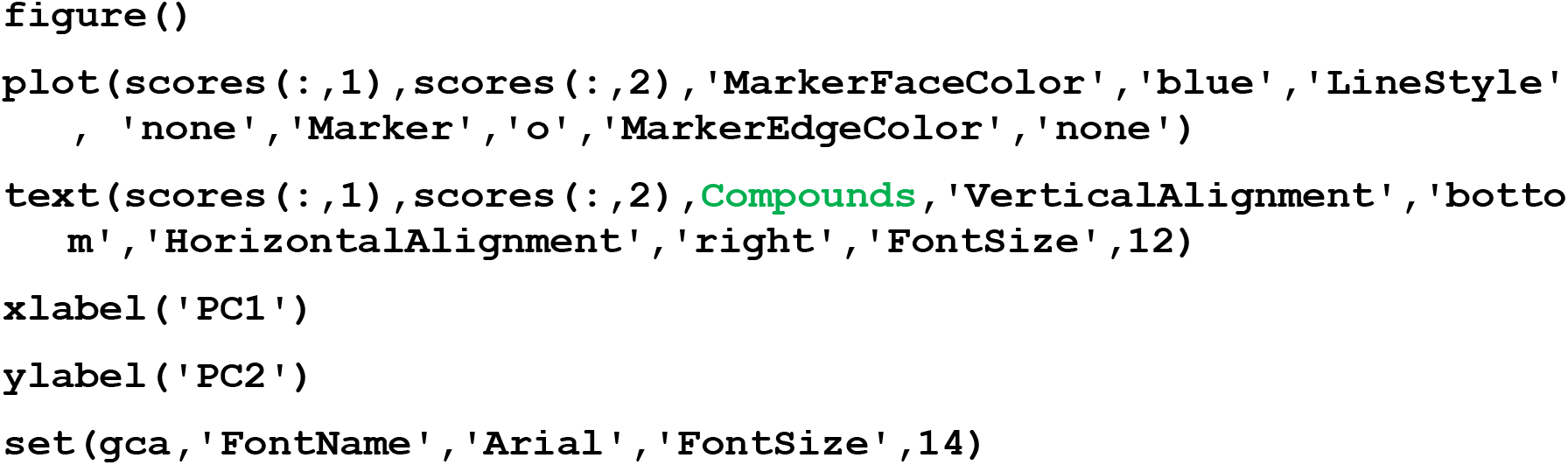

> *In the above command, the terms* `scores(:,1)` *and* `scores(:,2)` *specify to plot the coordinates (scores) of the 201 compounds with respect to PC1 (the data in column 1 of ‘scores’) and PC2 (the data in column 2 of ‘score’s). The elements could be plotted with respect to any pair of PCs by substituting the desired PC numbers for ‘1’ and ‘2’ in the command*.

A new Figure window will appear that contains the plot shown below (**Figure 2.7**).

> *This window can be brought to the front using the ‘Windows’ menu at the very top of your screen*.
>
> Figure 2.7: The 201 compounds from **X** plotted by their coordinates (scores) with respect to PCs 1 and 2. Each data symbol represents a different compound, plotted at coordinates (*PC1 score, PC2 score*). The numerical values of the PC scores can be found in ‘scores’, by looking at the cells in the row corresponding to the compound of interest (the rows being ordered as in **X**) and in the columns corresponding to PCs 1 and 2 (i.e. columns 1 and 2). The numbers that appear on the plot are the compound identifiers. The closer compounds are in this PC space, the more similar they are with respect to the 139 descriptors in **X**. 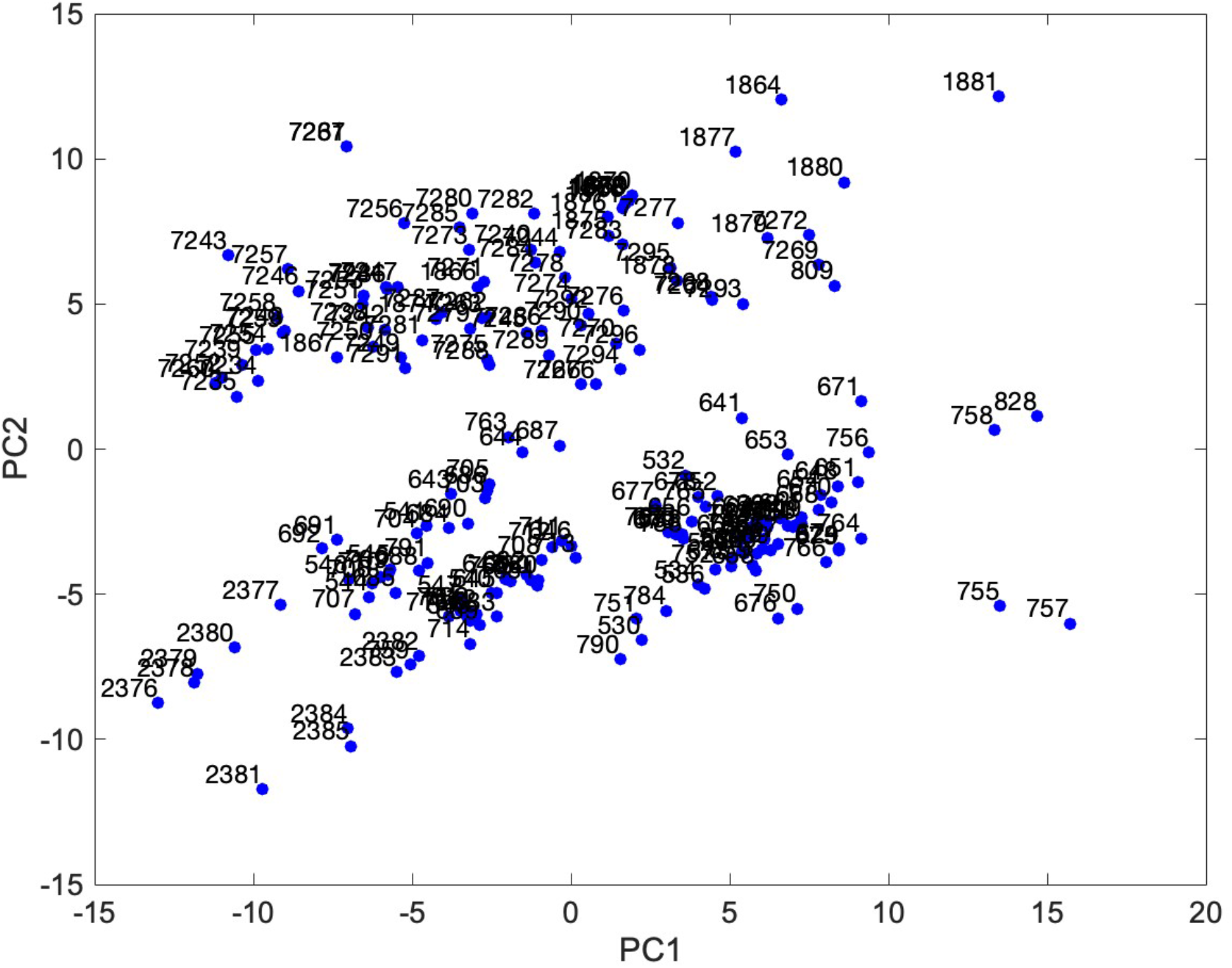
>
> *The data in Figure 2.7 would typically be plotted omitting the element labels (i.e. the compound ID numbers) to avoid making the figure too crowded. This would be done by omitting line 3, that starts ‘text*…*’ from the above command. We have included the compound IDs here to illustrate as clearly as possible the relationship between the compounds and their PC scores*.
>
> *For information on how to label the data points by color, for example to indicate by some other quality such as chemotype or membrane permeability, see Step 13 in Basic Protocol 3*.

#### 13. Plot the PCA results in three dimensions

Plot the scores for the 201 compounds with respect to Principal Components 1, 2 and 3.

~~~
figure()
scatter3(score(:,1),score(:,2),score(:,3),’MarkerFaceColor’,
  ‘blue’,’Marker’,’o’,’MarkerEdgeColor’,’none’)
xlabel(‘PC1’)
ylabel(‘PC2’)
zlabel(‘PC3’)
set(gca,’FontName’,’Arial’,’FontSize’,14)
~~~

A new Figure window will appear containing the plot shown in Figure 2.8.

> Figure 2.8: The 201 compounds from **X** plotted by their coordinates (scores) with respect to PCs 1, 2 and 3. Each data symbol represents a different compound, plotted at coordinates (*PC1 score, PC2 score, PC3 score*). The numerical values of the PC scores can be found in ‘scores’, by looking at the cells in the row corresponding to the compound of interest (the rows being ordered as in **X**) and in the columns corresponding to PCs 1, 2 and 3 (i.e. columns 1-3). Compared to Figure 2.7, this 3D representation provides a truer (but still incomplete) visualization of the relationships between the compounds, because PCs 1-3 capture somewhat more of the total variance in the compounds’ descriptor values than do just PCs 1 and 2 (see Figure 2.6). 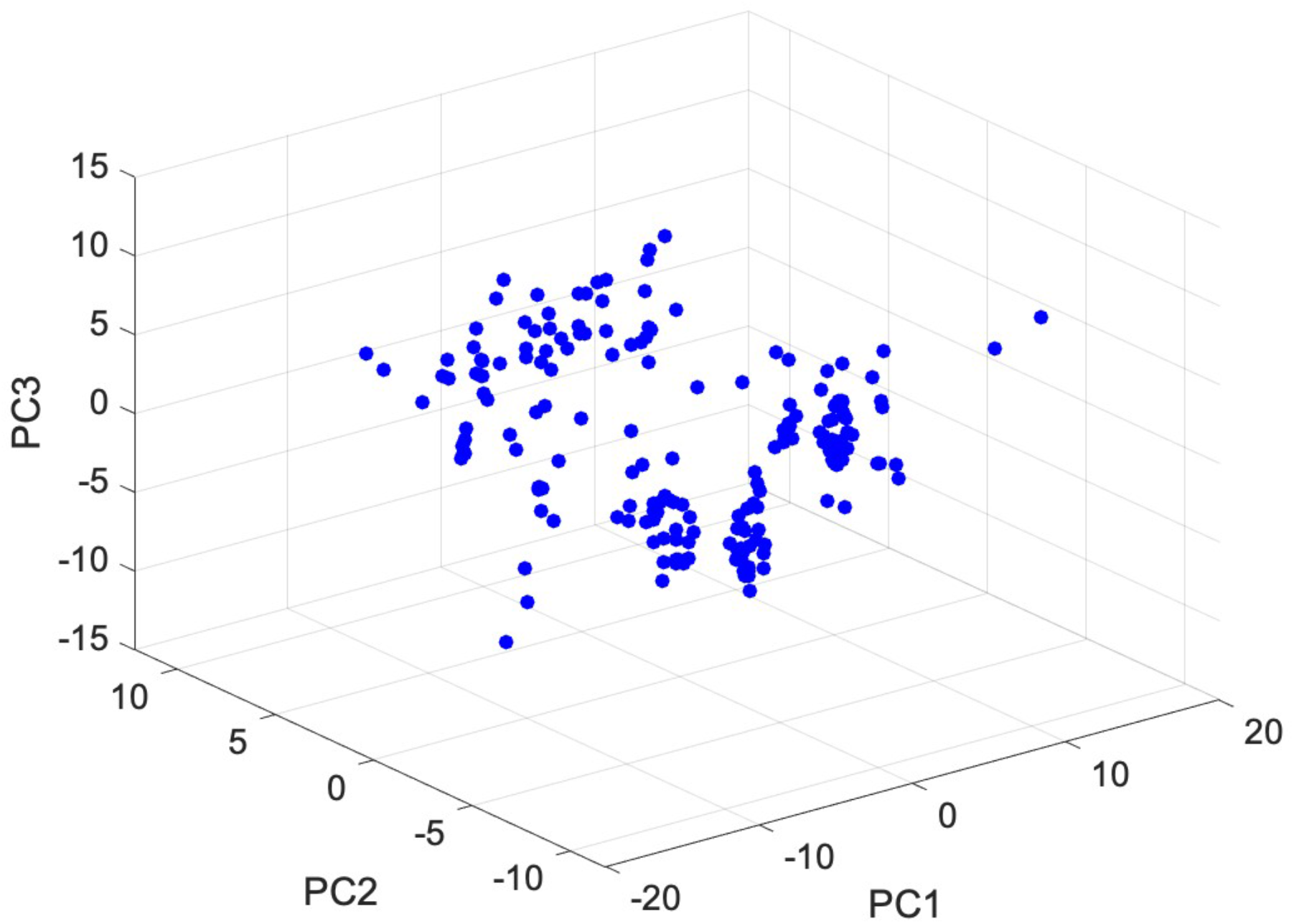
>
> *For simplicity, we have omitted the compound ID numbers from* ***Figure 2.8***. *To display them, include the following additional line in the command for Step 13:*

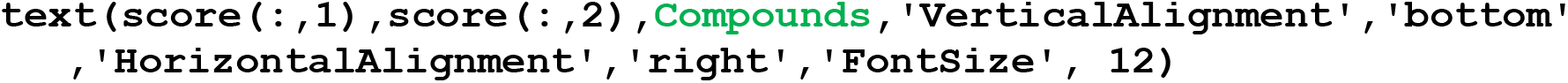

> *3D scatter plots such as* ***Figure 2.8*** *can be rotated in real time, for example to better view the data or to choose the optimal angle to use in a figure, by selecting the ‘3D rotation’ option from the ‘Tools’ menu at the top of the Figure window*.
>
> *The format and appearance of 3D scatter plots can be modified either by entering commands into the Command window or by using the menus at the of the Figure window. For more information type ‘doc scatter3’ in the Command window*.

#### 14. Create a biplot of the top 30 descriptors in PC1

> *A biplot is commonly used to visualize the results of PCA and related methods. It comprises a scatter plot of the scores for each element in the set with respect to PC1 and PC2, as described above in Step 12, but with the orthonormalized values of the loadings (coefficients) for each descriptor in PCs 1 and 2 superimposed on the same axes. A biplot thus displays, in a single plot, information both on how the elements are distributed in PC space, and which descriptors contribute most to PCs 1 and 2 and thus are most influential in defining the PC space*.

To plot only the **M** descriptors having the highest loadings (absolute values) for PC1, enter the following, where M is replaced with a number that represents the number of descriptors you wish to plot. Here, we use **M** = 30, to plot the 30 descriptors with the largest coefficients with respect to PC1.

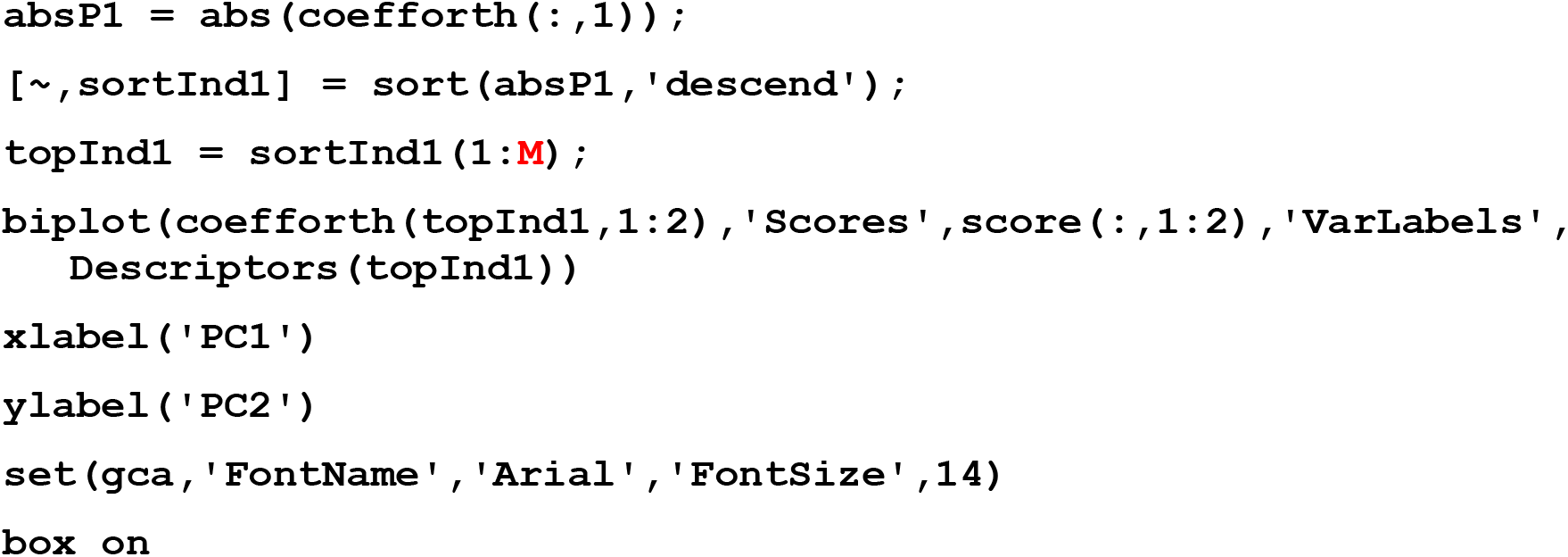

A new Figure window will appear containing the plot in Figure 2.9.

**Figure 2.9:**
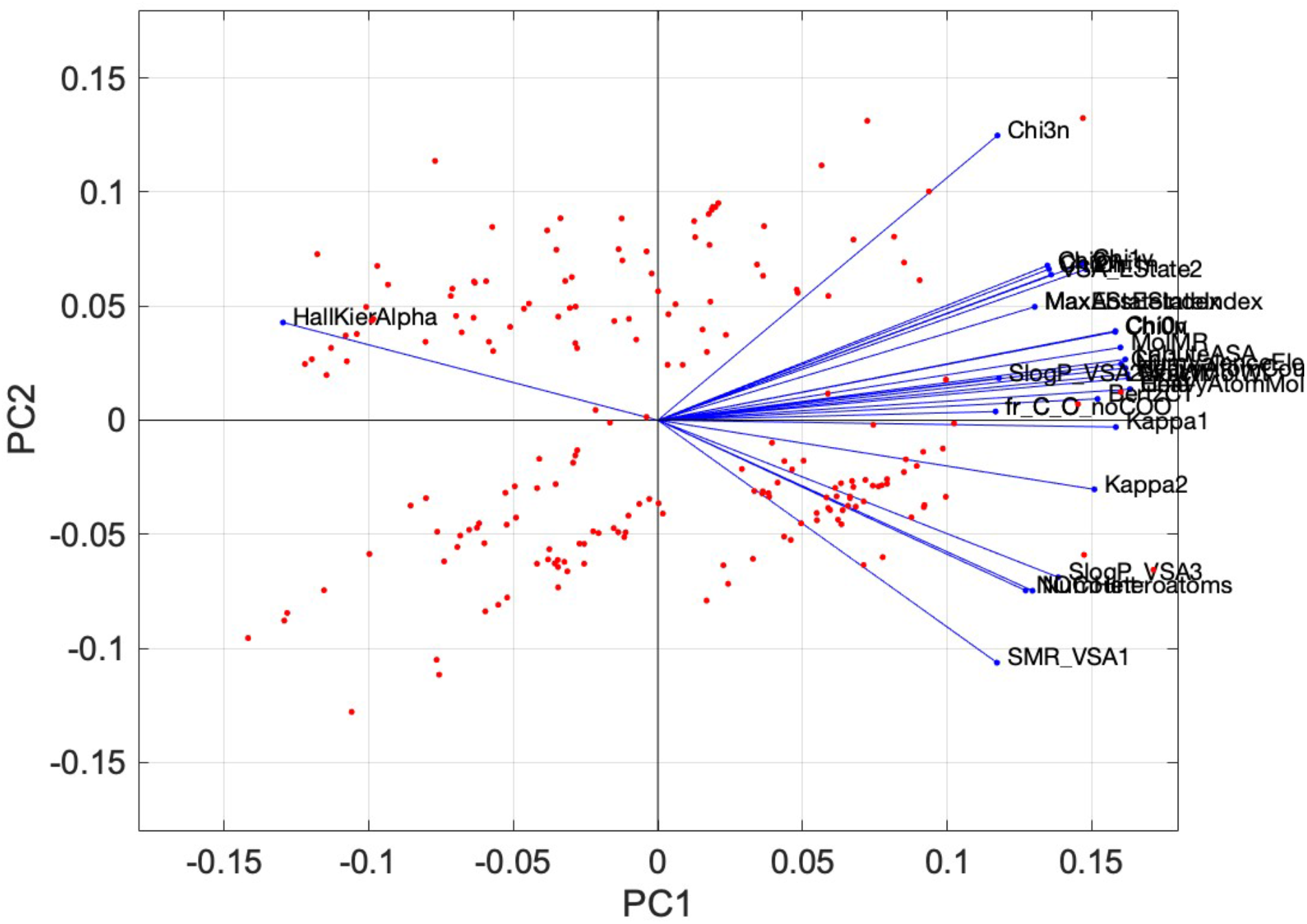
The 201 compounds from **X** plotted by their coordinates (scores) with respect to PCs 1 and 2 (red dots). Each data symbol represents a different compound, plotted at coordinates (*PC1 score, PC2 score*), exactly as in Figure 2.6 but with the compound ID numbers omitted. Superimposed on the same axes are the (ortho)normalized values of the coefficients for the 30 descriptors that make the greatest contribution to PC1. The higher the absolute value of the PC1 coordinate for the end of a given blue line, the more the descriptor influences that PC.

##### Interpreting a biplot

> *The magnitude of the contribution that each descriptor makes to PC1 is given by the x-axis (i.e. PC1 axis) coordinate for the blue data point that marks the far end of the blue line for that descriptor. This coordinate can be found by drawing a vertical line from the blue terminal data point to the PC1 axis and reading off the value. For example, the descriptor Kappa2 has an x-axis coordinate of approximately 0.15, indicating that the coefficient for this descriptor with respect to PC1 has an (orthonormalized) value of ∼ 0.15. Similarly, the coefficient for the descriptor HallKierAlpha, to the left of the plot, with respect to PC1 has a value of ∼-0.13*.
>
> *The values of the coefficients in PC2 are given by the y-axis (i.e. PC2 axis) coordinate for the blue data point that marks the far end of the blue line for each descriptor. These values can be found by visualizing a horizontal line from the terminal blue data point to the PC2 axis. Thus, the descriptor Chi3n has a coefficient in PC2 of ∼0.125, while Kappa2 has a coefficient in PC2 of approximately -0.25*.
>
> *Descriptors with a positive coefficient in PC1 are positively correlated with the PC – that is, a higher value for that descriptor will tend to confer a higher value for PC1. Descriptors with a negative coefficient, such as HallKierAlpha with respect to PC1, are anticorrelated with the PC, indicating that the higher the value of that descriptor for a given compound the lower the value of PC1. However*, <u>*when plotting a biplot the signs of the coefficients plotted can sometimes become switched as a group, so that coefficients with positive values terminate at negative values on a given PC axis and* vice versa. *Therefore, to know the sign of a coefficient it is necessary to look at the data variable ‘coefforth’ rather than reading off a biplot*.</u>
>
> *Like any graph, all of the information in a biplot can also be found in the data tables used to plot it (i.e. in ‘scores’ and ‘coefforth’*). *The biplot visually illustrates that the descriptors that contribute most strongly to PC1 are the cluster of descriptors at the far right of the plot, which all have coefficients in PC1 that have an absolute value of > 0.15. Moreover, several of the descriptors cluster close together, with only small angles between them, indicating that these descriptors are highly covariant. This high covariance can be confirmed by looking at the PCC values in the data variable ‘C’ that was created in Step 7. Examination of the properties of the descriptors that contribute most to PC1 could provide some insight into what over-arching feature of the compounds PC1 is capturing. Assigning functional labels to PCs is, however, an interpretation, rather than an actual result of PCA*.

#### 15. Create a biplot of the top 30 descriptors in PC2

Plot the **M** descriptors having highest coefficient (absolute value) in PC2. Here again, we have chosen **M** = 30 (**Figure 2.10**):

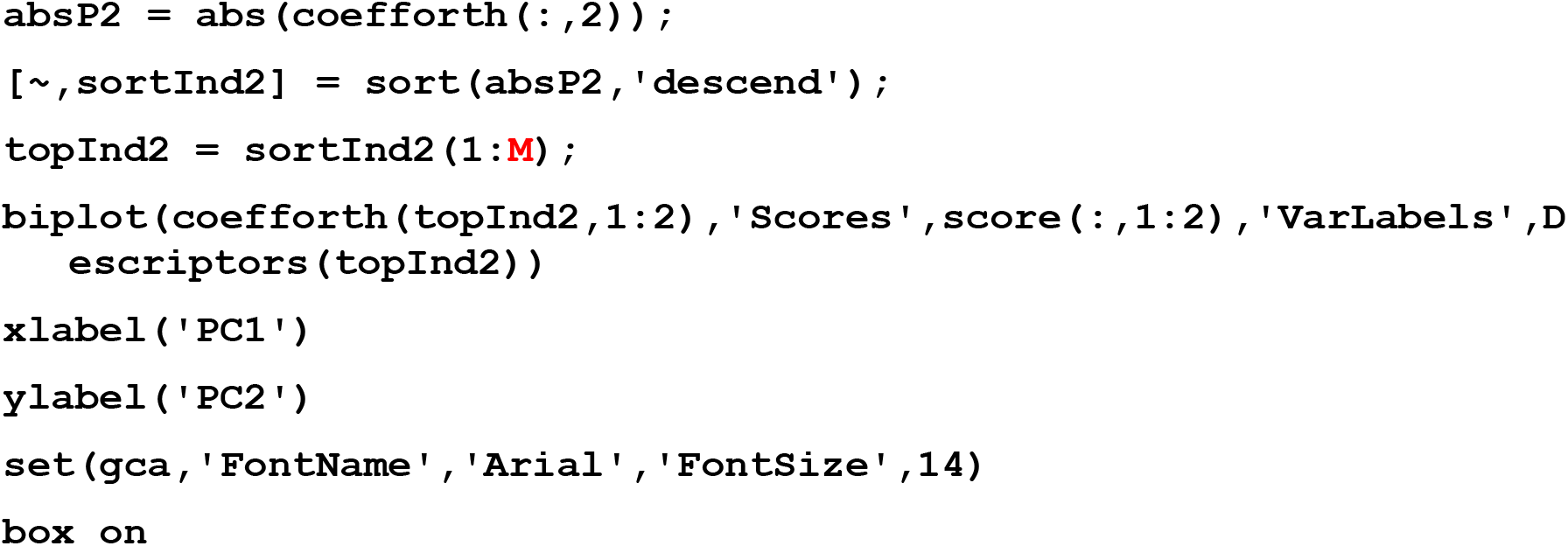

**Figure 2.10:**
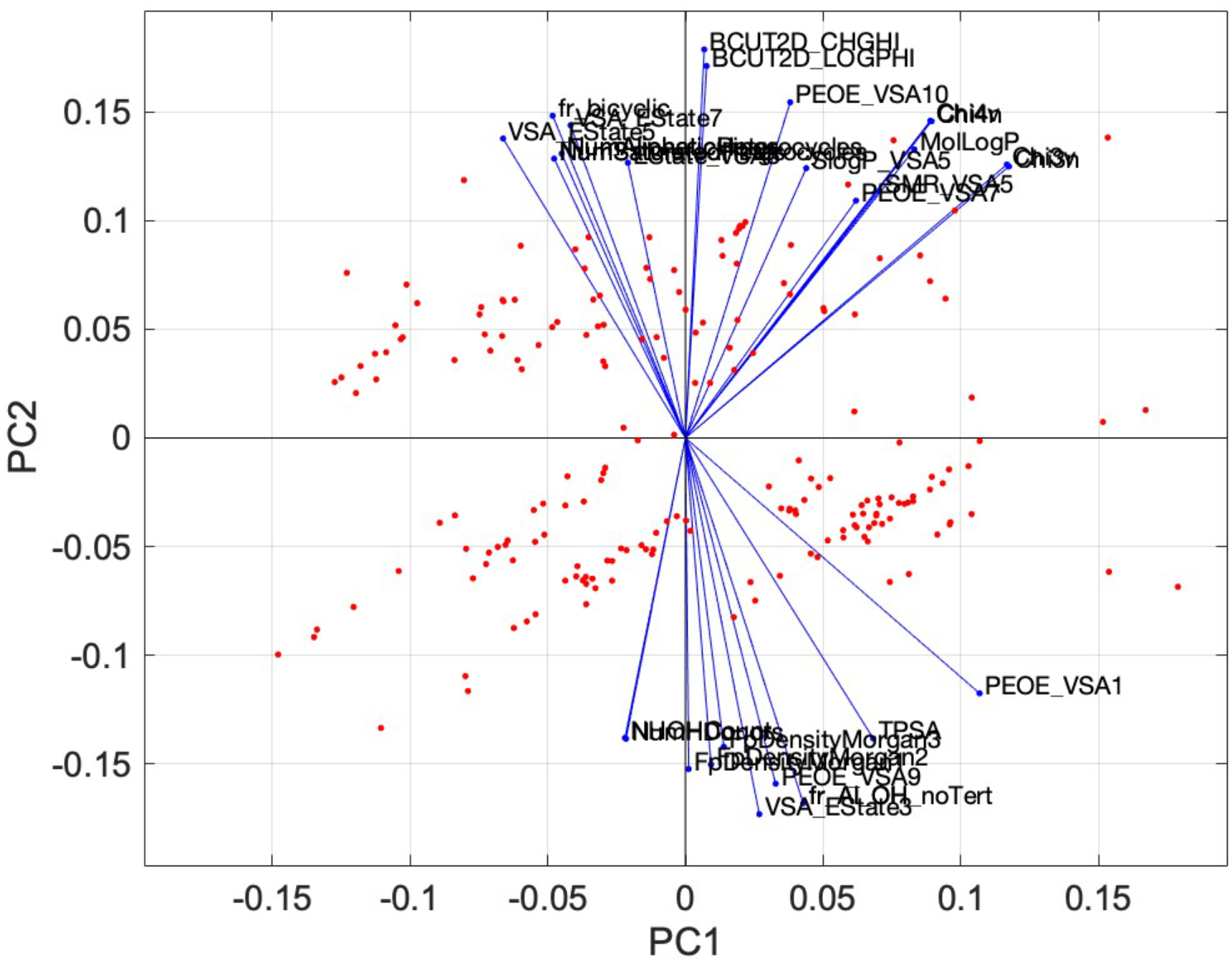
The 201 compounds from **X** plotted by their coordinates (scores) with respect to PCs 1 and 2 (red dots). Each data symbol represents a different compound, plotted at coordinates (*PC1 score, PC2 score*), exactly as in Figure 2.6 but with the compound ID numbers omitted. Superimposed on the same axes are the (ortho)normalized values of the coefficients for the 30 descriptors that make the greatest contribution to PC2. The higher the absolute value of the PC2 coordinate for the end of a given blue line, the more the descriptor influences that PC.

> *The descriptors that contribute most to PC2 are different from those that were most influential in PC1, shown in Figure 2.29. This is expected as PCs1 and 2 comprise orthogonal features of the data in X, and therefore it is not expected that any individual descriptor could be highly correlated with both PCs (or any two PCs)*.

#### 16. Identify the descriptors that most influence the first three PCs

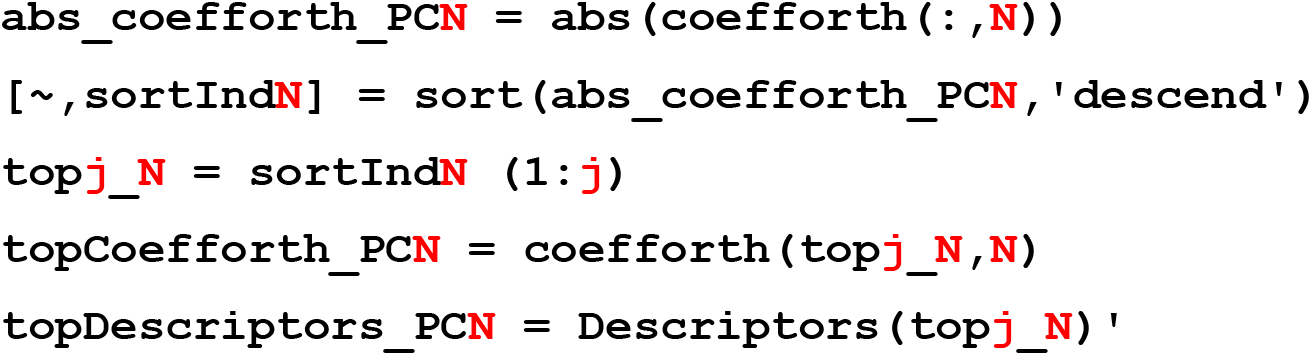
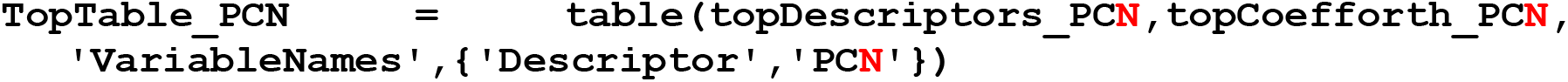

> *In the above command, the input* ***N*** *is the number of the PC for which you wish to identify the* ***j*** *descriptors that have the highest (absolute value) coefficients. Here, we performed the command three times, with N set at 1, 2, or 3, respectively, and with* ***j*** *= 10 in all cases. The results were manually collated into the table shown below. In the table, the signs of the coefficients were restored to indicate whether the value of the given PC increases or decreases as the value of the descriptor increases*.

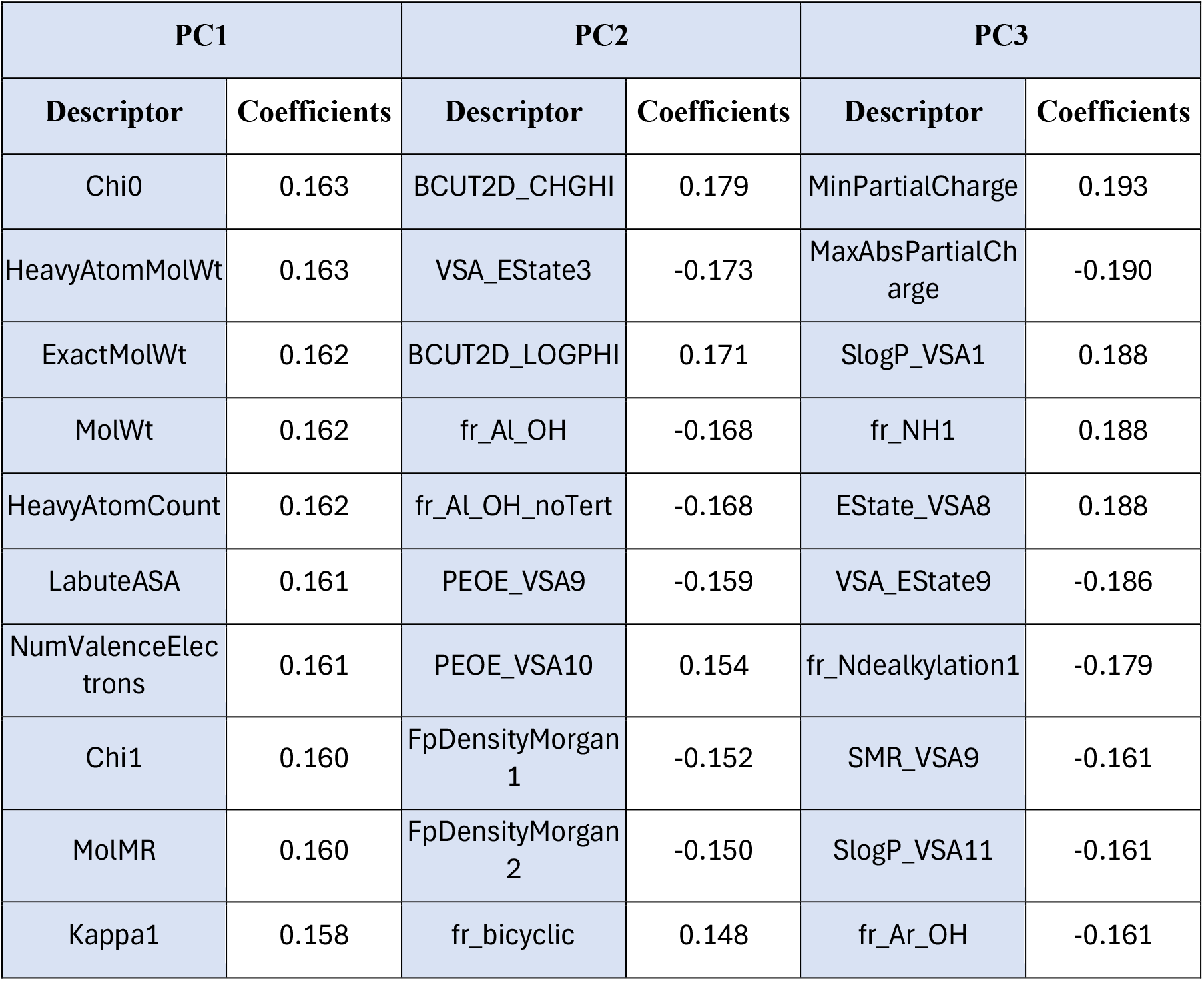

#### 17. Identify outlier compounds using the Hotelling’s T-squared statistic

Identify the outliers in the data set – i.e. those compounds that are most dissimilar to the bulk of the set – by ranking the compounds by their Hotellings T-squared values, which are contained in the data variable ‘tsquared’ created in Step 9.

> *As described in the Introduction to Basic Protocol 2, the Hotelling’s T-squared statistic is the distance in* n*-dimensional PC space of each element in the data set (i.e. each compound) from the center of mass of the data set. Thus, the compounds with the largest Hotelling’s T-squared value are the most dissimilar from the main body of compounds*.

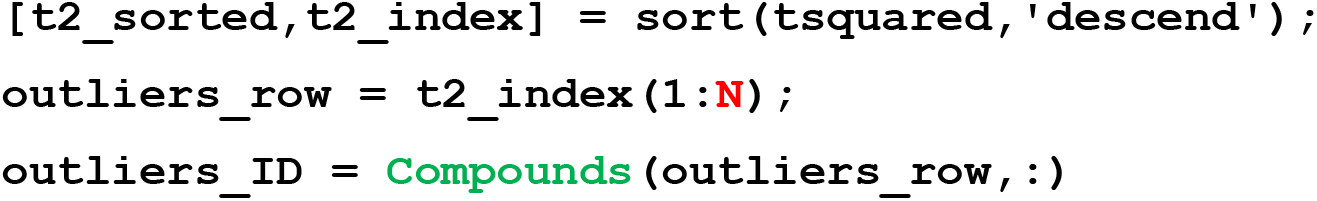

> *The ‘sort’ command sorts the values in ‘tsquared’ in descending order, creating two new data variables. The values in ‘tsquared’, sorted in order from highest to lowest, are placed in a* m *x 1 data variable called ‘t2_sorted’. A second* m *x 1 data variable, called ‘t2_index’, contains the row numbers in ‘tsquared’ re-ordered to match the order of their corresponding values in ‘t2_sorted’ (i.e. the row number in ‘tsquared’ that contained the highest value will appear in the first position in ‘t2_index’, and so on)*.
>
> *The second command identifies the first* ***N*** *entries in t2_’index’, corresponding to the* ***N*** *most outlying compounds in the data set, and places them in a new data object called ‘outliers_row’. For example, to find the ten most outlying compounds in the Training Set, here we set* ***N*** *= 10. The variable ‘outliers_row” will thus contain ten numbers corresponding to the row numbers of these compounds (in tsquared, X, etc.), ordered starting from the greatest outlier*.
>
> *The third command identifies the compound names/ID numbers for these* ***N*** *outlier compounds from the string array ‘Compounds’, created in Step 4, and places them in a new string array called ‘outliers_ID’*.

The above command returns a list of the compound IDs for the top **N** (in this case top 10) outliers, as determined by PCA:

~~~
7282
2385
809
828
757
763
790
755
1881
7258
~~~

> *Alternatively, a full list of all 80 compounds in the Training Set, ordered starting from the one with the greatest Hotelling’s T-squared statistic, can be created by entering:*

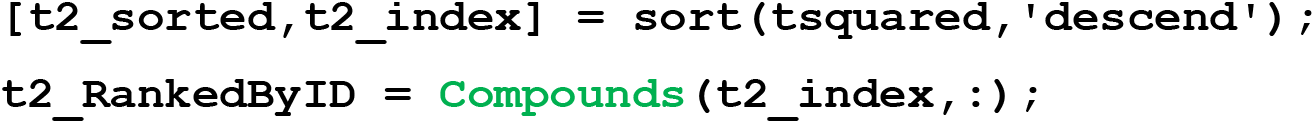

#### 18. Create a scatter plot to show how the compounds plotted in PC space break down into the different clusters identified by *k*-medoids clustering

> *In Basic Protocol 1c we showed how the compounds in this data set could be clustered into k subsets by the method of k-medoids clustering. One way of visualizing the results of such clustering is to plot the compounds in PC space, as per Figure 2.6, and color-code them according to the results of the clustering, using the following code:*

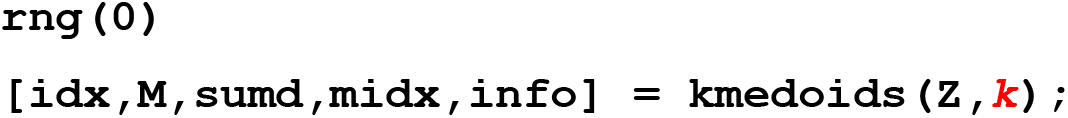
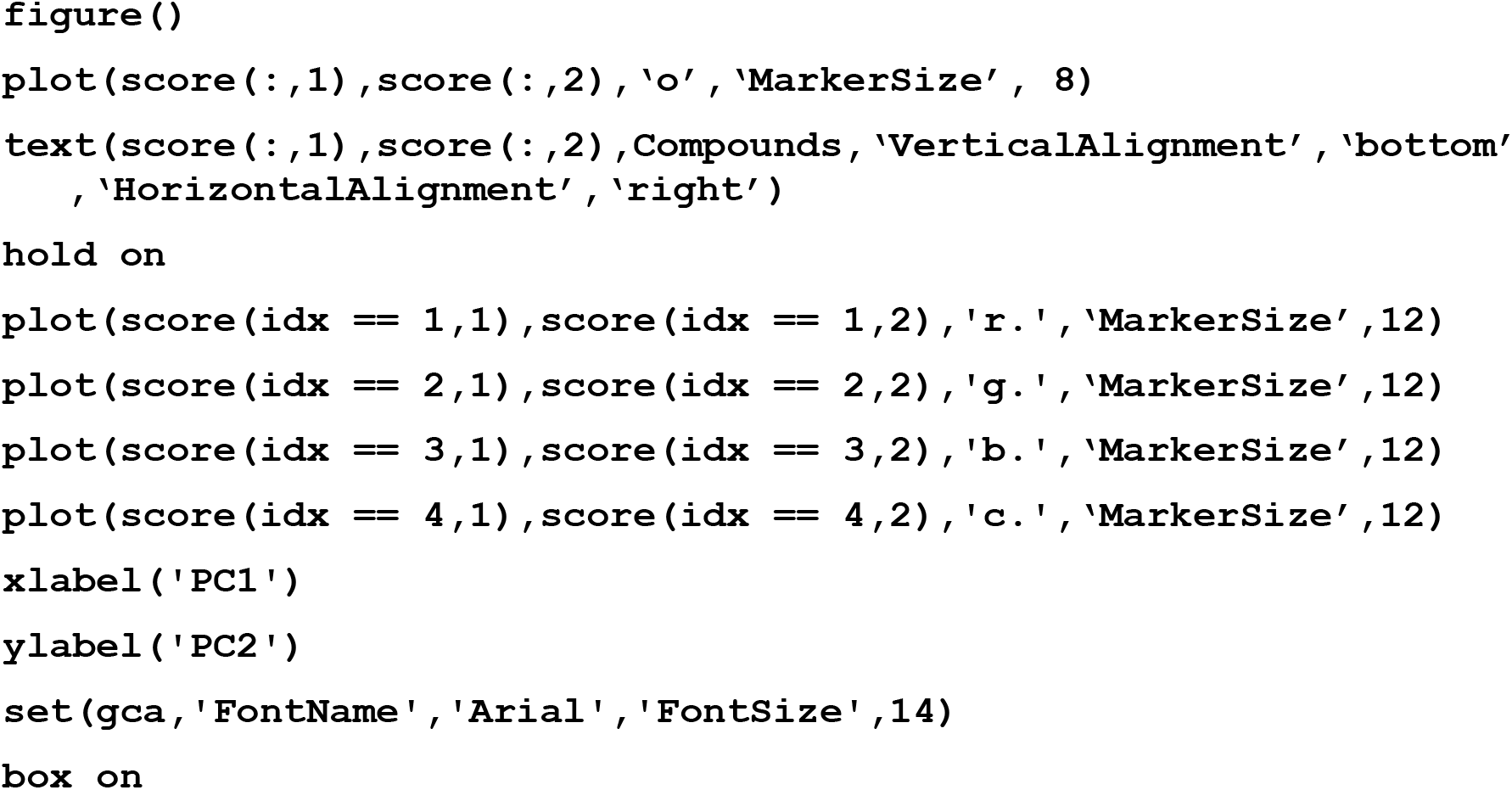

> *The variable* **k** *is the desired number of clusters – in this instance, the desired number of compounds to be selected for the representative subset. For the purpose of this protocol, we will use* **k *= 4***, *to cluster the compounds into 4 groups and identify the compound most representative of each cluster. For additional detail on* k*-medoids clustering, see Basic Protocol 1c*.
>
> *The method of* k*-medoids, like many clustering algorithms, starts by randomly selecting initial medoids (or cluster centers). Since this selection is random, different runs may start with different initial medoids, leading to different clustering results. By setting the seed (rng(0)), you will get the same result every time you run the code with that seed*.
>
> *The above command contains instructions for creating four different colored subsets of data points, based on our choice of* **k** *= 4 for the* k*-medoids clustering in line 1 of the above command. In the event it is desired to create and then color a different number of subsets, this can be done by increasing or decreasing the number of lines in the command of format:*

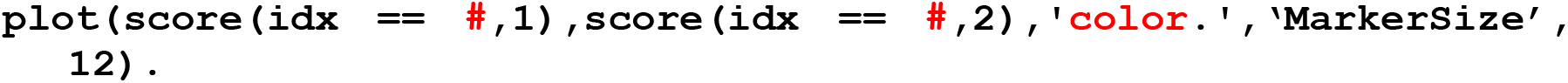

> *where each such line corresponds to the line in ‘idx’ containing the* x *and* y *coordinates for the subset of points to be colored, and specifies a different color*.

**Figure 2.11:**
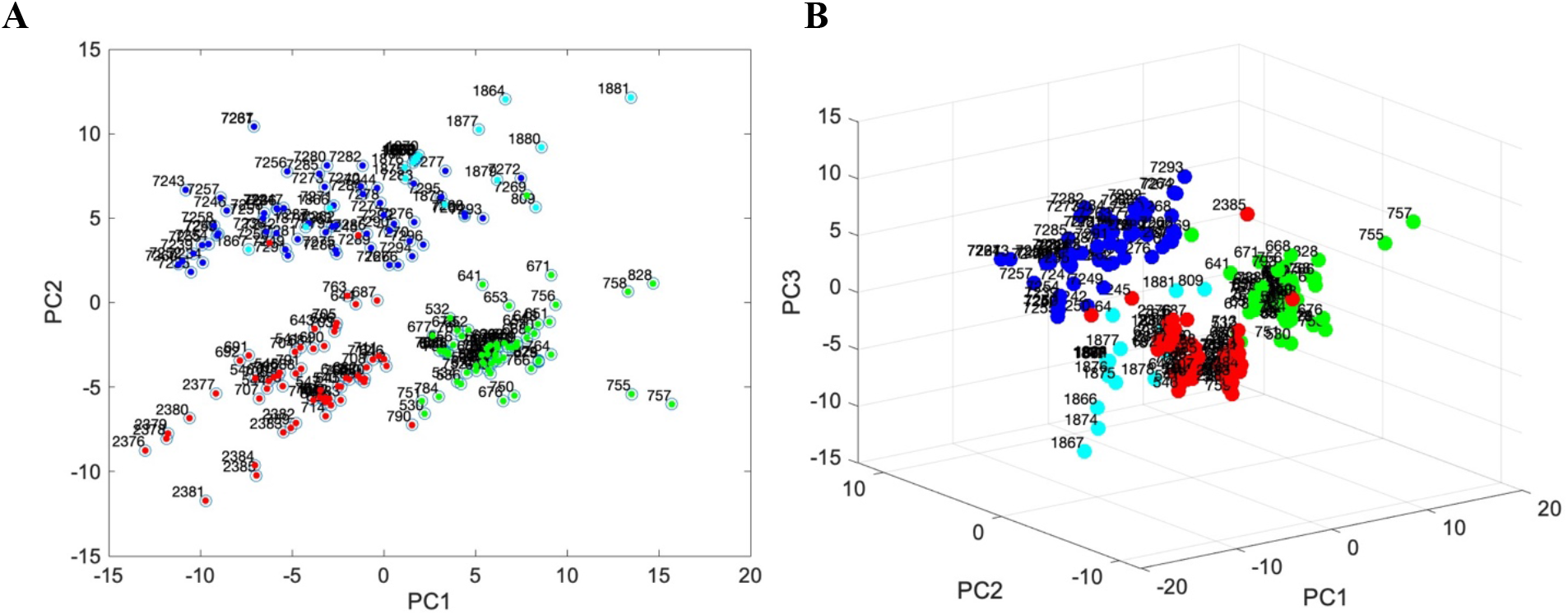
**(A)** The 201 compounds from **X** plotted by their coordinates (scores) with respect to PCs 1 and 2, as in Figure 2.7. Each data symbol represents a different compound, plotted at coordinates (*PC1 score, PC2 score*). The compounds that belong to clusters 1-4 from *k*-medoids clustering (Basic Protocol 1c, Steps 7 and 8) are colored red, green, blue, and cyan, respectively. The numbers that appear on the plot are the compound identifiers. **(B)** The compounds from (A) but plotted with respect to PCs 1-3.

A new Figure window will appear containing a plot similar to that shown in **Figure 2.11A**.

***Figure 2.11A*** *shows that division of the compounds into four groups by k-medoids clustering resulted in subsets that occupy distinct regions of property space, consistent with the notion that the method accurately clustered the compounds with respect to the descriptors provided. Although colors in certain regions of the plot appear to overlap, particularly the cyan and blue subsets, examination of the data plotted with respect to the first three PCs (****Figure 2.11B****) shows that these compounds are in fact fully separated in* n*-dimensional PC space*.

#### 19. Save your workspace as a .mat file

> *This file can be reopened to give a MATLAB workspace identical to the state that was saved*.

### Understanding the Results of Basic Protocol 2

#### What PCA does

PCA identifies patterns and trends in complex, multi-parameter data sets. The underlying notion is that to the extent two or more descriptors are covariant with each other, they are capturing the same information about the elements in the data set. Descriptors are rarely 100% covariant (i.e. PCC = +1 or -1), and it is only the covariant portion of the information from each that is redundant. PCA, an unsupervised ML method, simply looks for the greatest amount of covariant information in the input matrix *X* and calls this PC1. This process is equivalent to plotting all the data in *X* on a set of Cartesian axes, one axis for each of the *n* descriptors in *X*, and then drawing a straight line through the origin of the resulting *n*-dimensional cloud of data points in the direction that captures the most variance. Generally speaking, PC1 will pass through the longest axis of the data cloud. PC2 is a second new axis, orthogonal to (i.e. having zero covariance with) PC1, that captures as much as possible of the remaining variance that PC1 did not capture (i.e. passes through the second-longest dimension of the *n*-dimensional data cloud), and so on for PCs 3, 4, etc. Because each PC axis is mutually orthogonal to all the others, the information contained in each is uncorrelated with and thus independent of the information contained in any other PC. Each PC therefore reflects a distinct, nonredundant subset of information from *X*.

Examining the data in the variable ‘C’ created in Step 7, a portion of which is shown in **Figure 2.5**, shows a number of off-diagonal squares with high covariance, defined as a PCC value close to +1 or -1. For example, **Figure 2.5** shows that the count of NH + OH groups in a compound substantially covaries with molecular weight (PCC = 0.71). In other words, the bigger a compound is, the more NH + OH groups it contains, on average. As another example, this time of an anticorrelation, higher numbers of hydrogen bond donors are to some degree associated with lower lipophilicity as measured by MolLogP (PCC = -0.53). The existence of substantial covariance between the descriptors, as shown in the many substantial PCC values in ‘C’, indicates that there is a considerable redundancy among the descriptor set. This finding is consistent with the result from the Pareto plot in **Figure 2.6** showing that PCA was able to achieve a substantial degree of dimensionality reduction, from the 139 descriptors of the original data set to the 6 PCs required to capture 80% of the total information in **X** or the 10 descriptors required to capture >90%.

#### What can PCA be used for?

PCA can be used for several purposes. Chief among them is simply to reveal patterns of similarities and differences among the elements in a set with respect to a collection of provided properties or characteristics. By exploiting covariance among the input property values to reduce the dimensionality of the data, PCA can often allow the relationship between the data elements to be meaningfully visualized by plotting the data using just two or three principal component axes. Visualizing the data in this way can reveal which elements in the data set cluster most closely together in PC space, allowing identification of similarities and differences that would be difficult to discern from the original, high dimensionality data set. For example, in an analysis of this kind, data on measured signaling properties or expression of cell surface markers for a set of otherwise similar cells displaying a variety of disease phenotypes might reveal that certain diseases share similar profiles.

Another common use of PCA is to determine which properties are characteristic of a subset of elements in a data set that share a particular quality or behavior. An example of an analysis of this sort would be the use of PCA to identify which physicochemical properties are characteristic of the orally bioavailable compounds among a larger set of molecules. PCA addresses such questions by enabling comparison of the elements in the data set with respect to a large number of potentially relevant properties or characteristics. To the extent the elements showing the behavior of interest cluster together when plotted on PC axes, PCA will identify which properties contribute most to differentiating these elements from the larger set. Note that supervised ML methods, including PLSDA (described in Basic Protocol 3) provide other, arguably more direct, approaches to addressing this kind of question. PCA is nonetheless often used in this way.

#### Interpreting PCA scores

In PCA, the scores (calculated here in Step 12) are simply the coordinates of the various elements in the data set with respect to the different PC axes. If the scores were plotted on a set of *n* Cartesian axes corresponding to all *n* PCs, then the resulting cloud of data points in *n*-dimensional space would look identical to the result of plotting the compounds using the original *n* descriptors as axes. The difference is that, relative to the original descriptor axes, the PC axis set is rotated about the origin such that PC1 passes through the longest dimension of the data cloud, PC2 through the second-longest dimension that is orthogonal to PC1, etc. Importantly, because PCA gathers the largest portion of the variance in PC1, and decreasing amounts in each successive PC, after the first few PCs the amount of variance remaining to be captured typically becomes very small even when *n* is large. Consequently, the original data cloud which comprises all the information provided can often be represented by plotting the data with respect to only the first few PC axes with minimal loss of content, allowing the information to be manipulated, visualized, and analyzed much more simply compared to the original *n*-dimensional data set.

Plotting the scores for the first two or three PC axes reveals how the data points cluster in PC space, providing an approximation of how they cluster in the original *n*-dimensional property space and thus revealing which elements are more similar or dissimilar to others with respect to the *n* original properties. Examining which data points cluster together or lie far apart therefore reveals patterns of similarity and difference in the data that might be impossible or very difficult to visualize with the PCA transformation. The closer together two data points are in PCA space, the more similar are their characteristics with respect to the entire set of *n* original descriptors.

The key point in interpreting PCA scores is to consider what percentage of the total variance in *X* is captured by the first two or three PCs. This information is contained in the Pareto plot shown in **Figure 2.6**. If a high proportion (say >70 or 80%) of the variance is captured by these early PCs, then a 2D or 3D scatter plot should faithfully reflect the patterns and relationships that exist between the data elements. However, if the first two or three PCs collectively capture only a small fraction of the total variance in *X*, looking only at plots with axes of PC1-3 may give a misleading picture. While these early PCs will illustrate the most influential factors that distinguish the data elements from each other, other factors that are almost as important are being ignored.

In cases where >3 PCs are required to capture sufficient variance to make the analysis reasonably representative of the whole data set, the relationships can nonetheless be visualized by plotting the scores on a set of 2D plots systematically covering the desired number of PC axes. This idea is shown, for a simple illustrative case involving only 3 PCs, in **Figure 2.12A** and **B**. Panel A shows four hypothetical data points plotted in 3-dimensional PC space, which have positions that approximate a tetrahedral spatial arrangement. Panel B shows that the same information can be fully captured by a series of 2D plots that show the data points plotted with respect to all possible pairs of PC axes. Each 2D plot can be considered as the arrangement of the data points that would be observed upon looking at the 3D plot in (A) through a different face of the cube. Panel C shows how this methodology can be extended to any number of PC axes (the case for 4 PCs is illustrated), with each 2D plot again showing how the cloud of data points are arranged when the multidimensional data cloud is viewed through a different face of the hypercube defined by the axes. An example of such a treatment from our own work can be found in Figure 4 of reference *Viarengo et al*., *2021*. Representations of this kind can allow the representation of PCA results for which >3 PCs are required to capture the bulk of the variance in the data, ensuring that important relationships that are apparent only in higher-numbered but still significant PCs are not overlooked.

**Figure 2.12.**
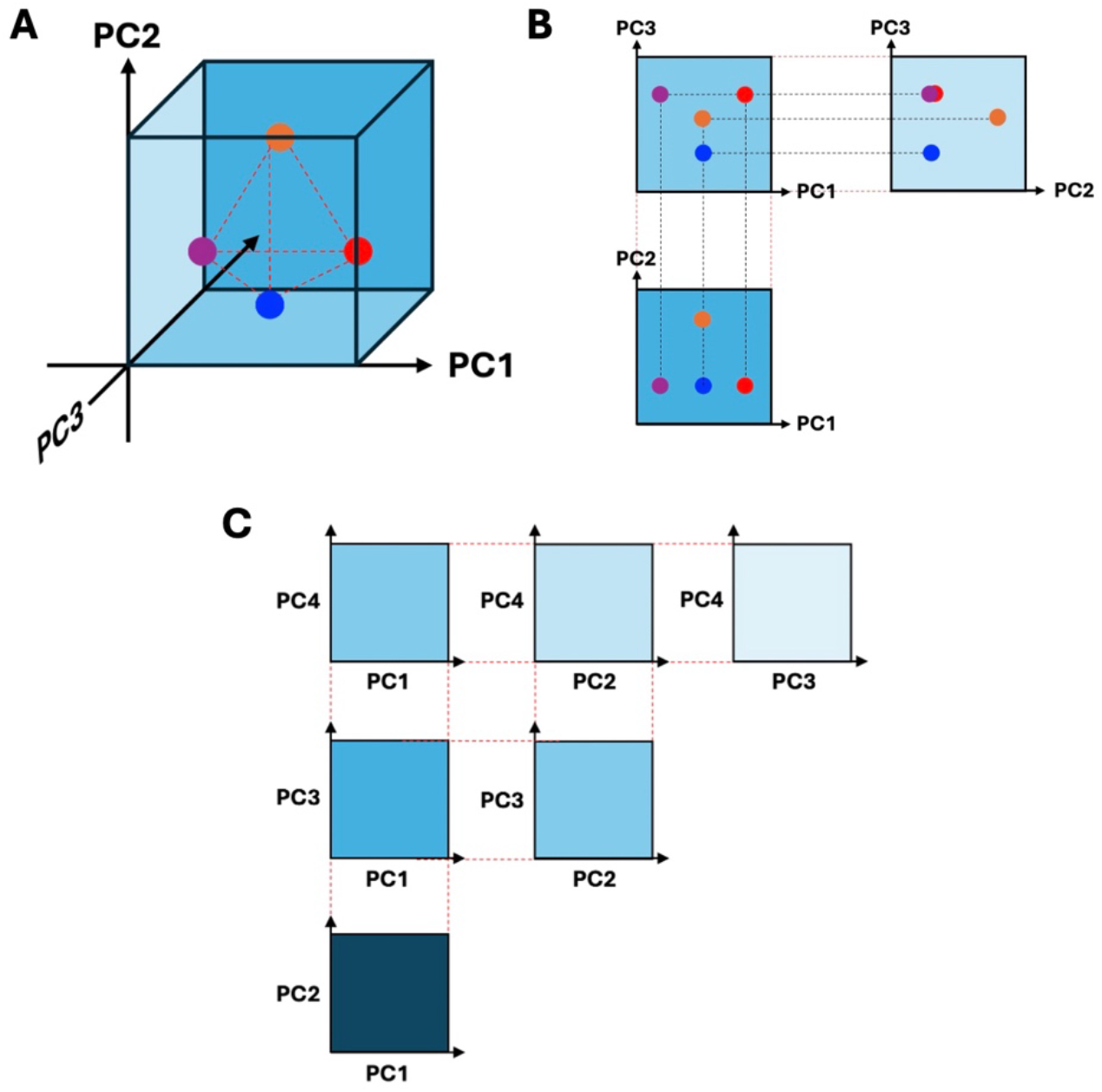
Representation of scores plotted in multi-dimensional PC space using a series of 2D plots. **A**. Hypothetical 3D plot of four tetrahedrally-arranged data points (colored circles), based on their scores with respect to PCs 1, 2 and 3. **B**. Set of 2D projections showing how the data points in (A) appear when viewed through the different faces of the cube that correspond to PC1 vs PC2 (front face of cube), PC1 vs PC3 (top face of cube), and PC3 vs PC2 (left face of cube, rotated 90 °C so that PC2 is the horizontal axis). **C**. Illustration of how the same principle can be applied to data plotted on an arbitrary number of PC axes, using as an example a data set plotted in PCs 1-4. Each 2D plot represents the view upon looking at the *n*-dimensional PC space from a different orthogonal viewpoint.

#### Interpreting PCA coefficients

With regard to interpreting the coefficient of the descriptors that dominate the low-numbered PCs from the worked example shown in the above protocol, we decided that such a discussion would not materially add to the value of this article. The descriptors we included in the example were chosen arbitrarily, for convenience in illustrating the method, and do not reflect the thoughtful construction of an input data matrix that would be required for a scientifically meaningful analysis of the compounds in question (see ‘Summary Comments on PCA’ for a general discussion of considerations for descriptor selection). The data we used, though real, are intended only to serve as a model system. Instead, we address the question of interpreting PCA coefficients directly, in the following paragraphs.

Interpreting which of the properties in the original input data are the main score drivers for the first few PCs can be a subtle business. Each PC represents a non-redundant subset of information from the input data that is orthogonal to (i.e. has zero covariance with) any other information in the data set. By examining which of the original properties have large coefficients (absolute value) for a particular PC, it is sometimes possible to discern a unifying conceptual connection between the properties that gives insight into what over-arching feature is driving the scores in that PC. So, using a hypothetical example, consider performing PCA on a set of compounds with respect to their physicochemical characteristics, and finding that one of the PCs showed the following properties to have coefficients with the largest absolute values: number of hydrogen bond donors and acceptors, polar surface area, dipole moment, and cLogP. Since all of these properties report in some way on the polarity of the compound, it might be reasonable to conclude that PC1 reflects the overall molecular polarity. This inference could be useful if, for example, the compounds that share a desired behavior, such as passive membrane permeability, cluster together with respect to their positions on this PC axis, and provides a way of assessing whether other compounds fall in the appropriate polarity window to also have a prospect of being membrane permeable assuming any other necessary conditions are met (*Viarengo et al*., *2021*).

Placing meaningful interpretations on a collection of descriptors that contribute strongly to a given PC can be very difficult, however. One issue is how to decide on the cut-off for the magnitude that a coefficient must have for a descriptor to be considered important for a given PC. Considering only the two or three descriptors with the highest (absolute value) coefficients may inspire a somewhat different interpretation to that indicated when a larger number of high coefficient descriptors is considered. Another difficulty is that some descriptors that have high coefficients in one PC may also be significant contributors to other high-ranking PCs. Such a result is perfectly reasonable; a property such as molecular weight, for example, can have significant covariance with quite disparate molecular features, such as e.g. polar surface area and nonpolar surface area, that have very different implications for the behavior of the molecule. For these and other reasons, deducing which features represent the themes of the high scoring PCs is often a matter of subjective interpretation, especially for PCs beyond the first two or three where the variance comprising the ‘signal’ of relevant information tends to become small compared to background variance resulting from ‘noise’ in the data. Unless the thematic commonality between the descriptors that dominate a given PC is very clear, such interpretations should be considered as speculative hypotheses subject to further experimental validation rather than as firm conclusions, and in our experience should be limited to only the first two or three PCs.

We have found two approaches useful in helping discern the over-arching feature represented by a given PC. One is to directly consider the covariance between the high-coefficient descriptors. This can be done by taking just the descriptors that have high (absolute value) coefficients in a given PC and constructing a covariance table as described above in Step 7. Considering highly covariant descriptors as a group can sometimes suggest the feature they collectively describe. To the extent that a given PC is dominated by two or more such groups of mutually covariant descriptors, the overlap or intersection of the inferred features may suggest a theme. For example, a PC might hypothetically have high coefficients for properties such as high sp^2^ content and low sp^3^ content, etc. – likely to be highly anticorrelated for molecules of similar size – and might also have high coefficients for, say, number of rotatable bonds and (a low) number of rings, which again will be highly covariant with each other but perhaps a little less so with the aforementioned descriptor pair. In considering what kinds of molecules would have both a high sp2/sp3 ratio and less constrained bond rotations, one possibility is that the PC in question in fact reflects molecular flexibility/rigidity. Such a hypothesis might be checked by looking to see whether other descriptors relating to flexibility appear towards the top of the list when the descriptors are ranked by the absolute magnitude of their coefficient in that PC, or by redoing the PCA but now including additional descriptors that more directly reflect molecular flexibility. Alternatively, the interpretation may be taken as a hypothesis for separate experimental validation. When making these judgments, it is clearly vital to have a sound understanding of what property each relevant descriptor represents, so that appropriate common themes can be identified. An example of this approach to interpreting PCA results can be found in (*Viarengo et al*., *2021*).

A second approach to solidifying interpretation of PC themes is to consider the signs of the coefficients. For example, in the hypothetical example discussed in the previous paragraph, if molecular flexibility is the correct interpretation of the PC in question, then the coefficients in that PC should have opposite signs for number of rotatable bonds and sp^3^ content, both of which tend to confer increased flexibility, compared to sp^2^ content and number of rings which will tend to rigidify a molecule. Passing such a test does not prove that molecular flexibility is the correct interpretation of what information is represented by the PC, but failing it likely indicates that the interpretation is wrong.

#### Interpreting the Hotellings t-squared statistic

The Hotelling’s t-squared statistics calculated in Step 9 are the Euclidean distances of each element from the origin in PC space (Hotelling, 1931). A compound with a low t-squared value thus lies close to the ‘center of mass’ of the cloud of data points seen when the scores are plotted in PC space, which defines the origin of PC space, while those elements with large t-squared values lie towards the edge of the data cloud. The elements with highest t-squared values can thus be considered as outliers from the greater mass of compounds with respect to the properties included in the original descriptor set. In the worked example shown in the protocol, the Hotelling’s t-squared values are found in the data variable called ‘tsquared’ that created in Step 9.

Regarding the results of the analysis illustrated in the protocol, it is noteworthy that the six biggest outliers identified by clustering in Basic Protocol 1a, compounds 2385, 790, 755, 575, 828, and 809, also appear in this list of top ten outliers as determined from the results of PCA. It is expected that compounds that differ substantially from the majority of the set will be identified by both methods. However, we do not expect that the outlier lists should necessarily be identical. This is because the distances identified by the clustering reflect pairwise separations between compounds or groups of compounds. Consequently, a chain of data points that reaches far out into property space may nevertheless have each element connected to its nearest neighbor by a short pairwise distance. In contrast, the Hotellings t-squared statistic quantifies distance from the center of mass of the compounds, so the terminal elements in a far-flung chain of compounds will have a high score regardless of how close its nearest neighbors are.

### Summary comments on PCA

#### Performing PCA meaningfully

Performing PCA in MATLAB or other appropriate software, is technically straightforward. However, to obtain meaningful information it necessary to carefully consider (i) what data points and properties to include in the analysis, (ii) whether and how to normalize the data, and (iii) how to meaningfully interpret the results. How data points cluster in PC space depends entirely on what elements and properties are included in the analysis. Important features will be missed if the input data don’t include information on that feature. For example, in using PCA to ask a question such “what compound types have been successful as oral drugs?”, one might consider any number of different characteristics in the analysis, and the compound groupings that result will depend on which characteristics are included. If a relevant molecular property is not captured by the descriptors in the input data, nothing can be directly learned about it. PCA is therefore often best used as an iterative exercise, where different sets of potentially relevant properties are tested, and the results compared to see what patterns in the data are most robust and informative. In the above example, for example, PCA could be repeated while adding or eliminating selected properties or groups of properties from the input data set. Identifying a set of input properties that cause the oral drugs to cluster together in PC space, away from the other compounds, are evidence that at least some of these properties have relevance to what distinguishes the oral drugs from the other compounds. The magnitudes of the coefficients of the different properties in the first few PCs can then help identify which properties are contributing most to this distinction.

Whether such a finding is useful versus trivial depends on both the compounds chosen to comprise the set and the properties included in the analysis. Consider, for example, PCA for a compound set comprising some oral drugs plus, say, a variety of naturally occurring sugars. It is entirely predictable that some of the top-ranking PCs will be dominated by factors such as polarity, aqueous solubility, number of hydroxyl groups, etc., as the presence of many hydroxyls and a high molecular polarity and solubility are strong characteristics of many sugars. This result from PCA would be true but trivial, not only because it is predictable based on prior knowledge, but also because it reflects a characteristic difference between the compound sets being compared that is intrinsic to which class the compound belongs to. In a good analysis, the compounds in the analysis set will be matched as closely as possible and yet still manifest a diversity of behavior that requires explanation. Using PCA to compare oral drugs with, for example, synthetic compounds from a drug company screening library would be much more likely to provide interesting and useful results, provided that that relevant molecular properties are included in the input data.

As should be apparent from the above discussion, each application of PCA in fact tests a specific hypothesis: that the similarities and differences among the data elements in the chosen set can be meaningfully explained in terms of the set of properties the investigator has chosen to describe them. A useful feature of PCA is that inclusion of additional properties that are irrelevant to distinguishing the elements is not a problem; any such descriptors will have close to zero coefficients in the top-ranking PCs, and thus will effectively be ignored. However, inclusion of properties that do differentiate between the data elements, but in ways that are irrelevant to the question the user is interested in, will pollute the PCA results with true but useless information. For example, using PCA to analyze the similarities and differences among a group of people can be done using a virtually infinite variety of personal characteristics, but if the purpose of the analysis is to understand how they compare in terms of e.g. athletic performance, then including data on how highly each person rates Mozart versus Beethoven is unlikely to prove informative, even though it may contribute strongly to distinguishing the individuals in the group and therefore show high coefficients in a low-numbered PC. This possibility highlights the fact that PCA reveals patterns of covariance, but any inferences concerning cause and effect are interpretations being placed on these patterns by the investigator based on additional external knowledge or intuition. It may well be that, among a small set of individuals, the better athletes happen to prefer Beethoven to Mozart, but absent a compelling rationale connecting this musical preference to athletic performance it would be unwarranted to conclude that one causes the other Any conclusions about causation that derive solely from PCA should be considered as hypotheses only, as the method does not evaluate causation. Having said this, unexpected PCA results of this sort can form the basis of a hypothesis to be prospectively tested in a properly constructed and controlled experiment. As a hypothesis generation tool PCA can be very powerful, due to its ability to compress a complex input data set into a smaller number of significant features.

As discussed in the Introduction to this protocol, inclusion of too many properties – the so-called Hughes Phenomenon – can lead to a decrease in the performance of PCA. The problem arises when the number of elements in the data set is insufficient to adequately sample the property space due to the high dimensionality of the latter. One way to mitigate against this problem is to use the covariance analysis from Step 7 to identify groups of properties that are highly covariant with each other with respect to the elements in the data set. A single descriptor from each group can then be chosen to capture the information for inclusion in the PCA data set, and the others descriptors from the group discarded. Thinning out the descriptor set in this way can result in a simpler model in which all of the included properties are well-sampled. Minimal information will be lost from the analysis provided that the covariance between the grouped descriptors is very high. However, the extent to which further dimensionality reduction can be achieved by the PCA itself will be decreased by the prior removal of redundant information.

The above discussion suggests several pitfalls to be avoided when using PCA.

i. It is important to avoid including properties in the input data set that cannot reasonably have any causal connection to the question of interest. Doing so will avoid the results being polluted with factors that co-vary with the behavior of interest but cannot be causative. It is perfectly fine to include properties that might or might not be causative with respect to the question of interest – indeed, doing so and seeing which ones matter is often the point of an analysis. Identifying which of these properties contribute to differentiating the elements from each other can reveal important new patterns and hypotheses for further investigation. But when deciding which properties to include, the investigator should ask themselves: ‘If this property were to be revealed as important, would that tell me anything relevant to the purpose of my analysis?’ If the answer is ‘no’ then the property should be excluded.
ii. How the property values are scaled is also very important. As described in a previous section, properties with large numerical value ranges will tend to have a greater influence on distances in PC space than will properties expressed in smaller numbers. Consider, for example, a set of three compounds, A, B, and C, with respective molecular weights of 300, 350 and 351 mass units, and clogP values of 1.0, 1.1, and 5.0. Compounds B and C will lie close together in PC space while A will appear as an outlier, because the difference of ∼50 mass units that separates A from B/C is numerically large compared to the differences in clogP values. Medicinal chemists will immediately see this result as counterintuitive, and even misleading, because the four-unit higher value for clogP of compound C versus A and B will likely have much more influence on molecular behavior than their 50-unit difference in MW. Another aspect of the property scaling problem is that the choice of units is often arbitrary; distances on the molecular scale can be expressed in pm, nm, Angstroms, or meters, energy in either Joules or calories (or kJ or kcal), dipole moments in Coulomb meters or in Debyes, etc. The choice of units will affect the absolute numerical range of the property values, and thus how compounds appear to cluster in PC space. One common solution to this problem is to z-score the data. Doing so rescales each property value to have a Standard Deviation of exactly one. Z-scoring is not always appropriate, however. To z-score is to implicitly assume that the original variances of the input properties are equally significant for the question at hand, which may or may not be scientifically valid. For example, a set of compounds to be analyzed may derive from a common chemotype, such that the molecular weights vary over only a small range. If the property values are z-scored, the resulting PCA will treat this small variation in MW, which is unlikely to have any real impact on the behavior of the molecules, as if it were just as significant as much more consequential differences in other properties. A solution to this problem is to attempt to normalize the properties to a common scale in which a value at the bottom of the scale (say, zero) and the top of the scale (say, 1, or 100) represent differences in each property that are of roughly comparable scientific importance for the question at hand. For applications to drug discovery, for example, for any given property the low and high ends of the scale can simply be set at e.g. the 10^th^ and 90^th^ percentile with respect to the values observed across all drugs (not the compounds in the current data set). Values outside this range (i.e. <0 or >100) can and will occur, but expressing them on this scale marks them as representing unusually low or high values with respect to drugs in general. For PCA addressing questions in other fields of molecular science, devising similarly rational property scales may require careful thought on the part of the investigator. Thankfully, it is not important to achieve perfect accuracy, but only to avoid egregious distortion of PC space by having property variances for which a unit difference would have grossly different significance for the question under study.
iii. The choice of the set of items or elements to be analyzed using PCA is also important to the outcome. When analyzing chemical compounds, for example, using compounds that derive from a single chemotypes will reveal information about which among the provided molecular properties are most important in distinguishing one compound of that chemotype from another. In contrast, if compounds from multiple chemotypes are used, the analysis will instead reveal what properties distinguish the chemotypes from each other and will reveal little about distinctions within a chemotype. The more distinct the chemotypes are in their properties, the more this will be the case. Quite different properties may be identified as important in these two analyses. It is important to recognize that these two analyses are in fact asking different questions, and to make sure the set of elements you choose for a given analysis is asking the question you wish to answer.

A related issue is the importance of using a data set of balanced composition when using PCA for certain purposes. If the goal is to understand what properties distinguish between elements of different types, such as different compound chemotypes, it is important that the data set contains a roughly similar number of representatives of each type. PCA itself is agnostic to any preconceptions the investigator may have concerning how the elements in the data set might be described as breaking down into subgroups. Instead, the analysis considers the differences between each pair of elements to no more or less important than that of any other pair. Consequently, if the data set comprises mostly elements of one type and contains only a few representatives of another, the analysis will be strongly biased towards factors that distinguish between the compounds in the larger set as these make up the lion’s share of the comparisons that can be made, which is not the question the investigator is trying to answer. One approach to equalizing the sizes of the subclasses of elements to be included in the overall data set is to first use the technique of *k*-medoids clustering (Basic Protocol 1c) on each subgroup of elements to select a subset of *k* representatives from each, where *k* is set as the number of elements in the smallest subset. The selected representatives from each subset can then be combined to form an overall data set that contains *k* representative members from each subset.

## VII. SUPERVISED ML METHODS: Partial Least Squares-Discriminant Analysis (PLSDA) and Partial Least Squares Regression (PLSR)

Partial Least Squares Discriminant Analysis (PLSDA) and Partial Least Squares Regression (PLSR) are supervised ML tools used to find relationships between two types of data matrices, denoted as the input and output variables (X and *y*). Like PCA, PLSDA and PLSR reduce the dimensionality of X by combining covariant information into a smaller number of composite axes called ‘components’ (or sometimes ‘latent variables’). However, unlike PCA, these supervised ML methods identify and rank the components based on their covariance with the output variable, *y*. Thus, PC1 is the composite axis that captures the highest possible fraction of the variance in *y*, PC2 is an orthogonal axis that contains as much as possible of the remaining variance in *y*, and so on. As a result, the methods allow the value of *y* to be modeled based on the contributions of the various descriptor values in X, allowing a value for the behavior *y* to be predicted for a new item if its input property values are known.

The main difference between PLSDA and PLSR is whether the outcome or behavior described in *y* takes the form of a categorical variable (i.e. falls into discrete classes, e.g. ‘yes’ versus ‘no’ or A versus B) or whether the behavior can be expressed on a continuous numeric scale (e.g. a response level, a time, etc.). PLSDA is designed to model categorical outcomes, while PLSR is a related method but for outcomes with continuous numerical values. Note that some outcomes that can be classified on a numeric scale (e.g. solubility) can also be expressed categorically (i.e. soluble versus insoluble, with respect to some chosen threshold). The rate at which a compound can permeate through a lipid membrane, which we use to illustrate these methods here, is one such behavior.

For both methods we use experimental permeability data for a set of cyclic peptide compounds, downloaded from the CycPeptMPDB database (http://cycpeptmpdb.com) (Li *et al*., 2023) as described in Section I. Compound permeabilities were measured using the Parallel Artificial Membrane Permeability Assay (PAMPA) method, as described by Kansy *et al. (1998)*. For the demonstration of PLSDA in Basic Protocol 3, we categorized the 50 compounds with the highest permeability, all of which fall above a threshold of P = 2.9 × 10^−5^ cm/s, as ‘Permeable’ whole the 50 least permeable compounds, all with permeability above P =1.3 × 10^−6^ cm/s, were simplistically classed as ‘Impermeable’. Compounds with intermediate permeability (i.e. between these thresholds) were excluded, to ensure that the two classes of compounds were distinctly different in behavior as is desirable for a categorical analysis.

PLSDA requires an input data matrix X and an output data matrix, *y*. In the current case, X comprises an *m* x *n* matrix containing the values of 139 different molecular properties for each of the compounds to be analyzed. This input data is similar to the data used for PCA in Basic Protocol 2, except that the set of compounds included is smaller due to the exclusion of borderline permeable compounds. For the output data matrix, *y*, we assigned a numerical label to each class of behavior: compounds rated as Impermeable were assigned the number 1 in the output vector y, while Permeable compounds were assigned the number 2. Thus, *y* comprises a *m* x 1 matrix, where *m* is the number of compounds, with each number in the matrix being either a 1 or a 2 depending on whether the compound in the corresponding row of X was classed as Impermeable or Permeable. For the PLSR analysis in Basic Protocol 4, we aim to predict the actual numeric permeability values, so no assignment to classes was needed. For Basic Protocol 4, therefore, X is an *m* x *n* matrix of 139 property values (columns) for the full set of 201 compounds (rows), while *y* is simply an *m* x 1 vector containing log_10_ of the experimentally determined permeability value for each compound in X, in units of cm/s.

As with other quantitative modeling methods, for PLSDA and PLSR it is good practice to reserve a subset (perhaps 20%) of the data to serve as a Test Set that can be used to validate the quality of the model that is developed. The remaining 80% of the compounds, which are used for model building, are called the Training Set. The compounds for the Test Set can be chosen randomly. However, when working with relatively small data sets random selection can lead to bias if, by chance, a number of very similar or highly atypical items happen to be selected. Alternatively, a representative subset can be selected for the Test Set, using a method such as *k*-medoids clustering (see Basic Protocol 1c).

Having designated the Training Set and Test Set, the PLSDA or PLSR analysis is performed on the Training Set, resulting in a set of Principal Components (PCs) – composite axes that contain the information in X and *y* after elimination of covariance in X and maximization of covariance with *y* – plus a set of scores that give the coordinates of each compound with respect to the PC axes. Each PC is defined by a set of coefficients, typically called ‘loadings’ in PLS methods, that quantify the extent to which each of the n original properties in X contributes to that PC; the large the absolute value of the loading, the greater the influence of that property on the PC in question. (For a more detailed description of PCs, scores, and coefficients/loadings, the reader is referred to the Introduction to PCA in Basic Protocol 2.)

An important step in PLSDA and PLSR is the decision of how many PCs to use in building the model of behavior *y*. The PCs are ranked in order of how much variance in *y* they account for (or ‘explain’, in the jargon of the field). Moreover, all PCs will contain a certain amount of ‘noise’, to the extent that the descriptor values in X are not perfectly accurate or, in some cases, are irrelevant to the behavior described in *y*. Consequently, the first few PCs will contain the most relevant information for explaining *y*, with the later PCs contain increasingly less useful information and proportionately more noise. Including too may PCs in the model risks beginning to fit this noise, a phenomenon called ‘overfitting’ that reduces the predictive accuracy of the model. One hallmark of an overfitted model is that, while its ability to account for the variations in *y* in the Training Set can be very good, when applied to the Test Set the model performs much less well. An efficient way to select the optimal number of PCs to include in the model is by using a process called ‘cross-validation’. In this process, a model is first built using only PC1, but leaving out a subset of the data points from the Training Set, and then the accuracy with which the model predicts the values of *y* for the omitted data points is calculated. This process is repeated, leaving out a different subset of the data points in turn, until all data points in the Training Set have been omitted once and their output property predicted. The root-mean-square (rms) prediction error across all predictions is calculated and taken as a measure of the predictive power of the model when built using only PC1. The same process is then repeated but using both PC1 and PC2 for the model building, then PCs 1-3, and so on. Plotting the rms prediction error as a function of the number of PCs included in the model reveals what number of PCs to include to minimize the prediction error and thus maximize predictive power. All of this can be done automatically using a single MATLAB command. Compared to this optimal number of PCs, using fewer PCs gives a model with poorer predictive power because relevant information is being excluded, and using more PCs gives poorer predictive power because the noise that is thereby introduced is more detrimental to the model than the benefit from any added relevant information. The model generated using this optimal number of PCs can then be applied to the separate Test Set of data for a more rigorous measure of predictive accuracy.

Once a PLSDA or PLSR model has been developed using the optimal number of PCs, it can be applied to predict the value of the output property *y* for any item for which the values of the input descriptors in X are provided.

## BASIC PROTOCOL 3: Partial Least Squares Discriminant Analysis (PLSDA)

### Purpose of method

To build a model that can take a set of elements, and can categorize the elements with respect to some observable or measurable outcome based on information about their properties. For PLSDA, the outcome is a categorical variable (behavior = class A versus class B, etc.), rather than being describable on a continuous numerical scale. For example, PLSDA could be used to analyze whether cells proliferate or die (the categories of behavior) based on, for example, the expression levels of a set of cell-surface receptors or the amplitude or timing of a set of intracellular signaling events (the properties).

### Background

PLSDA is conceptually related to the unsupervised ML method of Principal Component Analysis (PCA). It takes as input data the numerical values of a set of *n* properties for each member of a set of *m* elements and transforms these data to eliminate redundant information and thereby generate a smaller number of composite properties (Principal Components) that allow the information to be analyzed at a lower dimensionality. In our case, the input data (‘X’) comprise values of 139 different molecular properties for each of 80 different cyclic peptides in our compound set, and the output data (‘*y*’) are experimental values for the compounds’ passive permeability through a lipid membrane. As discussed in more detail in Basic Protocol 2, many of the molecular properties provided in X will likely be to some extent covariant, and thus contain a degree of redundancy in the information they contain about the molecules. By condensing this information into a smaller set of mutually orthogonal principal components, each of which corresponds to a distinct and nonredundant portion of the input data, the same information can be visualized and analyzed using fewer axes. The result of PLSDA is a model that can be used to predict the output behavior (i.e. permeability) of new elements (i.e. compounds) based on their input properties.

As is generally the case with supervised ML, the Protocol below generates a model using a Training Set comprising 80% of the available data. This model is then validated by testing how accurately it predicts the permeability of the 20% of the data that was held back to serve as a Test Set.

PLSDA is designed to predict categorical behaviors – that is, behaviors that can be assigned to a category. In the case described here, the individual compounds are categorized as ‘Permeable’ or ‘Not Permeable’ based on whether their experimentally measured permeation falls above or below designated threshold values. For modeling the value of a continuous numerical variable see the ML method of PLSR, described below in Basic Protocol 4.

Key considerations for PLSDA, as for PLSR, are (i) how to divide the available data into the Training and Test Sets, (ii) how many components to use in the final model building, (iii) how to assess the predictive accuracy of the model, (iv) how to judge the model’s ‘domain of applicability’ (i.e. for what kinds of compounds can the model be expected to accurately predict the output behavior?), and (v) and how to determine which properties are most important in driving differences in the output behavior.

#### Before you start

Be sure you have installed the necessary toolboxes mentioned in the ‘REQUIRED SOFTWARE’ section.

The stepwise instructions below assume that, for the Training Set and Test Set, you will be using the data we provide with this article (Supplementary file Data_PLSDA.xlsx). The Test and Training Sets used in this protocol were derived from the set of 201 compounds used in Basic Protocols 1 and 2, but with inclusion of the membrane permeability (PAMPA) data for the compounds. For Basic Protocol 3, we took the 50 compounds with the lowest permeability and the 50 most permeable. For the purpose of this protocol we will refer to these classes as ‘Impermeable’ and ‘Permeable’. We then used *k*-medoids clustering (Basic Protocol 1a) to select 10 representative compounds from the high permeability class and another 10 that are representative of the low permeability compounds. These 20 compounds were set aside to serve as the Test Set, while the remaining 80 compounds comprise the Training Set. A detailed description of how we generated the Training Set and Test Set in Data_PLSDA.xlsx is provided in the supplementary document (Data_prepared_SI.xlsx).

If you are using your own data rather than the supplementary file provided, be sure to format your external data files as described in ‘Section IV. DATA DOWNLOAD AND PREPARATION, and illustrated below in Figure 3.1, and to change the external filenames (blue text) in the following steps to match the names you have assigned to your files.

##### 1. Gather the input data

Details of the data download and preparation are given above in Section IV, and spreadsheets containing the data required to perform the protocols are included in the supplementary file ‘Data_PLSDA.xlsx’. For this protocol, the spreadsheets contain the values of 139 molecular features (descriptors) for a Training Set of 80 compounds, plus the corresponding data for a Test Set of 20 compounds that will be held back from model building and instead used to evaluate the predictive accuracy of the model once built. The supplementary file ‘Data_PLSDA.xlsx’ contains the data already formatted in this way, in the sheets named ‘PLSDA_Train’ and ‘PLSDA-Test’.

**Figure 3.1:**
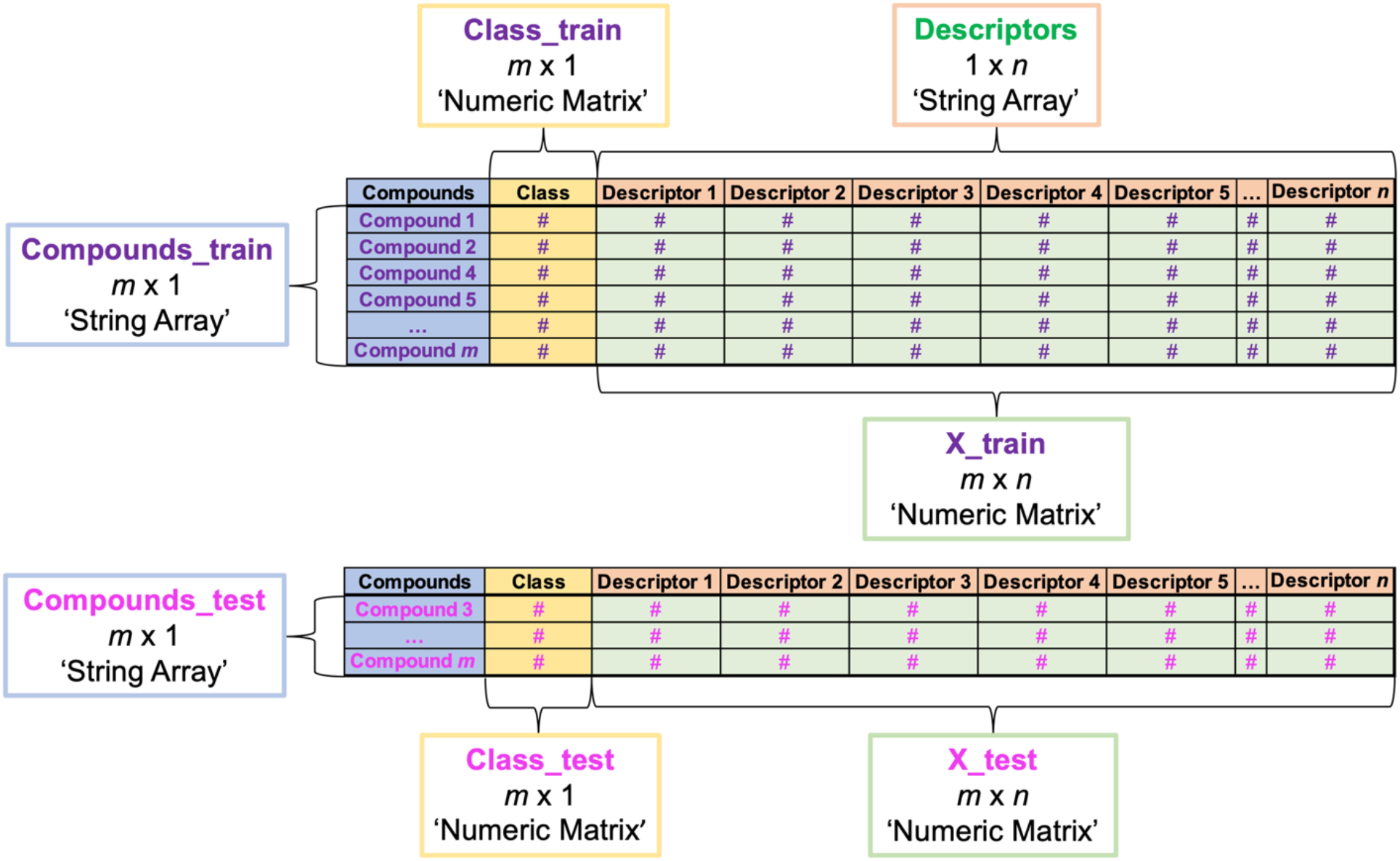
How to format data for import into MATLAB for PLSDA. Text in green indicates the names we assigned to the different data variables created during import. Black text in single quotes indicates what import file type to choose when importing each piece of data using the MATLAB Import Data window. If you are following the example in Protocol 3, you will simply import data we have already formatted, as shown above, from two tabs (train and test) in the supplementary file Data_PLSDA.xlsx.

> *If performing PLSDA on other types of data, the compound names in column 1 would be replaced by identifying labels for the different elements in the set, and the descriptor names in row 1 would be replaced by the names of the different properties that describe the elements*.

##### 2. Import the numerical input data, X_for the Training Set of compounds

i. Starting with a clear workspace, click on the Import Data button in the MATLAB Home tab (top of screen), navigate to where you have saved the Supplementary File ‘Data_PLSDA.xlsx’.
ii. Working in the Import Data window, open the tab labeled ‘PLSDA_train’. <u>Be sure you are on the correct sheet before you begin importing data</u>.
iii. Select the cells that contain the numerical data. <u>Be sure to exclude the column headers in Row 1 as well as the compound names, PAMPA logP values, and permeability class designations in columns 1-3</u>. The selected data should comprise 80 rows (reflecting the 80 compounds in the Training Set) and 139 columns (reflecting the 139 molecular descriptors we are using for this analysis). The selected data should include only numbers.
iv. Go to ‘Output Type’ at the top of the Data Import window and select ‘Numeric Matrix’ from the drop-down menu, then go to the Import Selection button at the top right of the Import Data window and click the green check mark (or select ‘Import Data’ from the drop-down menu). You should see a new data object with dimensions 80 × 139 in the Workspace window of your MATLAB desktop (you may need to move the Import Data window aside to see it).
v. Rename the 80 × 139 workspace variable. For the purpose of this exercise, you should call the new workspace variable **X_train**. You can do this in the open Data Import window by double clicking on the default name (same as the original Excel filename) that appears just below the top row of the window and typing in the desired new name. Alternatively, you can click on the newly created variable in the MATLAB Workspace window to highlight its default name, and type in the new name, **X_train**.

> *Data variables created in MATLAB can be renamed after the fact, as described in Step 2(v), but it is actually best practice to assign the desired name in the Data Import window before clicking ‘Import Data’ to create the variable. The reason is that the default name that MATLAB assigns to new data variables is the name of the external file from which the data were imported, making it easy to accidentally over-write a different data variable previously imported from the same external file if it was not assigned a unique name upon creation*.

##### 3. Import the output data, *y*, for the Training Set

Repeat the selection process from Step 2, but this time selecting the data from Column C, labeled ‘Class’, in the ‘PLSDA_train’ tab of Data_PLSDA.xlsx. Change the ‘Output Type” to ‘Numerical’, rename the data variable as **Class_train**, and then select ‘Import Data’. A new 80 × 1 data variable, **Class_train**, will appear in your Workspace window.

> *Import only the numerical values, being sure to exclude the header name (‘Class’) in Cell 1C*.

##### 4. Import the compound names for the Training Set

From the ‘PLSDA_train’ tab of Data_PLSDA.xlsx, import the compound names from Column A. Change the ‘Output Type’ of the matrix to ‘String Array’, and then select ‘Import Data’ to generate the array. Rename the new 80 × 1 string array that appears in the Workspace window as **Compounds_train**.

> *Exclude the column header ‘CycPeptMPDB_ID’ in cell A1, as this is not a compound name*.

##### 5. Import the descriptor names for the Training Set

From the ‘PLSDA_train’ tab of Data_PLSDA.xlsx, select the descriptor names (i.e. the column headers that appear in Row 1). Change the ‘Output Type’ of the matrix to ‘String’, assign the name **Descriptors** to the new data variable, and then select ‘Import Data’ to generate a new 1 × 139 string array in the Workspace window.

> *Be sure to exclude the column headers ‘CycPeptMPDB_ID’, ‘PAMPA’ and ‘Class’ in cells A1-C1, as these are not descriptor names. If you are using the Data_PLSDA.xlsx data file provided with this article, this means you should start your selection with cell D1 (‘MaxEStateIndex’)*.

##### 6. Import the numerical input data, compound names and output (*y*) data for the Test Set

Use the Data Import window to navigate to the sheet called ‘PLSDA_test’ in the file Data_PLSDA.xlsx, in the same manner as in steps 2-4, naming the data variables **X_test** (20 × 139), **Class_test** (20 × 1), and **Compounds_test** (20 × 1), respectively.

> *Be sure you are on the correct sheet, ‘PLSDA_test’, before importing these data. There is no need to import the descriptor names again, as these are identical for the Test Set and the Training Set and were already imported in Step 5*.
>
> *After performing Steps 1-6, your MATLAB workspace should contain the following 7 data variables:* ***Class_test, Class_train, Compounds_test, Compounds_train, Descriptors, X_test***, and ***X_train***.

##### 7. Save your MATLAB workspace as a .mat file (See Section V)

> *This file can be reopened to give a MATLAB workspace identical to the state that was saved*.

##### 8. Z-score (scale and normalize) the X_train values

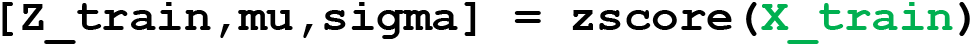

> *Z-scoring centers the values for each descriptor (i.e. each column in X) on zero and scales them to SD = 1. (See Protocol 1a, Step 7 for more details about z-scoring, including how to generate boxplots to visualize the results.)*

##### 9. Replace any ‘NaN’ (‘Not a Number’) results from Step 9 with zeroes

~~~
Z_train(isnan(Z_train)) = 0
~~~

> *NaN entries result from invalid mathematical operations such as division by zero, and their presence can prevent the proper operation of some subsequent commands. If you are using the data sets provided with this article, no NaN values will be generated. However, we include this step in case you are following this protocol with your own data set, for which NaN results could occur*.

##### 10. Select the optimal number of components (latent variables) for PLSDA

This is done by means of cross-validation, as described in the Introduction to Section VII, for which we use the ‘plsdacompsel’ command reported in *Ballabio and Consonni, 2013*.

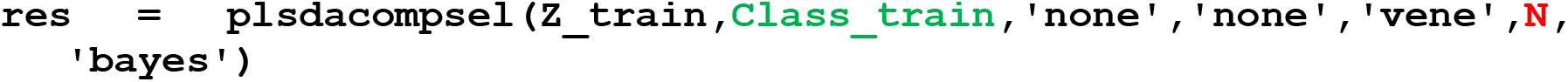

*The key inputs for this command are as follows:*

*‘Z_train’: Our z-scored X data for the Training Set, created in Step 8*.

***Class_train***: *Our output*, y, *data for the Training Set, imported in Step 3*.

*‘****N****’: The number of groups the data are to be divided into for cross-validation*.

*‘****N****’ has a value from 2 to* m, *where* m *is the number of elements in the set (in our case, 80). Setting ‘****N****’ = 2 will divide the data into two groups and run the modeling twice, leaving out each group in turn and then assessing the accuracy of the model generated using each half of the data set at predicted the permeability class of the other half of the compounds that were omitted. At the other extreme, setting ‘****N****’ =* m *will run the model* m *times, leaving out a single compound each time and measuring how well its permeability class is predicted by the model made using the data for the remaining* m*-1 elements. This latter procedure is known as ‘leave-one-out’ cross-validation. The fewer elements that are omitted in each round of cross-validation, the better the models in each round are likely to be. Thus, setting ‘****N****’ =* m *(i.e. ‘****N****’ = 80, in this case) is a safe choice. However, setting ‘****N****’ <* m *is sometimes advisable to reduce the computation time when working with large data sets*.

*The other inputs for the ‘plsdacompsel’ command are as follows:*

*preprocess_rows: Instructions for centering or scaling the data that comprise the rows in ‘Z_train’. None is wanted here, as our columns of descriptor values represent independent inputs, so the values in any given row have no statistical relationship to each other*.

*preprocess_columns: Instructions for centering or scaling the property values that comprise the columns in ‘Z_train’. None is required here because we have already scaled and centered these data by z-scoring in Step 8*.

*cv_type: Instructions for how the elements to be omitted from the model in each step of the cross-validation should be selected. We have selected the ‘Venetian blinds’ method (‘vene’). For other options, see [plsdacompsel documentation](https://viewer.mathworks.com/?viewer=plain_code&url=)* plsdacompsel documentation plsdacompsel documentation.

*assign_method: Instructions for how the element(s) omitted in each round of cross-validation is assigned to an output category (in this case, ‘Permeable’ or ‘Not permeable’) by the model. Here, we chose a method based on Bayesian Theory (‘bayes’). For the alternative option, see [plsdacompsel documentation](https://viewer.mathworks.com/?viewer=plain_code&url=)plsdacompsel documentation*.

> *As output the ‘plsdacompsel’ command generates a MATLAB structure object, called here ‘res’, that contains the following output data variables:*
>
> *‘er’: An n x 1 vector, where n is the total number of PCs = the total number of descriptors, which here = 139) containing the fraction of <u>incorrect predictions</u> observed in each round of cross-validation, starting from the round that used only PC1, and then the round that used PCs 1 and 2, and so on*.
>
> *‘ner’: An n x 1 vector containing the fraction of <u>incorrect predictions</u> observed in each round of cross-validation, starting from the round that used only PC1, and then the round that used PCs 1 and 2, and so on*.
>
> *‘not_ass’: An n x 1 vector containing the fraction of <u>elements that could not be assigned to either output category</u> observed in each round of cross-validation, starting from the round that used only PC1, and then the round that used PCs 1 and 2, and so on*.

##### 11. Create a plot to visualize the prediction accuracy revealed by cross-validation upon inclusion of each successive PC in the model building

~~~
figure()
plot(res.er,’Marker’,’o’,’LineWidth’,2,’DisplayName’,’Incorrect
  predictions’,’Color’,’red’)
hold on
plot(res.ner,’Marker’,’o’,’LineWidth’,2,’DisplayName’,’Correct
  predictions’,’Color’,’green’)
hold on
plot(res.not_ass,’Marker’,’o’,’LineWidth’,2,’DisplayName’,’Not
  assigned’,’Color’,’black’)
legend(‘show’)
legend(‘Location’,’best’)
set(gca,’FontName’,’Arial’,’FontSize’,14);
xlabel(‘Number of PCs’)
ylabel(‘Fraction of answers (predictions)’)
xlim([1 20])
~~~

A new Figure window will appear containing the plot shown in **Figure 3.2**.

**Figure 3.2.**
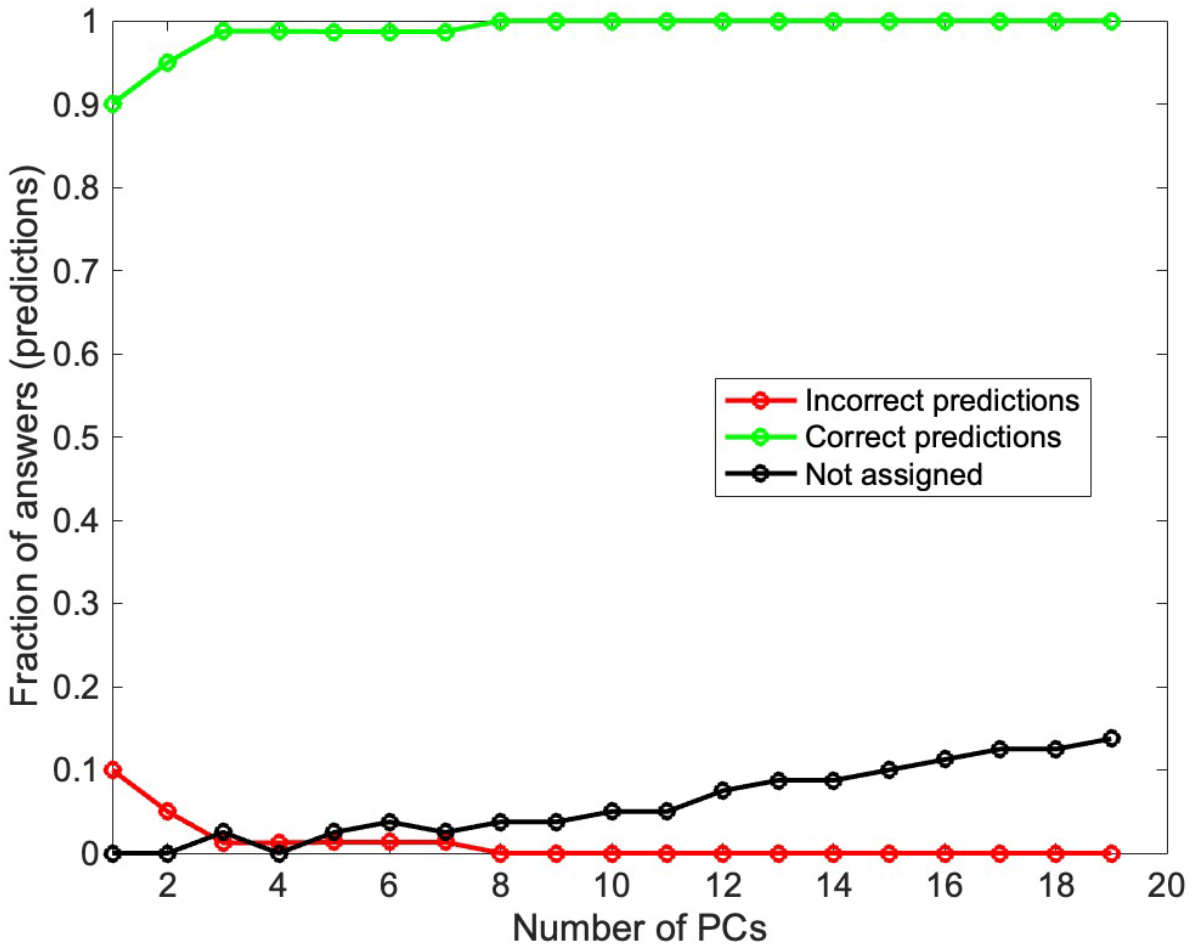
Results from leave-one-out cross-validation for PLSDA models generated using the data in the Training Set and including the number of PCs indicated on the x-axis. The model using four PCs gave the best performance, with a very low error rate and few compounds that could not be assigned to an output category. Inclusion of more PCs slightly reduced the already low error rate, but at the cost of increasing numbers of unassigned compounds.

##### 12. Generate a PLSDA model of the Training Set using the optimal number of PCs

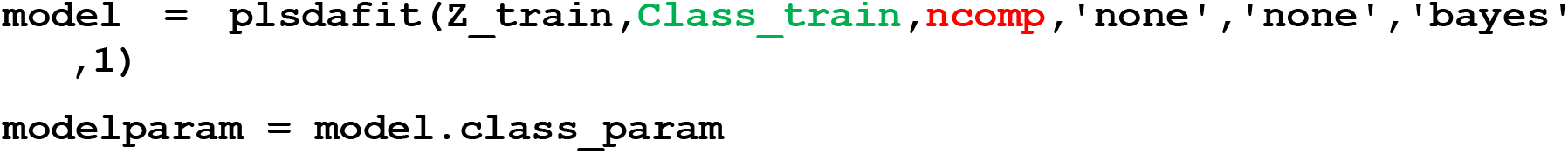

> *The user-selected input* ***ncomp*** *is the number of PCs to be used to build the model. Here, we set* ***ncomp*** *= 4, as this number was found to give the model with the greatest predictive accuracy as assessed by cross-validation in Step 11 (****Figure 3.2****). Other inputs for the command are ‘Z_train’*, ***Class_train***, *‘preprocess_rows’, ‘preprocess_columns’, ‘assign_method’, and ‘doqtlimit’. These inputs have the definitions and values used described in Step 10, except for ‘doqtlimit’ which is new. Setting ‘doqtlimit’ to ‘1’ has the command calculate Hotellings T-squared statistics (See Basic Protocols 2, Step 16) and Q limits, while setting it to ‘0’ omits these outputs. For additional explanation, see [plsdafit documentation](https://viewer.mathworks.com/?viewer=plain_code&url=)plsdafit documentation*.
>
> *As output the ‘plsdafit’ command generates a MATLAB structure object called here ‘model’ that contains the following key output files:*
>
> *‘yc’: An* m *x 2 matrix, where* m *is the number of compounds in the Training Set (i.e. 80), indicating the calculated outcome for each compound across two classes (Impermeable/Permeable). For example, if the compound belongs to Class 1, then the value in ‘yc’ would be higher in Class 1, with the difference in magnitude between the two numbers indicating the degree of confidence in the classification*.
>
> *‘T’: An* m *x* n *matrix giving the scores for each of the 80 compounds (rows) with respect to each of the 139 PCs (columns)*.
>
> *‘P’: An n x n matrix containing the coefficients for each of the 139 original descriptors (rows) in each of the 139 PCs (columns)*.
>
> *‘class_calc’: An m x 1 vector indicating which output class each compound was assigned to. A value of ‘1’ indicates the model assigned the compound as Impermeable’, ‘2’ indicates Permeable, and ‘0’ indicates ‘Not assigned’*.
>
> *‘class_param’: A MATLAB structure object containing a number of output variables relating to the accuracy of the model. These include ‘confmat’, ‘error_rate’, ‘specificity’ and ‘sensitivity’*.
>
> *‘confmat’: Represents the confusion matrix for the classification performed by the PLSDA model. It has a numerical matrix structure (in our case 2 × 3), where the rows correspond to the true (actual) class labels (1 and 2 in our example), and the columns correspond to the predicated class labels (1,2 and 0). The elements in the matrix represent the number of samples that fall into each actual-predicted class combination. The sum of elements in the matrix is equal to* m *(80 in our example). This output will be used in step 15 to visualize the confusion matrix*.
>
> *A number of other output variables are also generated by the ‘plsdafit’ command but are not discussed here as we do not use them in any subsequent steps. To learn about these other outputs, see [plsdafit documentation](https://viewer.mathworks.com/?viewer=plain_code&url=)*plsdafit documentation.

##### 13. Plot the scores in PCs 1 and 2 from the model as a 2D scatter plot

to visualize how well the model distinguishes the output classes.

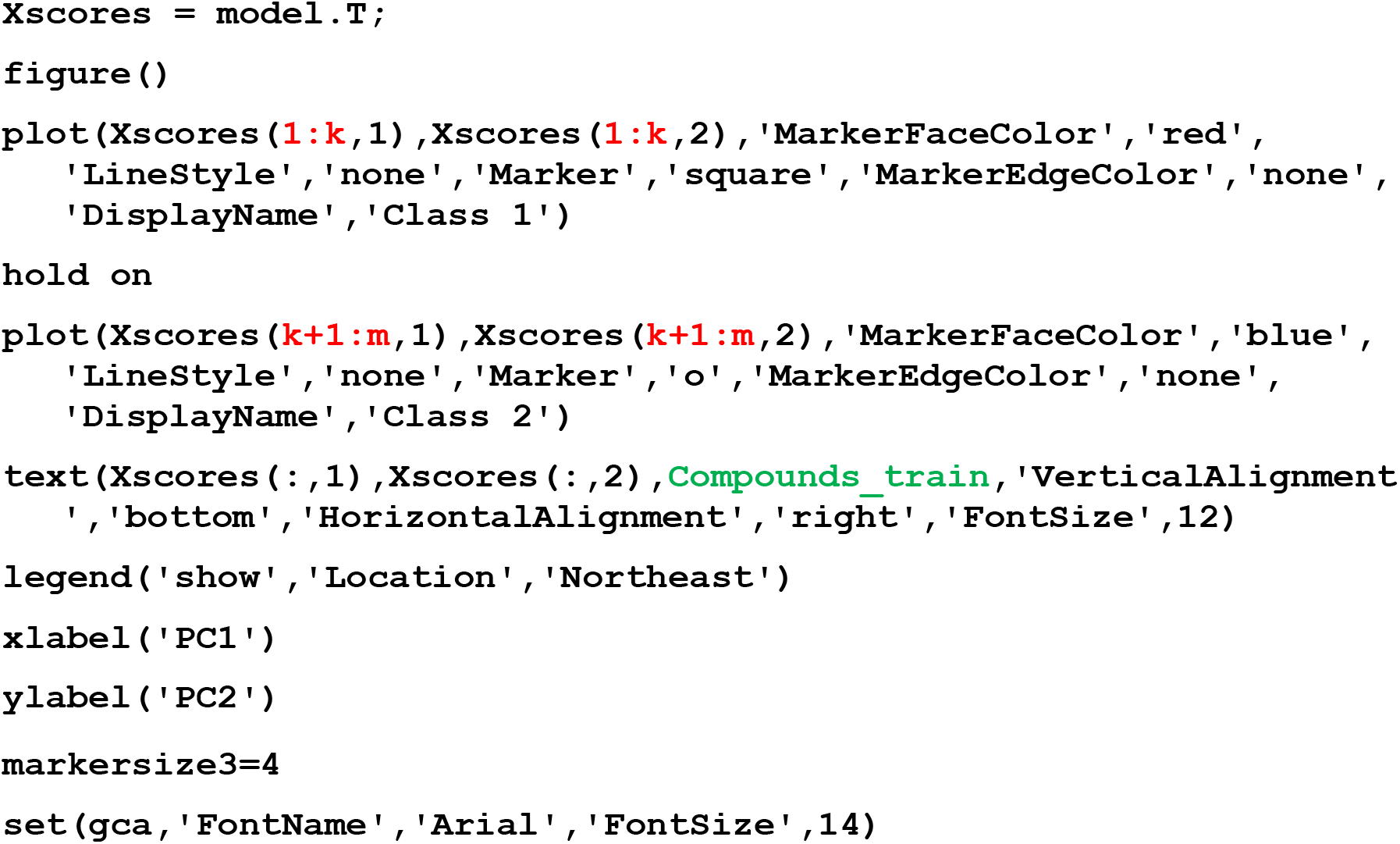

> *Lines 1-5 of the above command plot the data, while lines 6-11 serve to add the compound IDs, legend, axis labels, and to adjust the appearance of the plot*.
>
> *In the above command, the inputs ‘****k****’ and ‘****m****’ define the ranges of compounds (by row number in the original input files* ***X***, *Class_train, etc.) so that these data points can be plotted in a particular color. In our case, the Training Set was constructed such that rows 1-40 of the input data matrix* ***X*** *contain the Impermeable compounds (Class 1), and rows 41-80 contain the Permeable compounds (Class 2). (You can confirm these row numbers by looking at* ***Class_train****). In this command, therefore, setting* ***k*** *=40 and* ***m*** *= 80 allows us to color the data points corresponding to the permeable compounds in rows 1-40 red, and the impermeable compounds in rows 41-80 blue*.
>
> *For the purpose of visualization, the data points in* ***Figure 3.3*** *are labeled with the corresponding compound IDs, even though this makes for a crowded figure. To eliminate these labels, omit from the command the line that begins* `text(Xscores(:,1)`….
>
> Figure 3.3. The scores from the PLSDA model from Step 12, which used four PCs, plotted for PCs 1 and 2. The plot demonstrates good separation between the two classes of compounds along the 1st and 2nd PCs based on the information contained in the underlying descriptors. The Impermeable compounds are designated as Class 1 (red squares), while those that are Permeable are designated Class 2 (blue circles). 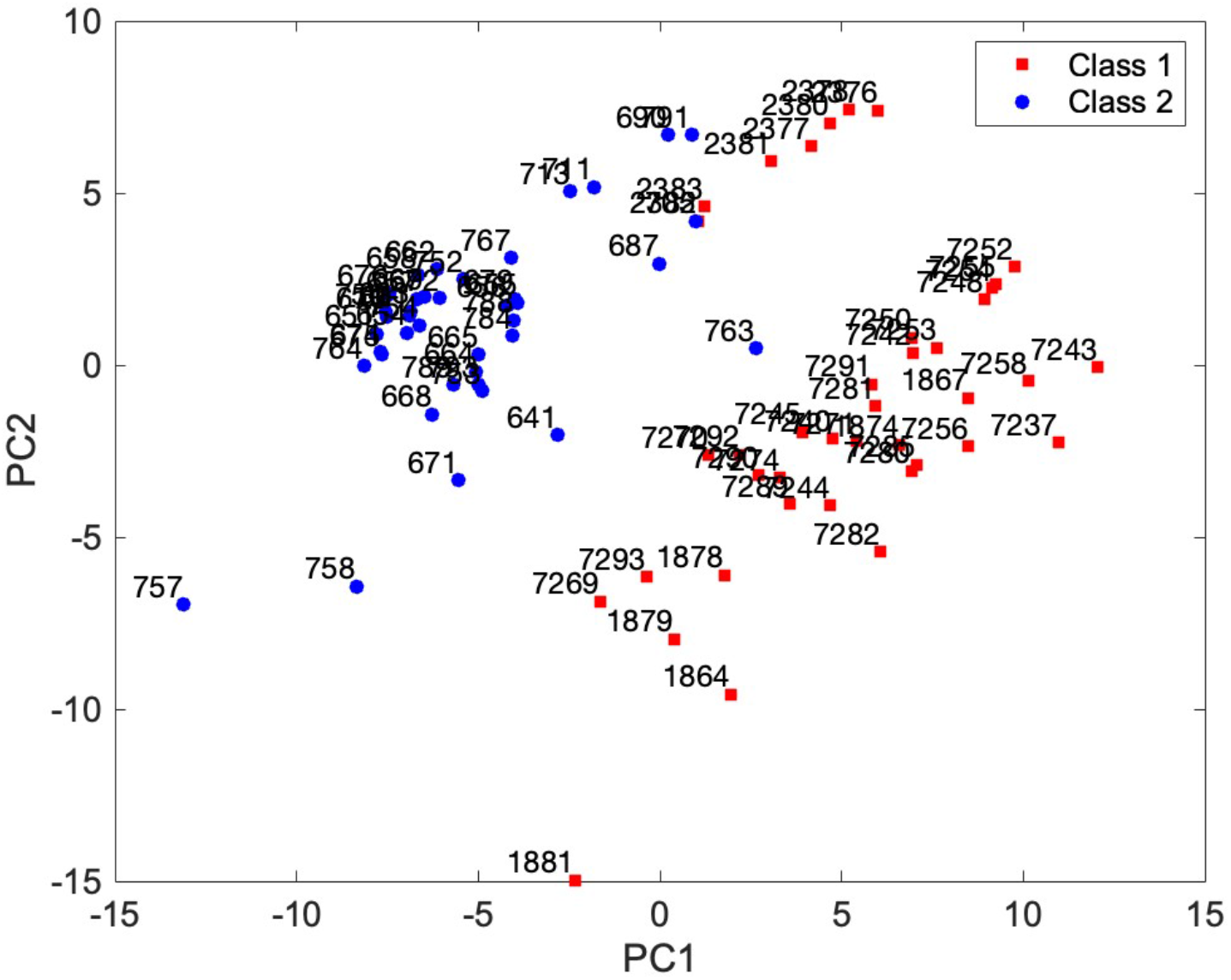
>
> *Examination of* ***Figure 3.3*** *shows that the red and blue data points, representing the Impermeable and Permeable compounds in the Training Set, are largely confined to different regions of PC space. This observation indicates that the model is achieving meaningful discrimination between the two outcome classes of compounds. In certain regions of the plot, however, the two outcome classes do appear to come very close together or even overlap. This appearance may reflect reality (i.e. that the model only imperfectly discriminates between the classes based on the descriptors provided), or it may instead be because in this figure we are looking at the data only with respect to their coordinates in PCs 1 and 2*. ***Figure 3.2*** *established that four PCs were required to achieve optimal discrimination, and that 2 PCs were not sufficient. Thus, some of the data points in* ***Figure 3.4*** *that appear to actually or nearly overlap as plotted may, in fact, be well separated in PC3 and/or PC4. To see if this is the case, we can plot the data in 3 PC dimensions to visualize PCs 1, 2, and 3. We do this in Step 14*.

##### 14. Plot the scores in PCs 1, 2 and 3 from the model as a 3D scatter plot

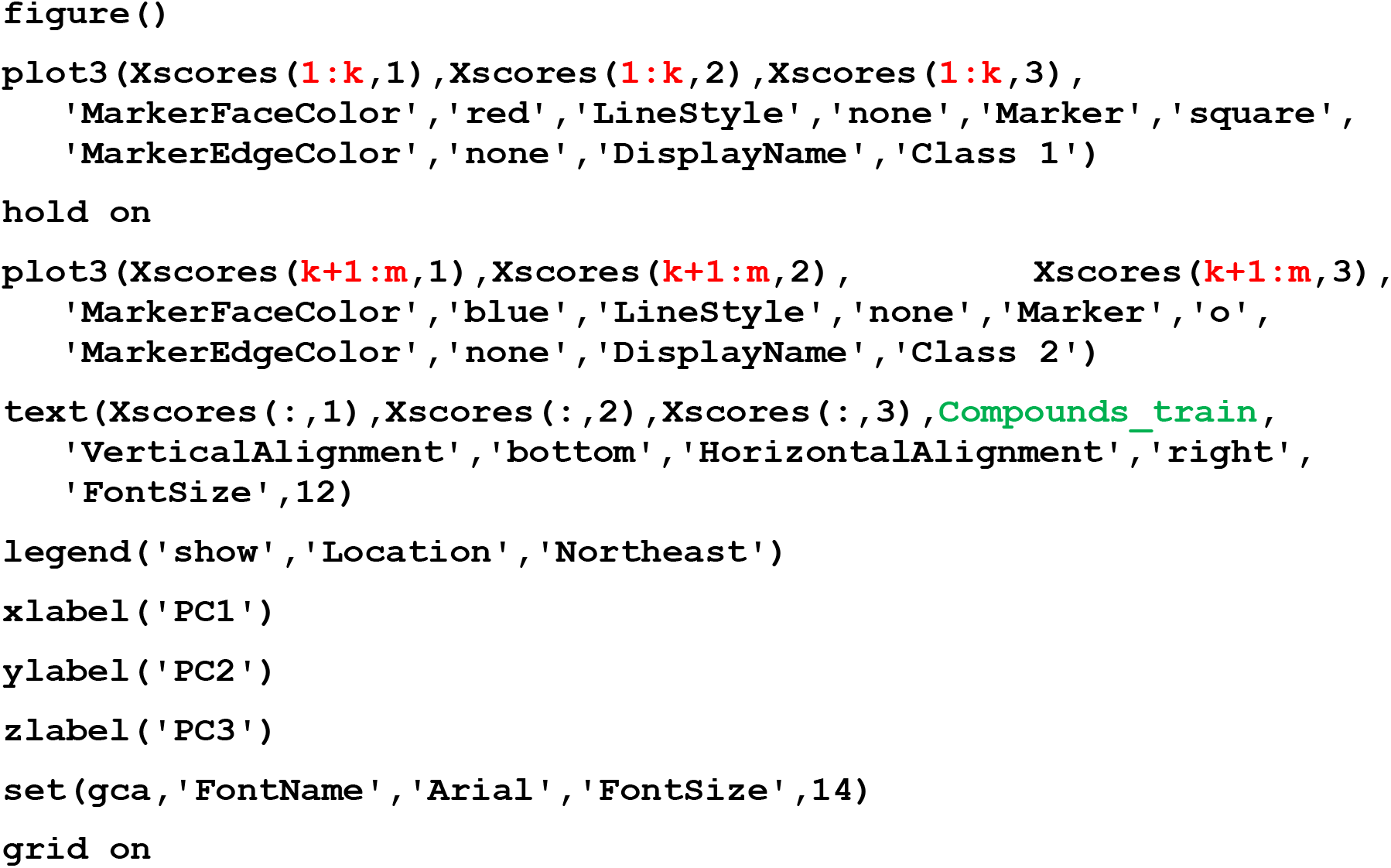

> *The values of* ***k*** *and* ***m*** *are set at 40 and 80, respectively, as explained in the preceding step*.
>
> Figure 3.4. The compound scores from the PLSDA model from Step 12, which used four PCs, plotted with respect to PCs 1, 2 and 3. The compounds that are Impermeable are designated as Class 1 (red squares), while those that are Permeable are designated Class 2 (blue circles). 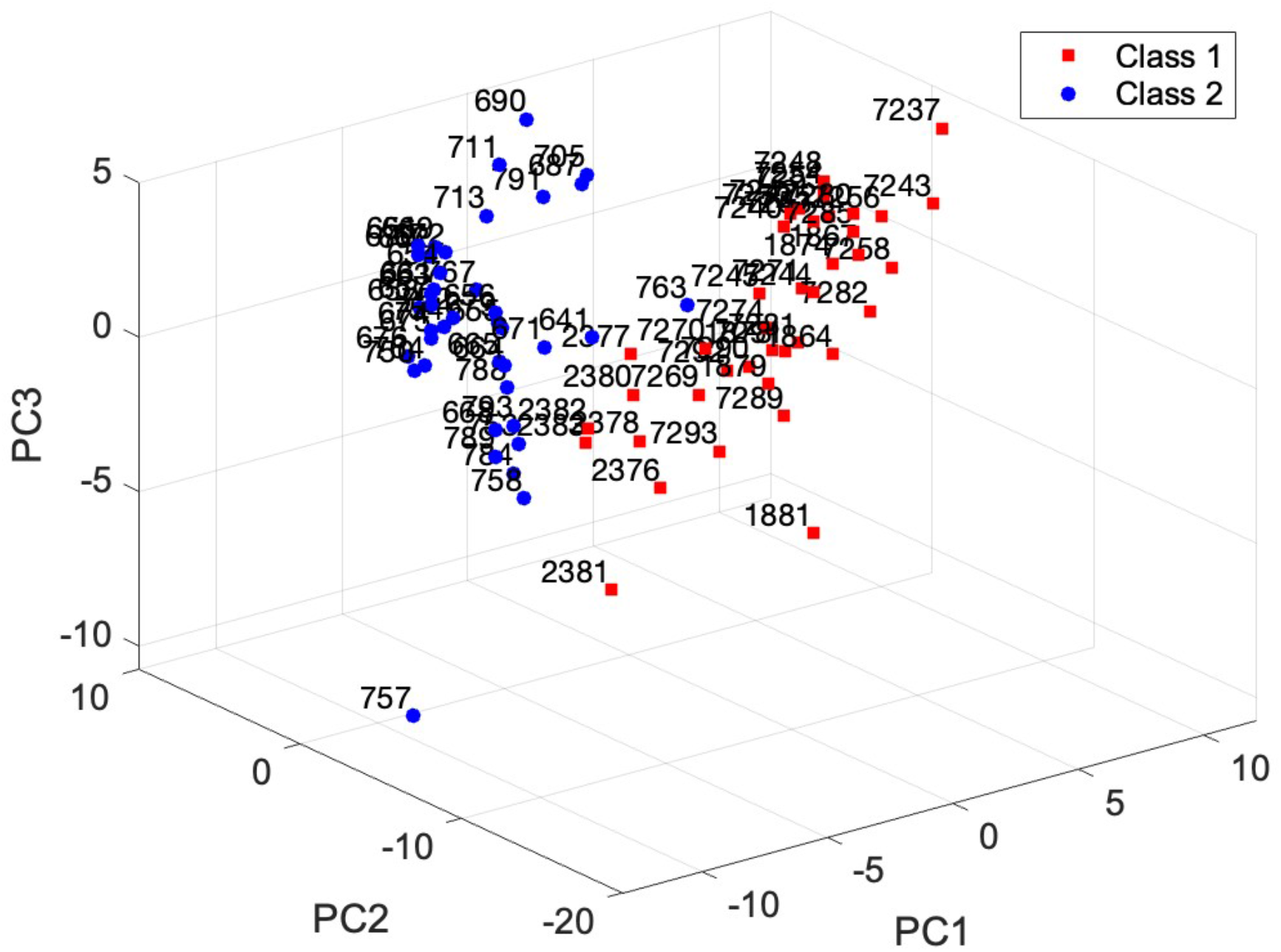
>
> *The plot in* ***Figure 3.4*** *shows that, when viewed with respect to the first 3 PCs, the two outcome classes do not overlap, and good discrimination is achieved. This finding shows that PC3 indeed separates the compounds from* ***Figure 3.3*** *that appeared to overlap. Given the results in* ***Figure 3.2***, *further contributions to discrimination from PC4 (not plotted here) are also expected*.

##### 15. Generate a confusion matrix (a.k.a. truth table)

to quantify the accuracy of the model with respect to correctly assigning the compounds in the Training Set:

~~~
Conf = [model.class_param.conf_mat];
Calc = [model.class_calc];
X_Labels = {‘Class 1’,’Class 2’,’Not Assigned’}
Y_Labels = {‘Class 1’,’Class 2’}
figure()
h = heatmap(X_Labels,Y_Labels,Conf,’Colormap’,[1 1 1],’FontSize’
  ,12)
h.ColorLimits = [min(Conf(:)),max(Conf(:))]
h.Colormap = [1 1 1]
h.FontSize = 16
h.ColorbarVisible = ‘off’
xlabel(‘Predicted’)
ylabel(‘Actual’)
~~~

> *This set of commands generates the output ‘Calc’ which contains the class assignment (i.e. ‘1’ or ‘2’) for the compounds from the Training Set. The command also plots a confusion matrix quantifying the model’s performance, as shown in* ***Figure 3.5***. *The results show that the model correctly assigned all 40 impermeable compounds to outcome Class 1, and all 40 Permeable compounds to Class 2. No compounds from the Training Set gave results that were ambiguous such that the model could not confidently assign the compound to either of the two classes*.
>
> Figure 3.5. The compound assignments made by the PLSDA model from Step 12, which used four PCs, for the Training Set, plotted as a confusion matrix (truth table). The actual (experimental) permeability class of each compound, imported in Step 3 as output variable **y**, is represented by their location in the top versus the bottom half of the matrix, while the class assignment arrived at by the model is represented by each compound’s position left to right. The absence of any compounds in the off-diagonal cells indicates that the model was 100% accurate in assigning the compounds in the Training Set. 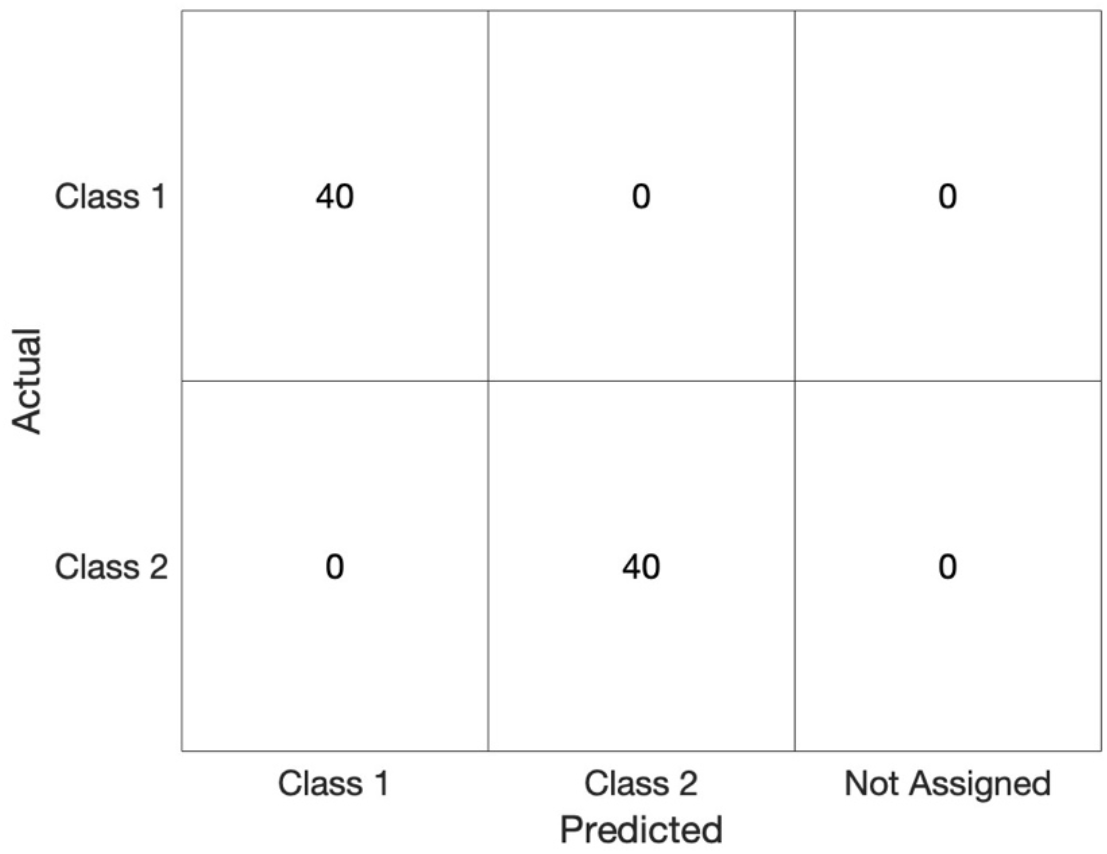
>
> *The perfect accuracy with which the PLSDA model assigns the compounds in the Training Set is a positive outcome. However, the true predictive accuracy of the model is established only when it is used to predict assignments for compounds that were not used in the model building, e.g. the compounds in the Test Set. We will do this test in a later step of the Protocol. First, we will illustrate some ways to understand the model*.

##### 16. Create a biplot of scores and loadings for the PLSDA model

~~~
biplot(model.P(:,1:2),’scores’,model.T(:,1:2),’varlabels’,
  Descriptors)
set(gca,’FontName’,’Arial’,’FontSize’,14)
box on
xlabel(‘PC1’)
ylabel(‘PC2’)
~~~

**Figure 3.6.**
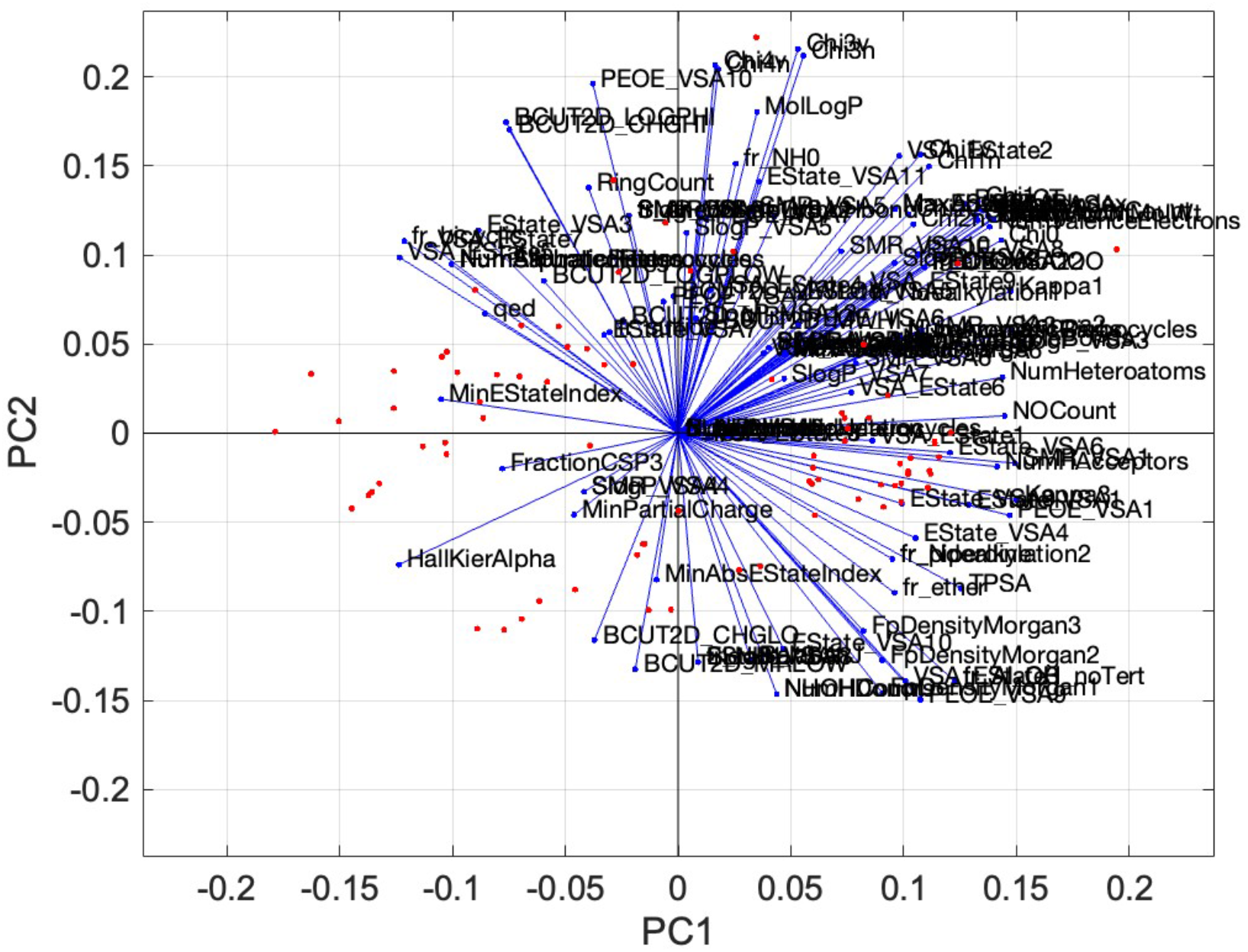
Biplot showing the 80 compound scores and 139 descriptor coefficients (loadings) from the PLSDA model (4 PCs) developed using the Training Set, plotted with respect to PC1 and PC2. Each red datapoint represents the position in PC space of a compound with respect to PCs 1 and 2. The blue lines, superimposed on the same axes, indicate the degree to which each descriptor contributes to PC1 and PC2. Specifically, the blue lines extend from the origin to a point in PC space that has a coordinate in PC1 that corresponds to the orthonormalized value of the descriptor’s coefficient in PC1, and a coordinate in PC2 corresponding to its coefficient in PC2.

*The descriptors that most influence a compound’s score in PC1 are those with the highest (absolute value) coefficient in PC1, identified by the blue lines with terminal points closest to the left and right edges of the plot. The descriptors that contribute most to PC2 are indicated by the blue lines that terminate closest to the top and bottom edges of the plot*. <u>*Important: when plotting a biplot, MATLAB sometimes inverts the signs of the loading (coefficient) values for some components so that the largest absolute Loading value is assigned as positive*. *Therefore, to know the sign of a coefficient it is necessary to look at the data variable ‘P’ rather than reading off the biplot*</u>. *(For additional discussion of biplots, see Basic Protocol 2, Step 14.)*

*The plot in Figure 3.6 is so crowded with descriptors and their labels it is difficult to interpret and has poor visual impact. In the following steps, we describe how to generate more useful biplots that show only a selected subset of descriptors*.

##### 17. Create a biplot showing only the M descriptors having the highest coefficients (absolute value) in PC1

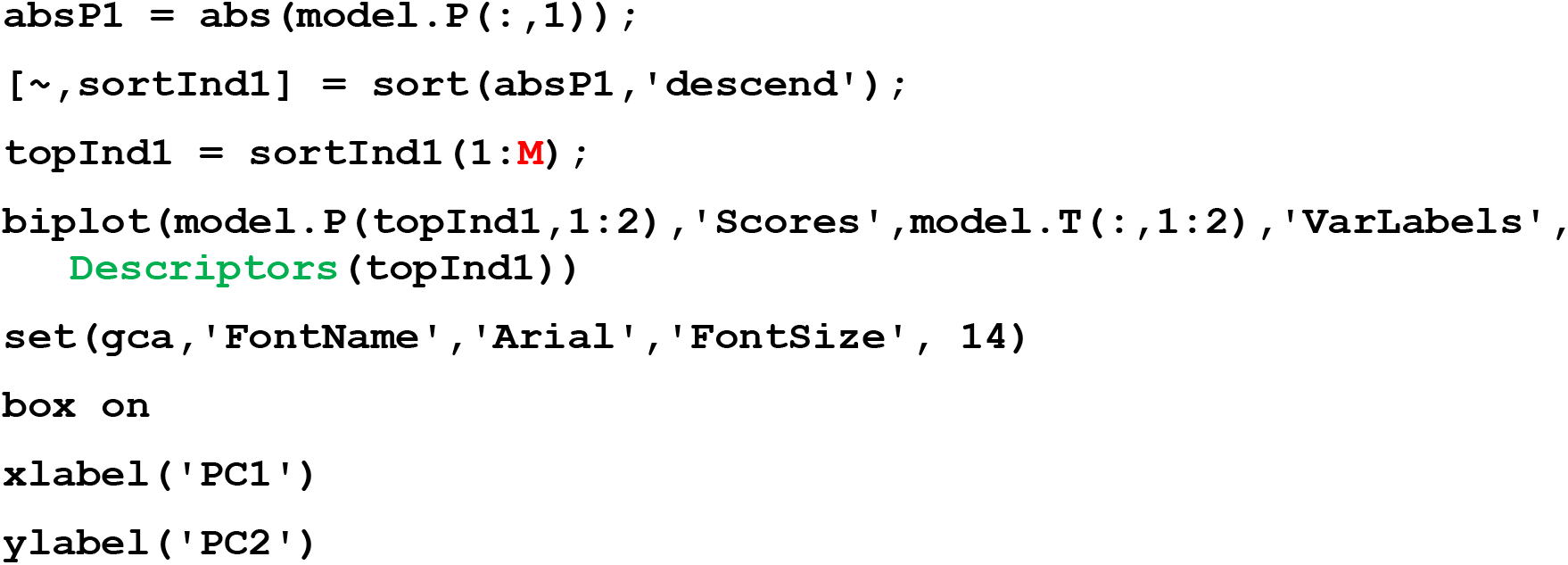

> *In plotting* ***Figure 3.7***, *we used* ***M*** *= 30 to display 30 descriptors*.

**Figure 3.7.**
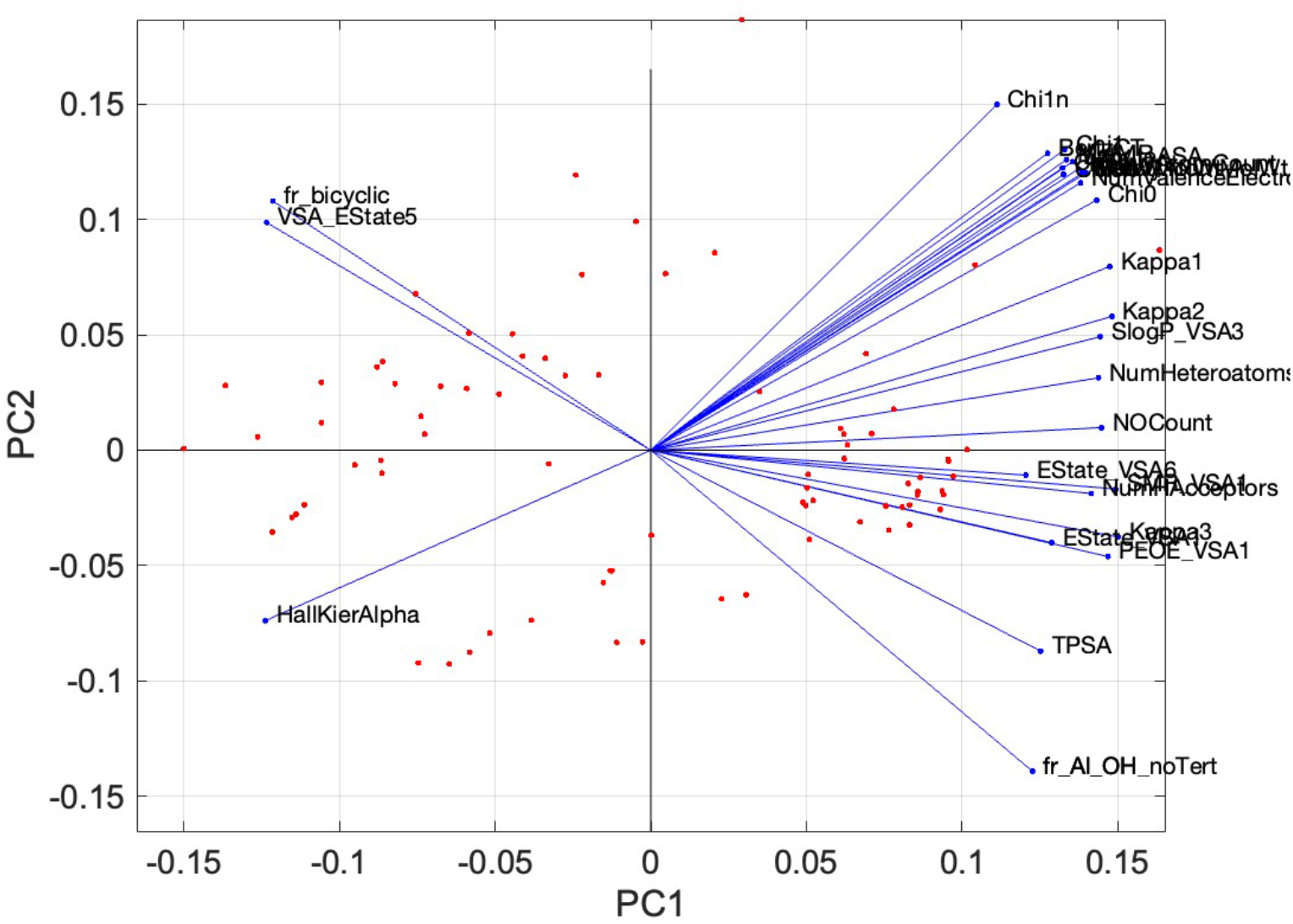
Biplot of the top 30 descriptors with the highest absolute loadings on PC1, as calculated by the PLSDA model from Step 12, indicating the most significant contributors to PC1 for distinguishing between the permeable and impermeable compounds.

##### 18. Create a biplot showing only the M descriptors having the highest coefficients (absolute value) in PC2

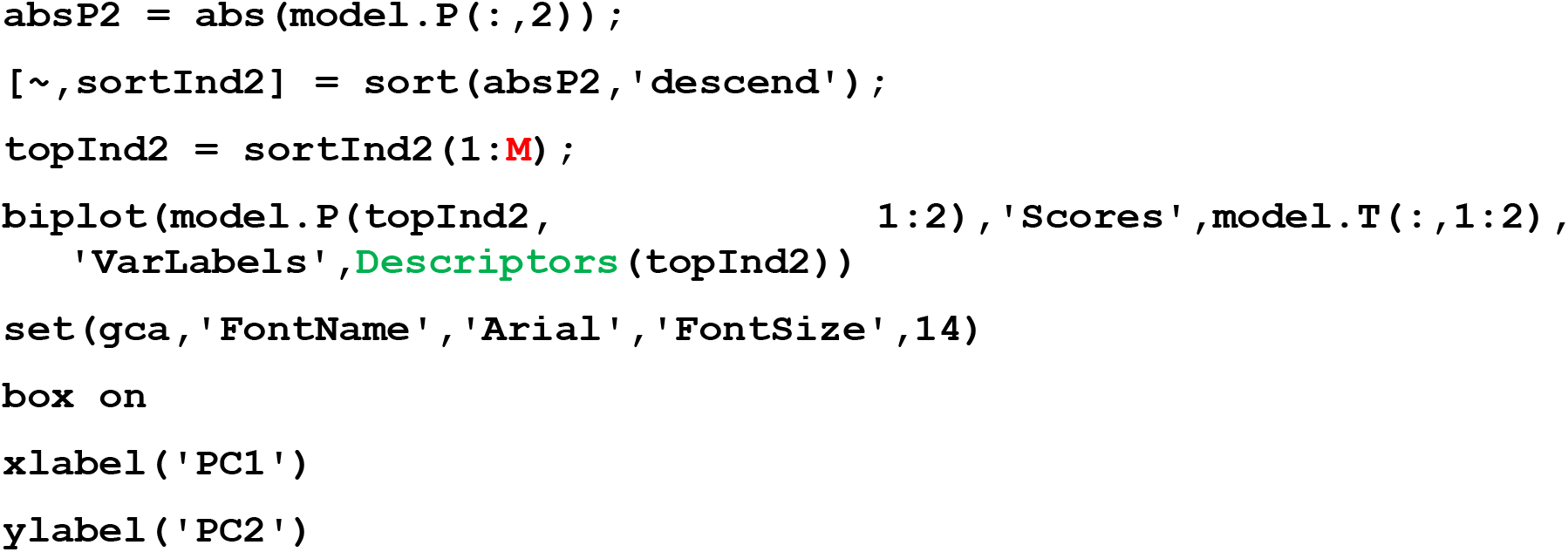

> *In plotting* ***Figure 3.8***, *we used* ***M*** *= 30 to display 30 descriptors*.

**Figure 3.8.**
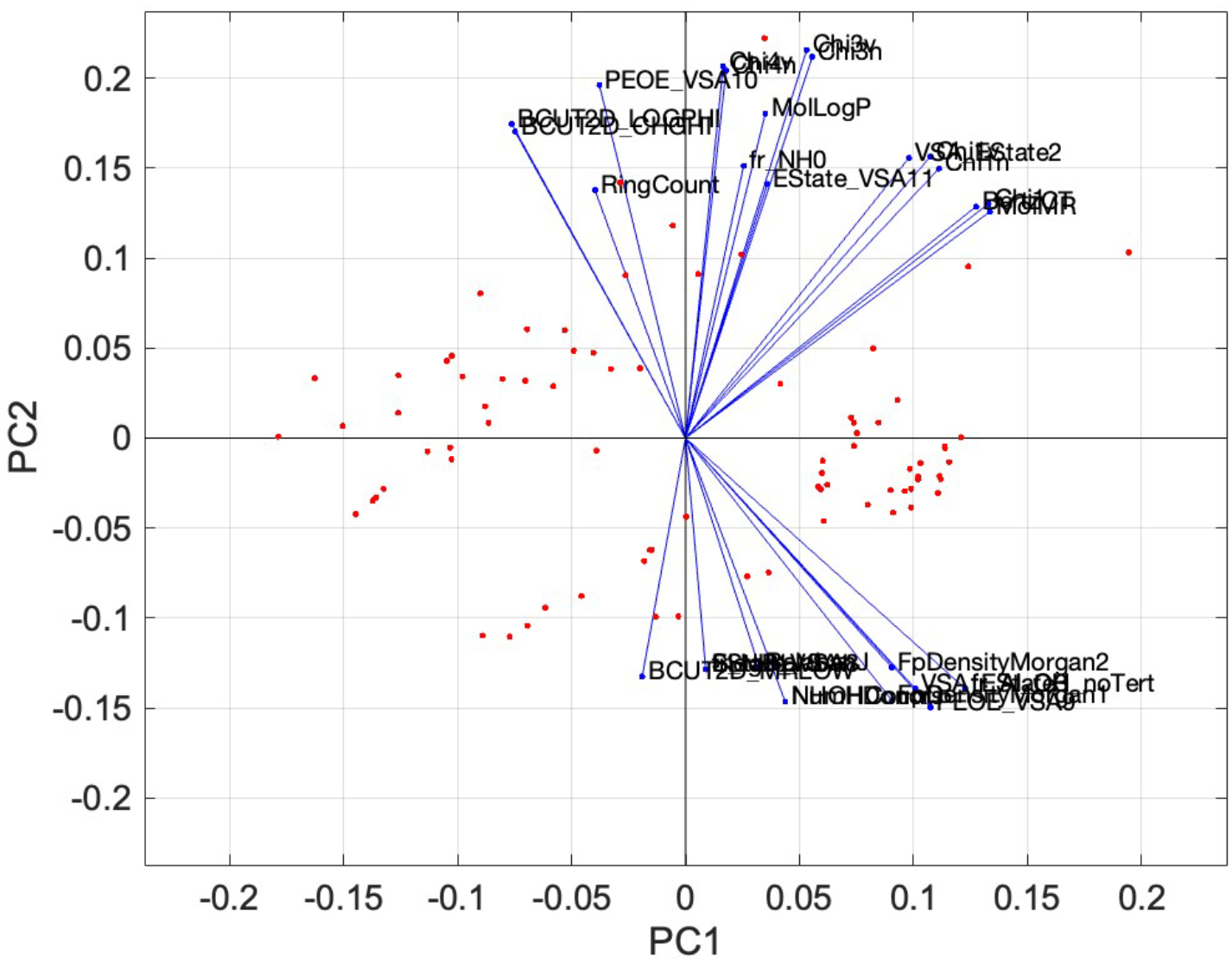
Biplot of the top 30 descriptors with the highest absolute loadings on PC2, as calculated by the PLSDA model from Step 12, indicating the most significant contributors to PC1 for distinguishing between the permeable and impermeable compounds.

##### 19. Create a table showing the descriptors with the 10 highest coefficients (loadings) in a given PC

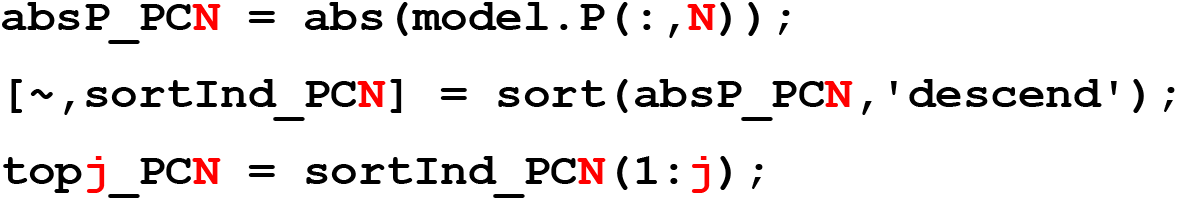
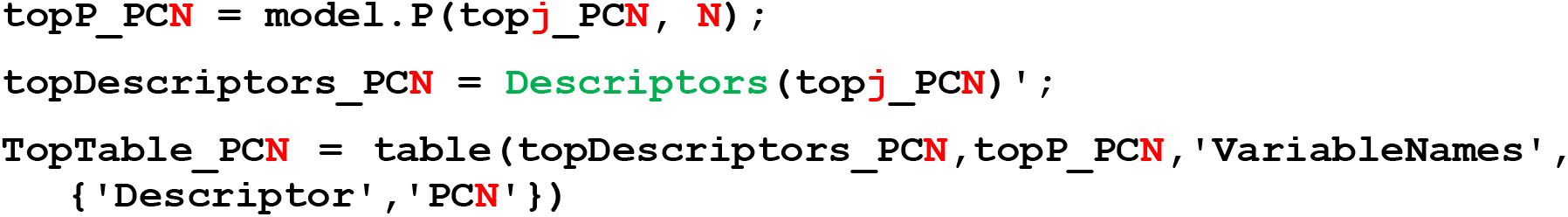

**N** *is the number of the PC in question;* ***j*** *is the number of top elements you wish to include in the table. The other major input is the variable ‘P’ that was generated during PLSDA model building (see Step 12), which is contained in the MATLAB structure ‘model’ that was an output from that step and so is identified here as ‘model.P’*.

> *This command creates a table containing the top 10 most influential descriptors in the PC numbered* ***N*** *and the corresponding coefficients (loadings) for each descriptor ranked by absolute value. We created the table below by running this command four times, once for each of PCs 1-4 (by changing* ***N*** *sequentially from 1 to 2, 3 etc), and combining the results as shown below (****Table 3.1****). The choice of 10 as the number of descriptors to tabulate is arbitrary; a different number of descriptors can be chosen by changing the number ‘10’ in line 3 of the code to the desired number*.
>
> Table 3.1. Descriptors giving the 10 highest (absolute value) coefficients for PCs 1-4. The signs associated with the descriptors have been restored in the table to indicate, to indicated whether an increased value for a given descriptor results in a higher or lower score for the PC. 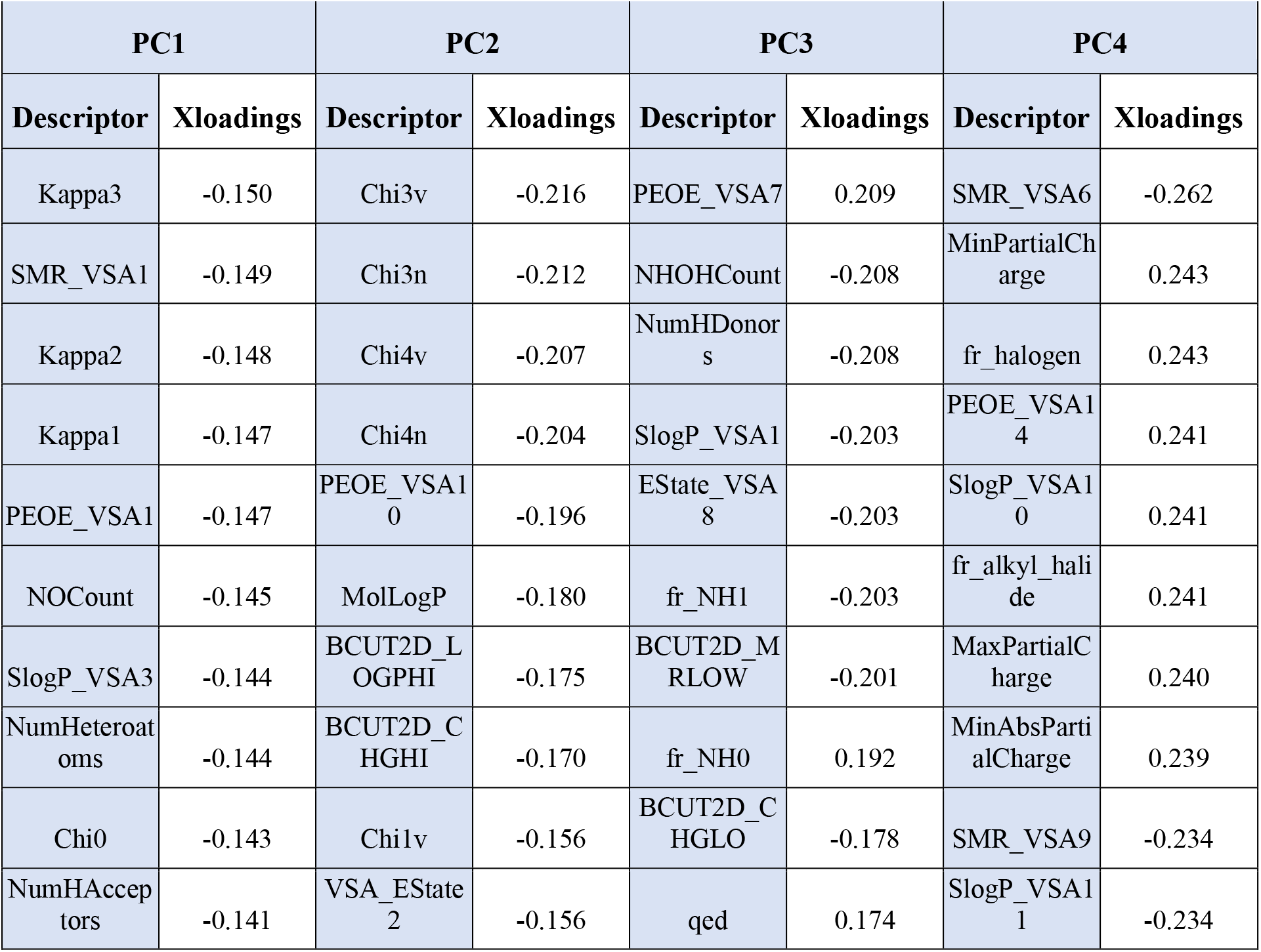

##### 20. Prepare the Test Set of data to use to evaluate the model developed above using the Training Set

(a) Scale and center test data to match the scaling done with the Training Set in Step 8.

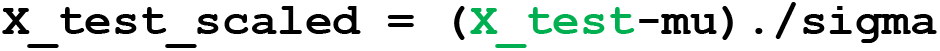

> *In MATLAB, placing a period immediately before a mathematical operation, as here in ‘./’, instructs that the operation be performed separately on each value in the matrix. If the period is omitted, the command would instead divide one matrix by the other via matrix division, which gives a very different result*.

###### Important

*When preparing input data for testing in an already-existing model, it is important to scale using the same mean (‘mu’) and standard deviation (‘sigma’) values that were used to scale the Training Set data when the model was originally developed. This is because the scaled descriptor values for the new data points must use the identical scale of the original data used during model building. If this is not done, then compounds from the Test and Training Sets that had, for example, identical unscaled value for a given descriptor would no longer have the same value once the descriptor was scaled. Hence, rather than simply z-scoring the data here, as we did when building the model in Step 8, we instead manually center and scale the data by subtracting from each descriptor value in* ***X_test*** *the mean descriptor value from the original scaling in Step 8 (‘mu’), and then divide by the standard deviation for that descriptor from Step 8 (‘sigma’)*.

(b) Replace any NaN with zeroes, in case NaNs were introduced during the scaling step

~~~
X_test_scaled(isnan(X_test_scaled)) = 0
~~~

##### 21. Apply the PLSDA model to the Test Set data to predict the permeability of these compounds

~~~
pred = plsdapred(X_test_scaled,model)
result = pred.class_pred
confidence = pred.prob
~~~

*The input data required for the command ‘plsdapred’ comprise (i) the scaled descriptor values in ‘X_test_scaled’, from Step 20(c), and (ii) the PLSDA model in ‘model’, from Step 12*.

*The output of the command is a MATLAB structure called ‘pred’ that contains several different data variables, of which the following are most relevant to our task:*

*‘yc’: A matrix of size* m *x #classes (i.e. 20 × 3 here) containing the predicted class assignments, ‘1’, ‘2’, or ‘0’, for each compound in the Test Set*.

*‘prob’: A matrix of size* m *x #classes containing the probability of belonging to each of the three classes, ‘1’, ‘2’, or ‘0’, for each compound in the Test Set*.

*‘class_pred’: An m x 1 vector giving the predicted class ‘1’, ‘2’, or ‘0’, for the compounds in order of the corresponding rows of* ***X_test***.

*‘class_pred_string’: An m x 1 vector giving the predicted class ‘Permeable’, ‘Not permeable’, or ‘Not Assigned’, for the compounds in order of the corresponding row of* ***X_test***.

*‘T’: An* m *x* n *(i.e. 20 × 139) matrix containing the predicted scores for the Test Set compounds in each PC*.

*‘Thot’: An* m *x 1 vector containing the Hotelling’s T2 statistic for each compound in the Test Set*.

*For information on the other variables in ‘pred’, see ‘plsdapred’ documentation*.

*The last three lines of the command rename some of the variables from ‘pred’ in a more intuitive way. Specifically, the class predictions for each compound from ‘pred.class_pred’ are placed in a new variable named ‘result’, and the probabilities underlying the predicted class assignments from ‘pred.prob’ are placed in a new variable called ‘confidence’*.

**Table 3.2.**
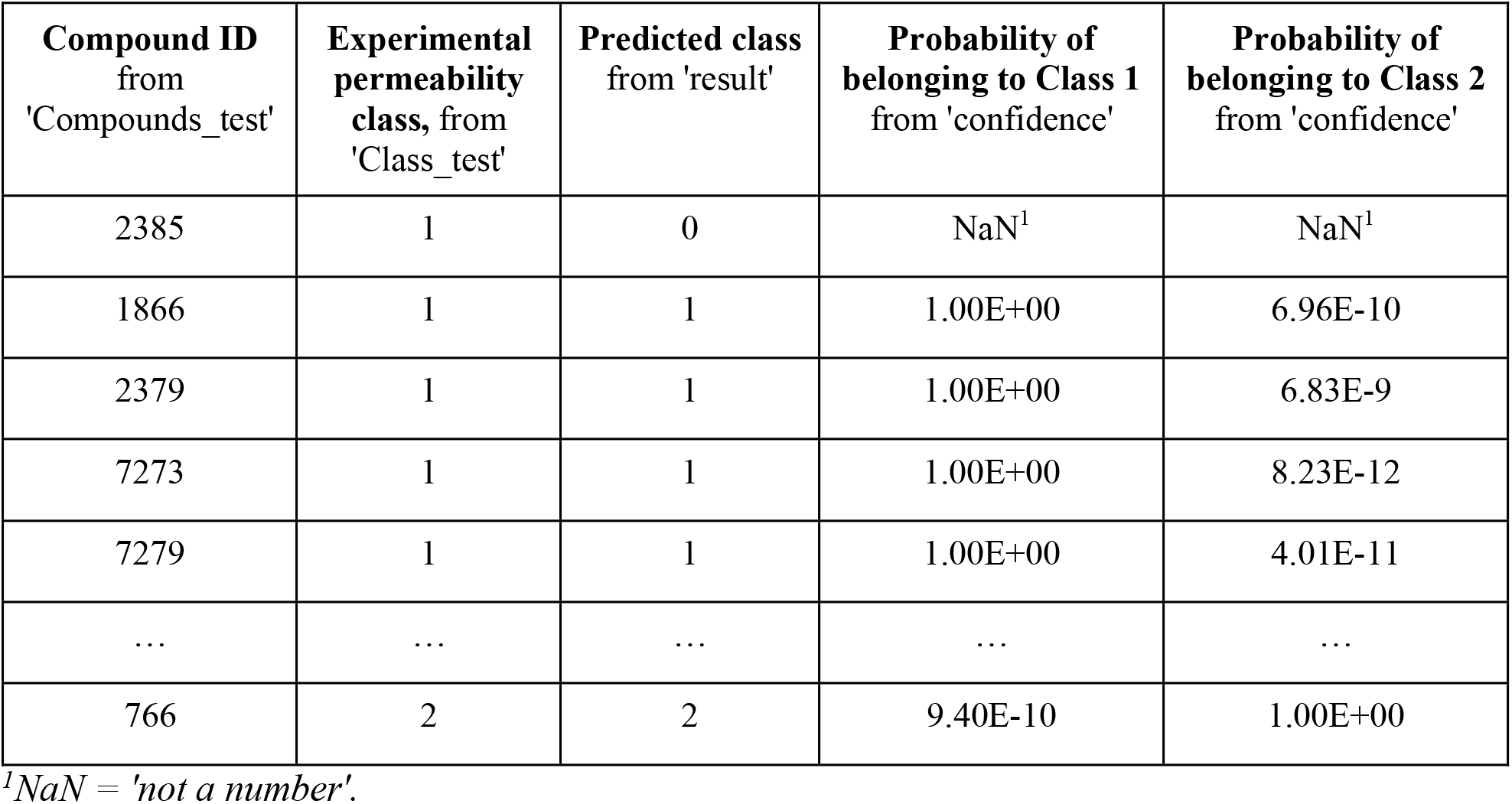
Truncated table showing selected results from the validation of the PLSDA model using the Test Set in Step 21. The table was constructed by putting together the information from the data variables named in each column heading.

| Compound ID<br>from<br>'Compounds_test' | Experimental<br>permeability<br>class, from<br>'Class_test' | Predicted class<br>from 'result' | Probability of<br>belonging to Class 1<br>from 'confidence' | Probability of<br>belonging to Class 2<br>from 'confidence' |
| --- | --- | --- | --- | --- |
| 2385 | 1 | 0 | NaN<sup>1</sup> | NaN<sup>1</sup> |
| 1866 | 1 | 1 | 1.00E+00 | 6.96E-10 |
| 2379 | 1 | 1 | 1.00E+00 | 6.83E-9 |
| 7273 | 1 | 1 | 1.00E+00 | 8.23E-12 |
| 7279 | 1 | 1 | 1.00E+00 | 4.01E-11 |
| ... | ... | ... | ... | ... |
| 766 | 2 | 2 | 9.40E-10 | 1.00E+00 |

<sup>1</sup>NaN = 'not a number'.

##### 22. Generate the confusion matrix (the truth table) to evaluate the robustness of the prediction

~~~
grouporder = [1 2 0];
ConfMatTest = confusionmat(Class_test,result,’Order’,grouporder); X_LabelsTest = {‘Class 1’,’Class 2’,’Not Assigned’};
Y_LabelsTest = {‘Class 1’,’Class 2’,’Not Assigned’};
figure
h = heatmap(X_LabelsTest,Y_LabelsTest,ConfMatTest,’Colormap’,[1 1
  1],’FontSize’,12);
h.ColorLimits = [0 max(ConfMatTest(:))];
h.Colormap = [1 1 1];
h.FontSize = 16;
h.ColorbarVisible = ‘off’;
xlabel(‘Predicted’);
ylabel(‘Actual’);
~~~

*The finding, from* ***Figure 3.9***, *that the PLSDA model was able to correctly assign the permeability of the compounds in the Test Set – which were not used in the model building – with 90% accuracy indicates that the model has good predictive power, at least for the kinds of cyclic peptide compounds included in this analysis. For a more detailed discussion of interpreting the results of PLSDA, see the following section*.

**Figure 3.9.**
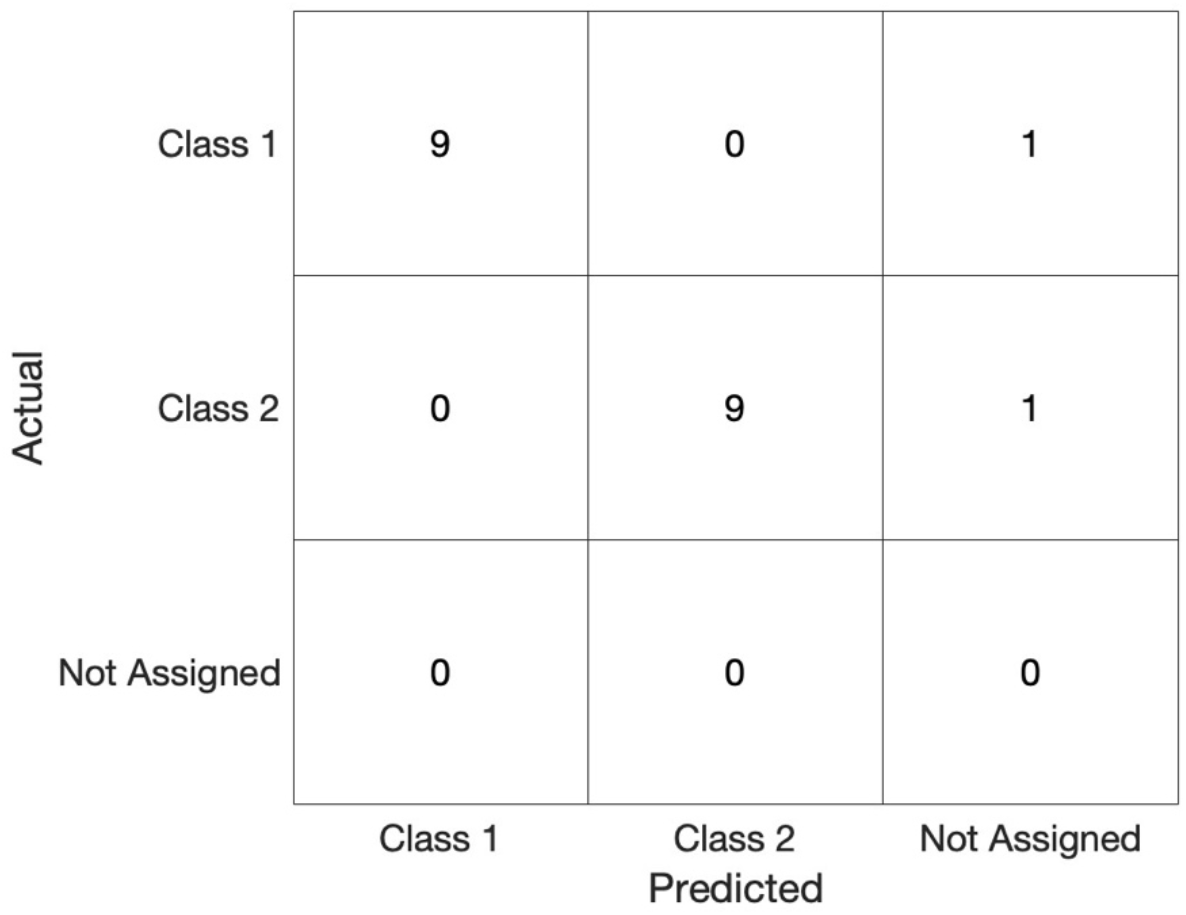
Confusion matrix showing the accuracy with which the PLSR model developed in Step 12 predicted the permeability of the compounds in the Test Set. The results show that 9 out of the 10 Impermeable compounds were correctly assigned as such, as were 9/10 of the Permeable compounds. No compounds were erroneously mis-assigned to the wrong set; the model was unable to assign the remaining two compounds.

*For the current situation the output behavior categories take the form of a positive versus a negative result (e.g. ‘Permeable’ versus ‘Not Permeable’). In such cases the performance of the model can be described in terms of its true positive rate, or ‘selectivity, which is the fraction of elements that in reality belong to the positive category that are rated by the model as positive, and its true negative rate or ‘sensitivity’, which corresponds to the fraction of actual negative elements the model correctly rates as negative. The truth table in* ***Figure 3.9*** *indicates that the PLSDA model developed here has a selectivity of 9/10 = 0.9, and a specificity that also equals 9/10 = 0.9*.

##### 23. Save your final workspace as a .mat file

*This file can be reopened to give a MATLAB workspace identical to the state that was saved*.

### Understanding the results of Basic Protocol 3

Many of the principles underlying PLSDA are related to PCA. For a full understanding of PLSDA, we recommend reading the ‘Introduction’ to Basic Protocol 2 and the additional sections ‘Understanding the Results of Basic Protocol 2’ and ‘Summary comments on PCA’ as background before tackling the following material.

### What PLSDA does

Like the technique of PCA discussed in Basic Protocol 2, PLSDA and the closely related method PLSR exploit covariance to extract information from complex data sets. The underlying notion is that to the extent two or more descriptors are covariant with each other, they are capturing the same information about the elements in the data set. Descriptors are rarely 100% covariant (i.e. PCC = +1 or -1), and so it is only the covariant portion of the information from each that is redundant. PCA, an unsupervised method, simply looks for the greatest amount of covariant information in the input matrix *X* and calls this PC1. This process is equivalent to plotting all the data in *X* on a set of Cartesian axes, one axis for each of the *n* descriptors in *X*, and then drawing a straight line through the origin of the resulting *n*-dimensional cloud of data points in the direction that captures the most variance – generally speaking, through the longest axis of the data cloud. PC2 is a second new axis, orthogonal to (i.e. having zero covariance with) PC1, that captures the largest amount of remaining variance that PC1 did not capture, and so on for PCs 3, 4, etc. As a supervised ML method, PLSDA instead looks for such redundant information in the input data matrix *X* that is most covariant with the data in the output data matrix *y*, on the grounds that this portion of information from *X* is most influential in determining (‘explaining’) what category of the outcome behavior *y* will be observed. The resulting new axis is called PC1. PC2 is a second new axis, orthogonal to PC1, that covaries as much as possible with the remaining unexplained variance in *y*, etc.

The main outputs of PLSDA are (i) the scores of the data elements with respect to PCs 1, 2, etc., (ii) the coefficients (loadings) that indicate how much each of the original descriptors contributes to each PC, (iii) the output behavior category (in this case, ‘Permeable’ or ‘Not Permeable’) that the model assigns to each element in the data set, and (iv) what confidence the model associates with the assignment of each element to the different possible output categories.

### Avoiding overfitting with PLSDA

As described in Step 11, cross-validation is an important step in choosing the number of PCs to include in model building that best balance utilization of the portion of the input data that truly contributes to discriminating the output behaviors versus the portion of the data that is irrelevant or stochastic noise. The optimum number of PCs depends in part on how much redundancy exists among the outcome-driving descriptors in the input data and, thus, how much dimensionality reduction can be achieved, as well as how distinct the properties of the elements in the different output categories are with respect to the provided descriptors. The sensitivity (see Step 22) of the model will tend to increase as more PCs are included, but selectivity will suffer as addition of more PCs increases the amount of noise is incorporated; 100% sensitivity can trivially be achieved by assigning all elements as positive! This phenomenon is known as ‘overfitting’. Cross-validation is thus critical to avoiding overfitting and achieving a model with the highest possible predictive accuracy. While the relative importance of sensitivity and selectivity for a model will depend on its purpose, in general the number of PCs that results in the confusion matrix that best captures the desired results is the number of PCs to use in model building.

### How to interpret PLSDA scores

As is the case with PCA (See Basic Protocol 2), the scores generated by PLSDA are simply the coordinates of the different elements in the input data set with respect to the PC axes. If the PLSDA model has been effective in differentiating the elements in the different output categories (i.e. the permeable and impermeable compounds), then the data points corresponding to these compound groupings should be well separated when plotted in PC space. **Figure 3.4** shows this to be the case for the Training Set data analyzed in this protocol. If the data sets were to overlap in PC space, this would indicate that the model does not perfectly discriminate between the different categories, and the greater the overlap the poorer the discrimination that was achieved. The scores plots, therefore, provide a visual assessment of the performance of the model, an assessment that is quantitatively captured in the confusion matrices in **Figures 3.5** for the Training Set and **3.9** for the Test Set.

### How to interpret PLSDA loadings (coefficients)

Like the related methods of PCA and PLS-DA, interpreting the coefficients in terms of the importance of which properties of the elements in the data set drive differences of interest – in this case, differences in the value of the output behavior *y* – is somewhat nuanced. The basic principle, that each PC represents a distinct feature of the elements in the set, and that the descriptors with the highest coefficients (often called ‘Loadings’ in PLS methods) in a given PC are driving the scores in that PC, are straightforward. But the reduction in dimensionality that is central to these ML techniques means that each PC is a composite of covariant information from a number of the original descriptors, often making it difficult to interpret what over-arching feature the PC represents.

The reader is referred to ‘Understanding the results of Basic Protocol 2’ for a more detailed discussion of approaches to using descriptor coefficients to discern the theme underlying a given PC. Here, we will only mention that for supervised ML methods other approaches for determining the overall importance of the contributing descriptors also exist. One such method is Variable Importance in Projection (VIP) (*Favilla et al*., *2013*). A VIP function has recently been added to MATLAB. In broad terms, VIP sums the influence of each of the original descriptors across all PCs used in model building to arrive at a ranked list of the cumulative influence of each descriptor on the model as a whole. The descriptors with the highest VIP scores (conventionally, scores >1) are considered most important for explaining the output behavior *y*. VIP does not solve the problem of redundancy among the original descriptors, still leaving it as a problem for the investigator to discern whether a given descriptor has a high VIP score because it is causative in determining the value of y, or just covaries with a causative descriptor. Nonetheless, it is undoubtedly a valuable method for determining the overall influence of the descriptors across all relevant PCs.

### Evaluating the predictive accuracy of PLSDA models

So-called ‘classification’ methods of supervised ML, such as PLSDA, serve to categorize the elements of the data set into discrete groups based on their output behavior. A common way to analyze the performance of classification models is by means of a ‘confusion matrix’, also known as a ‘truth table’, as illustrated in **Figures 3.5** and **3.9**. A confusion matrix comprises a grid, the squares of which contain digits that express the number of elements in the set that were assigned to their correct category versus erroneously assigned to an incorrect category versus could not be assigned with confidence. In the current case, when applied to the Training Set the model achieved perfect discrimination between the permeable and impermeable compounds, showing 100% sensitivity and 100% selectivity (see Step 22 for definitions of these terms). When applied to the Test Set of compounds, however, both sensitivity and selectivity dropped to 90%. It is quite typical for a model to perform less well with the Test Set compared to the Training Set that was used to build it. The extent to which these numbers differ between the Training and Test sets is to some extant a measure of the degree of overfitting in the model, but is also sensitive to the presence of extreme outliers especially when the data sets are small as in our case. The confusion matrix seen with the Test Set should be considered the best guide to the predictive accuracy of the model. In our case, the 90% for both sensitivity and selectivity achieved with the Test Set of compounds indicates that the model predicts membrane permeability with good accuracy, at least for compounds that closely resemble those present in our data set.

### Limitations of PLSDA

An important limitation of PLSDA is that the resulting model is reliable only for elements that closely resemble those in the Training Set. The extent to which different kinds of differences in the properties of the elements might lead to inaccuracy is unknowable, because this information was not available to the model. Thus, a PLSDA model will likely have the assessed level of predictive accuracy when used interpolatively, to assess the output behavior of elements with properties that fall within the property-space sampled by the elements contained in the Training Set data, but cannot be relied upon to extrapolate to predict the behavior of elements with properties that fall outside of this space. This property space is sometimes referred to as the model’s ‘domain of applicability’.

For example, in the analysis illustrated in this protocol, the only permeable compound that the model was unable to assign with confidence was compound ID2385, and the sole impermeable compound that could not be correctly assigned was ID828. Both of these compounds were among the top outliers identified by clustering (Basic Protocol 1a) and PCA (Basic Protocol 2). Taken together, these findings suggest that the reason these two compounds could not be categorized by the PLSDA model is because they are structurally somewhat distinct from the 80 compounds in the Training Set on which the model was trained. Thus, categorization for these two compounds failed likely because they fall outside the domain of applicability of the model. In general, elements for a data set that show as outliers from the Training Set in unsupervised ML analyses such as clustering or PCA are least likely to be correctly categorized by the model.

Another limitation of PLSDA, and all similar methods, is that the predictive accuracy of the model is critically dependent on choosing a set of input descriptors that contain the information required to explain the behavior of interest, For example, a model designed to predict which individuals will run 100 m in the shortest time is highly unlikely to be predictively accurate if the information provided about the individuals in question doesn’t include information on their health, physique, etc.; information about their music preferences, favorite authors, etc. will not lead to a useful model. While this point may seem trivial, when choosing what descriptors to use in PLSDA, it is critical to keep in mind that the descriptors embody a hypothesis that the output behavior can be *explained* in terms of the input data.

It is also important that the Training Set used to build the model contains similar numbers of elements in each output behavior category. A major imbalance in numbers will bias the model towards prediction of one class versus the other. The presence of extreme outliers will also hamper the development of an accurate model. In this sense, an outlier can be considered as a data element for which the output behavior depends on different properties than are drivers of the behavior of the bulk of the data set. Inclusion in the Training Set of a subset of elements that behave according to different principles than the majority of the set will necessarily obfuscate the factors important for the behavior of the majority cases.

Finally, is important to know that PLSDA embodies the tacit assumption that the different descriptors combine to explain the output behavior of the elements in the data set only in linear ways. Situations in which descriptor A is important when a second descriptor B has certain values but not when it has others are not well-handled by PLSDA or related methods such as PLSR.

## BASIC PROTOCOL 4: Partial Least Squares Regression (PLSR)

### Purpose of method

To build a model that can accurately quantify an observable or measurable behavior, based on the properties of each item, where the behavior to be predicted has a continuous numerical value rather than being a categorical outcome. For example, PLSR could be used to analyze the doubling time of bacterial growths (the output variable) based on a set of input variables such as the concentrations of different components in the growth medium, the growth temperature, pH, etc..

### Background

PLSR is conceptually related to the unsupervised ML method of Principal Component Analysis (PCA). It takes as input data the numerical values of a set of *n* properties for each member of a set of *m* elements, and transforms these data to eliminate redundant information and thereby generate a smaller number of composite properties (Principal Components) that allow the information to be analyzed at a lower dimensionality. In our case, the input data (‘**X**’) comprise values of 127 different molecular properties for each of the different cyclic peptides in our compound set, and the output data (‘**y**’) are experimental values for the compounds’ passive permeability through a lipid membrane. As discussed in more detail in Basic Protocol 2, many of the molecular properties provided in **X** will likely be to some extent covariant, and thus contain a degree of redundancy in the information they contain about the molecules. By condensing this information into a smaller set of mutually orthogonal Principal Components, each of which corresponds to a distinct and nonredundant portion of the input data, the same information can be visualized and analyzed using fewer axes. The result of PLSR is a model that can be used to predict the output behavior (i.e. permeability) of new elements (i.e. compounds), based on their input properties.

As is generally the case with supervised ML, the Protocol below generates a model using a Training Set comprising 80% of the available data, and this model is then validated by testing how accurately it predicts the permeability of the 20% of the data that was held back to serve as a Test Set.

Key considerations for PLSR, as for PLSDA, are (i) how to divide the available data into the Training and Test Sets, (ii) how many components to use in the final model building, (iii) how to assess the predictive accuracy of the model, (iv) how to judge the model’s ‘domain of applicability’ (i.e. for what kinds of compounds can the model be expected to accurately predict permeability?), and (v) and how to determine which properties are most important in driving differences in the output behavior.

#### Before you start

Be sure you have installed the necessary toolboxes mentioned in the ‘REQUIRED SOFTWARE’ section.

The stepwise instructions below assume that, for the Training Set and Test Set, you will be using the data we provide with this article (Supplementary file Data_PLSR.xlsx). If using your own dataset, make sure you have formatted the data for this analysis as described in the ‘DATA DOWNLOAD AND PREPARATION’ section (see **Figure 4.1**).

The Test and Training Sets used in this protocol were derived from the set of 201 compounds used in Basic Protocols 1 and 2, but with inclusion of the membrane permeability (PAMPA) data for the compounds. To make the Test Set, we used *k*-medoids clustering (Basic Protocol 1a) to select 40 compounds that were representative of the whole set. The remaining 161 compounds from the original set were designated as the Training Set. After selecting the compounds for the Test and Training Sets, we then reviewed the property values associated with both to ensure that no descriptor had identical values across all compounds. We did this because inclusion of descriptors with zero variance creates errors in the calculation. Twelve descriptors were found to have identical values across one or other of the compound sets. These were manually deleted from the data matrix, leaving 127 descriptors. The final dimensions of the Test and Training Set input data files are thus 40 × 127 and 161 × 127, respectively. A detailed description of how we generated the Training Set and Test Set in Data_PLSR.xlsx is provided the supplementary document (Data_prepared_SI.xlsx).

##### 1. Gather the input data

Details of the data download and preparation are given above in Section IV, and spreadsheets containing the data required to perform the protocols are included in the supplementary file ‘Data_PLSR.xlsx’. For this protocol, these comprise the values of 127 molecular features (descriptors) for a Training Set of 161compounds plus a Test Set of 40 compounds that will be held back from model building and instead used to evaluate the predictive accuracy of the model once built. The supplementary file ‘Data_PLSR.xlsx’ contains the data already formatted in this way, in the sheets named ‘PLSR_Train’ and ‘PLSR_Test’.

*If you are using your own data rather than the supplementary file provided, be sure to format your external data files as described in ‘Section IV. DATA DOWNLOAD AND PREPARATION, and illustrated in* ***Figure 4.1***, *and to change the external filenames (blue text) in the following steps to match the names you have assigned to your files*.

**Figure 4.1:**
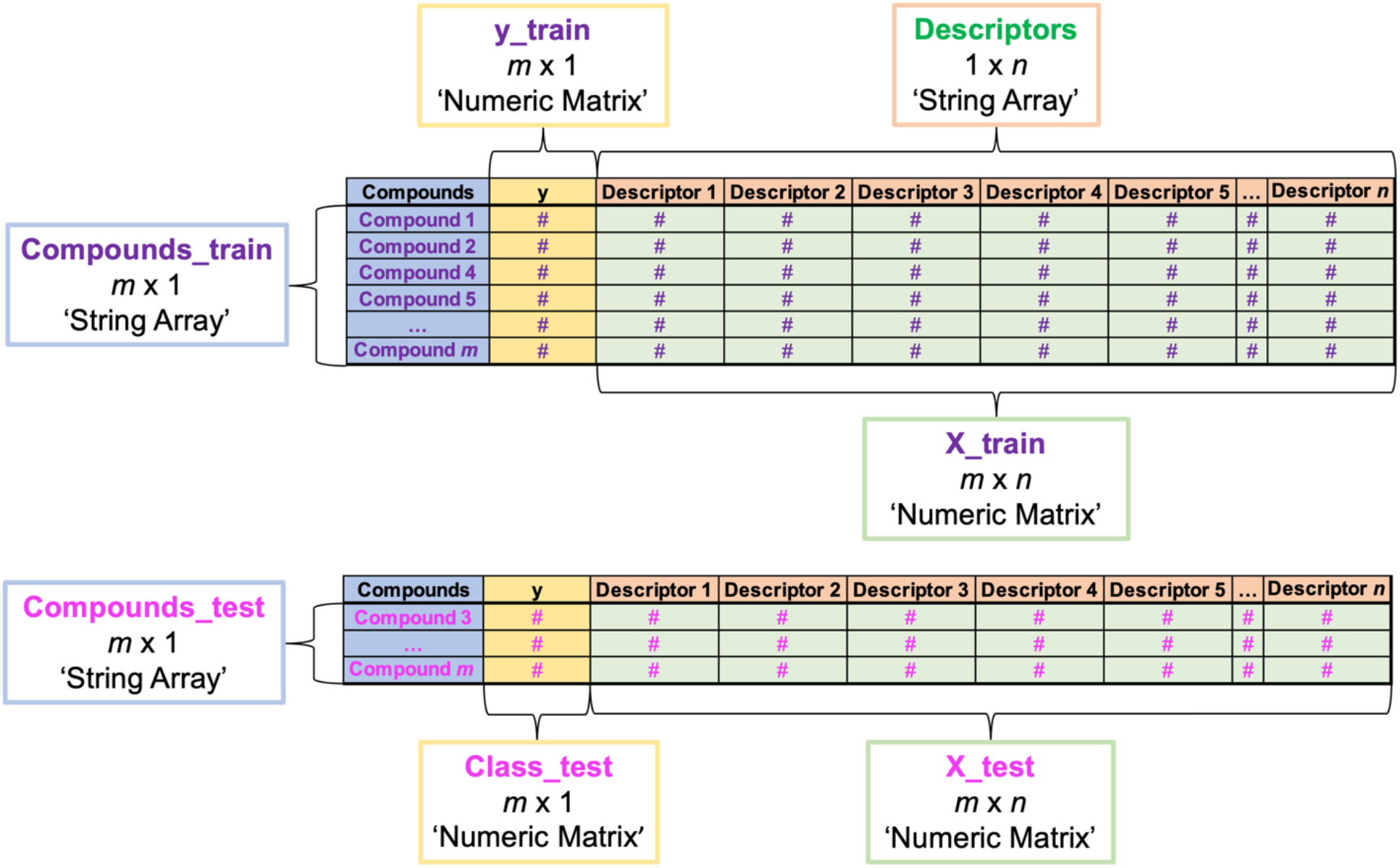
How to format data for import into MATLAB for PLSR analysis. Text in green indicates the filenames we assigned to the different data variables created during import, and that we refer to in Basic Protocol 4. Black text in single quotes indicates what import file type to choose when importing each piece of data using the MATLAB Import Data window. If you are following the example in the Protocol, you will simply import data we have already formatted as shown above from two tabs (train and test) in the supplementary file Data_PLSR.xlsx.

*If performing PLSDA on other types of data, the compound names in column 1 would be replaced by identifying labels for the different elements in the set, and the descriptor names in row 1 would be replaced by the names of the different properties that describe the elements*.

##### 2. Import the numerical input data, X_train, for the Training Set of compounds

(i) Starting with a clear workspace, click on the Import Data button in the MATLAB Home tab (top of screen), navigate to where you have saved the Supplementary File ‘Data_PLSR.xlsx’.
(ii) Working in the Import Data window, open the tab labeled ‘PLSR_train’. <u>Be sure you are on the correct sheet before you begin importing data.</u>
(iii) Select the cells that contain the numerical data. <u>Be sure to exclude the column headers in Row 1 as well as the compound names and PAMPA values in columns 1-2</u>. *The selected data should comprise 161 rows (reflecting the 161 compounds in the Training Set) and 127 columns (reflecting the 127 molecular descriptors we are using for this analysis). The selected data should include only numbers*.
(iv) Go to ‘Output Type’ at the top of the Data Import window and select ‘Numeric Matrix’ from the drop-down menu, then go to the Import Selection button at the top right of the Import Data window and click the green check mark (or select ‘Import Data’ from the drop-down menu). You should see a new data object with dimensions 161 × 127 in the Workspace window of your MATLAB desktop (you may need to move the Import Data window aside to see it).
(v) Rename the 161 × 127 workspace variable. For the purpose of this exercise, you should call the new workspace variable **X_train**. You can do this in the open Data Import window by double clicking on the default name (same as the original Excel filename) that appears just below the top row of the window and typing in the desired new name. Alternatively, you can click on the newly created variable in the MATLAB Workspace window to highlight its default name, and type in the new name, **X_train**.

*Data variables created in MATLAB can be renamed after the fact, as described in Step 2(v), but it is best practice to assign the desired name in the Data Import window before clicking ‘Import Data’ to create the variable. The reason is that the default name that MATLAB assigns to new data variables is the name of the external file from which the data were imported, making it easy to accidentally over-write a different data variable previously imported from the same external file if it was not assigned a unique name upon creation*.

##### 3. Import the output data, *y_train*, for the Training Set

Repeat the selection process from Step 2, but this time selecting the data from Column B, labeled ‘PAMPA’, in the ‘PLSR_train’ tab of Data_PLSR.xlsx. Change the ‘Output Type” to ‘Numerical’, rename the data variable as **y_train**, and then select ‘Import Data’. A new 161 × 1 data variable, **y_train**, will appear in your Workspace window.

*Import only the numerical values, being sure to exclude the header name (‘PAMPA’) in Cell 1B*.

##### 4. Import the compound names for the Training Set

From the ‘PLSR_train’ tab of Data_PLSR.xlsx, import the compound names from Column A. Change the ‘Output Type’ of the matrix to ‘String Array’, and then select ‘Import Data’ to generate the array. Rename the new 161 × 1 string array that appears in the Workspace window as **Compounds_train**.

*Exclude the column header ‘CycPeptMPDB_ID’ in cell A1, as this is not a compound name*.

##### 5. Import the descriptor names for the Training Set

From the ‘PLSR_train’ tab of Data_PLSR.xlsx, select the descriptor names (i.e. the column headers that appear in Row 1). Change the ‘Output Type’ of the matrix to ‘String’, assign the name **Descriptors** to the new data variable, and then select ‘Import Data’ to generate a new 1 × 127 string array in the Workspace window.

> *Be sure to exclude the column headers ‘CycPeptMPDB_ID’ and ‘PAMPA’ in cells A1-B1, as these are not descriptor names. If you are using the Data_PLSR.xlsx data file provided with this article, this means you should start your selection with cell C1 (‘MaxEStateIndex’)*.

##### 6. Import the numerical input data (X_test), output data (y_test), and compound names for the Test Set

Use the Data Import window to navigate to the sheet called ‘PLSR_test’ in the file Data_PLSR.xlsx, in the same manner as in steps 2-4, naming the data variables **X_test** (40 × 127), **y_test** (40 × 1), and **Compounds_test** (40 × 1), respectively.

> *Be sure you are on the correct sheet, ‘PLSR_test’, before importing these data. There is no need to import the descriptor names again, as these are identical for the Test Set and the Training Set*.
>
> *After performing Steps 1-6, your MATLAB workspace should contain the following 7 data variables:* ***Compounds_test, Compounds_train, Descriptors, X_test, X_train, y_test***, *and* ***y_train***.

##### 7. Save your MATLAB workspace as a .mat file

*This file can be reopened to give a MATLAB workspace identical to the state that was saved*.

##### 8. Scale and center the X_train data values

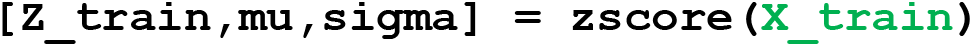

> *Z-scoring centers the values for each descriptor on zero and scales the variance to a range of SD = 1. (See Basic Protocol 1a, Step 7 for more information about z-scoring.)*
>
> *Refer to Basic Protocol 1a Step 8 for how to generate boxplots for visualizing the results of z-scoring*.

##### 9. Replace any ‘NaN’ (‘Not a Number’) results from Step 8 with zeroes

~~~
Z_train(isnan(Z_train)) = 0
~~~

*NaN entries result from invalid mathematical operations such as division by zero, and their presence can prevent the proper operation of some subsequent commands. If you are using the data sets provided with this article, no NaN values will be generated. However, we include this step in case you are following this protocol with your own data set, for which NaN results could occur*.

##### 10. Calculate the covariance between descriptors

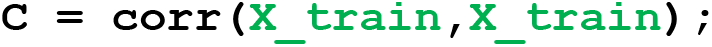

*This command creates a new data variable, C, that has the form of an* n *x* n *matrix (127 × 127 in our case) containing the Pearson Correlation Coefficients (PCC values) between each descriptor and every other descriptor in* ***X_train***. *Examination of this table shows which descriptors are most covariant with each other, and how covariant the information in the descriptor set is as a whole. The higher the covariance, the greater the redundancy in the input data, and the more PLSR will be able to reduce the dimensionality of the data to a small number of Principal Components containing the bulk of the information in* ***X_train*** *that is relevant to explaining the values in* ***y_train***. *(For a more detailed explanation of covariance and PCC, and of their relationship to dimensionality reduction, see the Introduction to Basic Protocol 2.)*

##### 11. Create two variables to represent the number of rows (compounds) and descriptors (columns)

~~~
[m,n] = size(Z_train)
~~~

*In our example, ‘m’ is the number of compounds in the training set (i.e. 161) and ‘n’ is the number of molecular descriptors (127)*.

##### 12. Calculate PLSR for numbers of Principal Components from 1-20

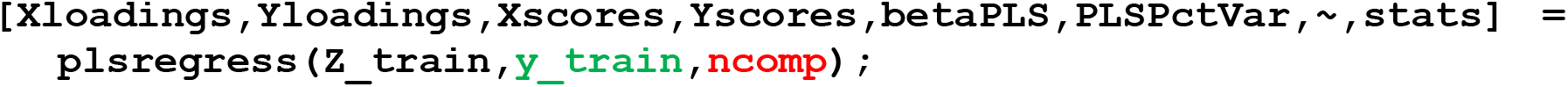

*As inputs, the ‘plsregress’ command requires (i) an* m *x* n *matrix of input data (i.e. the z-scored descriptor values in Z_train), (ii) an* m *x* q *matrix of values for the output property to be explained, where* q *is the number of columns in the output data matrix, y (here, the m x 1 vector* ***y_train*** *that contains the PAMPA values), and (iii) a number*, `ncomp`, *which is the maximum number of PCs to be used in the model, where N has a value between 1 and n. Here we set* `ncomp` *= 20 to create PLSR models for up to the first 20 PCs*.

*As outputs, the ‘plsregress’ command generates a number of new data variables, of which the most relevant here are as follows:*

*‘Xloadings’: A matrix of size* n *x* ***ncomp*** *containing the loadings (i.e. coefficients) of each of the n (i.e. 127) descriptors in the* ***ncomp*** *PCs chosen in the command*.

*‘Yloadings’: A matrix of size* 1 *x* ***ncomp*** *containing the loadings (i.e. coefficients) of the output variable* ***y_train*** *in the* ***ncomp*** *PCs chosen in the command. Note: if using PLSR with an output data set*, ***y***, *that contains more than a single column of numbers, then ‘Yloadings’ will have size # x* ***ncomp***, *where # is the number of columns in* ***y***.

*‘Xscores’: A matrix of size* m *x* ***ncomp*** *containing the scores (i.e. PC coordinates) of each of the m compounds in the training Set (i.e. 161) for each of the* ***ncomp*** *PCs chosen in the command. Note: the scores in ‘Xscores’ are orthonormalized, meaning that they have been normalized such that each component axis extends from a negative extreme of -1 to a positive extreme of +1*.

*‘Yscores’: A matrix of size* m *x* ***ncomp*** *containing the response scores (as opposed to the input scores) of each of the m compounds in the training Set (i.e. 161) for each of the* ***ncomp*** *PCs chosen in the command. The scores in ‘Yscores’ are not orthogonal or normalized*.

*‘betaPLS’: The regression coefficients for predicting* y *from X using the PLS components. ‘PLSPctVar’: A 2 x* ***ncomp*** *matrix in which the first row contains the percentage of the total variance in X (here*, ***X_train****) that each PC explains, and the second row contains the percentage*

*of the total variance in y (here*, ***y_train****) that each explains. Thus, row 1 indicates how much of the input information from the descriptors in X is captured by a PC, and row 2 indicates how much the PC explains the behavioral output given in y*.

*For more information and explanations about the inputs and outputs for the ‘plsregress’ command, type ‘doc plsregress’ into the Command window*.

##### 13. Plot a graph showing the percent variance explained in y_train versus the number of PCs

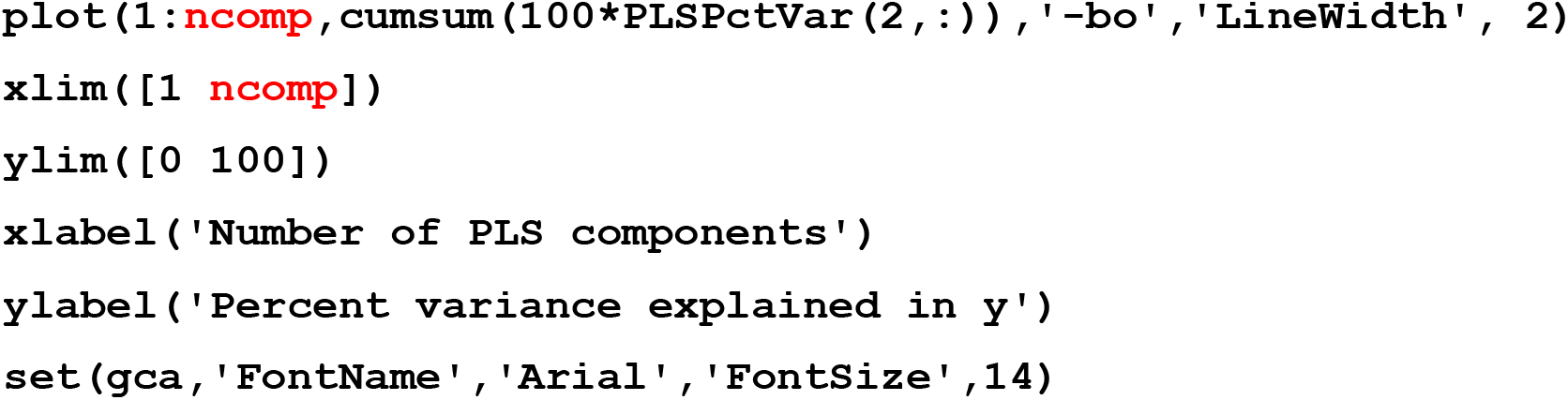

*As in Step 11, the input* ***ncomp*** *is the number of PCs, in this case the number to be shown on the plot. The value of* ***ncomp*** *should be no greater than* ***ncomp*** *from step 11. Here, we use* ***ncomp*** *= 20 to show the results for all 20 PCs for which the calculation was done in Step 11*.

*A new Figure window will open containing a graph resembling* ***Figure 4.2***.

**Figure 4.2.**
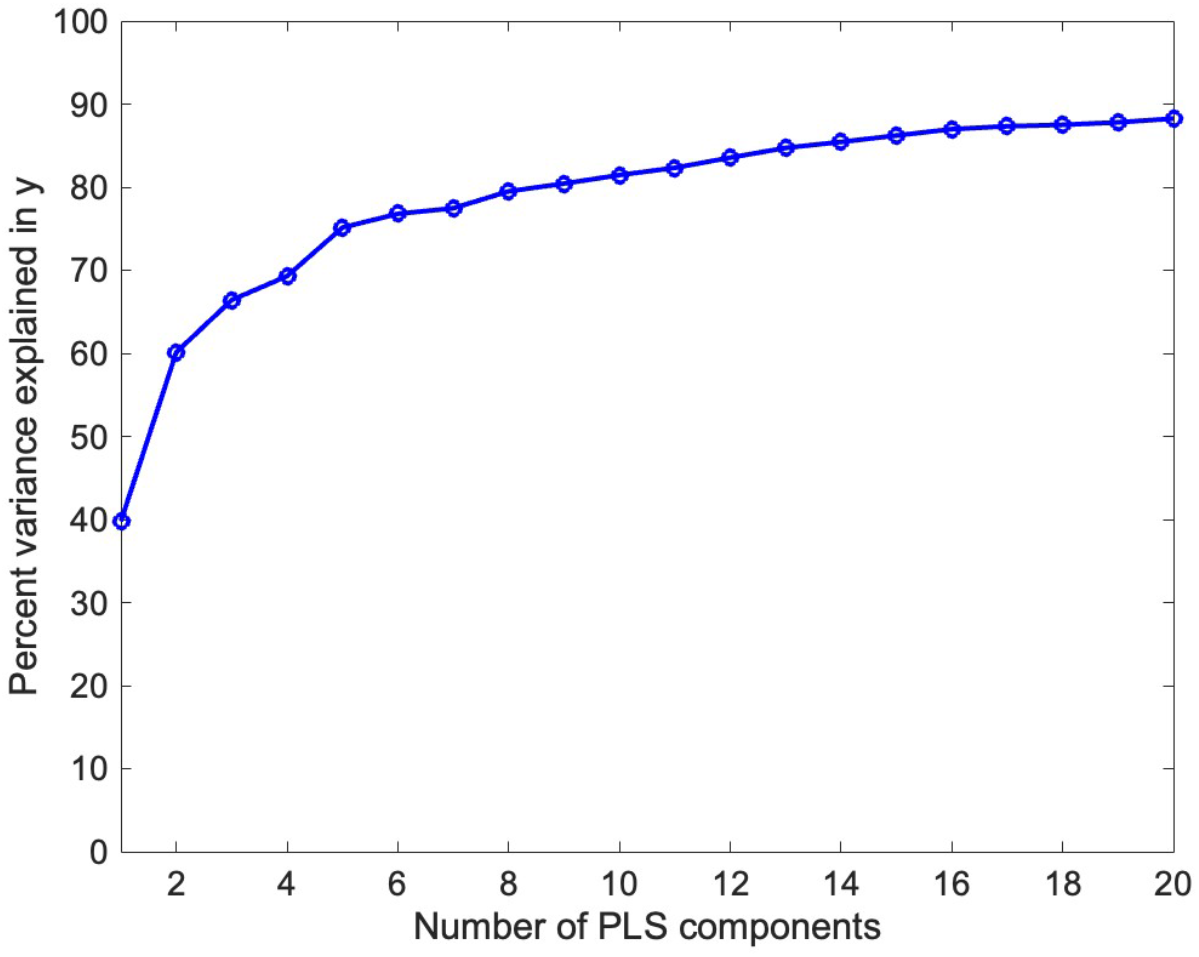
Plot showing how much of the variance in **y_train** is explained by the PLSR models is increasing numbers of PCs are included in the model building. Such a plot will typically asymptote at a value below 100%, because the combined input data in **X_train** may not contain all of the information required to fully explain variations in compound permeability. In this case, the asymptote appears to be close to 90%, implying that the model has been provided with sufficient information, in the form of the descriptor values in **X_train**, to account for most of the variation in the compound’s permeabilities. Indeed, the first 5 PCs alone capture sufficient information to account for ∼75% of the variance in **y_train**.

*The results from Figure **4.2** show that the information provided in X_train can account for the the lion’s share of variations in permeability among the Training Set compounds. If the asymptote was reached at a much lower value, this would indicate that the output behavior of compound permeability cannot be explained by the descriptors provided in X, suggesting that molecular properties important for permeability are absent from the descriptor set.*

*The observation that the first 5 PCs can account for 75% of the variance in* ***y_train*** *does not mean that 5 is the optimal number of PCs to include in the PLSR model. What matters is not how well the model explains the permeabilities of the compounds that were used to train the model, but rather how accurately the model can predict the permeabilities of compounds that it hasn’t seen before. As was the case with PLSDA, in Basic Protocol 3, we will assess the predictive accuracy of the model first by cross-validation and then by applying the model to a Test Set of compounds that were held back from model building*.

##### 14. Assess the predictive accuracy of the PLSR models using different numbers of PCs by cross-validation

*Cross-validation determines the optimal number of PCs to use in model building, to best balance the trade-off between useful information that is relevant to permeability versus noise in the data*.

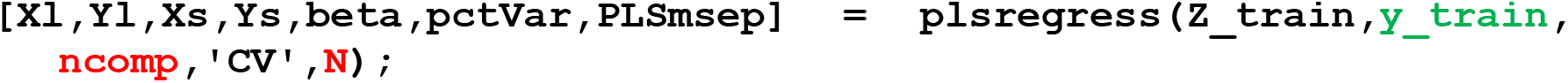

*As in Steps 11 and 12*, ***ncomp*** *should be replaced by the maximum number of PCs you wish to include in the cross-validation calculation. Here we have chosen to test up to 20 PCs, and therefore set* ***ncomp*** *= 20*.

*‘****N****’ has a value from 2 to* m, *where* m *is the number of elements in the set. Here, we set* ***N*** *= 161. Setting ‘****N****’ = 2 will divide the data into two groups and run the modeling twice, leaving out each group in turn and then assessing the accuracy of the model generated using each half of the data set at predicted the permeability class of the other half of the compounds that were omitted. At the other extreme, setting ‘****N****’ =* m *will run the model* m *times, leaving out a single compound each time and measuring how well its permeability class is predicted by the model made using the data for the remaining* m*-1 elements. This latter procedure is known as ‘leave-one-out’ cross-validation. The fewer elements that are omitted in each round of cross-validation, the better the models in each round are likely to be. Thus, setting ‘****N****’ =* m *(i.e. ‘****N****’ = 161, in this case) is a safe choice. However, setting ‘****N****’ <* m *is sometimes advisable to reduce the computation time when working with large data sets*.

*More information on the plsregress command and k-fold cross validation can be found at the following links: https://www.mathworks.com/help/stats/plsregress.html and https://www.mathworks.com/discovery/cross-validation.html.*

##### 15. Plot the cross-validation results

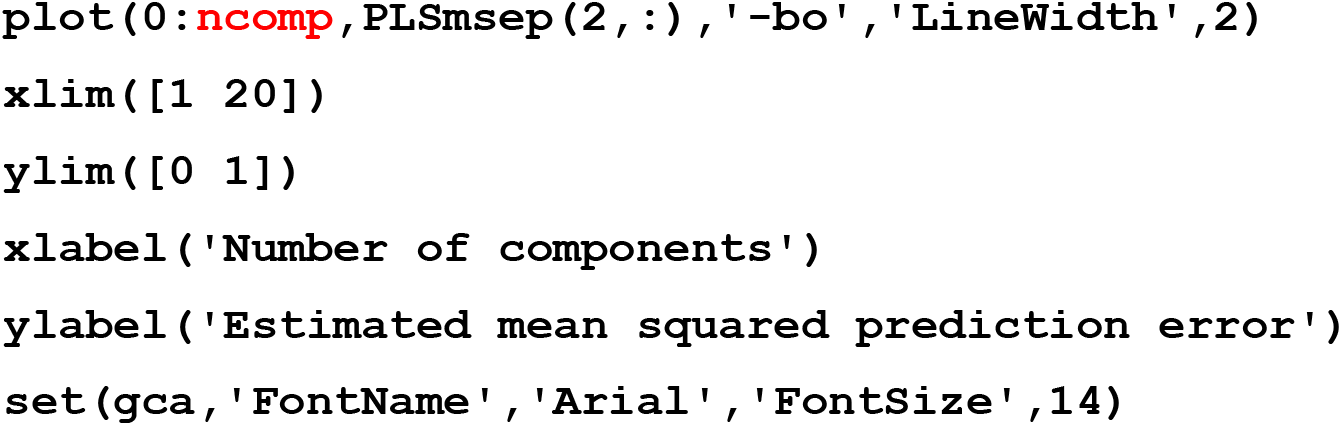

*As in Step 13, the input* ***ncomp*** *is the number of PCs, in this case the number to be shown on the plot. The value of* ***ncomp*** *should be no greater than* ***ncomp*** *from step 13. Here, we use* ***ncomp*** *= 20 to show the results for all 20 PCs for which the calculation was done in Step 13*.

**Figure 4.3.**
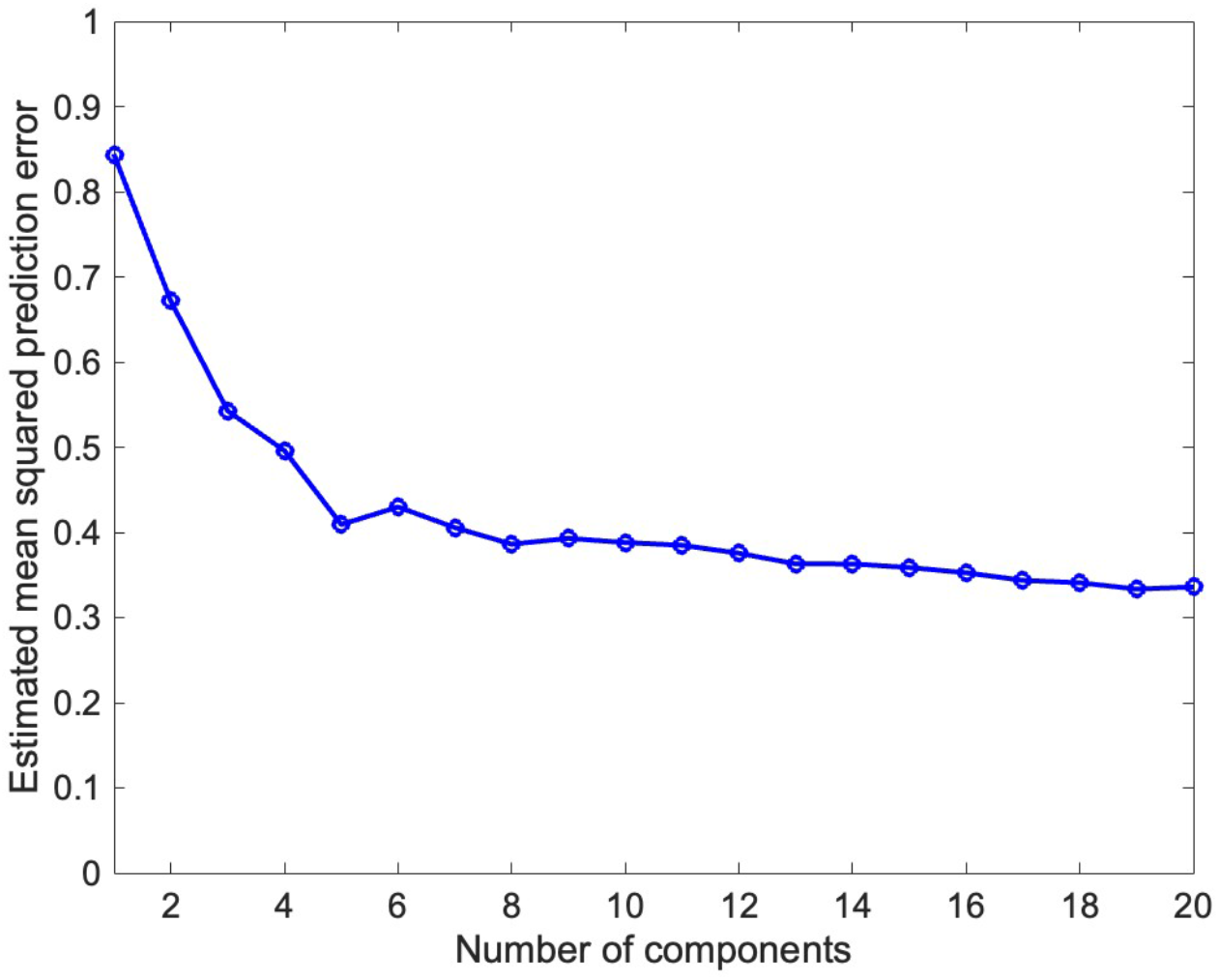
Results from leave-one-out cross-validation for PLSR models generated using the data in the Training Set and including the number of PCs indicated on the x-axis. The model using five PCs gave a local minimum in root-mean-square prediction error, with only incremental decreases in error seen upon including additional PCs in the model building. The results indicate that to develop a model with maximal predictive accuracy and minimal overfitting, the first five PCs should be used.

##### 16. Build a PLSR model for the Training Set using PCs 1-5

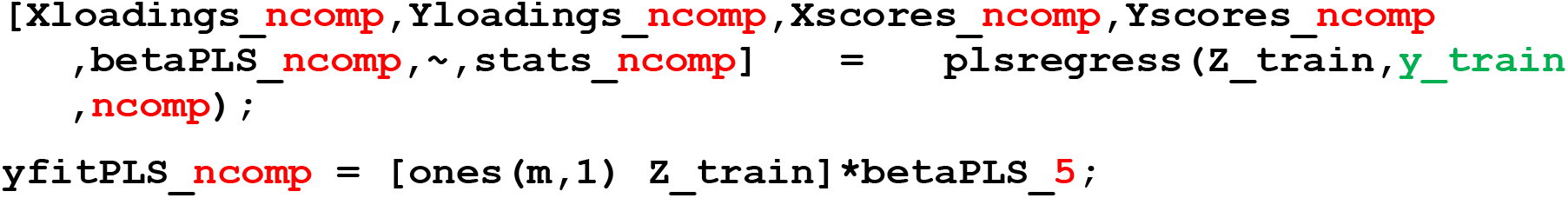

*Because* ***Figure 4.3*** *showed that the increase in predictive power is minimal beyond the first five principal components*,, *we chose* `ncomp` *= 5 to give the simplest model and to minimize overfitting. User judgment is required to decide when it is worth taking additional components into account to increase predictive power*.

*In the second command, be sure to leave a space between* ***ones(n,1)*** *and* ***Z_train***.

*The inputs and outputs for the ‘plsregress’ command are analogous to those described in Step 11*.

##### 17. Compare the modeled versus experimental permeability values for the Training Set compounds

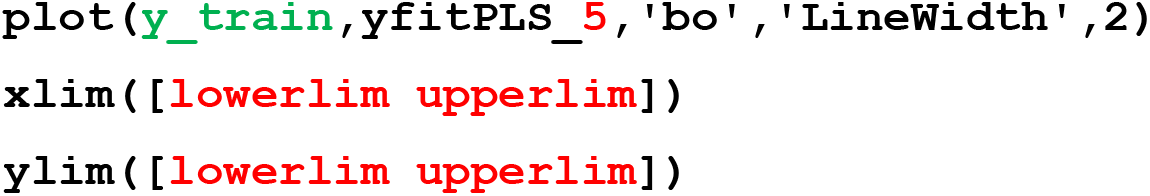
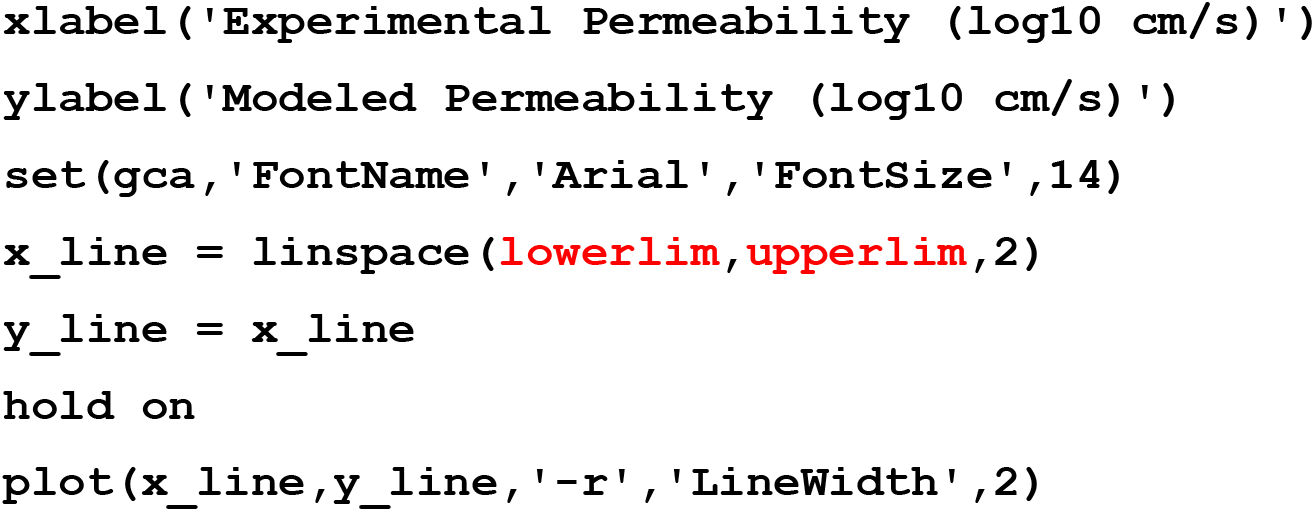

*The red ‘****5****’ in ‘yfitPLS_5’ need not be manually entered, as it was included in the name of this variable when created in Step 15. We highlight it in red here only to indicate that if a different value for* `ncomp` *is chosen in Step 15, then the number in this variable name will be different*.

*The inputs l****owerlim*** *and* ***upperlim*** *are used to set the lower and upper limits to the x- and y-axis scales. These limits should be set at values that encompass all experimental and modeled log Permeability values in* ***y_train*** *and* ***yfitPLS_5***. *Looking at these two data variables led us to choose l****owerlim*** *= -10 and* ***upperlim*** *= -3, which we found to include all the values in either variable*.

A new Figure window will appear that contains a plot resembling that in **Figure 4.4**.

**Figure 4.4.**
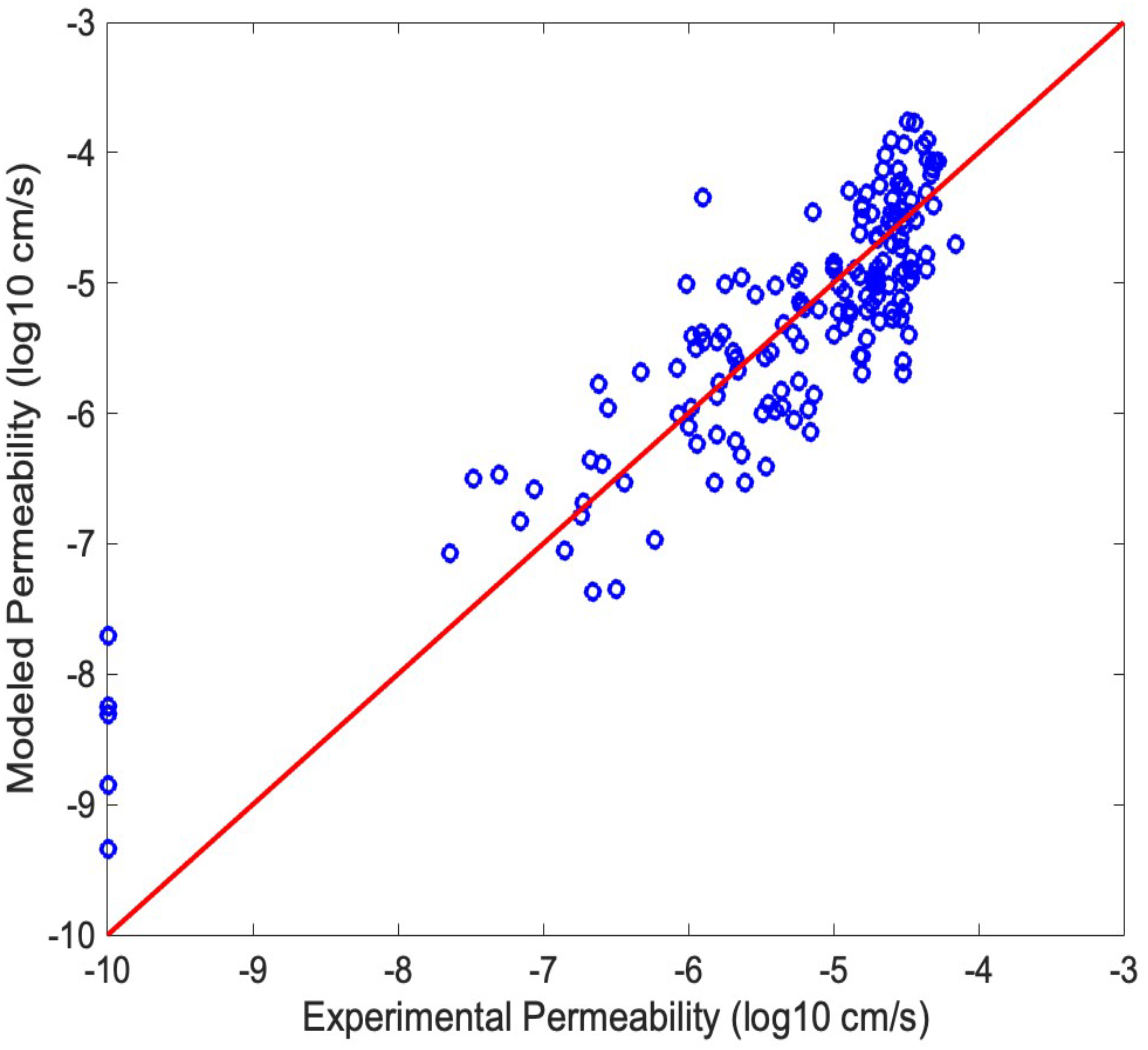
Scatter plot showing experimental versus modeled permeability values for the 161 compounds in the Training Set (blue circles). The x-axis shows the experimentally determined PAMPA value (from **y_train**), and the y-axis shows the modeled PAMPA values (from ‘yfitPLS_5’) based on the 5 PC PLSR model created in Step 15. The red line is the line of identity, y = x. The closer a data point lies to the line, the more accurately its permeability was modeled.

*Visual inspection of* ***Figure 4.4*** *shows a degree of correlation between the modeled and the experimental permeability values, implying that the PLSR model from Step 15 was somewhat effective in accounting for the permeability of the compounds in the Training Set. The predictive accuracy of the model can be quantified by calculating the Pearson Correlation Coefficient (R*^*2*^*) value between the measured and modeled permeability values. We do this in the next step*.

##### 18. Calculate r2 for the modeled versus the measured permeability values for the Training Set

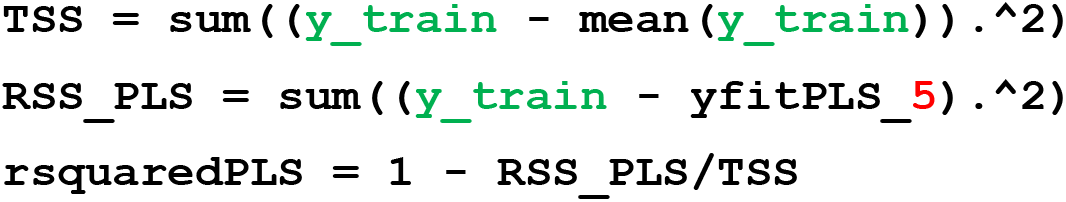

*The red ‘****5****’ in ‘yfitPLS_5’ need not be manually entered, as it was included in the name of this variable when created in Step 15. We highlight it in red here only to indicate that if a different value for* `ncomp` *is chosen in Step 15, then the number in this variable name will be different*.

*More information on the regression calculation can be found at the following link: https://www.mathworks.com/help/stats/partial-least-squares-regression-and-principal-components-regression.html*

The above command returns a value for ‘rsquaredPLS’ = 0.7514, indicating that the R^2^ value for the data fit in Figure 4.4 = 0.7514.

*The observant reader will note that this R*^*2*^ *value is essentially equivalent to the percent y-variance explained by PCs 1-5 in* ***Figure 4.2***. *Both numbers express what fraction of the variation in* ***y_train*** *can be explained by the information contained in the first five PCs calculated by PLSR*.

##### 19. Generate a biplot of the Training Set scores using the first two PCs

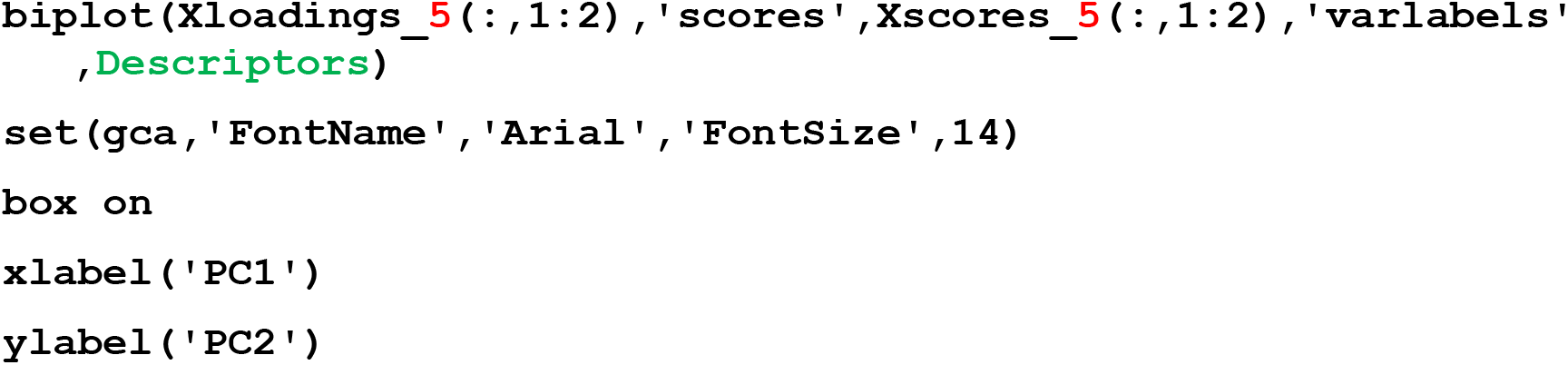

*The red ‘****5****’ in ‘Xloadings_5’ and in ‘Xscores_****5****’ need not be manually entered, as it was included in the name of this variable when created in Step 15. We highlight it in red here only to indicate that if a different value for* `ncomp` *is chosen in Step 15, then the number in this variable name will be different*.

**Figure 4.5.**
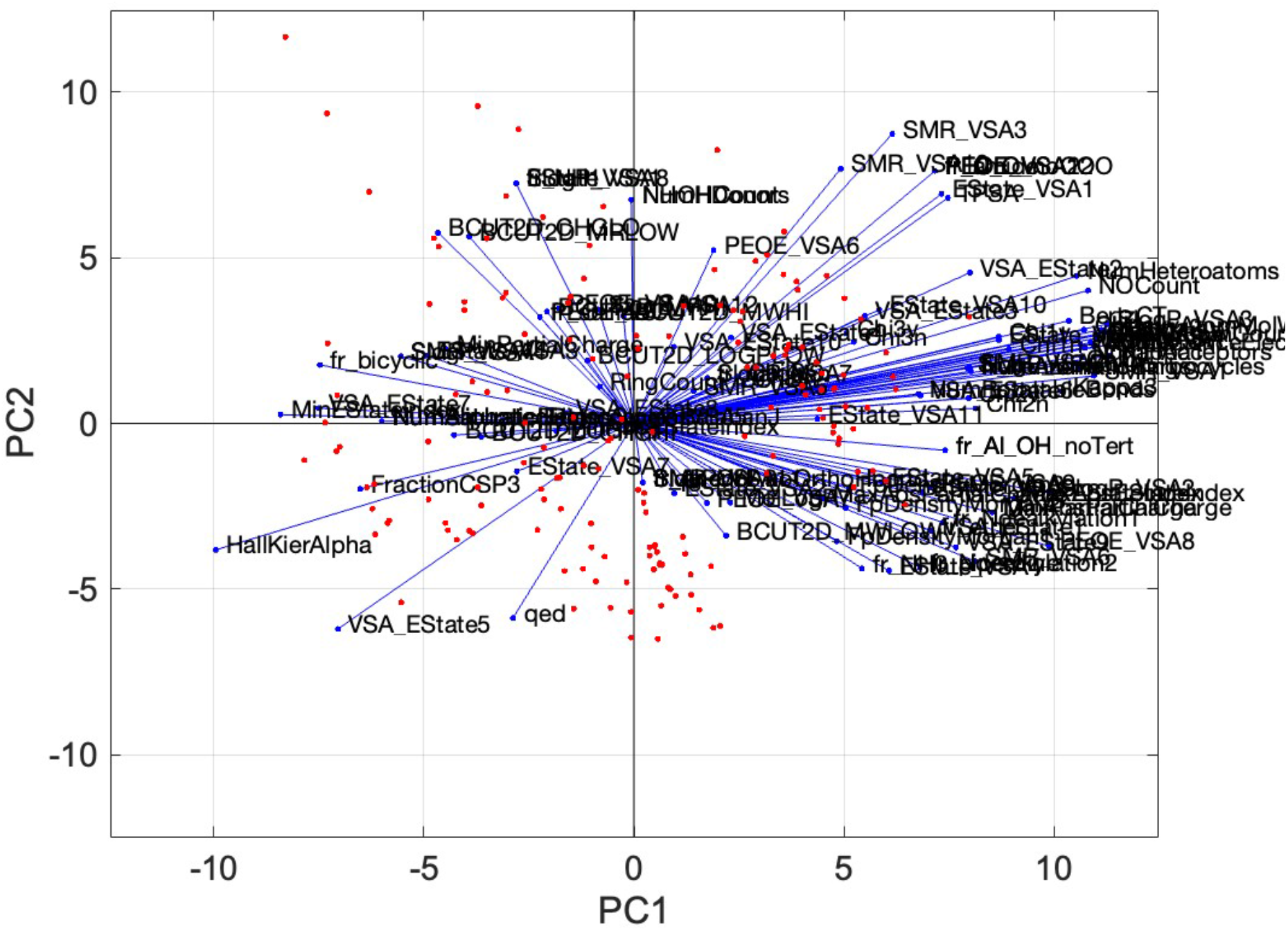
Biplot showing the 161 compound scores and 127 descriptor coefficients (loadings) from the PLSR model (5 PCs) developed using the Training Set, plotted with respect to PC1 and PC2. Each red datapoint represents the position in PC space of a compound with respect to PCs 1 and 2. The blue lines, superimposed on the same axes, indicate the degree to which each descriptor contributes to PC1 and PC2. Specifically, the blue lines extend from the origin to a point in PC space that has a coordinate in PC1 that corresponds to the orthonormalized value of the descriptor’s coefficient in PC1, and a coordinate in PC2 corresponding to its coefficient in PC2.

*The descriptors that most influence a compound’s score in PC1 are those with the highest (absolute value) coefficient in PC1, identified by the blue lines with terminal points closest to the left and right edges of the plot. The descriptors that contribute most to PC2 are indicated by the blue lines that terminate closest to the top and bottom edges of the plot*. *<u>Important: when plotting a biplot, MATLAB sometimes inverts the signs of the loading (coefficient) values for some components so that the largest absolute Loading value is assigned as positive. Therefore, to know the sign of a coefficient it is necessary to look at the data variable ‘P’ rather than reading off the biplot. (For additional discussion of biplot</u>, see Basic Protocol 2, Step 14.)*

*The plot in Figure 4.5 is so crowded with descriptors and their labels that its interpretability and visual impact are poor. In the following steps, we describe how to generate more useful biplots that show only a selected subset of descriptors*.

##### 20. Create a biplot showing only the M descriptors having the highest loadings (absolute value) in PC1

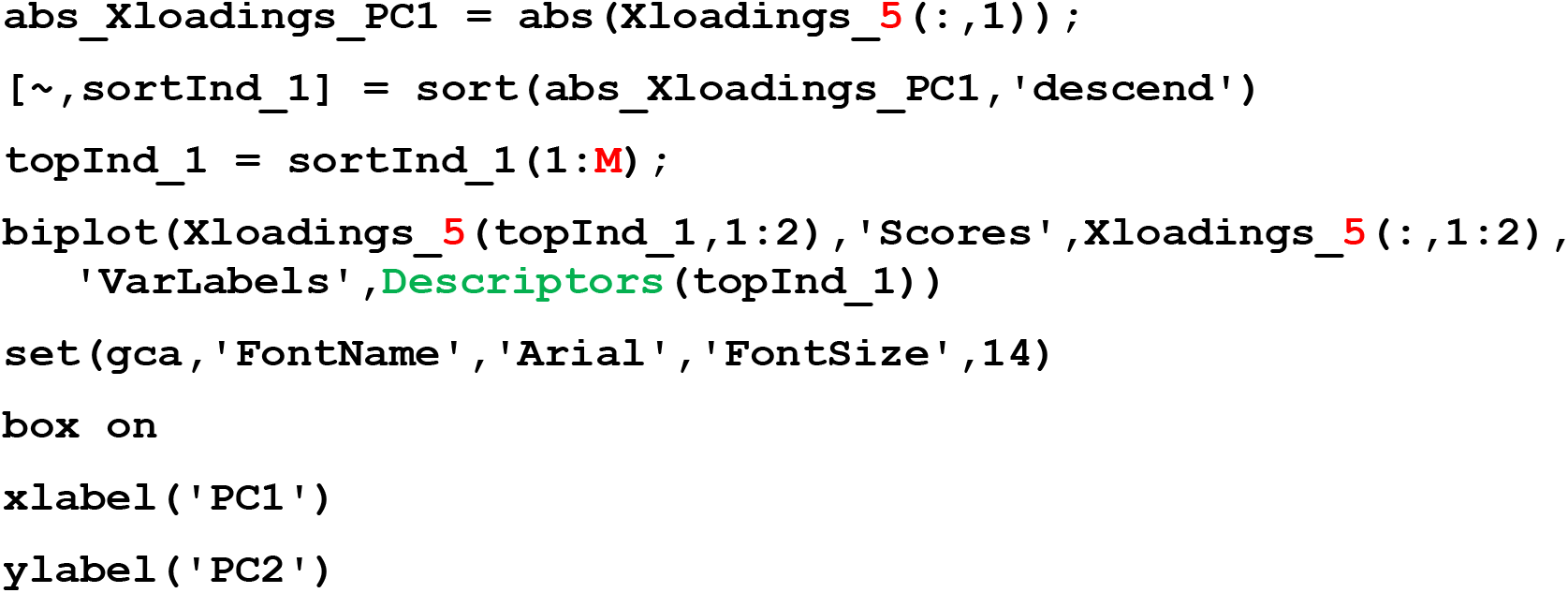

*The red ‘****5****’ in ‘Xloadings_5’ need not be manually entered, as it was included in the name of this variable when created in Step 15. We highlight it in red here only to indicate that if a different value for* `ncomp` *is chosen in Step 15, then the number in this variable name will be different*.

*In the plot shown in* ***Figure 4.6***, *we set* ***M*** *= 30 to display the 30 descriptors that most influence a compound’s score with respect to PC1*.

**Figure 4.6.**
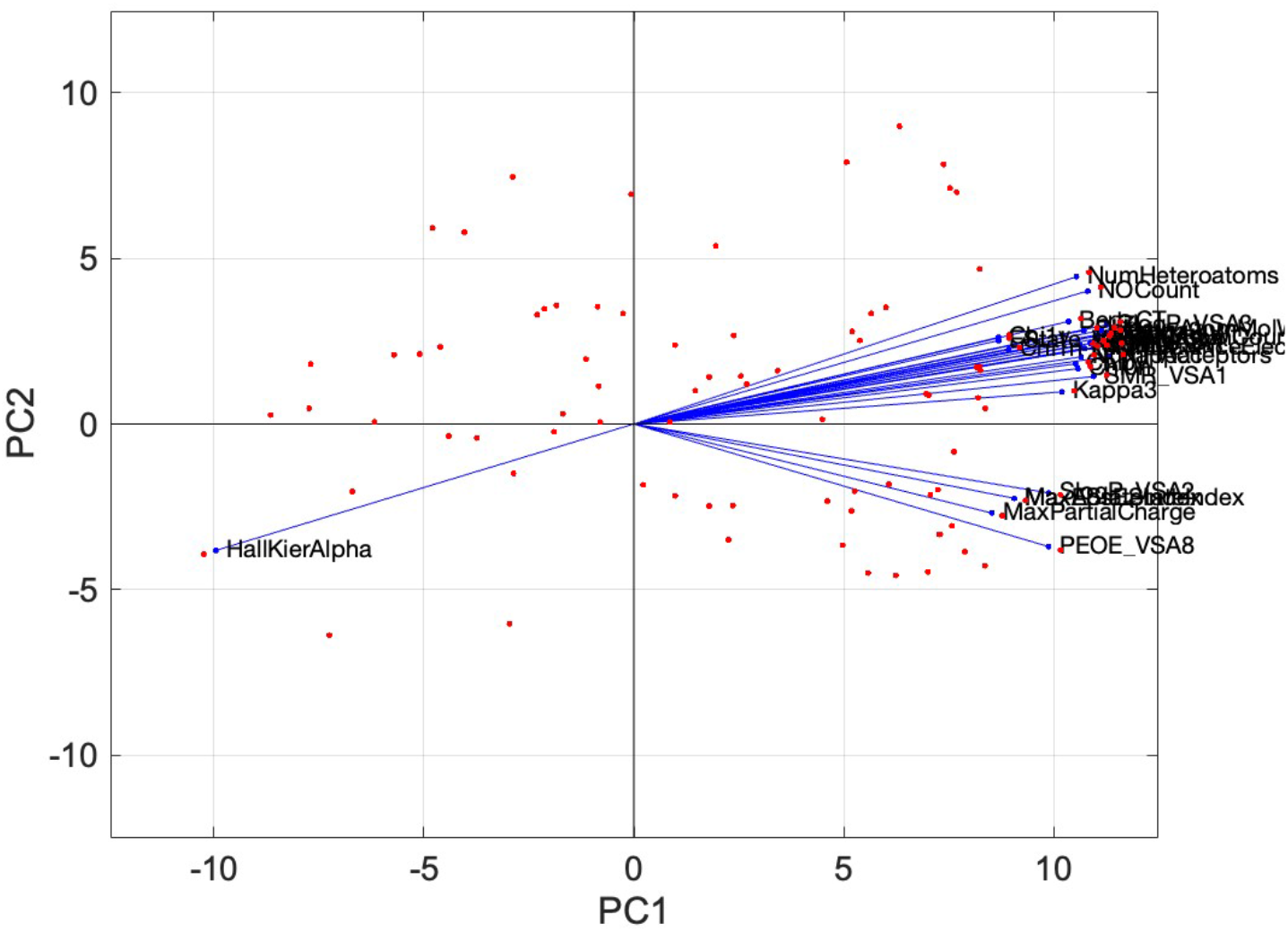
Biplot of the top 30 descriptors with the highest absolute loadings on PC1, as calculated by the PLSR model from Step 15, indicating the most significant contributors to PC1 for distinguishing between the permeable and impermeable compounds.

##### 21. Create a biplot showing only the M descriptors having the highest loadings (absolute value) in PC2

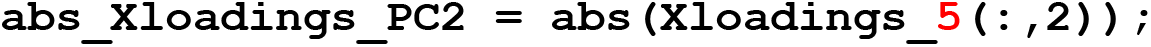
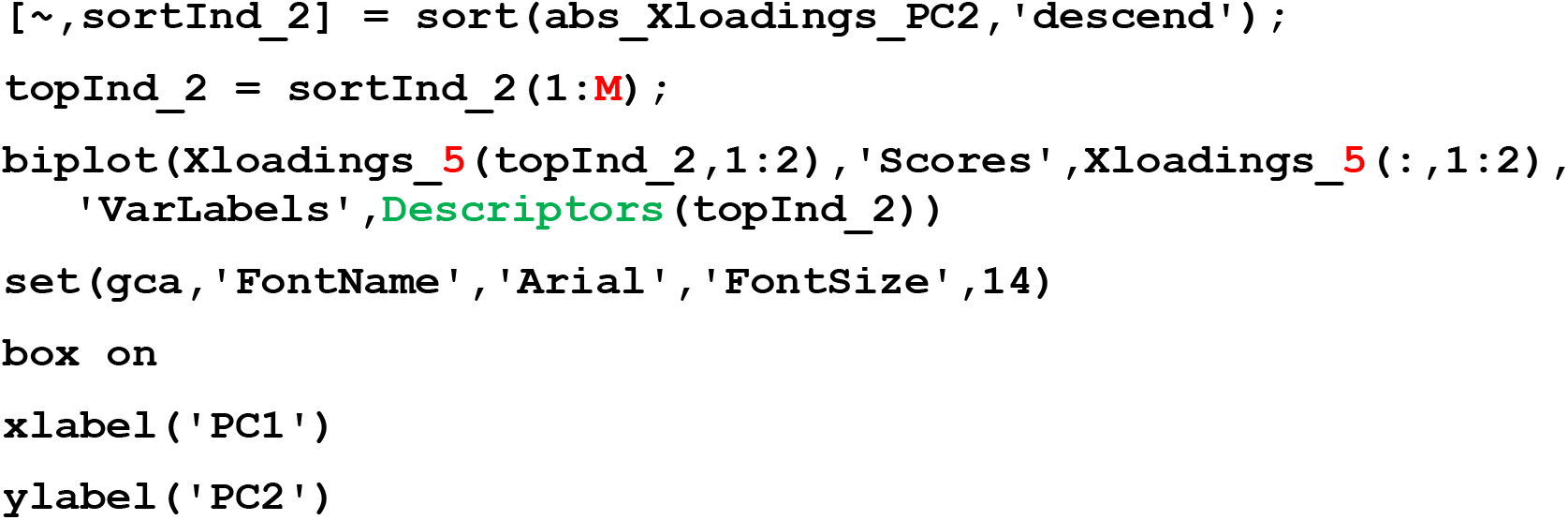

*The red ‘****5****’ in ‘Xloadings_5’ need not be manually entered, as it was included in the name of this variable when created in Step 15. We highlight it in red here only to indicate that if a different value for* `ncomp` *is chosen in Step 15, then the number in this variable name will be different*.

*As in the previous step, in the plot shown in* ***Figure 4.7***, *we set* ***M*** *= 30 to display the 30 descriptors that most influence a compound’s score with respect to PC2*.

**Figure 4.7.**
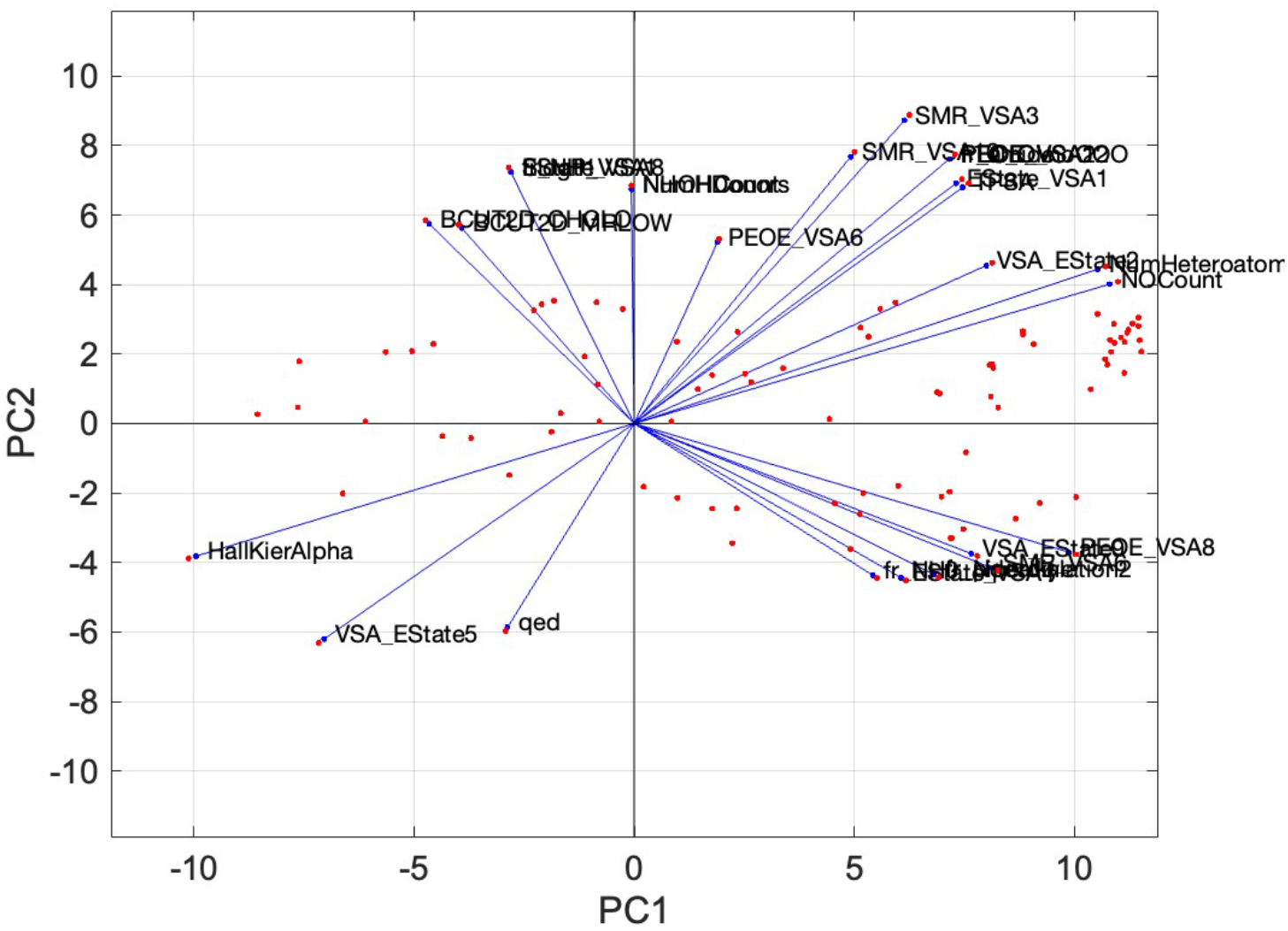
Biplot of the top 30 descriptors with the highest absolute loadings on PC2, as calculated by the PLSR model from Step 15, indicating the most significant contributors to PC2 for distinguishing between the permeable and impermeable compounds.

##### 22. Create a table of the loadings with the highest contribution to each PC

to understand how much each key descriptor contributes to each PC.

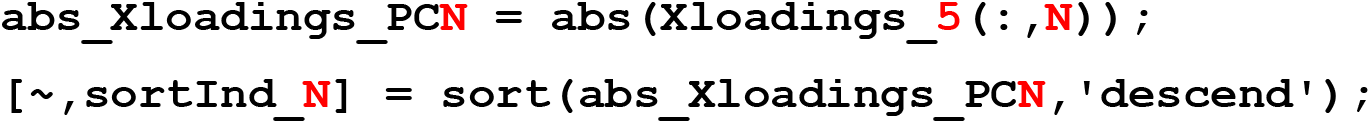
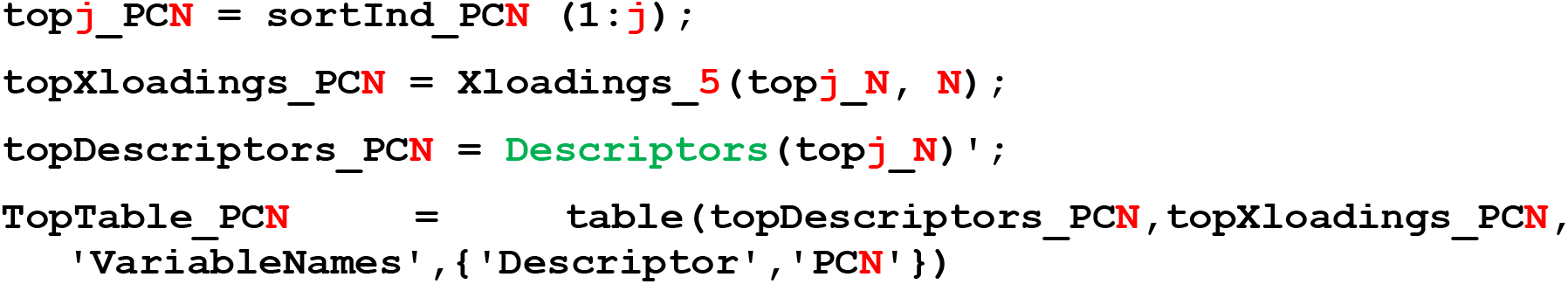

*Note that the red ‘****5****’ in ‘Xloadings_****5****’ need not be manually entered, as it was included in the name of this variable when created in Step 15. We highlight it in red here only to indicate that if a different value for* `ncomp` *is chosen in Step 15, then the number in this variable name will be different*.

*This command creates a table of the* ***j*** *descriptors with the highest coefficients in the specified PC (i.e. PC* ***N****). The command must be run separately for each PC of interest*.

*Line 1 of this command creates a new data variable of size* n *x 1 (i.e. 127 descriptors x 1 selected PC) which contains the absolute values of the coefficients for the descriptors in PC number* ***N***.

*Line 2 sorts the coefficients from ‘abs_Xloadings_PC****N****’ in descending order, and creates a new variable called ‘sortInd_PC****N****’ with the same size as ‘abs_Xloadings_PC****N****’. This new variable contains the row numbers from ‘abs_Xloadings_PC****N****’ that correspond to the newly sorted order. For example, if the coefficient with the largest absolute value in PC****N*** *was the number in row 57 of ‘abs_Xloadings_PC****ncomp****’, then the number ‘57’ would be in the top cell of ‘sortInd_PC****N****’*.

*Line 3 creates a new variable, top****j****_PC****N***, *that contains just the top* ***j*** *rows of ‘sortInd_PC****N****’. To tabulate just the 10 most important descriptors in each PC, here we set* ***j*** *= 10*.

*Line 4 uses the sorted row numbers that comprise the entries in ‘sortInd_PC****N****’ to identify the coefficients with the top j values in PC****N*** *from ‘Xloadings_****5****’ and copy them, in descending order, into a new data variable, ‘topXLoadings_PC****N****’. Note, because these values are pulled from ‘Xloadings_****5****’ and not from ‘abs_Xloadings_PC****5****’, each coefficient will have its original sign*.

*Line 5 uses the sorted row numbers that comprise the entries in ‘sortInd_PC****N****’ to identify the names of the coefficients with the top* ***j*** *values in ‘****Descriptors****’, imported in Step 5, and copy them, in the order matching the descriptor values in ‘topXLoadings_PC****N****’, into a new data variable, ‘topDescriptors_PC****N****’*.

*Line 6 creates a new variable of size* ***j*** *x 2, called ‘TopTable_PC****N****’, containing in the first column the names of the descriptors with the* ***j*** *highest (absolute) values in PC****N***, *and in the second column the numerical value of each descriptor, including its original sign*.

***Table 4.1*** *shows the results of running the above command five times, for each of PCs 1-5, and then combining the resulting data variables, ‘TopTable_PC****1****’, ‘TopTable_PC****2****’, ‘TopTable_PC****3****’, ‘TopTable_PC****4****’, and ‘TopTable_PC****5****’ into a single table*.

**Table 4.1.**
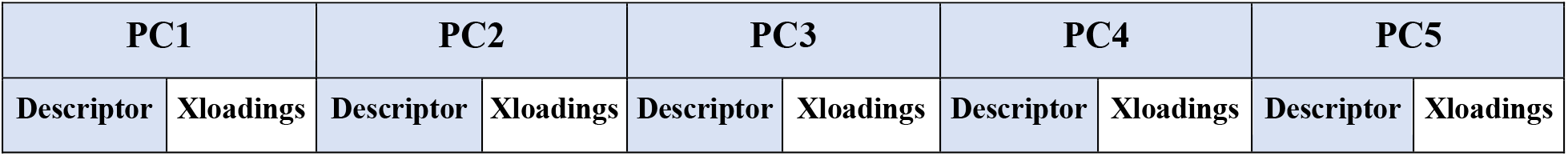
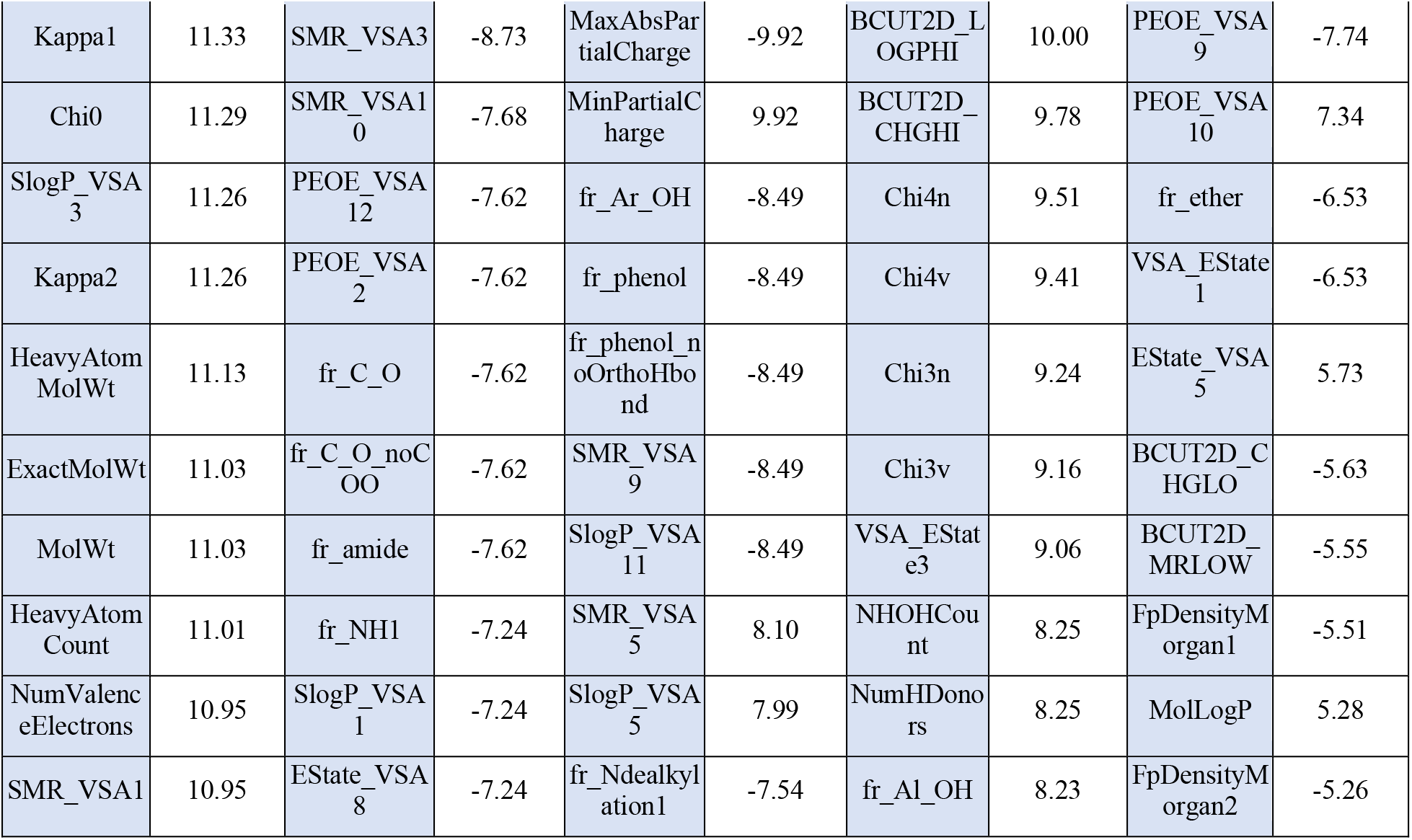
Names and values of the ten coefficients with the highest (absolute) values in each of PCs 1-5, for the PLSR model created in Step 15.

##### 23. Test the predictive accuracy of the PLSR model created in Step 15 by using it to predict the permeabilities of the compounds in the Test Set

*The 40 compounds in the Test Set, created in Step 6, have known experimental permeability values (contained in* ***y_test***, *from Step 6), but at no point were they used in building the PLSR model in Step 16. The ability of the model to use the descriptor values for these compounds (from* ***X_test****) to accurately predict their permeability values is therefore a stringent test of the predictive power of the model, at least with respect to the chemotypes present in the original compound set from which the Training and Test Sets were taken*.

(a) Scale and center the data in **X_test** to match the scaling done with the Training Set in Step 8.

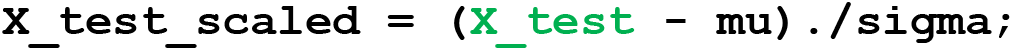

*In MATLAB, placing a period immediately before a mathematical operation, as here in ‘./’, instructs that the operation be performed separately on each value in the matrix. If the period is omitted, the command would instead divide one matrix by the other via matrix division, which gives a very different result*.

###### Important

*When preparing input data for testing in an already-existing model, it is important to scale using the same mean (‘mu’) and standard deviation (‘sigma’) values that were used to scale the Training Set data when the model was originally developed. This is because the scaled descriptor values for the new data points must use the identical scale of the original data used during model building. If this is not done, then compounds from the Test and Training Sets that had, for example, identical unscaled value for a given descriptor would no longer have the same value once the descriptor was scaled. Hence, rather than simply z-scoring the data here, as we did when building the model in Step 8, we instead manually center and scale the data by subtracting from each descriptor value in **X_test** the mean descriptor value from the original scaling in Step 8 (‘mu’), and then divide by the standard deviation for that descriptor from Step 8 (‘sigma’)*.

(b) Replace any NaN values with zeroes, in case NaNs were introduced during the scaling step

~~~
X_test_scaled(isnan(X_test_scaled)) = 0;
~~~

*NaN entries result from invalid mathematical operations such as division by zero, and their presence can prevent the proper operation of some subsequent commands. If you are using the data sets provided with this article, no NaN values will be generated. However, we include this step in case you are following this protocol with your own data set, for which NaN results could occur*.

(c) Apply the PLSR model to the Test Set data to predict the permeability of these compounds

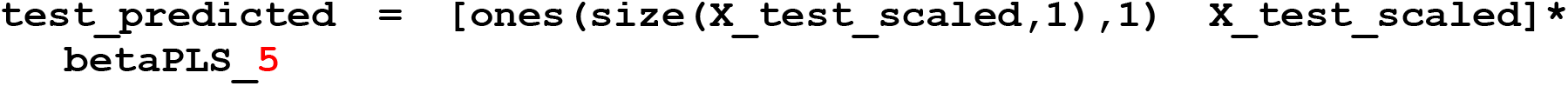

*The red ‘****5****’ in ‘betaPLS_****5****’ need not be manually entered, as it was included in the name of this variable when created in Step 15. We highlight it in red here only to indicate that if a different value for* `ncomp` *is chosen in Step 15, then the number in this variable name will be different*.

*This command creates a matrix of size* m *x 1 (i.e. 40 compounds in the Test Set = 40 rows x 1 column) called ‘test_predicted’, and populates it with the corresponding permeability values calculated using the scaled Test Set descriptor values in ‘X_test_scaled’ and the data from the PLSR model contained in ‘betaPLS_5’ created in Step 15 using the Training Set of compound data*.

*The resulting predicted permeabilities for the Test Set compounds can be visualized by creating a table that contains the predicted values from ‘test_predicted’ alongside the compound names/ID numbers from* ***Compounds_test*** *and the experimental permeabilities of the Test Set compounds from* ***y_test***, *both of which were imported in Step 6*.

**Table 4.2.**
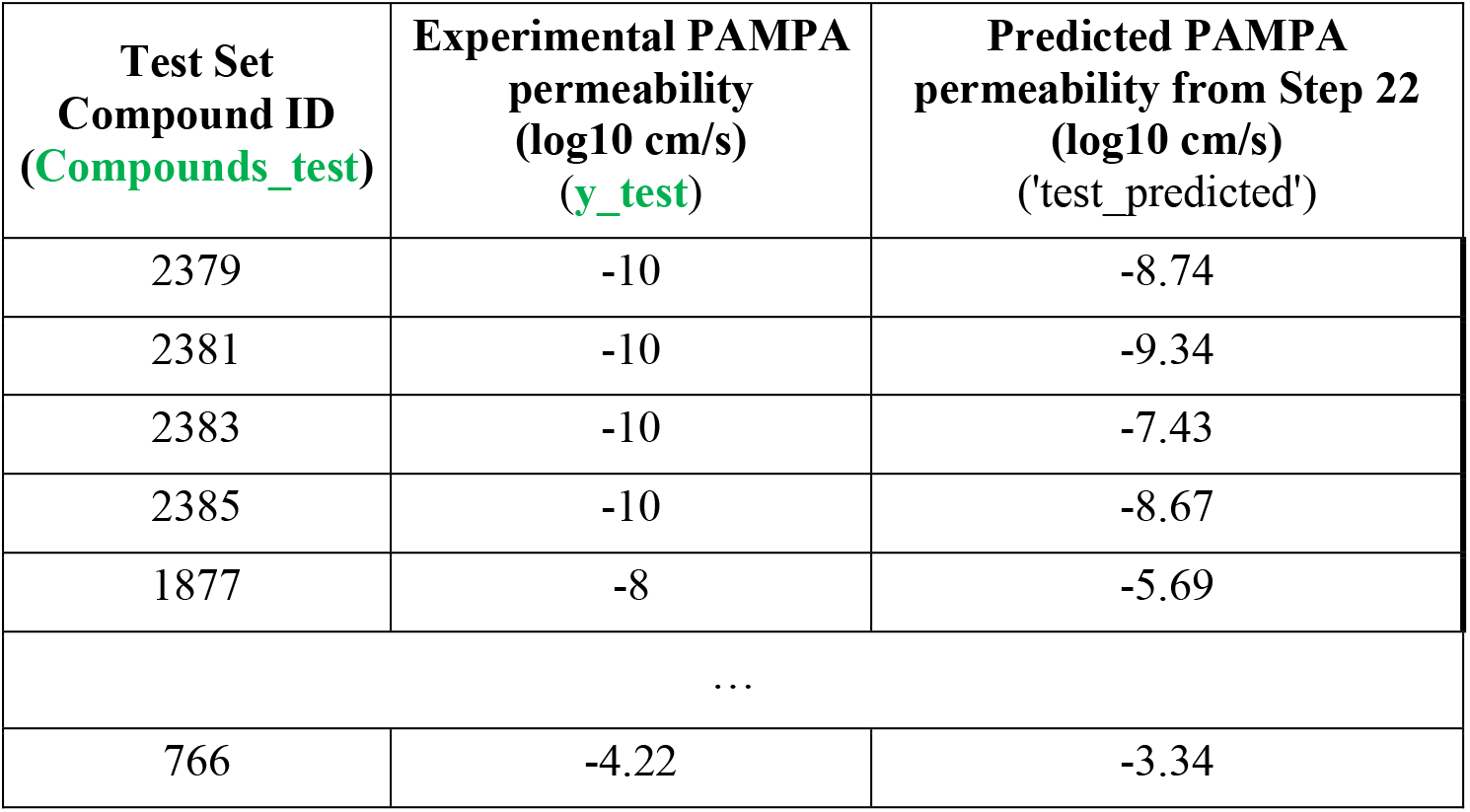
Predicted permeability values for the first few compounds from the Test Set, compared to their experimentally determined permeabilities. Each column in the table contains the data from a different MATLAB workspace variable, as indicated in the column headers. The full table, for all 40 Test Set compounds, can be found in the Data_PLSR.xlsx file provided with this article, under the PLSR_result tab.

##### 24. Plot the observed versus predicted response for the Test Set

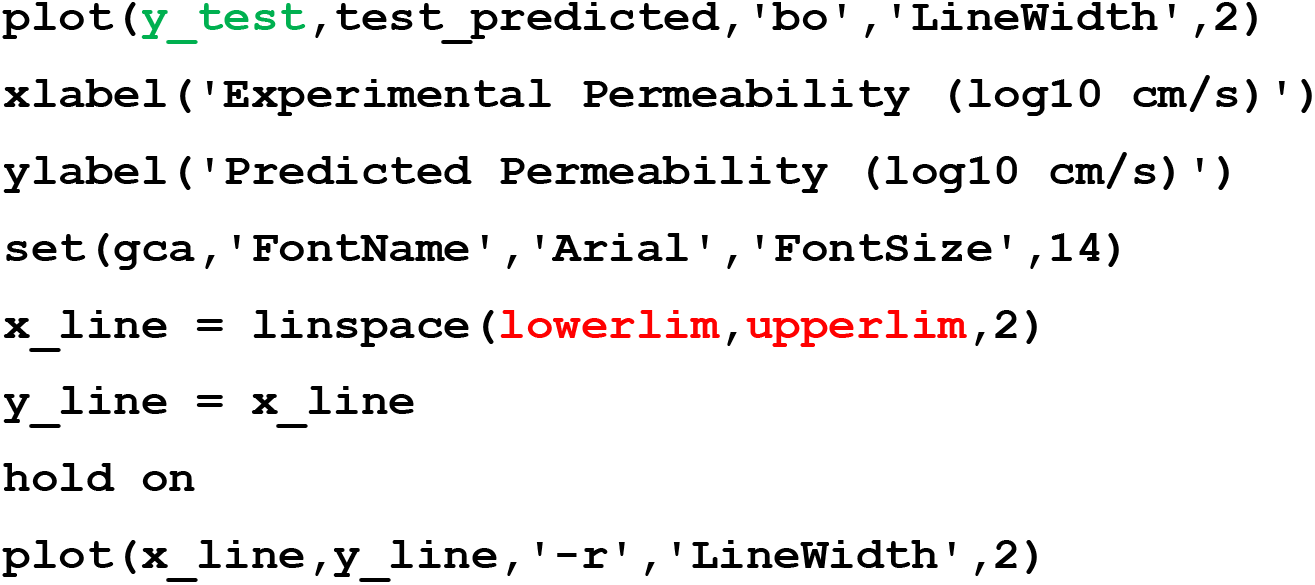

*The first line of this command creates a scatter plot of the data in* ***y_test*** *as x-variable and the data in ‘test_predicted’ as y-variable, and formats the appearance of the data points. Lines 2-4 add and format the axis labels. Lines 5 and 6 create a set of calculated data points corresponding to the line of identity, y = x. The inputs* ***lowerlim*** *= -10 and* ***upperlim*** *= -3 for this reference line were selected based on the range of permeability values found in* ***y_test***. *The line 7 command ‘hold on’ instructs that the following features are to be plotted on the existing plot, rather than on a new set of axes. Line 8 plots and formats the reference line of identity*.

*The resulting plot is shown in* ***Figure 4.8***.

**Figure 4.8.**
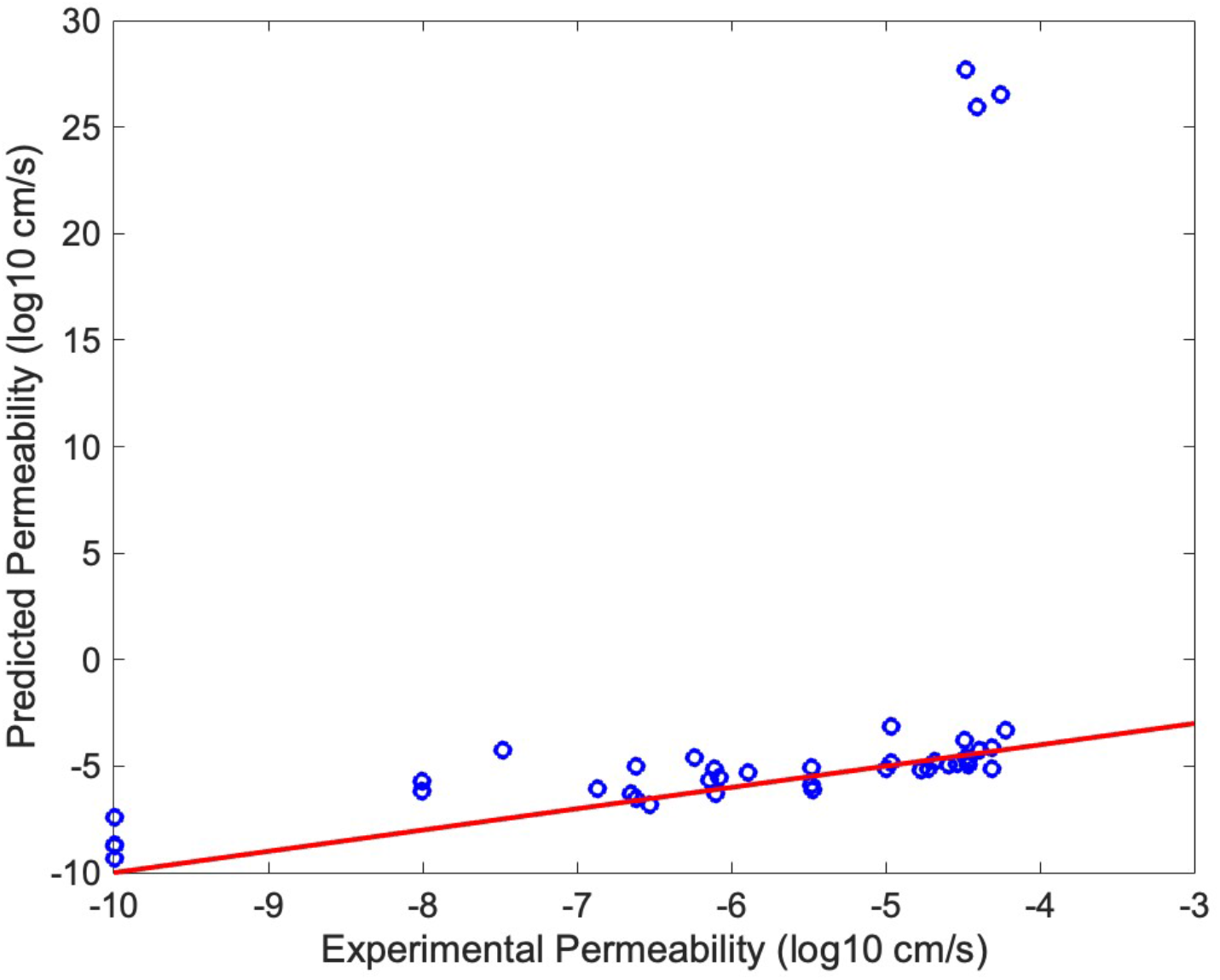
Scatter plot showing experimental versus predicted permeability values for the 40 compounds in the Test Set (blue circles). The x-axis shows the experimentally determined PAMPA value (from **y_test**), and the y-axis shows the predicted PAMPA values (from ‘test_predicted’) based on the 5 PC PLSR model created in Step 15. The red line is the line of identity, y = x. The closer a data point lies to the line, the more accurately its permeability was predicted.

***Figure 4.8*** *shows that the model was able to predict the permeability of some compounds in the Test Set with reasonable accuracy, but there are three egregious outliers for which the predicted permeability values are well above any reasonable range, and several other compounds for which permeability appears to be significantly overestimated*.

*A closer look at the 37 non-outlier compounds can be achieved by rescaling the y-axis of the plot to a scale of -10 to -3. This can be done by entering the following command, setting =* ***lowerlim*** *-10 and* ***upperlim*** *= -3:*

***ylim([lowerlim upperlim])***

*The resulting plot*, ***Figure 4.9***, *shows that, the three outlier compounds (no longer visible) aside, permeability is predicted reasonable accurately (± 1 log unit) for ∼75% of the compounds in the Test Set, but there is a general tendency for the model to overestimate the permeability of the remaining quarter of the compounds*.

**Figure 4.9.**
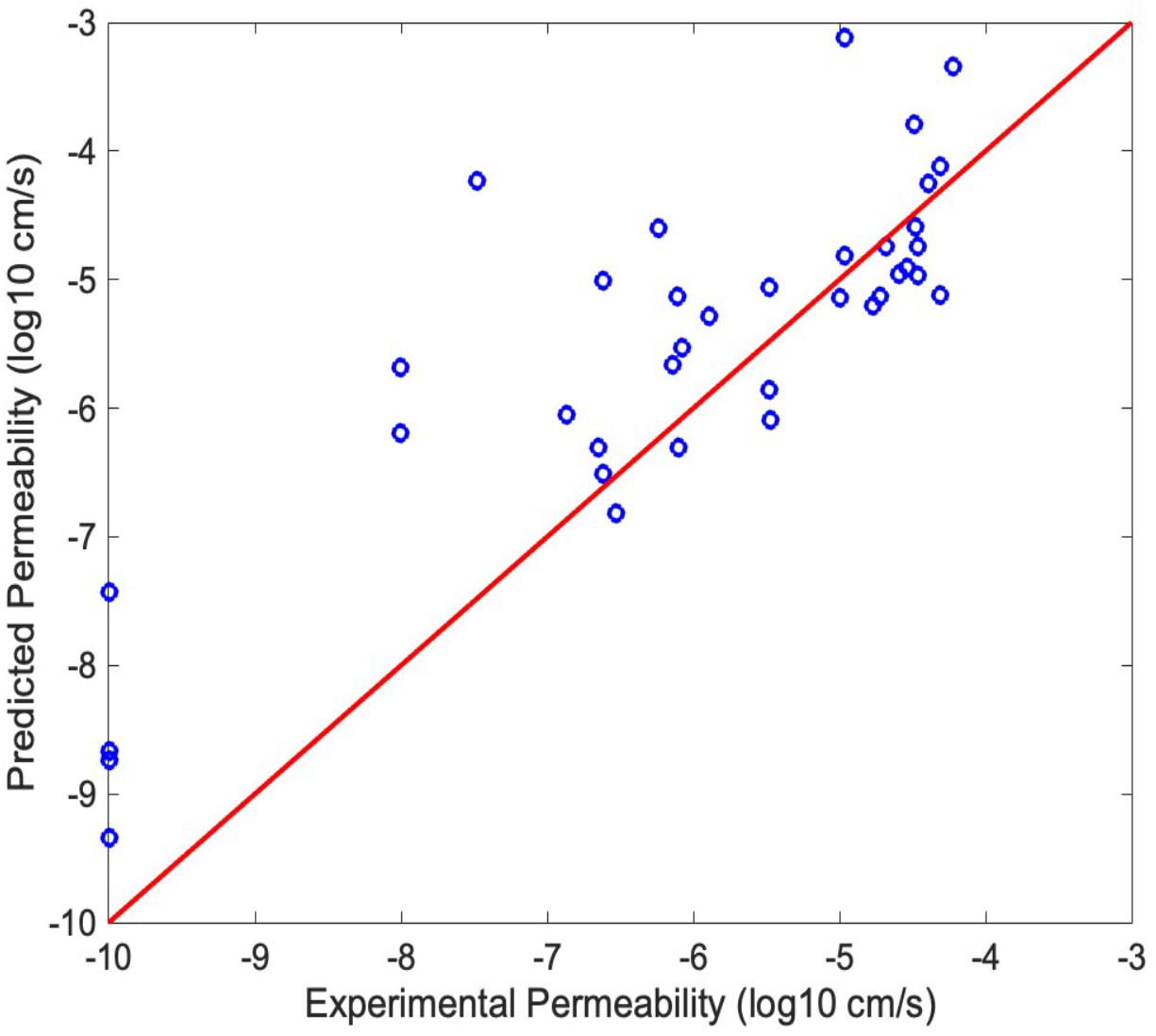
The data from Figure 4.8, but with the y-axis scale modified to extend from - 10 to -3. The blue circles show the experimental versus predicted permeability values for the 37 compounds in the Test Set that remain on-scale (the three high outliers seen in Figure 4.8 are no longer visible, being now off-scale). The red line is the line of identity, y = x. The closer a data point lies to the line, the more accurately its permeability was predicted.

##### 25. Calculate the predictive accuracy of the model

*The accuracy with which the model predicted the permeabilities of the compounds in the Test Set is conventionally quantified in terms of the Pearson Correlation Coefficient (PCC) between the predicted and experimental values. When applied to a Test Set, the PCC is often called a ‘q*^*2*^*’ value, to distinguish it from the ‘r*^*2*^*’ value calculated for the model’s ability to account for the permeabilities of the compounds from the Training Set with which the model was built*.

*The q*^*2*^ *value for the data in* ***Figure 4.8*** *can be calculated as follows:*

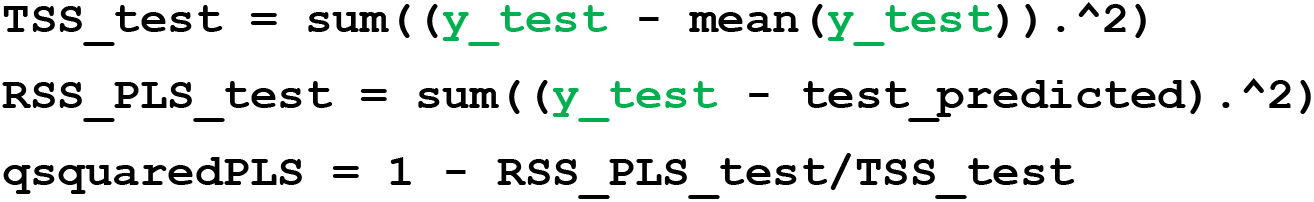

> *For the data shown in* ***Figure 4.8***, *the above command returns the following value:*
>
> *qsquaredPLS = 0.1811*
>
> *indicating that, across the 40 compounds in the Test Set, the model was able to explain ∼18 % of the variations in permeability*.
>
> *The q2 value is greatly influenced by the three extreme outliers visible in* ***Figure 4.8***. *Outliers of this magnitude are not reflective simply of poor predictions, but rather indicate a complete breakdown of the model; the model is simply not equipped to make a meaningful estimate of the permeabilities of these three compounds, for reasons that are discussed in the following section. A different view of the accuracy of the model could be obtained be calculating the q*^*2*^ *value after excluding these three outlier compounds. The resulting value of q*^*2*^ *= 0.98 gives a more accurate picture of the accuracy with which the model can predict the PAMPA permeabilities for compounds that resemble the majority of those used in the Training and Test Sets, but with the understanding that the predictions obtained for other kinds of compounds that differ from these in so-far undefined ways could be egregiously poor*.

##### 26. Save your final workspace as a .mat file

*This file can be reopened to give a MATLAB workspace identical to the state that was saved*.

### Understanding the Results of Basic Protocol 4

The techniques of PLSR is closely related to PLSDA, performing a similar calculation in essentially the same way. To avoid undue repetition, therefore, we refer the reader to the section on Understanding the Results of Basic Protocol 3 for a discussion of what these method do, how they work, interpretation of scores and loadings, and their limitations. Below, we make a few additional points that are specific to PLSR.

#### Percent variance explained

Among the outputs of PLSR are the percent of the variance in the input data matrix X, and in the outcome data matrix *y*, explained by each PC. These are contained in the data variable we called ‘PLSPctVar’ (see Step 11) and, in part, were the data used to plot **Figure 4.2**. For a supervised ML method such as PLSR, the goal is to use the data in X to explain (i.e. account for) the observed variations in the value of *y* among the elements in the data set. The more PCs are used in model building, the more variance in *y* is explained. There is an asymptote, however, and the maximum extent to which the model can explain *y* depends on whether the input data contains all the information required to do so. Thus, the extent to which the asymptote of ‘% *y* explained’ versus number of PCs falls short of 100% is a measure how much of the information required to explain *y* is in fact missing from X, perhaps due to the exclusion of key properties. This asymptote is also affected by the accuracy of the experimental values in *y*; even a perfect model will not and should not fit stochastic noise in the data. For these reasons, the asymptote of ‘% *y* explained’ versus number of PCs sets an upper limit to the r^2^ value that can be expected from the PLSR model. In our example, Figure 4.2 established that >70% of the variance in y could be explained by the first 5 PLS components, and the resulting model built using these PCs 1-5 achieved an r^2^ value of 0.75 (Figure 5C).

#### Overfitting

Successive PCs capture progressively less and less of the variance in *y*, and thus represent decreasingly influential features from X in determining the value of *y*. At some point, the incremental amount of relevant information contained in a high-numbered PC will be outweighed by the stochastic noise it brings int the model, and the performance of the model will begin to degrade as it strives to fit this noise. The phenomenon is known as ‘overfitting’. The number of PCs at which this balance between relevant information and noise cannot be discerned by looking at ‘% *y* explained’, because this measure does not distinguish whether the variations in *y* that are being explained, or the variations in X being used by the PC to explain them, are meaningful or stochastic. The best way to avoid overfitting is to directly assess the predictive power of the model. In PLSR, this assessment is done in two different steps in the protocol. During the model building itself, cycles of cross-validation are performed to determine how accurately a model built using a certain number of PCs can predict the *y* values for elements that are left out of the calculation. The number of PCs that give the model having the lowest prediction error in cross-validation is considered the optimal number of PCs to use to build the final model. The application of the model to predict the y values for a separate Test Set of compounds that were entirely withheld from the model building process provides a second and more stringent test of the predictive power of the model. Provided that the elements in the Test Set fall within the domain of applicability defined by (i.e. are sufficiently similar to) the Training Set, any overfitting will be revealed as a q^2^ value for the Test Set that is substantially lower than the r^2^ calculated for the Training Set.

In the event that excessive overfitting is found, there are several possible ways to mitigate it. First, it is important to check that poor q2 values does not reflect the presence in the Test Set of compounds that fall outside the domain of applicability of the model, as was the case in the example shown in the protocol. If there are concerns on this score, they can be addressed by checking the Test Set for the presence of property outliers, for example by clustering as we showed in Basic Protocol 1, and reconstituting the Test Set in a way that excludes such compounds. In general, if the Test Set is small, such that the overall results could depend strongly on the behavior of a few of its individual elements, it can be helpful to select a few different Test Sets from the original data pool and make sure the q^2^ value you associate with the model is representative of the model’s typical performance. If this is not the problem, the it is necessary to improve the quality of the data in X and/or *y*, or to find alternative or additional properties to include in X that better explain how *y* varies.

#### Evaluating the predictive accuracy of PLSDA models

As described above, the predictive accuracy of a PLSR model is best reflected in its q^2^ value; that is, Pearson Correlation Coefficient of the predicted versus the actual outcome values. The accuracy of a model is, however, inextricably linked to its domain of application. In general, a model will make the most accurate predictions when applied to elements that closely resemble (i.e. fall within the property ranges that are best sampled by) the elements of the Training Set with which the model is built. To the extent that the compounds in the Test Set are representative of those in the Training Set, it is the model’s accuracy for this type of interpolative application that is estimated by q^2^. Applying a PLSR model to predict the outcome behavior of an element that is very different in its properties from those in the Training Set is unlikely to give an accurate result, and the q^2^ value is irrelevant to such extrapolative applications. When developing a model, it is therefore crucial to ensure that, to the extent possible, the Training Set densely samples the property space for the types of elements it will subsequently be used to predict. To complicate matters further, it is impossible to know which deviations from the properties of the Training Set are likely to have a large or small effect on prediction accuracy and so, without doing additional tests with elements of known outcome, it can be difficult to judge whether a given application lies close to or far outside the domain of applicability.

A dilemma for the investigator is that a broad domain of applicability can be achieved only by using a diverse Training Set, and yet the more diverse the Training Set the more complex are the ways in which the input properties might influence the outcome behavior. Thus, the models with the highest q2 values will often be those that were trained on a very homogeneous Training Set and will have a very narrow domain of applicability. It is much more difficult to develop a model with a broad enough domain of applicability to make accurate predictions for elements with very different properties.

In the example used in Basic Protocol 4, the results show that three compounds, ID755, ID757 and ID790, were predicted to be orders of magnitude more permeable than was found experimentally, clearly indicating that these three compounds differ from the bulk of the Training Set compounds in ways that are sufficient to place them well outside the domain of applicability for the current model. It is noteworthy that these three compounds were identified as outliers both by clustering (Basic Protocol 1a) and by PCA (Basic Protocol 2). Nonetheless, it was not predictable that these particular compounds would necessarily cause the model to fail; other outlier compounds, such as ID2385, ID808, and ID1881, were more accurately predicted, presumably because the ways in which they differ from the compounds that typify the Training Set are less consequential for PAMPA permeability. We note that the three compounds that were not well predicted by the PLSR model were correctly categorized by PLSDA in Basic Protocol 3. However, this result is likely related to the fact that the three compounds were predicted by PLSR to be much more permeable than is the case in reality, an overestimate that would not change their categorization as ‘Permeable’ in PLSDA. Aside from these three outliers, the PLSR model performed quite well with respect to the Test Set, predicting the permeabilities of the remaining 37 compounds with an accuracy of q^2^ = 0.98.

As always in science, in reality the investigator must work with whatever data can practically be obtained. It is therefore crucial to give careful thought to what properties are likely to influence the outcome of interest (i.e. *y*), what range of the associated property space is effectively sampled by the Training Set, whether new cases to which the model is applied are likely to fall in or out of the model’s domain of applicability, and how this risk might be assessed.

## VIII. COMMENTARY

The main messages of this article are that ML comprises a powerful set of tools for helping experimental molecular scientists to organize and interpret data, formulate and test hypothesis, and plan and interpret experiments, and that many ML techniques can be learned and rigorously applied without requiring a background in statistics or computer science. Indeed, in our experience the process of formulating an ML-based approach and validating its results are analogous to the way we think about other experimental methods. In both spheres, it is important to have a firm conceptual grasp how the method works and what it does, to know errors to avoid, and to understand how to validate the results through appropriate controls and cross-validation with other methods and information. Most importantly, with ML as with any experimental method, a good scientist will not simply believe the output that the method delivers, but instead will be mindful of precisely what hypothesis is being tested by a given analysis, will critically evaluate possible sources of error and bias, and will consider how the results can be further tested and confirmed through other methods.

### When to z-score and when not to z-score?

A question that arose in several of the methods discussed is when it is appropriate to z-score or otherwise scale the input data. There is no universal answer to this question. Generally speaking, how to scale and normalize the data is more of an issue for unsupervised methods such as clustering and PCA than it is for the supervised methods. The reason is that the supervised methods evaluate the input data in terms of how strongly it covaries with a measured outcome. Consequently, properties that correlate strongly with the outcome will be rated as important in a manner that is insensitive to how the property is scaled. For the unsupervised methods, the higher the numerical range a given property has, the more likely it is to contribute strongly to separating the elements in *n*-dimensional property space, and so the numerical scaling matters.

Whether and how to compensate for this bias depends on the goals of the analysis. In making this decision, some guidelines that we have found useful are as follows: (i) Any result that depends strongly on the arbitrary choice of what units to use to express a given quantity (e.g. whether a length is expressed in mm versus km) is clearly suspect, and if observed indicates that some scaling is required. (ii) To z-score the input data is equivalent to assuming that the degree of variance present among the elements on the data set in any one descriptor is equally important to that in any other descriptor for defining the distribution of the elements in property space. To z-score before clustering or PCA, therefore, is to define outliers in the relative sense of how far from the main group are they in terms of standard deviations from the mean of each property, rather than by any absolute distance. Consequently, if one or more properties happen to vary across the elements in the set over only a small range, z-scoring the data will magnify these differences to the same importance as potentially more consequential property differences. (iii) As alternatives to z-scoring, other kinds of systematic weighting can be applied to the different properties, such as weighting by the reciprocal of the variance as illustrated in Basic Protocol 2. (iv) For purposes of molecular and cellular research, a good approach can be to consider from a scientific perspective what range of values for each property would have roughly comparable significance to the likely behavior of the elements in the set with respect to the question or phenomenon of interest for the study. It is perfectly valid to individually scale properties in this manner. Such judgments are often somewhat subjective but, for the purpose of eliminating gross unwanted biases, it is typically not important that the scaling be particularly precise.

Key lessons about the application of the discussed ML methods can be summarized as follows:

- For the ‘explainable’ ML methods described herein, each analysis is based on the hypothesis that the internal relationships or external behavior of interest for the elements in the data set can be explained in terms of the properties in the input data matrix. The less this is true, the less meaningful the results of the analysis will be. In short, these ML methods will not tell you anything that isn’t contained in the data you use as inputs; their value is their ability to simplify the data to extract meaningful patterns and relationships that would be impossible or very difficult detect in the raw data.
- The results of the analysis depend not only on the choice of which properties to include in the analysis, but also on the set of elements in the data set. An identical analysis can give very different results if the same elements are analyzed using a different set of properties, or if the same properties are analyzed across a different set of elements. For a meaningful analysis it is therefore important to think deeply about how the set of data elements and properties you plan to use relate to the precise question you are trying to ask. It is dangerous to apply these methods blindly. Instead, it is necessary to design the study carefully, being alert to the assumptions, biases, and limitations inherent in any data set.
- Having highlighted the need for thoughtful design of ML analyses, we wish to emphasize that the judgment required is largely scientific, rather than computational or statistical. Indeed, the application of these methods to scientific problems by experts in computer science or statistics who lack a firm and nuanced grasp of the scientific principles at issue is itself a problem. It is this scientific judgment that is required to formulate a specific question or hypothesis, and to properly evaluate what data elements and properties are best to use, and how the properties should be scaled to avoid unwanted bias in the results. The ML calculations themselves are straightforward, especially as implemented in sophisticated tools such as MATLAB or R. Any thoughtful experimental scientist can learn to apply these methods rigorously and meaningfully, thereby adding powerful new capabilities to their research toolkit.

### Time Considerations

Each of the above protocols was tested by a set of four students who had no prior experience with ML. The protocols took the novice students, on average, between 15 minutes to 1 hour to complete each protocol.

Computational processing time is negligible with moderately sized data sets on a personal desktop or laptop computer. We have not ourselves worked with large datasets (one hundred thousand or more cells) and therefore do not know how much time MATLAB requires to perform these protocols in such cases. For very large datasets, it may be necessary to run the protocols using a computing cluster.

## Supporting information

Data for Basic Protocols 1 and 2, Clustering and PCA

Data for Basic Protocol 3, PLSDA

Data for Basic Protocol 4, PLSR

Description of how data were prepared

Data prepared as described in Data_prepared_SI

## AUTHOR CONTRIBUTIONS

**AW, LVB:** Drafted initial protocols for PCA, PLSR, and clustering. **AB:** Drafted initial protocol for PLSDA. **MJKV** and **KAH:** Formal analysis, investigation, writing original draft, figure/table generation as related to Basic Protocols 1, 2 and 4; review of writing and editing. **AB:** Formal analysis, investigation, writing original draft, figure/table generation as related to Protocol 3; review of writing and editing. **DLP:** supervision, funding acquisition, reviewed and edited final draft. **AW:** Conceptualization, funding acquisition, writing original draft, supervision, writing review, editing, and writing final draft.

## DATA AVAILABILITY STATEMENT

No new data were produced during this study.

## ACKNOWLEDGMENTS

AW thanks Professor Douglas Lauffenburger and his research group for hosting him during his 2017 sabbatical and helping him begin to learn about machine learning. The authors thank the following individuals for testing the protocols to determine the time it takes for a novice in ML to complete them: Christina Alcaro, Katrina Nguyen, Haley Curtis, and Hannah Braverman. We also thank Madeline Bodor, Martin Dokholyan, and Amanda Newell for helping to verify all correct toolboxes needed for download through MATLAB. This work was supported by funding from the Department of Chemistry at Boston University. AW would like to acknowledge funding from Genentech Inc. that supported LVB in her development of some of the protocols described here. The work of AB was funded, in part, by grant ##### (to DLP).

