## Supplementary material for "Getting Started with Machine Learning for Experimental Biochemists and Other Molecular Scientists": Description of how data were prepared

**Supplementary for data preparation (Data_prepared.xlsx)**

1. Collect the input elements (compounds), features (descriptors), numerical X data (computed values) and numerical y data (only necessary for Basic Protocols 3 and 4 only, PAMPA values) in a single tab titled ‘data_prepared’ of a .xlsx file with no gaps or breaks between cells. File should be formatted with features listed at the top of each column in row 1 and input data element names at the start of each row in column A.

*To prepare our dataset: Download peptide data files for ‘Monomer Length’ 8 and 9 from http://cycpeptmpdb.com/download/ and save each file as a .xlsx. We have provided both datasets in two different tabs in the supplemental file RawData_CycPeptMPDB_DataSet_8and9.xlsx because we cannot guarantee these datasets will be available for download in the future.*

*There are 111 peptides in each file. Merge the peptides into one excel sheet by copying rows 2-112 of the ‘CycPeptMPDB_Peptide_Length_9’ tab to rows 113-223 of ‘CycPeptMPDB_Peptide_Length_8’ tab. Rename tab ‘data_prepared’. Sort dataset from highest to lowest PAMPA value.*

2. Start with a clear workspace in MATLAB. From the ‘data_prepared’ tab of the .xlsx file, select all consecutive cells starting from B2. Once cells with data have been highlighted, select ‘Numeric Matrix’ under ‘Output Type’, then press ‘Import Data’. Rename your file in the main MATLAB window directly after you import your data. In our example we use **X** to describe our numeric matrix.

*Data must be named before more selections are imported into MATLAB. If not, the numeric array will continually override the previous data file. We imported cells B2:IJ223 to get a 222 x 243 numeric matrix. The* ***X*** *import at this step includes the column of y data.*

3. From the ‘data_prepared’ tab of the .xlsx file, select all consecutive cells in row 1. Once cells with data have been highlighted, select ‘String Array’ under ‘Output Type’, then press ‘Import Data’. Rename your file in the main MATLAB window directly after you import your data. In our example we use **Descriptors** to describe our string array.

*We imported cells A2:IJ1 to get a 1 x 243 string array.*

4. From the ‘data_prepared’ tab of the .xlsx file, select consecutive cells in column A starting with row 2. Once cells with data have been highlighted, select ‘String Array’ under ‘Output Type’, then press ‘Import Data’. Rename your file in the main MATLAB window directly after you import your data. In our example we use **Compounds** to describe our string array.

*We imported cells A2:A223 to get a 222 x 1 string array.*

5. Remove any unnecessary information from your matrix to prepare **X** for analysis.

**ToDelete=[i:j, k, m]**

**X(:,ToDelete)=[]**

**Descriptors(:,ToDelete)=[]**

*We use* ***i*** *= 1,* ***j*** *= 18,* ***k*** *= 242 and* ***m*** *= 243. In our scenario, we are removing columns B through S, which contain irrelevant information pertaining to the sequence and naming of the residues in the macrocycles. We also remove columns II through IJ, which contain the PC1 and PC2 as these were non physicochemical descriptors from an analysis performed in the paper these were extracted from. After this step,* ***X*** *is a 222 x 223 numeric matrix and* ***Descriptors*** *is a 1 x 223 string array. This procedure may require adjustments if your input includes additional information that is not relevant to the analysis.*

6. Sort **X** and **Compounds** in ascending order by the y value.

**[X_sort, sortidx] = sortrows(X, 1,'ascend')**

**Compounds_sort = Compounds(sortidx)**

*This step is not essential, but sorting the dataset allows for simpler comparison between actual and predicted y values for the training set. Be sure to sort if following along with our example.*

7. Remove rows in **X** where y values are not available. Repeat to remove the same rows from **Compounds**.

**X_1=X_sort(~isnan(X_sort(:,1)),:)**

**Compounds_1=Compounds_sort(~isnan(X_sort(:,1)),:)**

*After this step, X_1 is a 201 x 223 numeric matrix and Compounds_1 is a 201 x 1 string array.*

8. Remove columns in **X** that show no variability, indicated by all values being zero. Repeat to remove the same columns from **Descriptors**.

**X_2=X_1(:,~all(X_1==0,1))**

**Descriptors_2=Descriptors(~all(X_1==0,1))**

*After this step, X_2 is a 201 x 154 numeric matrix and Descriptors_2 is a 1 x 154 string array.*

9. Remove columns in **X** that contain any NaN (Not a Number) values. Repeat to remove the same columns from **Descriptors**.

**X_3=X_2(:,~any(isnan(X_2),1))**

**Descriptors_3=Descriptors_2(~any(isnan(X_2),1))**

*After this step, X_3 is a 201 x 140 numeric matrix and Descriptors_3 is a 1 x 140 string array.*

10. Open a new .xlsx file. Copy 'Descriptors_3' from MATLAB and paste it as the first row in the .xlsx file starting in cell B2. Then, copy ‘Compunds_1’ and paste it in column A of the .xlsx file beginning in cell A2. Finally, copy ‘X_3' and paste it beginning in cell B2 of the .xlsx file. Save this file as Data_prepared.xlsx.

*In our example,* ***Descriptors_3*** *will have dimensions of 1 x 140,* ***Compounds_1*** *will have dimensions of 201 x 1 and* ***X_3*** *will have dimensions of 201 x 140.*

**Supplementary for PLSDA 80-20 generation (Data_PLSDA.xlsx)**

1. Open Data_prepared.xlsx. file and save as new .xlsx Data_PLSDA.xlsx.
2. Make a copy of tab ‘data_prepared’ tab and rename the copy as ‘data_prepared_plsda’.
3. After the 'CycPeptMPDB_ID' column (column A), add another column named ‘Class’, which will be used to categorize compounds into classes.
4. To the first 50 compounds (rows 2-51), add '1' to the column ‘Class’ to indicate the class of compounds with high permeability. To the bottom 50 compounds (rows 153-202), assign '2' to indicate the class of compounds with low permeability. Unassigned compounds (rows 52 to 152) can be deleted.

*For binary categorization in PLSDA use numbers 1 and 2 for classes. 0 is left as for the category that could not be assigned.*

5. Start with a clear workspace in MATLAB. From the ‘data_prepared_plsda’ tab of Data_PLSDA.xlsx, import consecutive cells of numerical *m* x *n* input data excluding column headers, sample names, and y data (‘PAMPA’ or ‘Class’) for rows corresponding to class 1 from step 4. Once cells with data have been highlighted, select ‘Numeric Matrix’ under ‘Output Type’. Once this is selected, press ‘Import Data’. Rename your file in the main MATLAB window directly after you import your data. In our example we use **X** to describe our *50* x 139 numeric matrix.

*Data must be named before more selections are imported into MATLAB. If not, the numeric matrix will continually override the previous imported workspace variable.*

11. Once **X** is selected, repeat the selection process with your *m* x 1 numeric output data by selecting the ‘Class’ column excluding the header name for class 1 compounds only. Select ‘Numeric Matrix’ under ‘Output Type’ then select ‘Import Data’. Name the output data matrix as needed. To use the same 80/20 generation script for both PLSDA and PLSR, we rename our 50 x 1 output data **y**.

*Output data will be named* **Class_test** *or* **Class_train** *in ‘Basic Protocol 3: PLSDA’.*

12. Select your *m* x 1 elements for class 1, excluding the column header, and import as ‘String Array’. Rename as needed. We rename our 50 x 1 string array of compound ID’s **Compounds**.
13. Repeat the selection process with your headers for each column of **X**, which are termed features, excluding the headers for ‘CycPeptMPDB_ID’, ‘PAMPA’ and ‘Class’. Change the input type of matrix to ‘String Array’ and then ‘Import Data’. Name the features as needed. For our example we rename our 1 x 139 string array **Descriptors**.
14. Z-score **X** and run k-medoids clustering on the new matrix **Data** to generate a 20% test set to omit from an 80% training set.

**[Z,mu,sigma] = zscore(X)**

**Z_train(isnan(Z_train)) = 0**

**rng(0)**

**[idx,M,sumd,D,midx,info]=kmedoids(Z,N)**

*Specify the number of compounds* **N** *that you need to select. The generated output* ***midx*** *will contain* **N** *rows selected for the testing set for class 1 or class 2. The numbers in* ***midx*** *correspond to the row number from the input matrix* **X***. In our case, we will use* ***N*** *= 10 which is 20% of our 50 compounds in each class.*

15. To manually generate the test and training set, create a copy of the ‘dataset_prepared_PLSDA’ tab and rename it as ‘PLSDA_train’. Remove rows with corresponding row numbers from **midx** output from the ‘PLSDA_train’ tab and paste to a new tab with the same headers named ‘PLSDA_test’.

*The row numbers from* ***midx*** *refer to* ***X****, which contains only numerical values for* ***Descriptors*** *without headers. If you are curating the testing set manually in a .xlsx file, make sure not to include headers in determining the rows that need to be removed for ‘X_test’.*

*Instead, to automatically generate the test and training sets, use the* generate_test_train.mlx *provided in the supplemental material for Step 15 for each class. The script will output* ***Descriptors*** *and test and train arrays for,* ***X****,* ***y*** *and* ***Compounds*** *for the respective class. Copy these variables into new ‘PLSDA_train’ and ‘PLSDA_test’ tabs as shown in Figure 3.1 in the main text.*

*If a pop up appears after selecting 'Run' under the 'EDITOR' tab, select 'Change folder' to continue.*

16. Clear the workspace and repeat steps 5-15 of this protocol for compounds from class 2 (rows 51-101 in ‘dataset_prepared_plsda’). If following the step for manual generation, remove and paste the 20% to the same ‘PLSDA_test’ tab below the rows selected from class 1. If using automatic generation with the script, copy and paste the *test and train arrays for,* ***X****,* ***Class*** *and* ***Compounds*** *for class 2 below class 1 in the* ‘PLSDA_train’ and ‘PLSDA_test’ tabs.

*As a result, you will have 80 rows of compounds (excluding headers) in ‘PLSDA_train’ and 20 rows of compounds in ‘PLSDA_test’. The headers will be the same in both .xlsx files.*

**Supplementary for PLSR 80-20 generation (Data_PLSR.xlsx)**

1. Open Data_prepared.xlsx. file and save as new .xlsx Data_PLSR.xlsx.
2. Start with a clear workspace. Import your numeric m x n input data excluding any column headers or sample names from the ‘dataset_prepared’ tab. Once cells with data have been highlighted, select 'Numeric Matrix' under ‘Output Type’. Once this is selected press ‘Import Data’. Ensure that you rename your file in the main MATLAB window directly after you import your data. In our example we use **X** to describe our 201 x 139 numeric matrix.

*Do not select your cells that you will be using as your* ***y*** *values yet. The selection should start in cell C2.*

*Data must be named before more selections are imported into MATLAB. If not, the numeric matrix will continually override the previous imported workspace variable.*

3. Once your data is selected repeat the selection process with your m x 1 numeric output data by selecting this column excluding any header names. Select ‘Numeric Matrix’ under ‘Output Type’ then select ‘Import Data’. Name the output data matrix as needed. For our example we label our *201* x 1 matrix as **y**.
4. Select your m x 1 data elements and import as ‘String Array’. Rename as needed. We label our 201 x *1* string array of compound ID’s as **Compounds**.
5. Repeat the selection process with your headers for each column of **X** which are termed features. Change the input type of matrix to ‘String Array’ and then 'Import Data'. Name the features as needed. For our example we label our 1 x *139* string array of descriptors as **Descriptors**.
6. Run k-medoids clustering on the z-scored **X** matrix to generate a 20% test set to omit from an 80% training set.

**[Data,mu,sigma] = zscore(X)**

**Z_train(isnan(Z_train)) = 0**

**rng(0)**

**[idx,M,sumd,D,midx,info]=kmedoids(Data,N)**

*The k-medoids command generates an N x 1 array of numbers which correspond to rows for* ***X****. N is the number of data elements you wish to have in your test set. Because we have 201 data elements, 20% of the set would be 40 so we used* ***N*** *= 40.*

7. To automatically generate the test and training sets, use the command line to run the ‘generate_test_train.mlx’ script provided in the supplemental material. The script will output test and train matrices for **X**, **y** and **Compounds**.

*If a pop up appears after selecting 'Run' under the 'EDITOR' tab, select 'Change folder' to continue.*

8. Remove columns from ‘X_train’, ‘X_test’ and **Descriptors** which are all 0’s or 1’s in the training set.

**X_train_2 = X_train(:, ~all(X_train == 0, 1))**

**X_test_2 = X_test(:, ~all(X_train == 0, 1))**

**Descriptors_2 = Descriptors( ~all(X_train == 0, 1))**

9. Copy variables ‘Compounds_train’, ‘X_train_2’, ‘y_train’ and ‘Descriptors_2’ into new ‘PLSR_train’ tab and variables ‘Compounds_test’, ‘X_test_2’, ‘y_test’ and ‘Descriptors_2’ into new ‘PLSR_test’ tab as shown in Figure 4.1 in the main text.

*As a result, you will have 161 rows of compounds (excluding headers) in ‘PLSDA_train’ and 40 rows of compounds in ‘PLSDA_test’. The headers will be the same in both .xlsx files.*

*The header name for cell A1 in both tabs is ‘CycPeptMPDB_ID’ and the header name for cell B1 in both tabs is ‘PAMPA’.*
